# Mobile version of the Battery for the Assessment of Auditory Sensorimotor and Timing Abilities (BAASTA): Implementation and adult norms

**DOI:** 10.1101/2023.07.21.550031

**Authors:** Simone Dalla Bella, Nicholas E.V. Foster, Hugo Laflamme, Agnès Zagala, Kadi Melissa, Naeem Komeilipoor, Mélody Blais, Simon Rigoulot, Sonja A. Kotz

## Abstract

Timing and rhythm abilities are complex and multidimensional skills that are highly widespread in the general population. This complexity can be partly captured by the Battery for the Assessment of Auditory Sensorimotor and Timing Abilities (BAASTA). The battery, consisting of 4 perceptual and 5 sensorimotor tests (finger tapping), has been used in healthy adults and in clinical populations (e.g., Parkinson’s disease, ADHD, developmental dyslexia, stuttering), and shows sensitivity to individual differences and impairment. However, major limitations for the generalized use of this tool are the lack of reliable and standardized norms and of a version of the battery that can be used outside the lab. To circumvent these caveats we put forward a new version of BAASTA on a tablet device capable of ensuring lab-equivalent measurements of timing and rhythm abilities. We present normative data obtained with this version of BAASTA from over 100 healthy adults between the ages of 18 and 87 years in a test-retest protocol. Moreover, we propose a new composite score to summarize beat-based rhythm capacities, the Beat Tracking Index (BTI), with close to excellent test-retest reliability. BTI derives from two BAASTA tests (beat alignment, paced tapping), and offers a swift and practical way of measuring rhythmic abilities when research imposes strong time constraints. This mobile BAASTA implementation is more inclusive and far-reaching, while opening new possibilities for reliable remote testing of rhythmic abilities by leveraging accessible and cost-efficient technologies.

## Introduction

Accurately perceiving the timing of auditory events is crucial for human adaptation to environmental demands. It enables us to cross a busy street safely, coordinate movements while dancing, and learn new skills such as speaking or playing a musical instrument. By extracting temporal regularities from external events, we can swiftly adjust and effectively coordinate our actions. The timing capacities that underlie auditory-motor skill learning play a critical role across the lifespan in both healthy and atypical populations (Dalla Bella et al., 2018; Rodriguez-Fornells et al., 2012; Trainor & Cirelli, 2015; Zatorre et al., 2007). Timing abilities are apparent in fine-grained beat perception in music, and the ability to move in synchrony with its regular rhythms. Musical features such as its regular temporal structure (rhythmic complexity, syncopation), but also its pitch structure (harmonic complexity) (Matthews et al., 2019; Witek et al., 2014) are particularly conducive to movement. This connection may be rooted in brain areas involved in motor control and audition (e.g., basal ganglia, motor, and pre-motor as well as temporal cortical areas) that are activated when we listen to rhythmic sequences without actually moving (Chen et al., 2008; Grahn & Brett, 2007; Janata et al., 2012; Kotz et al., 2018; Zatorre et al., 2007).

The majority of individuals are well equipped to process event timing, extract temporal regularities in the environment, and synchronize movement to an external beat (e.g., Damm et al., 2020; Grondin, 2010; Patel & Iversen, 2014; Sowiński & Dalla Bella, 2013; Tranchant et al., 2016). Among healthy individuals, however, there is considerable variability that manifests along a spectrum of huge individual differences in beat perception and synchronization (Fiveash et al., 2022; Sowiński & Dalla Bella, 2013; Tranchant et al., 2016). While many can extract the beat underlying a simple and complex rhythmic auditory sequence, some – “beat-deaf” or “poor synchronizers” – struggle with tests of beat perception, and/or when they are asked to move to the beat of an auditory sequence. These rhythm difficulties can take different forms and might be characterized by both poor rhythm perception and production (Palmer et al., 2014; Phillips-Silver et al., 2011), or show selective impairment of production (Sowiński & Dalla Bella, 2013) and/or perception (Bégel et al., 2017). Finally, poor timing and rhythmic abilities are found in clinical populations, including neurodegenerative and neurodevelopmental disorders (Allman & Meck, 2012; Lense et al., 2021), Parkinson’s disease (Benoit et al., 2014; Grahn & Brett, 2009; Merchant et al., 2008), ADHD (Noreika et al., 2013; Puyjarinet et al., 2017), and speech and language impairments (Bégel et al., 2022; Corriveau & Goswami, 2009; Falk et al., 2015; Ladányi et al., 2020).

Disrupted timing and rhythm abilities, when appearing early in life, may contribute to atypical developmental symptoms (Lense et al., 2021); they are also related to impaired performance in the elderly (e.g., motor disorders; Dalla Bella, 2020; Dalla Bella et al., 2017). Thus, a better characterization of timing and rhythm disorders in clinical populations is likely to play an important role in identifying risk factors in development (Ladányi et al., 2020) and in personalized intervention strategies (Dalla Bella et al., 2018; Dotov et al., 2019).

Capturing the complexity of timing and rhythmic abilities is a challenging task. These abilities are typically multidimensional, and are underpinned by a complex cognitive architecture involving several dimensions (e.g., beat-based, memory-based processes; Bonacina et al., 2019; Bouwer et al., 2020; Fiveash et al., 2022; Kasdan et al., 2022; Tierney & Kraus, 2015). This complexity is difficult to capture through isolated tasks and instead requires multiple assessments, as provided by comprehensive test batteries. Multiple tests afford precise measurement of timing and rhythm abilities, and the detection of individual profiles (Dalla Bella et al., 2024), allowing the isolation of the locus of impairment in individuals with rhythm disorders. Existing batteries for testing timing and rhythm abilities include both perceptual and sensorimotor tasks. The perceptual and performance dimensions of timing and rhythm abilities are not always correlated (Dalla Bella, Farrugia, et al., 2017; Fujii & Schlaug, 2013), suggesting that both abilities should tested to obtain a comprehensive profile of these abilities. Examples of tasks are perceptual judgments (e.g., deciding whether a metronome is aligned or not to a musical beat; Beat Alignment Test; Iversen & Patel, 2008), or finger tapping tests (Repp, 2005; Repp & Su, 2013). To systematically probe timing and rhythm abilities, batteries of perception and production tests have been devised such as the Battery for the Assessment of Auditory Sensorimotor and Timing Abilities (BAASTA, Dalla Bella, Farrugia, et al., 2017), and the Harvard Beat Alignment Test (H-BAT, Fujii & Schlaug, 2013). Here we will focus on BAASTA, which is an exhaustive battery including four perceptual tests and five sensorimotor tests (via finger tapping). One of the advantages of BAASTA is that it covers various timing and rhythm abilities, providing dedicated behavioral tests widely used in a wealth of studies (e.g., paced and unpaced finger tapping, detection of anisochronies, and beat alignment). The battery is sensitive to individual differences in timing and rhythm abilities. It can distinguish between individuals with and without musical training or informal music experience (Dalla Bella et al., 2024), as well as impaired rhythm perception and performance in healthy and patient populations (e.g., Bégel et al., 2017, 2022; Benoit et al., 2014; Dalla Bella, Farrugia, et al., 2017; Falk et al., 2015; Fiveash et al., 2022; Hidalgo et al., 2021; Oschkinat et al., 2022; Puyjarinet et al., 2017, 2022; Verga et al., 2021; Zagala et al., 2021). Hence, the battery can serve as a screening tool for rhythm disorders, and pave the way to individualized rhythm-based interventions (Dalla Bella et al., 2018).

Despite these advantages, BAASTA so far presents a few shortcomings. The battery is currently implemented for lab usage, thanks to tapping pads or sensors ensuring high temporal precision (≤ 1ms); this makes it quite unsuitable for use outside the lab, remotely, in a school, or for clinical use at a patient’s bedside. Interestingly, there have been developments of rhythmic tasks on mobile devices such as tablets and smartphones (Puyjarinet et al., 2017; Zanto et al., 2019), which are appealing and inclusive solutions for testing cognitive functions (Koo & Vizer, 2019). The problem with using off-the-shelf devices such as tablets or smartphones, however, is that they are not conceived as measurement tools in the first place. Timing inaccuracies resulting from unwanted delays (e.g., in the audio output, or between touch detection and recording) and imprecision due to the low sampling rate of the device touchscreen (between 60 and 240 Hz; Kousa, 2017) are common. A second shortcoming of BAASTA, and more generally with tests measuring timing and rhythmic abilities, is the lack of normative data across the lifespan. This limits our capacity to define cut-off scores for impaired performance and necessitates further data collection from a matched control group in separate studies. Finally, running participants through the full set of BAASTA tests is particularly time consuming (i.e., approximately 2 hours). Even though it may be justified in some cases to run all tests to obtain a thorough assessment of rhythmic abilities, this might be impractical or even unfeasible in clinical or educational setting.

Therefore, the objective of the current study is two-fold. Given the aforementioned limitations of mobile devices to record rhythmic performance, we first present a new solution, implementing BAASTA on a tablet device that ensures lab-comparable high accuracy and precision (Zagala et al., 2021). A prototype of this BAASTA tablet version was already used in several prior studies in healthy adults, children, and clinical populations (Bégel et al., 2018, 2022; Dauvergne et al., 2018; Puyjarinet et al., 2017, 2019). A further goal is i) to provide norms for BAASTA-tablet collected data from a group of over 100 adults across the lifespan (from 18 to over 80 years), ii) obtain test-retest reliability metrics for the battery, and iii) propose a composite score (i.e., the Beat Tracking Index) to summarize perceptual and sensorimotor rhythmic abilities, with a focus on beat-based abilities, within a minimal amount of time. This composite score is derived from two BAASTA tests: the Beat Alignment Test and paced tapping to music. These tasks can effectively represent beat-based rhythmic abilities, encompassing both beat perception and production (see Dalla Bella et al., 2024; Fiveash et al., 2022). Their performance is usually highly correlated, suggesting that they may tap onto similar beat-based mechanisms (Puyjarinet et al., 2017; Tranchant et al., 2021). Notably, poor performance in either or both of these tests is typically observed in the general population and clinical groups (for reviews, see Dalla Bella et al., 2024; Fiveash et al., 2022). Hence, a composite score, including both tests, is likely to provide a sensitive measure for capturing poor rhythmic performance in a parsimonious way. With this in mind, we introduced the Beat Tracking Index in a previous study where we tested rhythmic abilities in children and adults with ADHD as well as in neurotypical individuals (Puyjarinet et al., 2017). This composite score was useful in characterizing beat-based rhythmic abilities and in demonstrating a link between beat tracking and executive functions (inhibition control and flexibility) in ADHD. In summary, the Beat Tracking Index is expected to be a valuable tool for the inclusion in BAASTA norms. It has the potential to identify individual differences in beat-based rhythmic abilities in both neurotypical and clinical populations.

## Materials and methods

### Participants

A total of 116 adults, between 18 and 87 years of age and recruited in the Montreal area, started the experiment. Of these participants, 108 (74 female; 98 right-handed, 9 left-handed and 1 ambidextrous) completed the experiment. The other 8 participants (6.9% of the total) could not complete the experiment, or their data was discarded for technical reasons (intermittently recorded data), and consequently were not included in the analysis. We divided the final participants into four age groups: 18-21 years (*n* = 27; 18 female; *M* = 19.5 years; *SD* = 1.0), 22-29 years (*n* = 29; 19 female; *M* = 23.8 years; *SD* = 1.7), 30-54 years (*n* = 26; 16 female; *M* = 39.7 years; *SD* = 7.5), and 55-87 years (*n* = 26; 21 female; *M* = 67.9 years; *SD* = 8.5). The age range in each group was determined to ensure a comparable sample size in each group, based on the tested participants. A summary of participants grouped by age group, gender, and sample size at Test and Re-test is provided in Table 1. There was no difference in gender proportion across these age groups (*X*^2^(3) = 2.56, *p* = 0.46). We recruited participants who were fluent in English or French. We included only participants who were non-musicians (i.e., individuals who did not consider themselves as musicians, and with less than 5 years of formal musical training). Most participants (*n* = 72) indicated French as their first language; other first languages were Arabic (7), Spanish (7), English (6), Mandarin/Cantonese (4), Portuguese (3), Marathi (2), Bengali (1), Créole (1), Persian (1), Romanian (1), Russian (1), Telugu (1), and Turkish (1). Participants had a mean of 0.3 years of formal musical training (SD = 0.7 years; range: 0-4 years), and 1.5 years of total musical experience including both formal and informal musical activities (SD = 2.8 years; range: 0-19.5). Age groups did not differ in both measures of musical experience (*p* ≈ 1). Exclusion criteria for participating in the study were a history of alcohol or drug consumption, brain concussions, and reported visual or hearing deficits. All participants took part in a first testing session (*Test*) and 93 of them participated in a second session (*Re-test*) (64 female). The proportion of the participants in the Re-test across the age groups was comparable to that of the Test session. In the Re-test session 21 participants were in the 18-21 age group (15 female), 24 in the 22-29 group (15 female), 23 in the 30-54 group (14 female), and 25 in the 55-87 group (20 female). The study was approved by the ethics committee (certificate n.: CERAS-2018-19-096D) of the University of Montreal. Informed consent was obtained from all participants.

**Table 1.** Participant Demographics at Test (n = 108) and Re-test (n = 93)

| Age group | Test |  | Re-test |  |
| --- | --- | --- | --- | --- |
|  | N (female) | Mean age (SD) | N (female) | Mean age (SD) |
| 18-21 years | 27 (18) | 19.5 (1.0) | 21 (15) | 19.5 (1.1) |
| 22-29 years | 29 (19) | 23.8 (1.7) | 24 (15) | 23.5 (1.4) |
| 30-54 years | 26 (16) | 39.7 (7.5) | 23 (14) | 40.1 (7.7) |
| 55-87 years | 26 (21) | 67.9 (8.5) | 25 (20) | 67.6 (8.5) |

### Procedure and tests

Testing of participants involved two separate sessions (Test, Re-Test) conducted in sound-proof rooms at the BRAMS laboratory at the University of Montreal. In each session, we tested participants’ sensorimotor and timing abilities with the Battery for the Assessment of Auditory and Sensorimotor Timing Abilities (BAASTA; Dalla Bella et al., 2017). The tests and measures of perceptual and sensorimotor timing abilities adopted in this study are the same as in the original version of BAASTA. Note that in tests which required the calculation of a threshold (Duration discrimination, Anisochrony detection with tones, with music; see below), we used a staircase procedure (2 down / 1 up; Exp. 2, Dalla Bella et al., 2017) instead of a Maximum Likelihood Procedure (Exp. 1, Dalla Bella et al., 2017). The administration of the full BAASTA took approximately 2 to 2.5 hours in each session, and the sessions were separated by approximately 30 days. In a separate screening session, prior to Test and Re-Test, participants filled in a short questionnaire on basic demographic information and their musical experience.

#### Measures of perceptual and sensorimotor timing abilities

BAASTA includes 4 perceptual tasks and 5 production timing tasks. Perceptual tasks involved the discrimination of single durations (Duration discrimination), the detection of deviations from temporal regularity (beat isochrony) in tone and musical sequences (Anisochrony detection with tones, with music), and the detection of the alignment between the musical beat and a superimposed metronome sound (Beat Alignment Test). Sensorimotor tasks were finger tapping tests in the absence of stimulation (Unpaced tapping), synchronizing to the beat of tone and music sequences (Paced tapping with tones, with music), continuing tapping at the pace of a metronome (Synchronization-continuation), and adapting tapping to a tempo change (Adaptive tapping). The tasks were implemented via an app on a tablet device (see below for details), which recorded participants’ responses. Auditory stimuli were delivered via headphones (Beyer dynamic DT 770 PRO or DT 990 PRO). The sound pressure level was adapted for each participant at the beginning of the testing to achieve a comfortable level. The level was adjusted if needed in tasks across the battery, if explicitly requested by the participants. As BAASTA is formed by two test sets (perceptual and sensorimotor), the order of the sets was counter-balanced across participants; in the analyzed data set, 62 participants (57%) received perceptual tasks followed by sensorimotor tasks, while 46 (43%) received the opposite order. Task order within each set was fixed (Duration discrimination, Anisochrony detection with tones and music, BAT, for perceptual tasks; Unpaced tapping and Paced tapping to tones and music, for motor tasks). Each task was preceded by instructions and, except for the Unpaced tapping task, at least one practice trial. In addition, perceptual tasks included examples of stimuli (e.g., with or without a duration difference, anisochrony, or beat misalignment). Examples of stimuli presented in the different tasks are available at: https://www.biorxiv.org/content/10.1101/2023.07.21.550031 (in Supplementary material).

##### Perceptual tasks

In the *Duration discrimination* test we presented two tones (frequency = 1 kHz, envelope ramp of 10 ms) successively, one at a standard duration (600 ms) and the other at a comparison duration (between 600 and 1000 ms). The tones were separated by a silent interval of 600 ms. Participants judged whether the second tone lasted longer than the first. In the *Anisochrony detection with tones* task we tested the detection of a time shift in an isochronous tone sequence. We presented sequences of 5 piano tones (MIDI note = C6, tone frequency = 1047 Hz, duration = 150 ms; Inter-Onset Interval – IOI = 600 ms). Sequences were isochronous (i.e., with a constant IOI) or not (the 4^th^ tone was presented earlier than expected by up to 30% of the IOI). Participants judged whether the sequence was regular or not. In the *Anisochrony detection with music* task, we assessed the detection of a time shift in a musical sequence, namely an excerpt of two bars from Bach’s “Badinerie” orchestral suite for flute (BWV 1067) played with a piano timbre (inter-beat interval, IBI = 600 ms). As before, participants judged the regularity of the sequence. We tested beat perception with the *Beat Alignment Test* (BAT). We used 72 stimuli based on 4 regular musical sequences, including 20 beats each (beat = quarter note): two fragments from Bach’s “Badinerie”, and 2 from Rossini’s “William Tell Ouverture”. The stimuli were played with a piano timbre at three different tempi (with 450, 600, and 750-ms IBIs). These stimuli were fragments of longer excerpts from the same musical pieces used in the Paced tapping with music task (see below). A metronome (i.e., woodblock sound) was superimposed onto the music, and it was either aligned or non-aligned (out of phase, by +- 33% of the IBI, or with a different period, by +- 10% of the IBI) relative to the musical beat. Phase and period misalignments were equally likely in both directions (earlier/later; slower/faster). There were 18 stimuli (6 for each tempo) for each musical sequence: 4 stimuli with a metronome aligned to the beat, and 4 with a misaligned metronome (2 out of phase – 1 with a positive phase shift, 1 with a negative shift; 2 with a different period – 1 with a faster metronome, 1 with a slower metronome). Participants judged if the metronome was aligned or not with the musical beat. In all perceptual tasks except BAT, we used an adaptive two-alternative forced-choice staircase procedure (Leek, 2001; Levitt, 1971; Dalla Bella et al., 2017, for an implementation in BAASTA). The staircase protocol started with a large difference between standard and comparison stimuli (i.e., change in duration, IOI, or IBI). The difference changed adaptively during the trial, conditional to the participant’s response, and controlled by a 2 down / 1 up staircase procedure. Two consecutive positive responses to a difference were needed to reduce it by half in the following trial. A negative response led to a reverse in the change, and the difference was multiplied by a factor of 1.5. Every time the direction of a difference change reversed from up to down or from down to up, the value of the difference at which this occurred was recorded as a turnaround point. The trial ended after eight turnaround points and the threshold was calculated by averaging the last four. The obtained threshold corresponded to a probability of 70.7% for the 2 down / 1 up relative to the psychometric curve (Levitt, 1971). Notably, for each staircase sequence of stimuli there were 3 catch trials in which there was no change, which served to ensure that the participants paid attention during the task.

##### Sensorimotor tasks

In all sensorimotor tasks participants responded by tapping with the index finger of their dominant hand. To measure the participants’ preferred tapping rate and its variability in the *Unpaced tapping* task, we asked the participants to tap at their most natural rate for 60 seconds. In two additional unpaced conditions, we asked the participants to tap as fast or as slow as possible for 60 seconds. The unpaced tapping task at preferred tapping rate was repeated twice, at the very beginning and at the very end of the sensorimotor tasks. In the *Paced tapping with tones* task participants tapped to the sounds of a metronome (60 isochronously presented piano tones, MIDI note = E7, tone frequency = 2637 Hz, duration = 78 ms). There were three trials, each at one of 3 different tempi (600, 450, and 750-ms IOI). In the *Paced tapping with music* task participants tapped to the beat of two musical excerpts taken from Bach’s “Badinerie” and Rossini’s “William Tell Overture” (64 quarter notes; IBI = 600 ms). Participants repeated the paced tapping trials twice for each stimulus sequence (except 5 participants who did not perform the repetition due to a technical error). To test continuation tapping at the rate provided by a metronome, in the *Synchronization-continuation* task participants tapped with an isochronous sequence of 10 piano tones (MIDI note = E7, tone frequency = 2637 Hz, duration = 78 ms; at 600, 450, and 750 ms IOI), and continued tapping at the same rate, for a duration corresponding to 30 IOIs after the sequence stopped. Participants repeated the task twice at each tempo (except 5 participants). To assess the ability to adapt to a tempo change in a synchronization-continuation task, in the *Adaptive tapping* task participants tapped to an isochronous sequence (10 piano tones, MIDI note = C7, tone frequency = 2093 Hz, duration = 78 ms; at 600 ms IOI), at the end of which (last 4 tones) the tempo either increased, decreased (by 30 or 75 ms), or was kept constant (40% of the trials). Participants tapped to the tones in the sequence, and were instructed to adapt to the tempo change, and to continue tapping at the new tempo after the stimulus stopped. Moreover, at the end of the trial, participants judged whether they perceived a change in stimulus tempo (acceleration, deceleration) or not. In this task there were 10 experimental blocks of 6 trials each, presented in random order.

#### BAASTA tablet application

In previous studies, BAASTA was implemented as a computer version, making use of different software programs (Max MSP, and Matlab) and MIDI percussion pads (e.g., Benoit et al., 2014; Dalla Bella, Farrugia, et al., 2017). Here the battery took the form of an application on a tablet device (Samsung Galaxy Tab E running Android 7.1). The application supporting the tablet version of BAASTA will be available via the BeatHealth S.A. company (https://www.beathealth.tech/) on a Pay-Per-Use (PPU) basis. Participants’ responses in the perceptual tests were communicated verbally to the experimenter who entered them on the tablet device. Finger tapping performance was collected by having participants tap within a green rectangle (10.0 by 8.8 cm) on the tablet touchscreen (see Figure 1). As tablet devices are not precise measurement tools by design, we paid particular attention that the recording of unpaced and paced tapping timing complies with the highest lab standards. Timing inaccuracy in tapping tasks on tablet can arise from a delay in the audio output, the temporal uncertainty arising from the sampling rate of touch detection, and the processing delay between the touch detection and the recording of a tap (Zagala et al., 2021). These shortcomings are mostly due to the precision of the device touchscreen (sampling rate between 60 and 240 Hz) which is much coarser than lab measurements (1000 Hz or more). These issues are circumvented in the present BAASTA tablet implementation by relying on an audio recording of the sound the taps produce when they reach the touchscreen via the tablet integrated microphone. This procedure involves different phases, including the recording of the sound produced by the taps and of the contact time on the device touchscreen, and the alignment of the detected taps with the pacing stimulus to ensure the precise calculation of tap-to-beat synchronization (Dalla Bella & Andary, 2020), affording high temporal precision (≤1ms).

**Figure 1.**
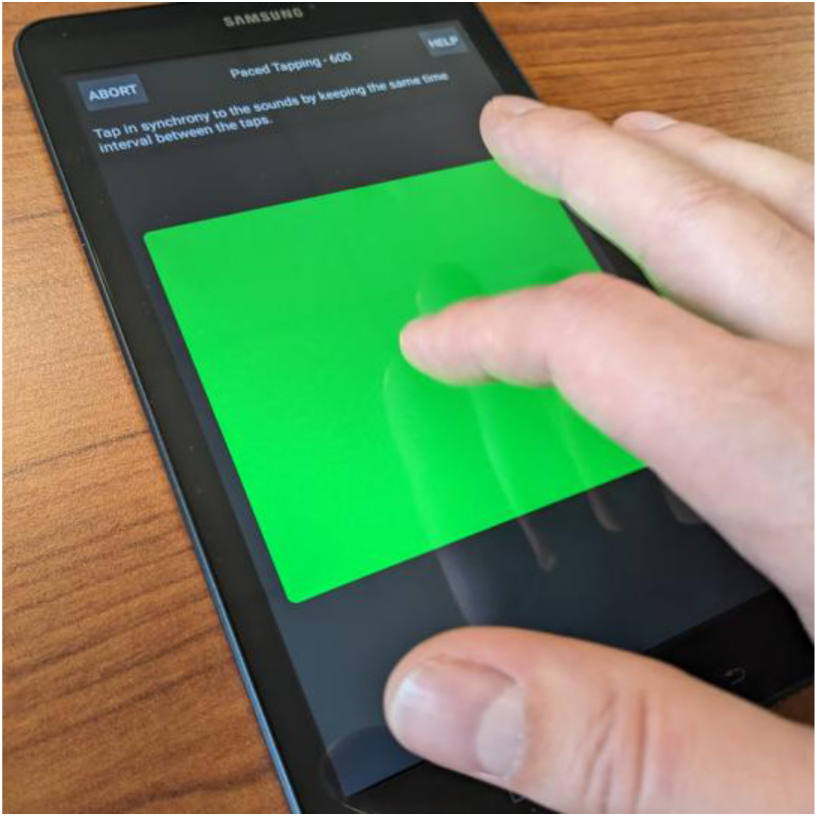
Photo of BAASTA testing with the Paced tapping task on tablet.

#### Data pre-processing and quality checks of BAASTA tablet application

Audio data collected via the BAASTA tablet app in sound-proof rooms at BRAMS laboratory were pre-processed and submitted to quality checks. As the recording of tapping data on tablet is based on audio recordings, the characteristics of the audio environment in which testing is carried out (sound-proof room in the laboratory vs. less-controlled environments for remote testing, such as schools, hospitals, at home) can potentially influence the measures of timing and synchronization. Moreover, the tapping strength and the ensuing sound can affect the capacity to detect the tapping time relative to the stimulus sound. Hence, data quality checks play an essential role to ensure that collected data are reliable and meet high measurement standards even when data collection cannot be realized in optimal lab sound-proof conditions. The sound recordings include a mixture of the tapping sounds and the stimuli, and are pre-processed by the following steps to obtain the tap times for scoring (Dalla Bella & Andary, 2020): find stimulus alignment; apply band-pass filter; find times of energy peaks; find alignment between peaks and touchscreen events; and perform windowed matching to select only the energy peaks that match a touchscreen event. A detailed description of the procedure is specified in International Patent No WO 2020/128088 A1 (Dalla Bella & Andary, 2020).

Subsequently, two diagnostic values were used to identify trials with potentially poor quality of tap detection. The first diagnostic value was the “match score”, defined as the percentage of touchscreen taps that were matched with a corresponding audio tap event. A value less than 100% can indicate overly faint tapping, and/or presence of excessive external noise in the recording such as talking, bumping of the testing apparatus, or vibration from nearby equipment. The second diagnostic value was the “match IQR”, defined as the interquartile range of the time differences between each corresponding pair of touchscreen and audio events. A value above 10 ms can indicate a misalignment of touchscreen events to energy peaks prior to matching, or signal-to-noise issues as described for the match score. A match score below 90% (because the tap scoring is tolerant to a small number of missing taps) or a match IQR exceeding 10 ms caused a trial to be flagged for manual review, and for subsequent exclusion if poor data quality was confirmed.

### Analyses of BAASTA data

For Duration discrimination, Anisochrony detection with tones and with music, we averaged the threshold obtained from the staircase procedure in the two trials. The threshold was expressed in percent of the standard duration or interval. Staircases including more than 1 false alarm among the 3 catch trials (i.e., the participant reported a difference when there was none) were rejected. In the BAT, we calculated the sensitivity index (*d’*) based on the proportions of Hits (correct detections) for misaligned metronomes, and of False alarms (when a misalignment was erroneously reported for aligned metronomes) for the entire set of 72 stimuli. Moreover, we calculated *d’* separately for each of the 3 tempi (medium, fast, and slow). Tapping data in all tasks were pre-processed following the same procedures adopted for computer-collected data (Dalla Bella, Farrugia, et al., 2017; Sowiński & Dalla Bella, 2013). We discarded taps leading to inter-tap intervals (ITIs) smaller than 100 ms (artifacts) and outliers. A tap was an “outlier” if the ITI between the actual tap and the preceding tap was smaller than Q1 – 3*Interquartile range (IQR) or greater than Q3 + 3*IQR, where Q1 is the first quartile and Q3 is the third quartile (as in Dalla Bella et al., 2017). We calculated the mean inter-tap interval (ITI, in ms) and the coefficient of variation of the ITI (CV ITI – namely, the (*SD* of the ITI) / mean ITI) as a measure of motor variability. Moreover, we computed measures of phase synchronization in the Paced tapping tasks using circular statistics (Berens, 2009; Fisher, 1993; for use in BAASTA see Dalla Bella, Farrugia, et al., 2017). Tap times in each sequence were coded as unitary vectors with angles relative to the pacing event (tone or musical beat) on a polar scale, where 360° corresponds to the inter-stimulus interval. We obtained the mean resultant vector *R* from the unit vectors corresponding to all the taps in a sequence. We used two measures of phase synchronization: consistency and accuracy. Consistency corresponds to the length of vector *R*, ranging from 0 to 1, where 0 refers to a uniform distribution of angles around the circle (i.e., lack of synchronization), and 1 to maximum consistency (no variability). Accuracy corresponds to the angle of vector R (*θ* or relative phase, in degrees), and it indicates whether participants tapped before (negative angle) or after (positive angle) the pacing event. Accuracy was calculated only if participants’ synchronization performance was above chance (null hypothesis = random distribution of data points around the circle), as assessed with the Rayleigh test for circular uniformity (Fisher, 1995; Wilkie, 1983). The null hypothesis is rejected when *R* vector length is sufficiently large according to this test. For simplicity, the vector direction in degrees was converted into percent of IOI (vector direction %) on a linear scale. Values in percent can be submitted to linear statistics that are more standard in many fields than directional statistics. Vector direction % is reported in the graphs and in the Summary norm tables in Annex 1; vector direction % and in degrees are reported in the full norm tables in Annex 2. Vector length values were submitted to a logit transformation as done in previous studies (e.g., Dalla Bella et al., 2024; Falk et al., 2015; Fiveash et al., 2022; Puyjarinet et al., 2017) to reduce data skewness, which is typical of synchronization data (e.g., Kirschner & Tomasello, 2009; Sowiński & Dalla Bella, 2013). In both Paced tapping and Synchronization-continuation tasks, the results in the two trials were averaged.

In the Adaptive tapping task, an adaptation index (as in Schwartze et al., 2011) was calculated separately for acceleration trials (i.e., faster tempi with final sequence IOIs < 600 ms) and deceleration trials (slower tempi with final sequence IOIs > 600 ms). When the value of the adaptation index is 1, the adaptation is perfect; lower and higher values than 1 indicate under correction and over correction, respectively, relative to the new metronome tempo. In the same task we calculated the sensitivity index (*d’*) for detecting tempo changes based on the proportions of Hits (when a tempo acceleration or deceleration was correctly detected) and False Alarms (when a tempo acceleration or deceleration was reported while there was no change or the opposite change).

### Statistical analyses

Statistical analyses were performed in R (version 4.2.2; R Core Team, 2022). The measure score distribution shapes were described in terms of skewness and excess kurtosis (i.e., kurtosis – 3) using the *moments* package in R. In order to reduce reliance on normal distributions in the task scores, robust rank-based versions of t-tests, ANOVAs and regressions were used where possible, using the *Rfit* package in R (Kloke & McKean, 2012). Each BAASTA variable was first tested for an effect of gender, and then separately for the effects of age (in years) and age group (categorically) using the *rfit* function. P-values were calculated from the resulting *t* statistic for effects of gender and age, and as an omnibus test via the “drop in dispersion test” for the effect of age group (Kloke & McKean, 2012).

In several tasks, additional comparisons were made to test for score differences across tempi or task conditions (unpaced initial vs. final; music 1 vs music 2). As these were within-participant comparisons, they were tested using mixed-effects models including a random participant intercept using the R packages *lmerTest* (Kuznetsova et al., 2017) and *lme4* (Bates et al., 2015), yielding *F* and *p* values calculated from Wald tests using Kenward-Roger approximated degrees of freedom (Halekoh & Højsgaard, 2014). Where applicable, effect sizes were calculated as partial omega-squared (ω_p_²), a less-biased estimator of population partial eta-squared, using the R package *effectsize* (Ben-Shachar et al., 2020).

#### Calculation of the Beat Tracking Index (BTI)

Beat Tracking Index (BTI; for usage in a previous study, Puyjarinet et al., 2017) is a global measure of beat tracking skills computed by considering the performance of both the BAT and Paced tapping with music tasks from BAASTA. The source data to compute the BTI were the overall sensitivity index (d’) obtained from the BAT, and synchronization consistency in paced tapping averaged across the two music conditions (i.e., “Sensitivity index (d’) for all trials” in the BAT, and “Consistency (logit of vector length), music 1”, and “Consistency (logit of vector length), music 2” in Paced tapping). First, values were converted to z-scores according to each measure’s mean and standard deviation across all participants (see Table S2 and Table S6), where z-score = (value – mean)/SD. A combined music score was obtained by taking the mean of each participant’s “music 1” and “music 2” z-scores. Finally, the BTI was calculated by averaging each participant’s BAT z-score with their combined music score. Note that although the BTI was calculated from z-scores, it is not itself a z-score, given that its average is not exactly 0 nor its SD is 1. Thus, BTI values are provided on an arbitrary scale.

#### Test-retest reliability

We compared the first and the second testing sessions by Wilcoxon-Mann-Whitney test (*wilcox.test* R function, “paired”). Rejection of the null hypothesis implies a significant difference between the two sessions and reveals a systematic error that may result from an experimental bias such as the effect of weariness or learning. Correlation of scores between the two sessions was calculated by Spearman rank correlation (using *cor.test* R function). To measure the agreement of BAASTA measures between test and re-test sessions, we calculated the intra-class correlation coefficient (ICC3,1), using the *icc* function from the *irr* R package (Gamer et al., 2019). The ICC3,1 (2-way model; Weir, 2005) was selected as a measure of intra-class correlation, as it does not include the variance associated with systematic error (bias) (Brozek & Alexander, 1947; Fleiss, 1986; Weir, 2005). For this reason, ICC3,1 is considered a good index of test-re-test reliability with the same experimenter. The ICC is commonly used as a measure of relative reliability, namely, the consistency of the relative position of one individual in a group (Vaz et al., 2013). Following Koo and Li (2016), ICC values were interpreted as ≥ 0.90, excellent; 0.75–0.90, good; 0.50–0.75, moderate and < 0.50, poor. The other form of reliability is absolute reliability (Fleiss, 1986; Weir, 2005), which deals with the degree to which measurements vary and provides an index of the expected trial-to-trial noise in the data. SEM, SEM%, and coefficient of reliability (CR, also referred to as the smallest real difference, or minimum difference) were used to assess absolute reliability. The SEM is expressed in the same unit as the measurement of interest and quantifies the variability between the two sessions. It is calculated as the square root of the within-subjects error variance. CR represents the 95% confidence interval, a value for which any retest score outside that interval would be interpreted to represent a change beyond the measurement error of the test (Weir, 2005). It is calculated by multiplying the SEM by 2.77 (√2 times 1.96). For comparison between tasks in different units, the SEM% and CR% were calculated by dividing the SEM or the CR by the mean of all measurements from both sessions and multiplied by 100.

## Results

### Preliminary analyses

Data were submitted to quality checks and preliminary analyses. Following a manual review of tapping trials based on automatic diagnostic measures, a total of 7 trials were excluded from scoring because of talking in the middle of the trial, and one trial was excluded because of background noise that interfered with audio-based tap detection. The mean match score of tapping trials submitted to scoring was 98.9, and the mean match IQR was 5.0. After the subsequent application of scoring criteria (see Methods, “Analyses of BAASTA data”), score completeness (percent of participants having a score on a given measure included for analysis) was 91-94% for Duration discrimination and Anisochrony detection, 100% for BAT, 95-99% for Unpaced tapping, 90-100% for Paced tapping to tones^1^, 83-99% for Paced tapping to music^1^, 100% for Synchronization-continuation, 97% for Adaptive tapping, and 94% for the BTI. In sum, diagnostic scores indicate that the data entering in the calculation of the norms were of high quality.

We ran preliminary analyses on the final dataset to test whether there were effects of gender on any task measures, finding only a greater hit rate for males on the BAT medium tempo trials (t(106) = 2.14, *p* = .035). As no further gender effects were found on the main BAT d’ measures (*p* > .15), nor across any other measures (*p* > .09), data were not separated by gender for subsequent analyses.

### Norm tables

We analyzed data collected at the Test phase to generate norm tables. The measures of timing and rhythmic abilities reported in the tables are summarized in Table 2, with reference to the norm summary tables and full tables, in which the norms are presented. The norm summary tables (Tables S1-S10 located in Annex 1) report skewness and excess kurtosis for the distributions, the mean and SD, as well as percentile scores (0, 25^th^, 50^th^, 75^th^, and 100^th^ percentile) for the overall group of participants, and separately for the four age groups (18 to 21, 22 to 29, 30 to 54, and 55 to 87 years). Significant differences between the four age groups are indicated in these summary tables, as well as differences between tempos or other comparisons within the same task (unpaced initial vs. final; music 1 vs music 2). Finally, a significant regression between the reported scores and age is reported. Normative data for perceptual tasks are presented in Table S1 (Duration discrimination, Anisochrony detection with tones, Anisochrony detection with music) and Table S2 (BAT). Data for sensorimotor tasks are reported in Tables S3 through S10 (Table S3-S4: Unpaced tapping; Table S5-S7: Paced tapping with tones and with music; Table S8: Synchronization continuation; Table S9: Adaptive tapping). More detailed tables for each task measure, including all percentile scores (from 0 through 100), and values of statistical tests are available in Tables S11-S67 (located in Annex 2). It can be seen that age group differences are found only in very few cases, namely in Anisochrony detection with tones (*F*(3,96) = 3.51, *p* < .05, ω_p_^2^ = 0.07), Unpaced tapping (for tapping rate only, in fast tapping condition, *F*(3,103) = 3.11, p < .05, ω_p_^2^ = 0.06), Paced tapping (consistency, with tones, IOI= 450ms, *F*(3,103) = 2.95, p < .05, ω_p_^2^ = 0.05; with music 1, *F*(3,102) = 2.94, p < .05, ω_p_^2^ = 0.05; and with music 2, *F*(3,98) = 3.95, *p* = .01, ω_p_^2^ = 0.08). Differences among tempo trials were found in the BAT (*F*(2,214) = 24.47, *p* < .001, ω_p_^2^ = 0.18), Paced tapping with tones (variability, *F*(2,212.4) = 17.60, *p* < .001, ω_p_^2^ = 0.13; consistency, *F*(2,212.5) = 4.12, *p* < .05, ω_p_^2^ = 0.03; and accuracy, *F*(2,200.5) = 8.45, *p* < .001, ω_p_^2^ = 0.07), and Synchronization continuation (variability, *F*(2,214.0) = 4851.15, *p* < .001, ω_p_^2^ = 0.02), showing a general tendency to perform better at medium or slow tempos. Moreover, we found differences between the initial and last trials in Unpaced tapping (for rate, in the spontaneous condition, *F*(1,103.2) = 11.32, *p* = .001, ω_p_^2^ = 0.09), and between the two musical stimuli in Paced tapping (consistency, *F*(1,101.8) = 9.90, *p* < .01, ω_p_^2^ = 0.08; and accuracy, *F*(1,91.3) = 47.06, *p* < .001, ω_p_^2^ = 0.34), showing that synchronization is more difficult for music 2 than for music 1. Finally, a significant regression between the scores and age was found in most tasks, namely in Anisochrony detection with tones (*t*(107) = 2.47, *p* < .05, ω_p_^2^ = 0.04) and with music (*t*(107) = 2.43, *p* < .05, ω_p_^2^ = 0.04), in the BAT (fast tempo trials, *t*(107) = -2.48, *p* < .05, ω_p_^2^ = 0.05), in Unpaced tapping (variability, in spontaneous tapping, *t*(107) = -2.05, *p* < .05, ω_p_^2^ = 0.03; and fast tapping, *t*(107) = -2.06, *p* < .05, ω_p_^2^ = 0.03), Paced tapping (consistency in tapping with tones with IOI = 450ms, *t*(107) = 2.32, *p* < .05, ω_p_^2^ = 0.04; accuracy in tapping with tones with IOI = 450ms, *t*(107) = 2.11, *p* < .05, ω_p_^2^ = 0.03; consistency in tapping to music 2, *t*(107) = 2.84, *p* < .01, ω_p_^2^ = 0.06), in Synchronization-continuation (rate, at the fastest tempo, *t*(107) = 3.00, *p* < .01, ω_p_^2^ = 0.07; variability, at the slower tempo, *t*(107) = -2.39, *p* < .05, ω_p_^2^ = 0.04), and in Adaptive tapping (adaptation index, for deceleration trials, *t*(107) = -2.17, *p* < .05, ω_p_^2^ = 0.03).

**Table 2.** Description of BAASTA Measures.

| Variable name | Task | Task category | Outcome measure | Figure | Norm table | Full table |
| --- | --- | --- | --- | --- | --- | --- |
| DurDisc_threshold | Duration discrimination | Perceptual | Threshold (% of IOI) | Figure 2 | Table S1 | Table S11 |
| Anisochrony_tones_threshold | Anisochrony detection with tones | Perceptual | Threshold (% of IOI) | Figure 2 | Table S1 | Table S12 |
| Anisochrony_music_threshold | Anisochrony detection with music | Perceptual | Threshold (% of IOI) | Figure 2 | Table S1 | Table S13 |
| BAT_all_dprime | Beat Alignment Test | Perceptual | Sensitivity index ($d'$) for all trials | Figure 3 | Table S2 | Table S14 |
| BAT_fast_dprime | Beat Alignment Test | Perceptual | Sensitivity index ($d'$) for fast tempo trials | Figure 3 | Table S2 | Table S15 |
| BAT_med_dprime | Beat Alignment Test | Perceptual | Sensitivity index ($d'$) for medium tempo trials | Figure 3 | Table S2 | Table S16 |
| BAT_slow_dprime | Beat Alignment Test | Perceptual | Sensitivity index ($d'$) for slow tempo trials | Figure 3 | Table S2 | Table S17 |
| BAT_all_hits | Beat Alignment Test | Perceptual | Hit rate for all trials | — | — | Table S18 |
| BAT_fast_hits | Beat Alignment Test | Perceptual | Hit rate for fast tempo trials | — | — | Table S19 |
| BAT_med_hits | Beat Alignment Test | Perceptual | Hit rate for medium tempo trials | — | — | Table S20 |
| BAT_slow_hits | Beat Alignment Test | Perceptual | Hit rate for slow tempo trials | — | — | Table S21 |
| BAT_all_fa | Beat Alignment Test | Perceptual | False alarm rate for all trials | — | — | Table S22 |
| BAT_fast_fa | Beat Alignment Test | Perceptual | False alarm rate for fast tempo trials | — | — | Table S23 |
| BAT_med_fa | Beat Alignment Test | Perceptual | False alarm rate for medium tempo trials | — | — | Table S24 |
| BAT_slow_fa | Beat Alignment Test | Perceptual | False alarm rate for slow tempo trials | — | — | Table S25 |
| Unpaced_spontaneous_right_mean_iti | Unpaced tapping | Sensorimotor | Rate (ITI in ms) of spontaneous tapping, initial trial | Figure 4 | Table S3 | Table S26 |
| Unpaced_spontaneous_final_right_mean_iti | Unpaced tapping | Sensorimotor | Rate (ITI in ms) of spontaneous tapping, final trial | Figure 4 | Table S3 | Table S27 |
| Unpaced_fast_mean_iti | Unpaced tapping | Sensorimotor | Rate (ITI in ms) of fast tapping | Figure 4 | Table S3 | Table S28 |
| Unpaced_slow_mean_iti | Unpaced tapping | Sensorimotor | Rate (ITI in ms) of slow tapping | Figure 4 | Table S3 | Table S29 |
| Unpaced_spontaneous_right_CV_iti | Unpaced tapping | Sensorimotor | Variability (CV of ITI) of spontaneous tapping, initial trial | Figure 4 | Table S4 | Table S30 |
| Unpaced_spontaneous_final_right_CV_iti | Unpaced tapping | Sensorimotor | Variability (CV of ITI) of spontaneous tapping, final trial | Figure 4 | Table S4 | Table S31 |
| Unpaced_fast_CV_iti | Unpaced tapping | Sensorimotor | Variability (CV of ITI) of fast tapping | Figure 4 | Table S4 | Table S32 |
| Unpaced_slow_CV_iti | Unpaced tapping | Sensorimotor | Variability (CV of ITI) of slow tapping | Figure 4 | Table S4 | Table S33 |
| Paced_metro_450_mean_CV_iti | Paced tapping with tones | Sensorimotor | Variability (CV of ITI), fast tempo | Figure 5 | Table S5 | Table S34 |
| Paced_metro_600_mean_CV_iti | Paced tapping with tones | Sensorimotor | Variability (CV of ITI), medium tempo | Figure 5 | Table S5 | Table S35 |
| Paced_metro_750_mean_CV_iti | Paced tapping with tones | Sensorimotor | Variability (CV of ITI), slow tempo | Figure 5 | Table S5 | Table S36 |
| Paced_metro_450_vecLenLogit | Paced tapping with tones | Sensorimotor | Consistency (logit of vector length), fast tempo | Figure 5 | Table S6 | Table S37 |
| Paced_metro_600_vecLenLogit | Paced tapping with tones | Sensorimotor | Consistency (logit of vector length), medium tempo | Figure 5 | Table S6 | Table S38 |
| Paced_metro_750_vecLenLogit | Paced tapping with tones | Sensorimotor | Consistency (logit of vector length), slow tempo | Figure 5 | Table S6 | Table S39 |
| Paced_metro_450_vecDirPct | Paced tapping with tones | Sensorimotor | Accuracy (vector direction %), fast tempo | Figure 5 | Table S7 | Table S40 |
| Paced_metro_600_vecDirPct | Paced tapping with tones | Sensorimotor | Accuracy (vector direction %), medium tempo | Figure 5 | Table S7 | Table S41 |
| Paced_metro_750_vecDirPct | Paced tapping with tones | Sensorimotor | Accuracy (vector direction %), slow tempo | Figure 5 | Table S7 | Table S42 |
| Paced_metro_450_mean_vector_direction | Paced tapping with tones | Sensorimotor | Accuracy (vector direction degrees), fast tempo | — | — | Table S43 |
| Paced_metro_600_mean_vector_direction | Paced tapping with tones | Sensorimotor | Accuracy (vector direction degrees), medium tempo | — | — | Table S44 |
| Paced_metro_750_mean_vector_direction | Paced tapping with tones | Sensorimotor | Accuracy (vector direction degrees), slow tempo | — | — | Table S45 |
| Paced_music_badine_mean_CV_iti | Paced tapping with music | Sensorimotor | Variability (CV of ITI), music 1 | Figure 5 | Table S5 | Table S46 |
| Paced_music_ross_mean_CV_iti | Paced tapping with music | Sensorimotor | Variability (CV of ITI), music 2 | Figure 5 | Table S5 | Table S47 |
| Paced_music_badine_vecLenLogit | Paced tapping with music | Sensorimotor | Consistency (logit of vector length), music 1 | Figure 5 | Table S6 | Table S48 |
| Paced_music_ross_vecLenLogit | Paced tapping with music | Sensorimotor | Consistency (logit of vector length), music 2 | Figure 5 | Table S6 | Table S49 |
| Paced_music_badine_vecDirPct | Paced tapping with music | Sensorimotor | Accuracy (vector direction %), music 1 | Figure 5 | Table S7 | Table S50 |
| Paced_music_ross_vecDirPct | Paced tapping with music | Sensorimotor | Accuracy (vector direction %), music 2 | Figure 5 | Table S7 | Table S51 |
| Paced_music_badine_mean_vector_direction | Paced tapping with music | Sensorimotor | Accuracy (vector direction degrees), music 1 | — | — | Table S52 |
| Paced_music_ross_mean_vector_direction | Paced tapping with music | Sensorimotor | Accuracy (vector direction degrees), music 2 | — | — | Table S53 |
| SyncCont_metro_450_mean_mean_iti | Synchronization-continuation | Sensorimotor | Rate (ITI in ms), fast tempo | Figure 6 | Table S8 | Table S54 |
| SyncCont_metro_600_mean_mean_iti | Synchronization-continuation | Sensorimotor | Rate (ITI in ms), medium tempo | Figure 6 | Table S8 | Table S55 |
| SyncCont_metro_750_mean_mean_iti | Synchronization-continuation | Sensorimotor | Rate (ITI in ms), slow tempo | Figure 6 | Table S8 | Table S56 |
| SyncCont_metro_450_mean_CV_iti | Synchronization-continuation | Sensorimotor | Variability (CV of ITI), fast tempo | Figure 6 | Table S8 | Table S57 |
| SyncCont_metro_600_mean_CV_iti | Synchronization-continuation | Sensorimotor | Variability (CV of ITI), medium tempo | Figure 6 | Table S8 | Table S58 |
| SyncCont_metro_750_mean_CV_iti | Synchronization-continuation | Sensorimotor | Variability (CV of ITI), slow tempo | Figure 6 | Table S8 | Table S59 |
| Adaptive_plus_75_dprime2 | Adaptive tapping | Sensorimotor | Sensitivity index ($d'$) of perceiving tempo deceleration (IOI + 75ms) | Figure 7 | Table S9 | Table S60 |
| Adaptive_plus_30_dprime2 | Adaptive tapping | Sensorimotor | Sensitivity index ($d'$) of perceiving tempo deceleration (IOI + 30ms) | Figure 7 | Table S9 | Table S61 |
| Adaptive_minus_75_dprime2 | Adaptive tapping | Sensorimotor | Sensitivity index ($d'$) of perceiving tempo acceleration (IOI - 75ms) | Figure 7 | Table S9 | Table S62 |
| Adaptive_minus_30_dprime2 | Adaptive tapping | Sensorimotor | Sensitivity index ($d'$) perceiving a tempo acceleration (IOI - 30ms) | Figure 7 | Table S9 | Table S63 |
| Adaptive_iso_600_CV_iti | Adaptive tapping | Sensorimotor | Variability (CV of ITI) of continuation tapping (isochronous condition) | Figure 7 | Table S9 | Table S64 |
| Adaptive_adaptation_index_acceleration | Adaptive tapping | Sensorimotor | Adaptation index (acceleration trials) | Figure 7 | Table S9 | Table S65 |
| Adaptive_adaptation_index_deceleration | Adaptive tapping | Sensorimotor | Adaptation index (deceleration trials) | Figure 7 | Table S9 | Table S66 |
| BTI | BTI | Composite | Beat Tracking Index | Figure 8 | Table S10 | Table S67 |

The results obtained in the Test phase with the mobile version of BAASTA were compared to the original version (Dalla Bella, et al., 2017) implemented on a computer. The comparison revealed very minor differences, none of which attained statistical significance (see Supplementary materials).

### Composite score: Beat Tracking Index

Individual performances for all participants in the Test phase for the sensorimotor and perceptual components of the BTI are presented in Figure 2 (panel A). These perceptual and sensorimotor components are positively related (*R*^2^ = .17; *F*(1,99) = 20.0; *p* < .001). However, the dispersion of the individual performances around the regression line is quite wide, and in some cases high performance on the perceptual component is associated with poor performance on the sensorimotor component, or vice-versa. Hence one of the component and associated tasks may be more capable than the other to capture poor or good performance in a rhythmic task. Calculating a composite score – the BTI (see distribution, Figure 2, Panel B) – can thus allow maximizing the capacity to capture individual variability. For example, in this sample (Test phase), 5 participants fell below 2 *SD* from the mean BTI (< -1.60), a threshold often used to determine whether a participant is impaired in beat perception or synchronization (e.g., Sowiński & Dalla Bella, 2013). This may indicate altered beat-based abilities, which would deserve further inquiry. Normative scores for the BTI are provided in Tables S10 and S57.

**Figure 2.**
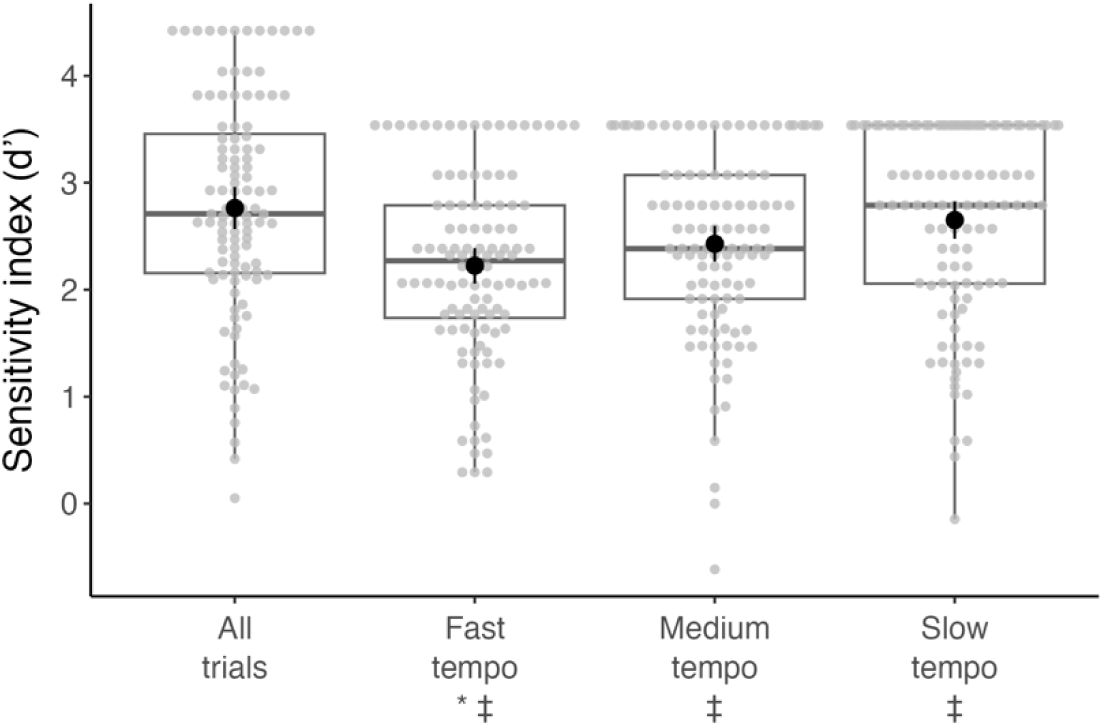
Distribution of perceptual thresholds for Duration discrimination, Anisochrony detection with tones, and Anisochrony detection with music. Means and 95% confidence intervals are indicated with solid circles and vertical lines, and boxplots indicate sample quartiles. The symbol * indicates an age regression (*p* < .05), † a difference among age groups (p < .05), and ≈ no difference between Anisochrony detection conditions (p > .05). Additional details on the Duration discrimination task are presented in Tables S2 and S11, and for the Anisochrony detection task in Tables S1, S12 and S13.

**Figure 3.**
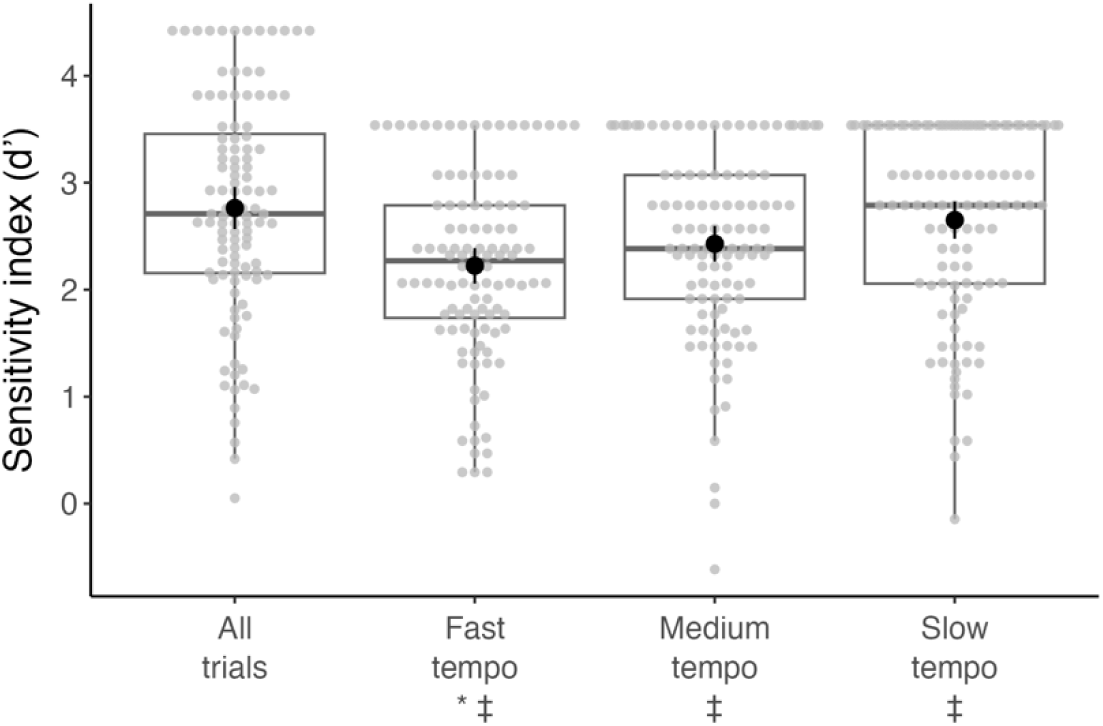
Beat alignment test (BAT) score distributions for the sensitivity index. Means and 95% confidence intervals are indicated with solid circles and vertical lines, and boxplots indicate sample quartiles. The symbol * indicates an age regression (*p* < .05), and ‡ a difference among tempo conditions (p < .05). Additional details on the BAT task are presented in Tables S2 and S14-S25.

**Figure 4.**
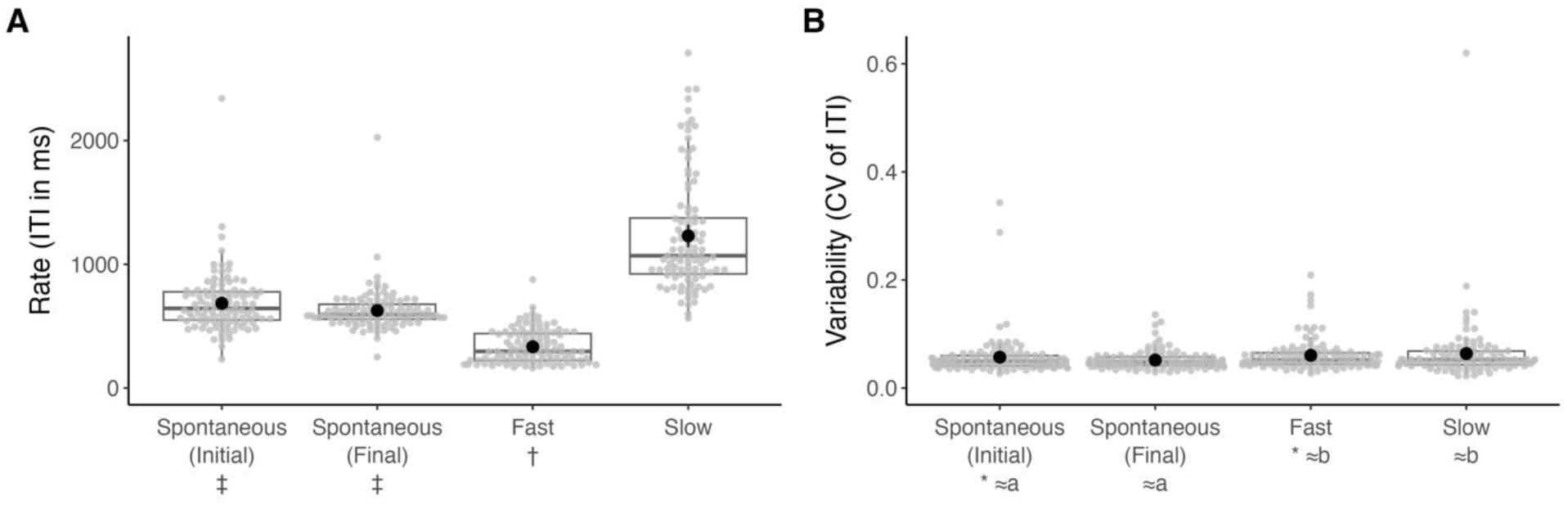
Unpaced tapping score distributions for (A) tapping rate and (B) tapping variability. Means and 95% confidence intervals are indicated with solid circles and vertical lines, and boxplots indicate sample quartiles. The symbol * indicates an age regression (p < .05), † a difference among age groups (p<.05), ‡ a difference between initial and final spontaneous conditions (p < .05), ≈*a* no difference between initial and final spontaneous conditions (p > .05), and ≈*b* no difference between fast and slow conditions (p > .05). Additional details on the Unpaced tapping task are presented in Tables S3, S4, and S26-S33.

**Figure 5.**
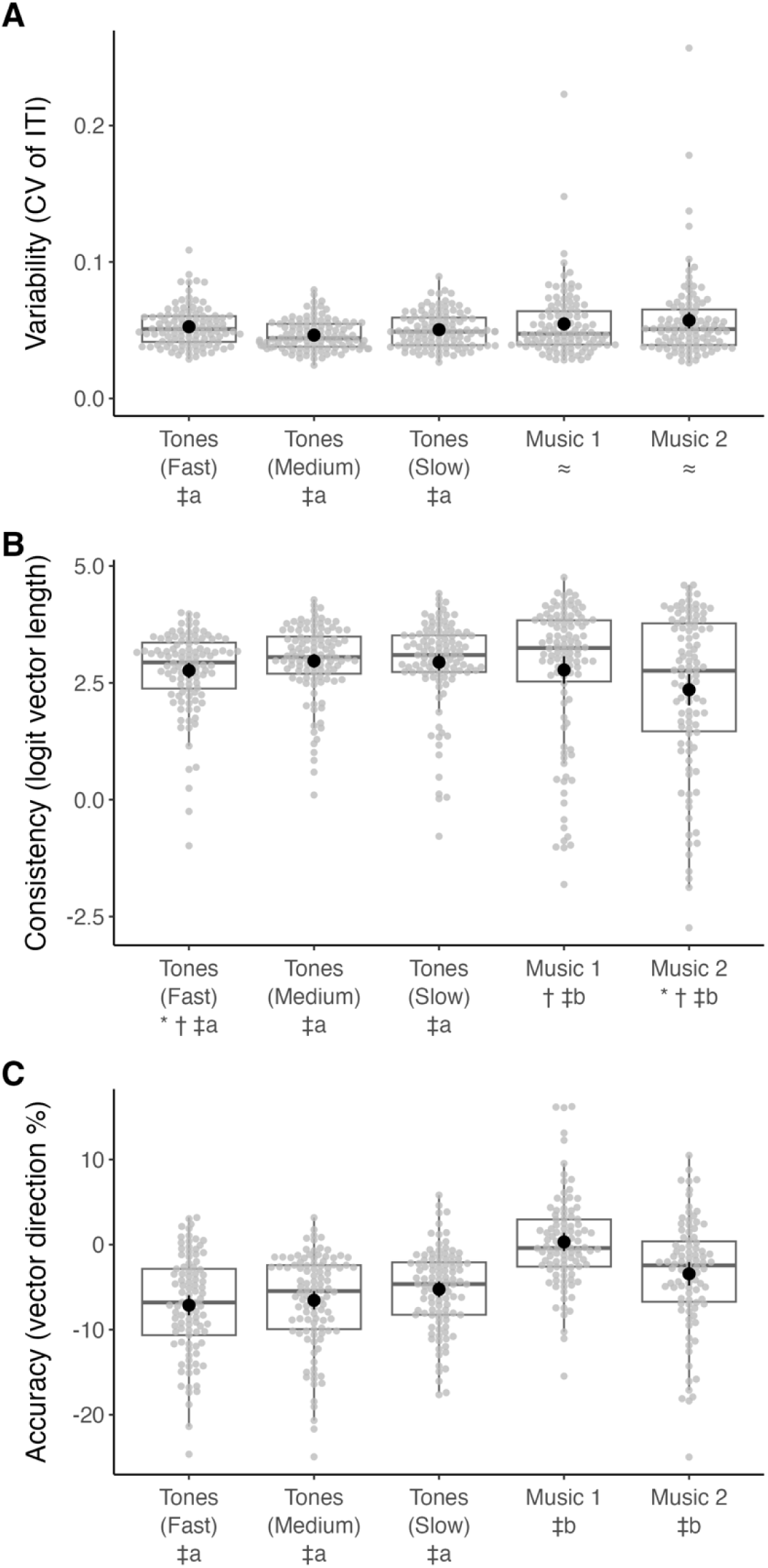
Paced tapping score distributions for (A) tapping variability, (B) synchronization consistency, and (C) synchronization accuracy. Means and 95% confidence intervals are indicated with solid circles and vertical lines, and boxplots indicate sample quartiles. Reference values for consistency (range for non-transformed vector length: 0-1) are: non-transformed 0.2 = -1.39 after logit transformation; 0.5 = 0.00; 0.75 = 1.10; 0.90 = 2.20; 0.95 = 2.94; 0.99 = 4.60. The symbol * indicates an age regression (p < .05), † a difference among age groups (p<.05), ‡*a* a significant difference among tempos (p < .05), ‡*b* a difference between music conditions (p < .05), and ≈ no difference between music conditions (p > .05). Additional details on the Paced tapping task are presented in Tables S5-S7 and S34-S53.

**Figure 6.**
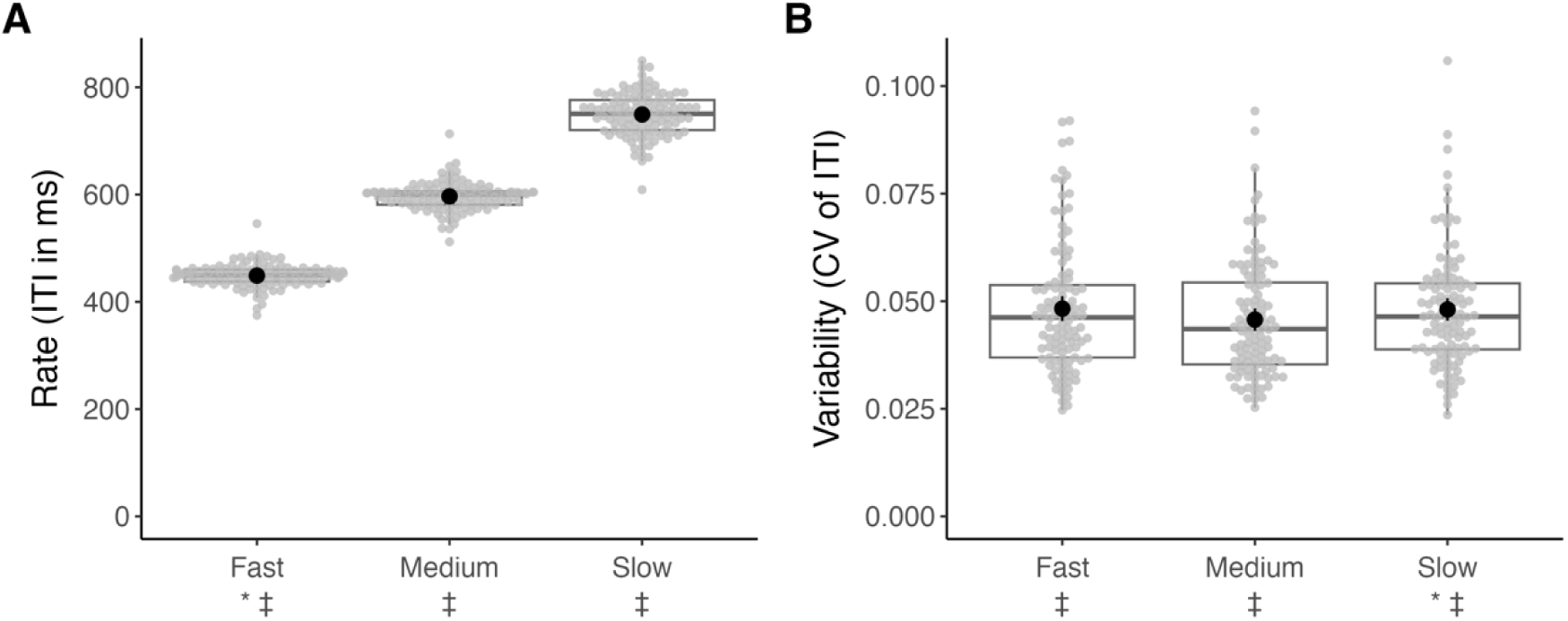
Synchronization continuation score distributions for (A) tapping rate and (B) tapping variability. Means and 95% confidence intervals are indicated with solid circles and vertical lines, and boxplots indicate sample quartiles. The symbol * indicates an age regression (p < .05), ‡a a difference of rate among tempo conditions (p < .05), and ‡b a difference of variability among tempo conditions (p < .05). Additional details for the Synchronization-continuation task are presented in Tables S8 and S54-S59.

**Figure 7.**
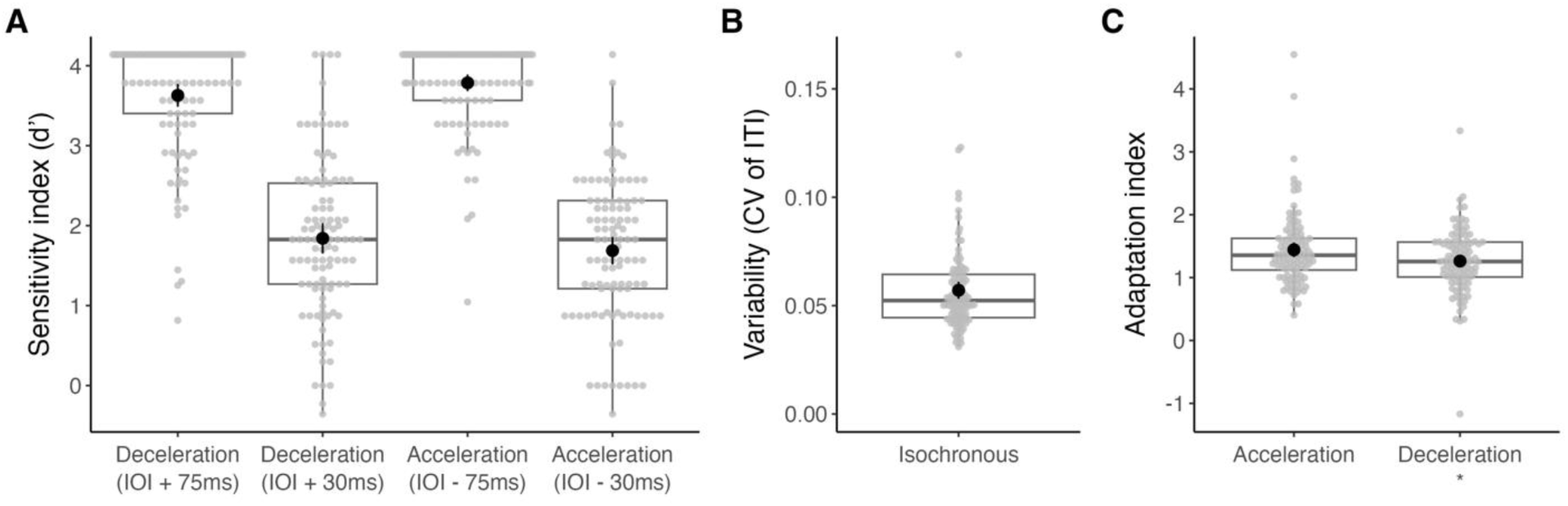
Adaptive tapping task score distributions for (A) perceptual sensitivity index, (B) tapping variability in the isochronous condition, and (C) tapping adaptation index. Means and 95% confidence intervals are indicated with solid circles and vertical lines, and boxplots indicate sample quartiles. The symbol * indicates an age regression (p < .05). Additional details for the Adaptive tapping task are presented in Tables S9 and S60-S66.

**Figure 8.**
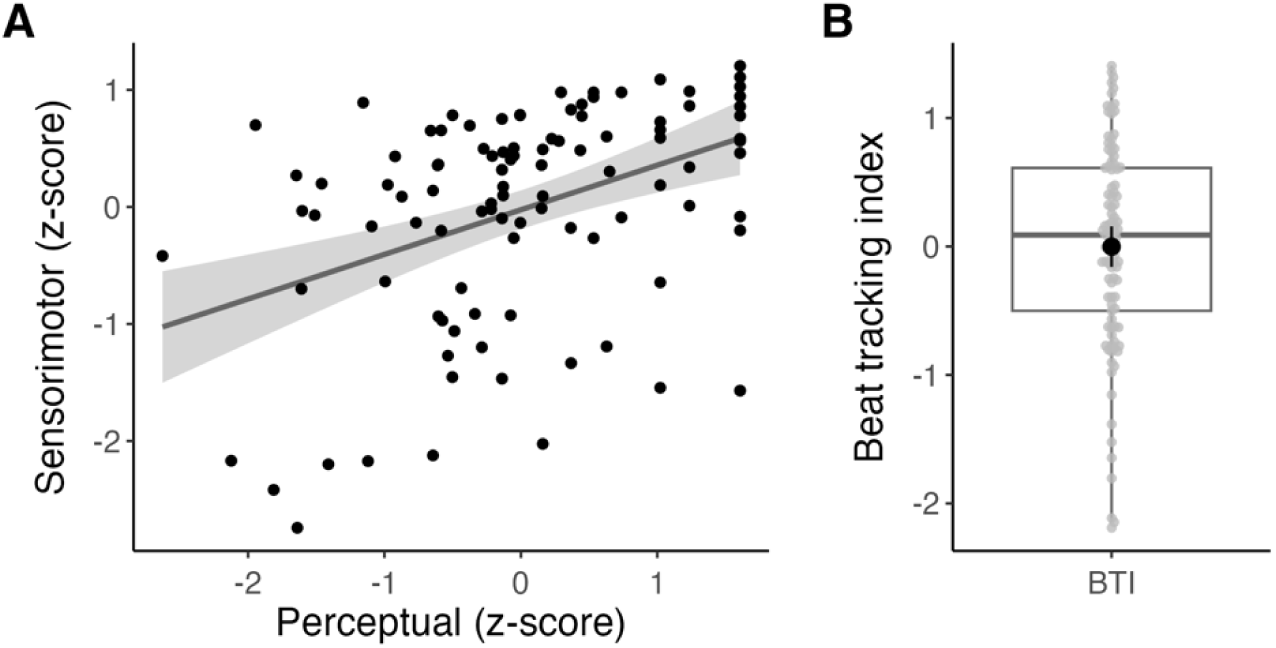
The Beat Tracking Index (BTI). (A) Relation between the BTI perceptual and sensorimotor components by participant (*n* = 101). The perceptual component is based on the overall sensitivity index (d’) obtained from the BAT, and the sensorimotor component is based on paced tapping (synchronization consistency averaged across the two music conditions). Shaded area indicates 95% confidence interval of the linear regression. B) Distribution of the BTI score. Means and 95% confidence intervals are indicated with a solid circle and vertical lines, and the boxplot indicates sample quartiles. Additional details for the BTI are presented in Tables S10 and S67.

### Test-retest reliability measures

Measures of test-retest reliability are reported in Table 3 for all the variables reported in each task and for BTI scores. Systematic error tests reveal only a few cases in which we observed a significant change of the performance from Test to Re-test, namely in the BAT (improvement at medium tempo only), Unpaced tapping (improvement of variability at spontaneous tapping, increased rate of slow tapping), Paced tapping with tones (improvement at medium tempo only), Synchronization-continuation (decreased rate at slow tempo), and Adaptive tapping (slight worsening of variability of continuation tapping in the isochronous condition). Scores at Test and Re-test are always significantly correlated, as shown by Spearman’s rank correlation (*rho* ranging between 0.28 and 0.82). Test-retest reliability for perceptual tests ranged from poor to good altogether (ICC from 0.47 to 0.77). In particular, the sensitivity index for BAT showed good test-retest reliability (ICC = 0.77). Similarly, most of the scores in sensorimotor tasks displayed moderate to good test-retest reliability, but with a few exceptions. Synchronization consistency in Paced tapping with music (with both musical stimuli) displayed good test-retest reliability (with ICC = 0.76, average across the two musical stimuli). In contrast, poor test-retest reliability was observed for variability (CV ITI) in Unpaced tapping, Paced tapping with music (but not with tones), and Synchronization-continuation (at the slow tempo). Notably, the BTI boasted good, close to excellent, test-retest reliability (ICC = 0.87). This is not surprising as the scores entering in calculation of the BTI showed themselves good to test-re-test reliability.

**Table 3.** Test-Retest Reliability.

| Task | Variable | Session 1<br>Mean (SD) | Session 2<br>Mean (SD) | Relative reliability indices |  |  | Absolute reliability indices |  |  |  |
| --- | --- | --- | --- | --- | --- | --- | --- | --- | --- | --- |
|  |  |  |  | Wilcoxon p | rho | ICC | SEM | SEM% | CR | CR% |
| Beat Alignment Test | Sensitivity index (d') for all trials | 2.81 (1.01) | 2.84 (0.99) | 0.587 | 0.72 * | 0.77 | 0.48 | 16.89 | 1.32 | 46.81 |
| Beat Alignment Test | Sensitivity index (d') for fast tempo trials | 2.27 (0.87) | 2.24 (0.87) | 0.662 | 0.60 * | 0.67 | 0.50 | 22.32 | 1.39 | 61.86 |
| Beat Alignment Test | Sensitivity index (d') for medium tempo trials | 2.45 (0.83) | 2.60 (0.84) | 0.025 * | 0.71 * | 0.73 | 0.42 | 16.64 | 1.17 | 46.14 |
| Beat Alignment Test | Sensitivity index (d') for slow tempo trials | 2.68 (0.89) | 2.69 (0.87) | 0.804 | 0.68 * | 0.70 | 0.49 | 18.12 | 1.35 | 50.23 |
| Duration discrimination | Threshold (% of IOI) | 28.2 (11.9) | 30.0 (14.1) | 0.149 | 0.67 * | 0.64 | 7.85 | 26.97 | 21.75 | 74.77 |
| Anisochrony detection with tones | Threshold (% of IOI) | 11.7 (4.9) | 11.5 (4.9) | 0.259 | 0.74 * | 0.76 | 2.42 | 20.92 | 6.72 | 57.98 |
| Anisochrony detection with music | Threshold (% of IOI) | 11.4 (6.9) | 12.3 (6.8) | 0.366 | 0.52 * | 0.47 | 4.97 | 41.93 | 13.78 | 116.22 |
| Unpaced tapping | Variability (CV of ITI) of spontaneous tapping, initial trial | 0.056 (0.034) | 0.050 (0.016) | 0.019 * | 0.55 * | 0.22 | 0.02 | 44.42 | 0.06 | 123.14 |
| Unpaced tapping | Variability (CV of ITI) of fast tapping | 0.062 (0.031) | 0.056 (0.022) | 0.112 | 0.28 * | 0.12 | 0.02 | 42.33 | 0.07 | 117.34 |
| Unpaced tapping | Variability (CV of ITI) of slow tapping | 0.057 (0.026) | 0.055 (0.041) | 0.103 | 0.44 * | 0.32 | 0.03 | 51.49 | 0.08 | 142.73 |
| Unpaced tapping | Rate (ITI in ms) of spontaneous tapping, initial trial | 693.9 (251.8) | 671.7 (255.6) | 0.167 | 0.47 * | 0.72 | 133.14 | 19.50 | 369.06 | 54.05 |
|  |  |  |  | Wilcoxon p | rho | ICC | SEM | SEM% | CR | CR% |
| Unpaced tapping | Rate (ITI in ms) of fast tapping | 333.1 (132.2) | 342.1 (114.7) | 0.144 | 0.68 * | 0.63 | 75.13 | 22.26 | 208.25 | 61.69 |
| Unpaced tapping | Rate (ITI in ms) of slow tapping | 1214.3 (453.4) | 1101.2 (385.0) | 0.002 * | 0.65 * | 0.65 | 238.20 | 20.57 | 660.26 | 57.03 |
| Paced tapping with tones | Variability (CV of ITI), fast tempo | 0.052 (0.015) | 0.052 (0.016) | 0.896 | 0.75 * | 0.77 | 0.01 | 14.29 | 0.02 | 39.62 |
| Paced tapping with tones | Variability (CV of ITI), medium tempo | 0.046 (0.012) | 0.045 (0.014) | 0.049 * | 0.76 * | 0.69 | 0.01 | 15.76 | 0.02 | 43.69 |
| Paced tapping with tones | Variability (CV of ITI), slow tempo | 0.050 (0.013) | 0.050 (0.015) | 0.355 | 0.75 * | 0.71 | 0.01 | 15.22 | 0.02 | 42.18 |
| Paced tapping with tones | Accuracy (vector direction %), fast tempo | -7.1 (6.0) | -7.0 (5.7) | 0.891 | 0.70 * | 0.63 | 3.55 | -50.39 | 9.85 | -139.67 |
| Paced tapping with tones | Accuracy (vector direction %), medium tempo | -6.4 (5.3) | -4.6 (4.9) | 0.002 * | 0.55 * | 0.47 | 3.59 | -65.43 | 9.96 | -181.35 |
| Paced tapping with tones | Accuracy (vector direction %), slow tempo | -5.4 (4.8) | -4.5 (5.5) | 0.064 | 0.62 * | 0.57 | 3.33 | -67.45 | 9.24 | -186.97 |
| Paced tapping with tones | Consistency (logit of vector length), fast tempo | 2.77 (0.85) | 2.81 (0.90) | 0.964 | 0.62 * | 0.60 | 0.56 | 20.01 | 1.55 | 55.47 |
| Paced tapping with tones | Consistency (logit of vector length), medium tempo | 2.95 (0.81) | 3.11 (0.85) | 0.008 * | 0.65 * | 0.58 | 0.53 | 17.40 | 1.46 | 48.24 |
| Paced tapping with tones | Consistency (logit of vector length), slow tempo | 2.95 (0.95) | 2.95 (0.85) | 0.955 | 0.68 * | 0.66 | 0.53 | 17.86 | 1.46 | 49.51 |
| Paced tapping with music | Variability (CV of ITI), music 1 | 0.054 (0.026) | 0.054 (0.048) | 0.203 | 0.64 * | 0.23 | 0.03 | 63.34 | 0.09 | 175.57 |
| Paced tapping with music | Variability (CV of ITI), music 2 | 0.056 (0.030) | 0.053 (0.022) | 0.184 | 0.77 * | 0.57 | 0.02 | 31.36 | 0.05 | 86.94 |
|  |  |  |  | Wilcoxon p | rho | ICC | SEM | SEM% | CR | CR% |
| Paced tapping with music | Accuracy (vector direction %), music 1 | 0.6 (4.9) | 1.0 (4.2) | 0.133 | 0.60 * | 0.47 | 3.33 | 411.61 | 9.22 | 1140.94 |
| Paced tapping with music | Accuracy (vector direction %), music 2 | -2.7 (6.0) | -2.1 (5.3) | 0.233 | 0.67 * | 0.52 | 3.92 | -161.99 | 10.86 | -449.02 |
| Paced tapping with music | Consistency (logit of vector length), music 1 | 2.86 (1.48) | 2.78 (1.65) | 0.691 | 0.78 * | 0.79 | 0.73 | 25.74 | 2.01 | 71.35 |
| Paced tapping with music | Consistency (logit of vector length), music 2 | 2.46 (1.63) | 2.61 (1.56) | 0.103 | 0.81 * | 0.74 | 0.81 | 32.10 | 2.26 | 88.98 |
| Synchronization-continuation | Variability (CV of ITI), fast tempo | 0.048 (0.015) | 0.049 (0.015) | 0.325 | 0.66 * | 0.58 | 0.01 | 20.28 | 0.03 | 56.21 |
| Synchronization-continuation | Variability (CV of ITI), medium tempo | 0.044 (0.013) | 0.044 (0.012) | 0.315 | 0.61 * | 0.66 | 0.01 | 16.73 | 0.02 | 46.38 |
| Synchronization-continuation | Variability (CV of ITI), slow tempo | 0.047 (0.013) | 0.049 (0.025) | 0.370 | 0.63 * | 0.31 | 0.02 | 35.24 | 0.05 | 97.68 |
| Synchronization-continuation | Rate (ITI in ms), fast tempo | 449.2 (21.9) | 448.3 (23.0) | 0.724 | 0.68 * | 0.75 | 11.19 | 2.49 | 31.01 | 6.91 |
| Synchronization-continuation | Rate (ITI in ms), medium tempo | 597.9 (26.1) | 597.9 (26.9) | 0.753 | 0.65 * | 0.79 | 12.33 | 2.06 | 34.17 | 5.71 |
| Synchronization-continuation | Rate (ITI in ms), slow tempo | 748.9 (39.5) | 761.9 (39.8) | 0.007 * | 0.52 * | 0.43 | 29.08 | 3.85 | 80.61 | 10.67 |
| Adaptive tapping | Adaptation index (acceleration trials) | 1.434 (0.599) | 1.421 (0.535) | 0.769 | 0.47 * | 0.62 | 0.35 | 24.67 | 0.98 | 68.39 |
| Adaptive tapping | Adaptation index (deceleration trials) | 1.254 (0.405) | 1.205 (0.495) | 0.457 | 0.41 * | 0.44 | 0.34 | 27.62 | 0.94 | 76.56 |
|  |  |  |  | Wilcoxon p | rho | ICC | SEM | SEM% | CR | CR% |
| Adaptive tapping | Variability (CV of ITI) of continuation tapping (isochronous condition) | 0.056 (0.020) | 0.058 (0.021) | 0.020 * | 0.75 * | 0.80 | 0.01 | 15.89 | 0.02 | 44.04 |
| Adaptive tapping | Sensitivity index (d') perceiving a tempo acceleration (IOI - 30ms) | 1.77 (0.87) | 1.74 (0.98) | 0.742 | 0.53 * | 0.52 | 0.64 | 36.61 | 1.78 | 101.49 |
| Adaptive tapping | Sensitivity index (d') of perceiving tempo acceleration (IOI - 75ms) | 3.79 (0.54) | 3.76 (0.56) | 0.476 | 0.51 * | 0.71 | 0.30 | 7.83 | 0.82 | 21.69 |
| Adaptive tapping | Sensitivity index (d') of perceiving tempo deceleration (IOI + 30ms) | 1.89 (0.97) | 1.99 (0.91) | 0.332 | 0.54 * | 0.56 | 0.63 | 32.26 | 1.74 | 89.42 |
| Adaptive tapping | Sensitivity index (d') of perceiving tempo deceleration (IOI + 75ms) | 3.65 (0.75) | 3.67 (0.64) | 0.854 | 0.38 * | 0.50 | 0.49 | 13.44 | 1.36 | 37.26 |
| BTI | Beat Tracking Index | 0.033 (0.820) | -0.011 (0.865) | 0.266 | 0.82 * | 0.87 | 0.31 | —<sup>a</sup> | 0.85 | —<sup>a</sup> |

\* $p < .05$ . <sup>a</sup> The SEM% and CR% are not presented for the BTI because its values are centered on zero. ICC: intra-class correlation coefficient;
CR: coefficient of reliability.

## Discussion

The goals of the present study were to present a mobile solution for BAASTA (app on a tablet device) introduced in previous studies (Bégel et al., 2018, 2022; Dauvergne et al., 2018; Puyjarinet et al., 2017, 2019), affording an accurate measure of perceptual and sensorimotor timing abilities. Moreover, we provide norms and test-retest reliability measures for this version of the battery, and a summary metric for beat-based timing, reflecting both perceptual and motor skills. The first two goals address the need for a portable and reliable solution when testing timing and rhythmic abilities outside of the lab, and for normative data from a healthy adult population. The last goal addresses the problem inherent in BAASTA’s testing time (around 2 hours), which may prevent the use of the full battery when only a general estimate of rhythmic abilities is needed.

Since mobile devices have limitations regarding the measurement of motor performance using a touchscreen, we paid particular attention to provide a method capable of lab-quality recording performance. The proposed method is based on audio analysis, by exploiting the capacity of mobile devices to record sound in the nearby environment, including the tapping sound on the touchscreen (Dalla Bella & Andary, 2020). This solution benefits from the high resolution of the recorded auditory signal, affords high temporal precision (≤1ms), is device-independent (for a similar web-based implementation, see also Anglada-Tort et al., 2022), and does not require prior calibration. This tablet version of BAASTA could successfully replicate the results obtained with paced tapping on a lab computer (Zagala et al., 2021), and provides similar results as compared to the original version of BAASTA (Dalla Bella et al., 2017). Despite these advantages, however, one of the shortcomings of relying on auditory recordings is the sensitivity of the measurement to ambient noise. Even though BAASTA tests should be ideally administered in a sound-proof room, this condition may be harder to meet when the tests are performed outside of the lab (in a school class, hospital), or in more ecological settings. Thus, we developed methods affording data quality checks for ensuring the reliability of the collected data when BAASTA-tablet is used in less-than-ideal testing settings. To this aim, we proposed two metrics (match score and match IQR) which can attest whether data quality is excellent, identify any trials needing reviewing, and provide a basis to select the best quality trials in situations where we know that the quality of audio recordings may be affected (e.g., testing in ecological settings). These quality checks, provided on a trial-by-trial basis, are critical for BAASTA users to make informed decisions about the data before they are included in further analyses or used for diagnosis purposes in clinical studies.

To our knowledge, this is the first battery of timing and sensorimotor tests to be validated on a mobile device in the adult population across the lifespan. Norms play an important role in defining cut-off scores for impaired performance in various clinical groups, and for ensuring comparability across the lifespan and studies. The norms are provided for four age groups of comparable sample size. Even though we observed between-group age differences only in a few cases, we found relations between most BAASTA tasks and age (as a continuous dimension). These findings suggest that age should be considered to compare data collected in future studies with the battery, by choosing the appropriate age group and/or testing condition or stimulus material. Test-retest reliability measures reveal differences among BAASTA tasks, spanning from poor to good reliability. For example, among perceptual tests, the BAT (Iversen & Patel, 2008), and among sensorimotor tests, paced tapping to music (e.g., Repp, 2005; Repp & Su, 2013), both display good test-retest reliability (with ICC ∼ 0.76). Only one measure across tasks revealed quite low test-retest reliability, namely motor variability (CV of ITI; e.g., ICC < 0.35 in unpaced tapping tests, or in individual conditions in paced tapping with music or synchronization-continuation). This suggests that measures such as accuracy/consistency in paced tapping are preferable over other measures (e.g., motor variability) to provide a robust assessment of rhythmic abilities, in particular when re-testing is a critical part of the experimental design (e.g., in longitudinal protocols, clinical trials).

Notably, the aforementioned tasks showed good test-retest reliability (BAT and paced tapping with music) and are also those found to capture most variability in rhythmic capacities across different BAASTA tests as shown in a recent machine-learning study (Dalla Bella et al., 2024). This finding supports the choice of the two tasks to compute a composite score – the BTI – as a summary measure of beat-based rhythmic abilities (see also, Puyjarinet et al., 2017 for a first presentation of the BTI). Two measures from these tasks (i.e., the sensitivity index from BAT, and synchronization consistency from paced tapping with music) are significantly related, but quite weakly (< 20% of accounted variance; a stronger correlation was found in previous studies focusing on children and adults with neurodevelopmental disorders, Puyjarinet et al., 2017, and on adults with altered beat synchronization, Tranchant et al., 2021). In a few cases, participants show dissociations between the two tasks. That the performance in beat perception and synchronization tasks can sometimes dissociate is not a novel finding, as shown in single-case evidence of poor synchronization to the beat in the presence of spared beat perception (Sowiński & Dalla Bella, 2013), or the opposite dissociation (Bégel et al., 2017). Altogether these findings suggest that the mechanisms underlying beat perception and production (Cannon & Patel, 2021; Patel & Iversen, 2014) may not be completely overlapping. Having a perceptual and a sensorimotor score both contribute to a composite score like the BTI is particularly appealing to capture individual differences in rhythmic abilities (see also Fiveash et al., 2022), while maximizing the chances of detecting poor performance in each of the components. In a previous study, we used BTI to express individual differences in rhythmic abilities in children and adults with and without neurodevelopmental disorders (ADHD; Puyjarinet et al., 2017). BTI could successfully capture relations between rhythm abilities and executive functions, showing that among children and adults with ADHD, those with the lowest scores in BTI (poor beat trackers) had poorer performance in flexibility and inhibition tasks. Owing to BTI’s excellent test-retest reliability (ICC = 0.87) and efficiency (< 15 minutes of testing, with a minimal set of conditions), we advise a more extensive use of this metric. Using only two BAASTA tests, with a gain in time over 80% relative to the full battery (two hours; Dalla Bella, Farrugia, et al., 2017) is also particularly appropriate in situations where testing time is very limited, such as in large clinical studies, in which rhythmic abilities are tested among many other cognitive capacities.

In sum, the adaptation of BAASTA for use on a mobile device, the availability of norms for the adult population, and the possibility of using BTI as a valuable composite score is expected to increase the use of the battery in adult healthy individuals to quantify individual differences (e.g., Dalla Bella et al., 2024; Fiveash et al., 2022), and in clinical populations with poor timing and rhythmic abilities (e.g., Parkinson, Benoit et al., 2014; Grahn & Brett, 2009; ADHD, Puyjarinet et al., 2017; speech and language impairement, Bégel et al., 2022; Corriveau & Goswami, 2009; Falk et al., 2015; Ladányi et al., 2020; Lense et al., 2021; Autism Spectrum Disorder, Allman et al., 2011). A mobile solution for assessing timing and rhythmic abilities, particularly appealing due to its portability and accessibility to the public, facilitates remote testing of participants with disabilities who cannot reach the laboratory, while ensuring high data reliability (with the caveats linked with the use of audio signal). This development is in line with a more general tendency to exploit technological applications for assessment and intervention purposes (e.g. of mobile medicine, (Cerrato & Halamka, 2019), recently extending to musical assessment and rhythmic training (for reviews, Agres et al., 2021; Dalla Bella, 2022). Eventually, it is expected that mobile tasks like the ones presented in BAASTA will be integrated in mobile applications devoted to rhythm-based training programs, with the purpose of personalizing the intervention (e.g., Dotov et al., 2019).

## Supporting information

Annex 1 - summary norm tables

Annex 2 - full norm tables

BAASTA stimuli examples

## Acknowledgements and funding information

The research was funded by a maturation grant from French SATT AxLR (YOURHYTHM project), and by funding from a Discovery Grant (RGPIN-2019-05453) from the Natural Sciences and Engineering Research Council of Canada (NSERC), and by the Canada Research Chair program (CRC in music auditory-motor skill learning and new technologies) to SDB. We wish to thank Camille Gaillard, Paulina Herrera, Gabrielle Tessier, Anais Bédard, Justine Daoust, and Mathilde Rochette for their help in participant recruitment and data collection.

## Open practices statement

The anonymized data collected in this experiment and used to generate BAASTA norms are available as open data via the BioRXiv online data repository: https://www.biorxiv.org/content/10.1101/2023.07.21.550031v2.supplementary-material. The experiment was not preregistered.

## Footnotes

1 In the paced tapping tasks, accuracy is not calculated when a participant’s tapping consistency is extremely low (i.e., when the Rayleigh test is not significant), causing some accuracy scores to be absent for participants at the lower margin of performance.

