## Supplementary material for "Mobile version of the Battery for the Assessment of Auditory Sensorimotor and Timing Abilities (BAASTA): Implementation and adult norms": Annex 1 - summary norm tables

#### Annex 1: Norm summary tables

### Norm summary tables by task

***Table S1:*** *Norms for Thresholds of Duration Discrimination and Anisochrony Detection*

| | | | | | Percentiles | | | | |
|---|---|---|---|---|---|---|---|---|---|
| **Group** | **N (N female)** | **Skewness** | **Excess kurtosis** | **Mean (SD)** | **0** | **25** | **50** | **75** | **100** |
| **Duration discrimination – Threshold (% of IOI)** | | | | | | | | | |
| All | 101 (70) | 0.89 | 0.71 | 29.1 (12.7) | 7.6 | 19.3 | 26.9 | 35.4 | 67.6 |
| Age 18 to 21 | 25 (16) | 1.21 | 2.81 | 26.4 (9.7) | 10.3 | 19.7 | 26.0 | 30.4 | 58.1 |
| Age 22 to 29 | 29 (19) | 0.56 | -0.05 | 29.2 (11.5) | 11.8 | 20.7 | 29.8 | 35.6 | 59.3 |
| Age 30 to 54 | 25 (16) | 0.95 | -0.14 | 30.9 (17.0) | 8.5 | 18.4 | 26.7 | 43.1 | 67.6 |
| Age 55 to 87 | 22 (19) | -0.11 | -1.06 | 29.9 (12.0) | 7.6 | 19.8 | 32.4 | 38.9 | 48.6 |
| **Anisochrony detection with tones – Threshold (% of IOI) ≈** | | | | | | | | | |
| All * | 100 (69) | 0.41 | -0.69 | 12.4 (5.1) | 3.8 | 8.4 | 11.6 | 16.3 | 24.0 |
| Age 18 to 21 † | 26 (18) | 0.47 | -0.74 | 12.2 (5.5) | 3.8 | 8.1 | 11.1 | 16.4 | 23.2 |
| Age 22 to 29 | 26 (16) | 0.41 | -0.85 | 10.9 (4.3) | 4.7 | 7.5 | 10.1 | 14.1 | 19.6 |
| Age 30 to 54 | 23 (15) | 0.37 | -0.88 | 11.4 (4.3) | 5.3 | 8.1 | 10.5 | 14.5 | 19.8 |
| Age 55 to 87 | 25 (20) | -0.01 | -0.87 | 15.2 (5.3) | 4.4 | 11.3 | 14.4 | 19.0 | 24.0 |
| **Anisochrony detection with music – Threshold (% of IOI) ≈** | | | | | | | | | |
| All * | 98 (68) | 0.95 | 0.11 | 12.0 (7.1) | 3.2 | 6.5 | 10.6 | 15.9 | 32.0 |
| Age 18 to 21 | 26 (18) | 1.17 | 0.36 | 11.0 (7.6) | 3.6 | 5.1 | 7.8 | 13.9 | 29.2 |
| Age 22 to 29 | 25 (17) | 1.25 | 1.25 | 11.0 (5.4) | 4.1 | 7.7 | 9.7 | 12.3 | 26.1 |
| Age 30 to 54 | 24 (14) | 0.82 | -0.58 | 11.7 (7.8) | 3.2 | 5.2 | 9.5 | 15.9 | 27.9 |
| Age 55 to 87 | 23 (19) | 0.74 | 0.07 | 14.6 (7.0) | 4.7 | 9.9 | 13.4 | 19.1 | 32.0 |

*Note.* Additional details for Duration discrimination task are presented in Table S11, and for the Anisochrony detection task in Tables S12-S13.

* Significant age regression ($p < .05$). † Significant difference among age groups ($p < .05$). ≈ No difference between Anisochrony detection conditions ($p > .05$).

***Table S2:*** *Norms for Beat Alignment Test Sensitivity Index*

| | | | | | Percentiles | | | | |
|---|---|---|---|---|---|---|---|---|---|
| **Group** | **N (N female)** | **Skewness** | **Excess kurtosis** | **Mean (SD)** | **0** | **25** | **50** | **75** | **100** |
| **Beat Alignment Test – Sensitivity index (d') for all trials** | | | | | | | | | |
| All | 108 (74) | -0.23 | -0.40 | 2.76 (1.03) | 0.05 | 2.15 | 2.71 | 3.46 | 4.42 |
| Age 18 to 21 | 27 (18) | -0.86 | 0.41 | 3.02 (1.06) | 0.42 | 2.62 | 2.93 | 3.82 | 4.42 |
| Age 22 to 29 | 29 (19) | 0.16 | -1.03 | 2.83 (1.10) | 0.89 | 2.13 | 2.54 | 3.82 | 4.42 |
| Age 30 to 54 | 26 (16) | -0.58 | 0.06 | 2.69 (1.07) | 0.05 | 2.22 | 2.84 | 3.31 | 4.42 |
| Age 55 to 87 | 26 (21) | 0.22 | -0.39 | 2.51 (0.87) | 1.06 | 1.88 | 2.58 | 3.03 | 4.42 |
| **Beat Alignment Test – Sensitivity index (d') for fast tempo trials ‡** | | | | | | | | | |
| All * | 108 (74) | -0.24 | -0.44 | 2.22 (0.87) | 0.29 | 1.74 | 2.27 | 2.79 | 3.54 |
| Age 18 to 21 | 27 (18) | -1.16 | 1.34 | 2.46 (0.80) | 0.29 | 2.19 | 2.57 | 3.07 | 3.54 |
| Age 22 to 29 | 29 (19) | -0.13 | -0.91 | 2.31 (0.96) | 0.29 | 1.60 | 2.32 | 3.54 | 3.54 |
| Age 30 to 54 | 26 (16) | -0.20 | -0.09 | 2.12 (0.83) | 0.29 | 1.67 | 2.06 | 2.52 | 3.54 |
| Age 55 to 87 | 26 (21) | 0.24 | -0.25 | 1.98 (0.82) | 0.59 | 1.66 | 1.98 | 2.34 | 3.54 |
| **Beat Alignment Test – Sensitivity index (d') for medium tempo trials ‡** | | | | | | | | | |
| All | 108 (74) | -0.72 | 0.68 | 2.43 (0.87) | -0.62 | 1.91 | 2.38 | 3.07 | 3.54 |
| Age 18 to 21 | 27 (18) | -1.65 | 2.47 | 2.61 (1.04) | -0.62 | 2.32 | 2.79 | 3.54 | 3.54 |
| Age 22 to 29 | 29 (19) | -0.17 | -0.89 | 2.44 (0.84) | 0.59 | 1.77 | 2.38 | 3.07 | 3.54 |
| Age 30 to 54 | 26 (16) | -0.36 | -0.22 | 2.30 (0.93) | 0.00 | 1.78 | 2.35 | 2.79 | 3.54 |
| Age 55 to 87 | 26 (21) | 0.11 | -0.88 | 2.36 (0.63) | 1.31 | 1.92 | 2.38 | 2.79 | 3.54 |
| **Beat Alignment Test – Sensitivity index (d') for slow tempo trials ‡** | | | | | | | | | |
| All | 108 (74) | -0.83 | -0.18 | 2.65 (0.91) | -0.15 | 2.06 | 2.79 | 3.54 | 3.54 |
| Age 18 to 21 | 27 (18) | -1.00 | -0.03 | 2.83 (0.86) | 0.59 | 2.22 | 3.07 | 3.54 | 3.54 |
| Age 22 to 29 | 29 (19) | -0.83 | 0.17 | 2.73 (0.83) | 0.44 | 2.06 | 2.79 | 3.54 | 3.54 |

| Group | N (N female) | Skewness | Excess kurtosis | Mean (SD) | Percentiles | | | | |
|---|---|---|---|---|---|---|---|---|---|
| | | | | | **0** | **25** | **50** | **75** | **100** |
| Age 30 to 54 | 26 (16) | -1.05 | 0.05 | 2.62 (1.08) | -0.15 | 2.10 | 3.07 | 3.54 | 3.54 |
| Age 55 to 87 | 26 (21) | -0.20 | -1.33 | 2.42 (0.86) | 1.02 | 1.52 | 2.68 | 3.07 | 3.54 |

*Note.* Additional details for the Beat Alignment Test are presented in Tables S14-S25.

* Significant age regression ($p < .05$). ‡ Significant difference between tempo conditions ($p < .05$).

***Table S3:*** *Norms for Unpaced Tapping Rate*

| Group | N (N female) | Skewness | Excess kurtosis | Mean (SD) | Percentiles 0 | 25 | 50 | 75 | 100 |
|---|---|---|---|---|---|---|---|---|---|
| **Unpaced tapping – Rate (ITI in ms) of spontaneous tapping, initial trial ‡** | | | | | | | | | |
| All | 106 (72) | 3.22 | 18.96 | 685.3 (243.0) | 233.9 | 549.4 | 643.8 | 777.8 | 2341.5 |
| Age 18 to 21 | 26 (17) | 0.51 | -0.11 | 721.1 (225.8) | 336.3 | 536.1 | 684.6 | 876.4 | 1304.6 |
| Age 22 to 29 | 29 (19) | 0.78 | 1.13 | 661.1 (152.7) | 386.4 | 549.0 | 645.1 | 745.1 | 1109.7 |
| Age 30 to 54 | 26 (16) | 2.76 | 9.19 | 718.9 (394.5) | 233.9 | 477.9 | 629.1 | 842.5 | 2341.5 |
| Age 55 to 87 | 25 (20) | 0.38 | -1.24 | 641.0 (102.2) | 488.8 | 560.2 | 607.9 | 736.5 | 839.5 |
| **Unpaced tapping – Rate (ITI in ms) of spontaneous tapping, final trial ‡** | | | | | | | | | |
| All | 104 (72) | 4.86 | 35.75 | 626.6 (177.3) | 250.6 | 556.7 | 593.4 | 677.2 | 2025.4 |
| Age 18 to 21 | 26 (17) | 0.62 | -0.42 | 626.3 (119.6) | 459.0 | 536.3 | 590.5 | 709.8 | 896.7 |
| Age 22 to 29 | 28 (19) | 0.49 | -0.23 | 605.0 (88.9) | 450.3 | 550.1 | 594.6 | 658.2 | 814.9 |
| Age 30 to 54 | 25 (16) | 4.10 | 16.56 | 655.9 (297.4) | 401.7 | 570.3 | 586.7 | 648.2 | 2025.4 |
| Age 55 to 87 | 25 (20) | 0.56 | 2.73 | 621.8 (145.8) | 250.6 | 542.6 | 613.9 | 699.4 | 1058.6 |
| **Unpaced tapping – Rate (ITI in ms) of fast tapping** | | | | | | | | | |
| All | 107 (73) | 0.96 | 1.03 | 333.9 (134.5) | 161.0 | 223.0 | 296.2 | 440.4 | 876.1 |
| Age 18 to 21 † | 27 (18) | 0.44 | -0.83 | 345.3 (132.7) | 176.2 | 238.6 | 310.5 | 452.9 | 652.1 |
| Age 22 to 29 | 29 (19) | 1.63 | 3.36 | 334.3 (153.5) | 168.8 | 223.1 | 296.2 | 426.4 | 876.1 |
| Age 30 to 54 | 26 (16) | 1.01 | -0.27 | 276.6 (105.0) | 161.0 | 197.8 | 243.2 | 307.1 | 520.4 |
| Age 55 to 87 | 25 (20) | 0.16 | -1.13 | 380.7 (126.0) | 195.5 | 269.9 | 363.0 | 466.3 | 603.3 |
| **Unpaced tapping – Rate (ITI in ms) of slow tapping** | | | | | | | | | |
| All | 103 (71) | 1.18 | 0.66 | 1230.1 (465.8) | 564.3 | 922.4 | 1069.4 | 1374.5 | 2708.4 |
| Age 18 to 21 | 25 (16) | 0.86 | -0.09 | 1325.9 (394.6) | 835.7 | 1008.0 | 1244.8 | 1473.6 | 2338.6 |
| Age 22 to 29 | 28 (19) | 2.14 | 4.73 | 1107.3 (440.5) | 687.6 | 820.2 | 961.1 | 1160.9 | 2708.4 |

| | | | | | Percentiles | | | | |
|---|---|---|---|---|---|---|---|---|---|
| **Group** | **N (N female)** | **Skewness** | **Excess kurtosis** | **Mean (SD)** | **0** | **25** | **50** | **75** | **100** |
| Age 30 to 54 | 25 (16) | 0.63 | -0.99 | 1351.7 (545.3) | 599.9 | 924.8 | 1207.3 | 1760.0 | 2416.7 |
| Age 55 to 87 | 25 (20) | 1.42 | 1.58 | 1150.3 (448.8) | 564.3 | 935.3 | 1039.0 | 1276.3 | 2413.1 |

*Note.* Additional details for the Unpaced tapping task are presented in Tables S26-S33.

† Significant difference among age groups ($p<.05$). ‡ Significant difference between initial and final spontaneous conditions ($p < .05$).

***Table S4:*** *Norms for Unpaced Tapping Variability*

| | | | | | Percentiles | | | | |
|---|---|---|---|---|---|---|---|---|---|
| **Group** | **N (N female)** | **Skewness** | **Excess kurtosis** | **Mean (SD)** | **0** | **25** | **50** | **75** | **100** |
| **Unpaced tapping – Variability (CV of ITI) of spontaneous tapping, initial trial ≈ [a]** | | | | | | | | | |
| All * | 106 (72) | 5.59 | 34.90 | 0.057 (0.039) | 0.027 | 0.041 | 0.050 | 0.060 | 0.343 |
| Age 18 to 21 | 26 (17) | 4.17 | 17.00 | 0.066 (0.059) | 0.036 | 0.043 | 0.053 | 0.062 | 0.343 |
| Age 22 to 29 | 29 (19) | 4.32 | 18.88 | 0.062 (0.046) | 0.035 | 0.044 | 0.054 | 0.059 | 0.288 |
| Age 30 to 54 | 26 (16) | 0.57 | -0.61 | 0.051 (0.015) | 0.031 | 0.040 | 0.049 | 0.061 | 0.085 |
| Age 55 to 87 | 25 (20) | 2.68 | 8.97 | 0.049 (0.016) | 0.027 | 0.042 | 0.047 | 0.052 | 0.114 |
| **Unpaced tapping – Variability (CV of ITI) of spontaneous tapping, final trial ≈ [a]** | | | | | | | | | |
| All | 104 (72) | 2.26 | 6.60 | 0.052 (0.018) | 0.029 | 0.042 | 0.048 | 0.058 | 0.136 |
| Age 18 to 21 | 26 (17) | 1.91 | 4.03 | 0.054 (0.020) | 0.031 | 0.042 | 0.049 | 0.058 | 0.123 |
| Age 22 to 29 | 28 (19) | 2.24 | 5.52 | 0.055 (0.022) | 0.032 | 0.044 | 0.050 | 0.059 | 0.136 |
| Age 30 to 54 | 25 (16) | 2.24 | 6.56 | 0.049 (0.018) | 0.029 | 0.037 | 0.045 | 0.055 | 0.117 |
| Age 55 to 87 | 25 (20) | 1.17 | 1.20 | 0.049 (0.012) | 0.032 | 0.042 | 0.045 | 0.053 | 0.082 |
| **Unpaced tapping – Variability (CV of ITI) of fast tapping ≈ [b]** | | | | | | | | | |
| All * | 107 (73) | 2.67 | 8.64 | 0.061 (0.029) | 0.027 | 0.044 | 0.052 | 0.065 | 0.209 |
| Age 18 to 21 | 27 (18) | 2.56 | 5.98 | 0.067 (0.039) | 0.036 | 0.046 | 0.058 | 0.068 | 0.209 |
| Age 22 to 29 | 29 (19) | 1.87 | 3.90 | 0.065 (0.025) | 0.036 | 0.048 | 0.060 | 0.067 | 0.152 |
| Age 30 to 54 | 26 (16) | 2.46 | 6.15 | 0.056 (0.028) | 0.032 | 0.041 | 0.049 | 0.057 | 0.162 |
| Age 55 to 87 | 25 (20) | 1.37 | 1.80 | 0.053 (0.019) | 0.027 | 0.042 | 0.047 | 0.062 | 0.111 |
| **Unpaced tapping – Variability (CV of ITI) of slow tapping ≈ [b]** | | | | | | | | | |
| All | 103 (71) | 7.46 | 64.15 | 0.064 (0.061) | 0.022 | 0.043 | 0.052 | 0.068 | 0.620 |
| Age 18 to 21 | 25 (16) | 4.23 | 17.18 | 0.085 (0.115) | 0.022 | 0.045 | 0.053 | 0.076 | 0.620 |
| Age 22 to 29 | 28 (19) | 3.06 | 10.60 | 0.056 (0.031) | 0.029 | 0.041 | 0.048 | 0.056 | 0.189 |

| | | | | | | | | | |
|---|---|---|---|---|---|---|---|---|---|
| Age 30 to 54 | 25 (16) | 1.66 | 3.37 | 0.055 (0.026) | 0.023 | 0.037 | 0.053 | 0.065 | 0.140 |
| Age 55 to 87 | 25 (20) | 0.71 | 0.03 | 0.060 (0.019) | 0.032 | 0.047 | 0.057 | 0.070 | 0.109 |

*Note.* Additional details for the Unpaced tapping task are presented in Tables S26-S33.

* Significant age regression ($p < .05$). ≈[a] No difference between initial and final spontaneous conditions ($p > .05$). ≈[b] No difference between fast and slow conditions ($p > .05$).

***Table S5:*** *Norms for Paced Tapping to Tones and Music, Tapping Variability*

| Group | N (N female) | Skewness | Excess kurtosis | Mean (SD) | Percentiles 0 | 25 | 50 | 75 | 100 |
|---|---|---|---|---|---|---|---|---|---|
| **Paced tapping with tones – Variability (CV of ITI), fast tempo ‡** | | | | | | | | | |
| All | 107 (73) | 1.00 | 1.41 | 0.053 (0.015) | 0.029 | 0.042 | 0.051 | 0.060 | 0.109 |
| Age 18 to 21 | 27 (18) | 1.63 | 3.31 | 0.056 (0.015) | 0.038 | 0.046 | 0.054 | 0.061 | 0.109 |
| Age 22 to 29 | 29 (19) | 0.93 | 0.57 | 0.053 (0.014) | 0.031 | 0.046 | 0.052 | 0.060 | 0.091 |
| Age 30 to 54 | 26 (16) | 0.43 | -0.88 | 0.052 (0.015) | 0.029 | 0.038 | 0.049 | 0.065 | 0.086 |
| Age 55 to 87 | 25 (20) | 0.78 | 0.88 | 0.049 (0.013) | 0.032 | 0.037 | 0.049 | 0.057 | 0.085 |
| **Paced tapping with tones – Variability (CV of ITI), medium tempo ‡** | | | | | | | | | |
| All | 108 (74) | 0.62 | -0.01 | 0.046 (0.011) | 0.024 | 0.038 | 0.044 | 0.055 | 0.080 |
| Age 18 to 21 | 27 (18) | 0.90 | 1.14 | 0.048 (0.009) | 0.036 | 0.041 | 0.047 | 0.053 | 0.076 |
| Age 22 to 29 | 29 (19) | 0.62 | 0.14 | 0.046 (0.012) | 0.024 | 0.037 | 0.045 | 0.056 | 0.080 |
| Age 30 to 54 | 26 (16) | 0.63 | -0.64 | 0.047 (0.012) | 0.029 | 0.039 | 0.043 | 0.055 | 0.071 |
| Age 55 to 87 | 26 (21) | 0.51 | -0.85 | 0.044 (0.010) | 0.030 | 0.036 | 0.042 | 0.051 | 0.066 |
| **Paced tapping with tones – Variability (CV of ITI), slow tempo ‡** | | | | | | | | | |
| All | 107 (73) | 0.51 | -0.28 | 0.050 (0.013) | 0.027 | 0.039 | 0.049 | 0.059 | 0.089 |
| Age 18 to 21 | 27 (18) | 1.02 | 0.82 | 0.052 (0.013) | 0.033 | 0.044 | 0.047 | 0.058 | 0.089 |
| Age 22 to 29 | 29 (19) | 0.25 | -0.76 | 0.051 (0.012) | 0.034 | 0.039 | 0.050 | 0.059 | 0.077 |
| Age 30 to 54 | 26 (16) | 0.27 | -1.21 | 0.050 (0.013) | 0.031 | 0.040 | 0.049 | 0.063 | 0.072 |
| Age 55 to 87 | 25 (20) | 0.48 | -0.62 | 0.049 (0.014) | 0.027 | 0.038 | 0.049 | 0.059 | 0.079 |
| **Paced tapping with music – Variability (CV of ITI), music 1 ≈** | | | | | | | | | |
| All | 106 (72) | 3.34 | 17.42 | 0.055 (0.026) | 0.028 | 0.039 | 0.047 | 0.064 | 0.223 |
| Age 18 to 21 | 27 (18) | 0.69 | -0.75 | 0.054 (0.018) | 0.028 | 0.041 | 0.047 | 0.068 | 0.090 |
| Age 22 to 29 | 29 (19) | 0.19 | -1.01 | 0.055 (0.018) | 0.028 | 0.037 | 0.054 | 0.068 | 0.092 |

| | | | | | Percentiles | | | | |
|---|---|---|---|---|---|---|---|---|---|
| **Group** | **N (N female)** | **Skewness** | **Excess kurtosis** | **Mean (SD)** | **0** | **25** | **50** | **75** | **100** |
| Age 30 to 54 | 24 (14) | 2.78 | 7.30 | 0.059 (0.043) | 0.030 | 0.039 | 0.045 | 0.057 | 0.223 |
| Age 55 to 87 | 26 (21) | 1.38 | 2.03 | 0.052 (0.017) | 0.029 | 0.040 | 0.046 | 0.060 | 0.106 |
| **Paced tapping with music – Variability (CV of ITI), music 2 ≈** | | | | | | | | | |
| All | 102 (70) | 3.63 | 18.32 | 0.057 (0.031) | 0.026 | 0.039 | 0.051 | 0.065 | 0.257 |
| Age 18 to 21 | 24 (17) | 1.80 | 2.91 | 0.060 (0.026) | 0.031 | 0.044 | 0.055 | 0.068 | 0.137 |
| Age 22 to 29 | 28 (18) | 0.74 | -0.06 | 0.057 (0.018) | 0.034 | 0.043 | 0.054 | 0.069 | 0.102 |
| Age 30 to 54 | 25 (15) | 2.56 | 6.39 | 0.063 (0.052) | 0.026 | 0.034 | 0.044 | 0.079 | 0.257 |
| Age 55 to 87 | 25 (20) | 0.20 | -0.81 | 0.049 (0.011) | 0.030 | 0.039 | 0.050 | 0.054 | 0.070 |

*Note.* Additional details for the Paced tapping tasks are presented in Tables S34-S53.

‡ Significant difference among tempo conditions ($p < .05$). ≈ No difference between music conditions ($p > .05$).

***Table S6:*** *Norms for Paced Tapping to Tones and Music, Synchronization Consistency*

| Group | N (N female) | Skewness | Excess kurtosis | Mean (SD) | Percentiles 0 | 25 | 50 | 75 | 100 |
|---|---|---|---|---|---|---|---|---|---|
| **Paced tapping with tones – Consistency (logit of vector length), fast tempo ‡[a]** | | | | | | | | | |
| All * | 107 (73) | -1.66 | 3.82 | 2.77 (0.86) | -0.98 | 2.38 | 2.94 | 3.36 | 4.00 |
| Age 18 to 21 † | 27 (18) | -2.52 | 8.30 | 2.58 (0.87) | -0.98 | 2.28 | 2.75 | 3.07 | 3.52 |
| Age 22 to 29 | 29 (19) | -1.00 | 1.16 | 2.60 (0.93) | -0.25 | 2.08 | 2.85 | 3.22 | 4.00 |
| Age 30 to 54 | 26 (16) | -1.29 | 0.69 | 2.82 (1.01) | 0.25 | 2.60 | 3.16 | 3.52 | 3.97 |
| Age 55 to 87 | 25 (20) | -0.13 | -0.12 | 3.11 (0.43) | 2.09 | 2.78 | 3.16 | 3.39 | 3.94 |
| **Paced tapping with tones – Consistency (logit of vector length), medium tempo ‡[a]** | | | | | | | | | |
| All | 108 (74) | -1.23 | 1.76 | 2.97 (0.78) | 0.10 | 2.70 | 3.05 | 3.49 | 4.28 |
| Age 18 to 21 | 27 (18) | -1.23 | 1.36 | 2.93 (0.68) | 1.01 | 2.74 | 2.96 | 3.43 | 3.81 |
| Age 22 to 29 | 29 (19) | -0.96 | 0.93 | 2.98 (0.73) | 0.84 | 2.61 | 3.05 | 3.47 | 3.95 |
| Age 30 to 54 | 26 (16) | -0.96 | 0.49 | 2.78 (0.97) | 0.10 | 2.17 | 3.01 | 3.39 | 4.16 |
| Age 55 to 87 | 26 (21) | -1.73 | 4.79 | 3.19 (0.72) | 0.59 | 2.91 | 3.29 | 3.63 | 4.28 |
| **Paced tapping with tones – Consistency (logit of vector length), slow tempo ‡[a]** | | | | | | | | | |
| All | 107 (73) | -1.65 | 3.17 | 2.94 (0.95) | -0.78 | 2.73 | 3.10 | 3.52 | 4.41 |
| Age 18 to 21 | 27 (18) | -1.95 | 4.10 | 2.99 (0.89) | 0.12 | 2.77 | 3.14 | 3.53 | 4.29 |
| Age 22 to 29 | 29 (19) | -1.79 | 3.75 | 2.98 (0.84) | 0.05 | 2.83 | 3.08 | 3.43 | 4.07 |
| Age 30 to 54 | 26 (16) | -1.18 | 0.97 | 2.84 (1.00) | 0.02 | 2.52 | 3.18 | 3.46 | 4.18 |
| Age 55 to 87 | 25 (20) | -1.65 | 3.44 | 2.94 (1.10) | -0.78 | 2.76 | 3.08 | 3.58 | 4.41 |
| **Paced tapping with music – Consistency (logit of vector length), music 1 ‡[b]** | | | | | | | | | |
| All | 106 (72) | -1.30 | 0.74 | 2.78 (1.52) | -1.81 | 2.53 | 3.25 | 3.84 | 4.76 |
| Age 18 to 21 † | 27 (18) | -1.02 | 0.03 | 2.60 (1.70) | -1.81 | 1.34 | 3.11 | 3.86 | 4.76 |
| Age 22 to 29 | 29 (19) | -0.63 | -0.99 | 2.18 (1.80) | -1.03 | 0.78 | 2.84 | 3.40 | 4.42 |

| | | | | | Percentiles | | | | |
|---|---|---|---|---|---|---|---|---|---|
| **Group** | **N (N female)** | **Skewness** | **Excess kurtosis** | **Mean (SD)** | **0** | **25** | **50** | **75** | **100** |
| Age 30 to 54 | 24 (14) | -1.80 | 2.38 | 3.23 (1.37) | -0.97 | 3.10 | 3.78 | 3.99 | 4.44 |
| Age 55 to 87 | 26 (21) | -1.19 | 2.06 | 3.21 (0.68) | 1.07 | 2.96 | 3.29 | 3.61 | 4.15 |
| **Paced tapping with music – Consistency (logit of vector length), music 2 ‡[b]** | | | | | | | | | |
| All * | 102 (70) | -0.90 | 0.09 | 2.35 (1.71) | -2.74 | 1.46 | 2.76 | 3.77 | 4.59 |
| Age 18 to 21 † | 24 (17) | -0.49 | -0.88 | 1.92 (1.61) | -1.54 | 0.82 | 2.39 | 3.09 | 3.99 |
| Age 22 to 29 | 28 (18) | -0.42 | -0.83 | 1.62 (2.05) | -2.74 | 0.37 | 1.67 | 3.38 | 4.27 |
| Age 30 to 54 | 25 (15) | -0.98 | 0.40 | 2.96 (1.41) | -0.93 | 2.10 | 3.45 | 4.08 | 4.59 |
| Age 55 to 87 | 25 (20) | -1.60 | 3.71 | 2.98 (1.20) | -1.18 | 2.34 | 3.13 | 3.82 | 4.59 |

*Note.* Additional details for the Paced tapping tasks are presented in Tables S34-S53. Reference values for consistency (range for non-transformed vector length: 0-1) are: non-transformed 0.2 = -1.39 after logit transformation; 0.5 = 0.00; 0.75 = 1.10; 0.90 = 2.20; 0.95 = 2.94; 0.99 = 4.60.

* Significant age regression ($p < .05$). † Significant difference among age groups ($p < .05$). ‡[a] Significant difference among tempo conditions ($p < .05$). ‡[b] Significant difference between music conditions ($p < .05$).

***Table S7:*** *Norms for Paced Tapping to Tones and Music, Synchronization Accuracy*

| | | | | | Percentiles | | | | |
|---|---|---|---|---|---|---|---|---|---|
| **Group** | **N (N female)** | **Skewness** | **Excess kurtosis** | **Mean (SD)** | **0** | **25** | **50** | **75** | **100** |
| **Paced tapping with tones – Accuracy (vector direction %), fast tempo ‡[a]** | | | | | | | | | |
| All | 97 (65) | -0.43 | -0.23 | -7.1 (5.9) | -24.6 | -10.6 | -6.8 | -2.8 | 3.2 |
| Age 18 to 21 | 23 (15) | -1.02 | 0.94 | -7.9 (5.9) | -24.6 | -9.8 | -6.8 | -4.6 | 1.1 |
| Age 22 to 29 | 25 (15) | -0.14 | -0.99 | -7.5 (6.1) | -18.8 | -12.3 | -7.9 | -3.4 | 3.0 |
| Age 30 to 54 | 24 (15) | 0.23 | -0.48 | -7.4 (5.0) | -17.4 | -10.9 | -7.8 | -4.4 | 3.2 |
| Age 55 to 87 | 25 (20) | -0.70 | -0.37 | -5.8 (6.4) | -21.4 | -10.1 | -4.2 | -0.1 | 2.4 |
| **Paced tapping with tones – Accuracy (vector direction %), medium tempo ‡[a]** | | | | | | | | | |
| All | 106 (73) | -0.94 | 0.58 | -6.5 (5.6) | -24.9 | -9.9 | -5.5 | -2.4 | 3.2 |
| Age 18 to 21 | 27 (18) | -0.61 | -0.66 | -7.3 (5.3) | -19.1 | -10.7 | -6.1 | -2.8 | 0.8 |
| Age 22 to 29 | 29 (19) | -0.82 | -0.15 | -6.4 (5.7) | -20.7 | -10.1 | -4.6 | -2.5 | 1.8 |
| Age 30 to 54 | 24 (15) | -1.14 | 0.82 | -6.4 (5.4) | -21.7 | -9.0 | -5.4 | -1.9 | -0.6 |
| Age 55 to 87 | 26 (21) | -1.22 | 1.98 | -6.1 (6.1) | -24.9 | -8.9 | -5.2 | -1.6 | 3.2 |
| **Paced tapping with tones – Accuracy (vector direction %), slow tempo ‡[a]** | | | | | | | | | |
| All | 104 (70) | -0.36 | 0.02 | -5.3 (4.7) | -17.6 | -8.3 | -4.6 | -2.1 | 5.8 |
| Age 18 to 21 | 27 (18) | -0.85 | -0.05 | -6.3 (5.1) | -17.6 | -9.0 | -4.9 | -2.5 | 1.2 |
| Age 22 to 29 | 28 (18) | 0.20 | -0.56 | -3.8 (3.8) | -10.8 | -6.2 | -3.6 | -1.5 | 4.6 |
| Age 30 to 54 | 24 (14) | -0.29 | -0.15 | -3.9 (5.1) | -14.9 | -6.4 | -3.9 | -1.1 | 5.8 |
| Age 55 to 87 | 25 (20) | -0.13 | -1.07 | -7.0 (4.3) | -14.6 | -10.5 | -7.1 | -4.0 | -0.4 |
| **Paced tapping with music – Accuracy (vector direction %), music 1 ‡[b]** | | | | | | | | | |
| All | 99 (70) | 0.54 | 1.54 | 0.3 (5.4) | -15.5 | -2.6 | -0.4 | 3.0 | 16.2 |
| Age 18 to 21 | 25 (17) | -0.11 | 2.56 | 0.5 (5.7) | -15.5 | -0.9 | 0.5 | 2.2 | 16.1 |
| Age 22 to 29 | 25 (18) | 0.45 | 0.01 | 0.2 (6.6) | -11.0 | -4.5 | 0.4 | 3.3 | 16.2 |

| | | | | | Percentiles | | | | |
|---|---|---|---|---|---|---|---|---|---|
| **Group** | **N (N female)** | **Skewness** | **Excess kurtosis** | **Mean (SD)** | **0** | **25** | **50** | **75** | **100** |
| Age 30 to 54 | 23 (14) | 1.66 | 2.79 | 0.2 (5.2) | -7.4 | -2.5 | -0.9 | 0.9 | 16.2 |
| Age 55 to 87 | 26 (21) | 0.39 | -0.17 | 0.3 (4.1) | -7.8 | -2.4 | 0.3 | 2.2 | 9.5 |
| **Paced tapping with music – Accuracy (vector direction %), music 2 ‡[b]** | | | | | | | | | |
| All | 90 (62) | -0.66 | 0.77 | -3.4 (6.6) | -25.0 | -6.7 | -2.4 | 0.4 | 10.5 |
| Age 18 to 21 | 19 (13) | -1.10 | 1.83 | -4.1 (7.4) | -25.0 | -6.8 | -2.3 | 0.2 | 8.8 |
| Age 22 to 29 | 23 (15) | -0.48 | -0.52 | -4.0 (7.6) | -18.4 | -7.3 | -2.2 | 0.9 | 7.6 |
| Age 30 to 54 | 24 (15) | -0.35 | 0.08 | -2.0 (6.6) | -17.9 | -5.1 | -1.5 | 1.9 | 10.5 |
| Age 55 to 87 | 24 (19) | -0.69 | 0.15 | -3.8 (4.9) | -15.8 | -6.5 | -2.8 | -0.8 | 3.9 |

*Note.* Additional details for the Paced tapping tasks are presented in Tables S34-S43.

‡[a] Significant difference among tempo conditions ($p < .05$). ‡[b] Significant difference between music conditions ($p < .05$).

***Table S8:*** *Norms for Synchronization-Continuation, Tapping Rate and Variability*

| | | | | | Percentiles | | | | |
|---|---|---|---|---|---|---|---|---|---|
| **Group** | **N (N female)** | **Skewness** | **Excess kurtosis** | **Mean (SD)** | **0** | **25** | **50** | **75** | **100** |
| **Synchronization-continuation – Rate (ITI in ms), fast tempo ‡[a]** | | | | | | | | | |
| All * | 108 (74) | 0.17 | 4.10 | 448.8 (21.6) | 374.6 | 437.5 | 449.4 | 459.8 | 545.7 |
| Age 18 to 21 | 27 (18) | 2.09 | 6.26 | 451.1 (24.6) | 417.4 | 436.5 | 447.3 | 459.1 | 545.7 |
| Age 22 to 29 | 29 (19) | -1.02 | 0.88 | 443.4 (21.0) | 387.2 | 433.0 | 449.8 | 456.3 | 477.9 |
| Age 30 to 54 | 26 (16) | -1.32 | 2.79 | 444.2 (21.1) | 374.6 | 435.8 | 447.6 | 459.3 | 480.6 |
| Age 55 to 87 | 26 (21) | 0.13 | -0.98 | 457.3 (17.0) | 428.8 | 444.7 | 456.0 | 468.0 | 485.0 |
| **Synchronization-continuation – Rate (ITI in ms), medium tempo ‡[a]** | | | | | | | | | |
| All | 108 (74) | 0.43 | 3.74 | 596.5 (26.1) | 511.7 | 581.1 | 599.7 | 606.4 | 713.1 |
| Age 18 to 21 | 27 (18) | 2.32 | 7.14 | 597.7 (29.2) | 564.4 | 578.8 | 599.5 | 604.8 | 713.1 |
| Age 22 to 29 | 29 (19) | -0.80 | 0.46 | 592.7 (31.7) | 511.7 | 589.9 | 602.8 | 608.7 | 658.4 |
| Age 30 to 54 | 26 (16) | 0.04 | -0.34 | 594.5 (19.7) | 553.7 | 580.6 | 595.3 | 607.6 | 638.6 |
| Age 55 to 87 | 26 (21) | 0.65 | -0.03 | 601.6 (21.3) | 563.0 | 588.8 | 595.9 | 615.1 | 652.6 |
| **Synchronization-continuation – Rate (ITI in ms), slow tempo ‡[a]** | | | | | | | | | |
| All | 108 (74) | -0.28 | 0.61 | 749.3 (40.2) | 609.0 | 720.2 | 750.5 | 776.4 | 849.4 |
| Age 18 to 21 | 27 (18) | 0.21 | -0.25 | 761.2 (36.3) | 687.9 | 740.0 | 760.3 | 786.9 | 837.6 |
| Age 22 to 29 | 29 (19) | -0.53 | -0.55 | 749.2 (37.0) | 672.1 | 729.8 | 755.6 | 772.8 | 808.3 |
| Age 30 to 54 | 26 (16) | -0.31 | 0.27 | 742.7 (52.9) | 609.0 | 709.6 | 750.4 | 768.3 | 849.4 |
| Age 55 to 87 | 26 (21) | 0.34 | -0.92 | 743.9 (31.4) | 685.4 | 719.5 | 735.0 | 770.7 | 802.0 |
| **Synchronization-continuation – Variability (CV of ITI), fast tempo ‡[b]** | | | | | | | | | |
| All | 108 (74) | 1.00 | 0.58 | 0.048 (0.015) | 0.025 | 0.037 | 0.046 | 0.054 | 0.092 |
| Age 18 to 21 | 27 (18) | 0.50 | -0.32 | 0.051 (0.016) | 0.025 | 0.041 | 0.048 | 0.062 | 0.087 |
| Age 22 to 29 | 29 (19) | 1.13 | -0.04 | 0.052 (0.018) | 0.032 | 0.039 | 0.047 | 0.054 | 0.092 |

| | | | | | Percentiles | | | | |
|---|---|---|---|---|---|---|---|---|---|
| **Group** | **N (N female)** | **Skewness** | **Excess kurtosis** | **Mean (SD)** | **0** | **25** | **50** | **75** | **100** |
| Age 30 to 54 | 26 (16) | 0.91 | 0.16 | 0.044 (0.013) | 0.028 | 0.035 | 0.042 | 0.050 | 0.075 |
| Age 55 to 87 | 26 (21) | 0.35 | -0.77 | 0.045 (0.012) | 0.026 | 0.035 | 0.044 | 0.053 | 0.071 |
| **Synchronization-continuation – Variability (CV of ITI), medium tempo ‡[b]** | | | | | | | | | |
| All | 108 (74) | 1.05 | 1.16 | 0.046 (0.014) | 0.025 | 0.035 | 0.044 | 0.054 | 0.094 |
| Age 18 to 21 | 27 (18) | 1.49 | 2.88 | 0.048 (0.014) | 0.030 | 0.039 | 0.046 | 0.054 | 0.094 |
| Age 22 to 29 | 29 (19) | 0.94 | 0.83 | 0.046 (0.015) | 0.025 | 0.035 | 0.044 | 0.056 | 0.090 |
| Age 30 to 54 | 26 (16) | 0.50 | -0.97 | 0.046 (0.014) | 0.028 | 0.035 | 0.041 | 0.058 | 0.075 |
| Age 55 to 87 | 26 (21) | 1.41 | 2.31 | 0.042 (0.012) | 0.027 | 0.033 | 0.039 | 0.048 | 0.081 |
| **Synchronization-continuation – Variability (CV of ITI), slow tempo ‡[b]** | | | | | | | | | |
| All * | 108 (74) | 1.22 | 2.59 | 0.048 (0.014) | 0.024 | 0.039 | 0.046 | 0.054 | 0.106 |
| Age 18 to 21 | 27 (18) | 1.60 | 2.78 | 0.053 (0.017) | 0.028 | 0.044 | 0.049 | 0.057 | 0.106 |
| Age 22 to 29 | 29 (19) | 0.60 | -0.03 | 0.048 (0.012) | 0.026 | 0.040 | 0.047 | 0.054 | 0.076 |
| Age 30 to 54 | 26 (16) | 0.48 | -0.03 | 0.049 (0.015) | 0.024 | 0.039 | 0.049 | 0.054 | 0.085 |
| Age 55 to 87 | 26 (21) | 0.43 | -1.18 | 0.042 (0.009) | 0.031 | 0.035 | 0.039 | 0.050 | 0.058 |

*Note.* Additional details for the Synchronization-continuation task are presented in Tables S54-S59.

* Significant age regression ($p < .05$). ‡[a] Significant difference of rate among tempo conditions ($p < .05$). ‡[b] Significant difference of variability among tempo conditions ($p < .05$).

***Table S9:*** *Norms for Adaptive Tapping, Sensitivity Index and Tapping Variability*

| Group | N (N female) | Skewness | Excess kurtosis | Mean (SD) | Percentiles 0 | 25 | 50 | 75 | 100 |
|---|---|---|---|---|---|---|---|---|---|
| **Adaptive tapping – Sensitivity index (d') of perceiving tempo deceleration (IOI + 75ms)** | | | | | | | | | |
| All | 105 (72) | -1.74 | 2.72 | 3.63 (0.74) | 0.82 | 3.40 | 4.14 | 4.14 | 4.14 |
| Age 18 to 21 | 26 (17) | -1.06 | 0.41 | 3.47 (0.74) | 1.45 | 3.00 | 3.78 | 4.14 | 4.14 |
| Age 22 to 29 | 29 (19) | -1.87 | 2.25 | 3.68 (0.86) | 1.25 | 3.78 | 4.14 | 4.14 | 4.14 |
| Age 30 to 54 | 26 (16) | -2.47 | 6.50 | 3.70 (0.75) | 0.82 | 3.62 | 4.14 | 4.14 | 4.14 |
| Age 55 to 87 | 24 (20) | -1.05 | 0.06 | 3.66 (0.58) | 2.22 | 3.37 | 3.78 | 4.14 | 4.14 |
| **Adaptive tapping – Sensitivity index (d') of perceiving tempo deceleration (IOI + 30ms)** | | | | | | | | | |
| All | 105 (72) | 0.23 | -0.02 | 1.84 (0.98) | -0.36 | 1.27 | 1.83 | 2.53 | 4.14 |
| Age 18 to 21 | 26 (17) | 0.40 | -0.64 | 1.69 (1.19) | -0.36 | 0.84 | 1.57 | 2.16 | 4.14 |
| Age 22 to 29 | 29 (19) | 0.69 | 0.94 | 1.75 (0.83) | 0.30 | 1.27 | 1.71 | 2.22 | 4.14 |
| Age 30 to 54 | 26 (16) | -0.79 | 0.62 | 2.01 (0.88) | -0.23 | 1.63 | 2.07 | 2.56 | 3.27 |
| Age 55 to 87 | 24 (20) | 0.54 | 0.11 | 1.93 (1.01) | 0.00 | 1.25 | 1.78 | 2.54 | 4.14 |
| **Adaptive tapping – Sensitivity index (d') of perceiving tempo acceleration (IOI - 75ms)** | | | | | | | | | |
| All | 105 (72) | -2.21 | 6.27 | 3.79 (0.54) | 1.04 | 3.57 | 4.14 | 4.14 | 4.14 |
| Age 18 to 21 | 26 (17) | -1.56 | 1.77 | 3.75 (0.55) | 2.09 | 3.62 | 3.96 | 4.14 | 4.14 |
| Age 22 to 29 | 29 (19) | -1.65 | 1.38 | 3.92 (0.39) | 2.91 | 3.78 | 4.14 | 4.14 | 4.14 |
| Age 30 to 54 | 26 (16) | -2.01 | 4.09 | 3.61 (0.74) | 1.04 | 3.34 | 3.78 | 4.14 | 4.14 |
| Age 55 to 87 | 24 (20) | -1.20 | 0.26 | 3.86 (0.39) | 2.96 | 3.78 | 4.14 | 4.14 | 4.14 |
| **Adaptive tapping – Sensitivity index (d') perceiving a tempo acceleration (IOI - 30ms)** | | | | | | | | | |
| All | 105 (72) | -0.09 | -0.07 | 1.69 (0.88) | -0.36 | 1.21 | 1.83 | 2.31 | 4.14 |
| Age 18 to 21 | 26 (17) | 0.76 | 0.56 | 1.54 (0.83) | 0.00 | 0.88 | 1.42 | 1.80 | 3.78 |
| Age 22 to 29 | 29 (19) | 0.09 | 0.18 | 1.79 (0.97) | -0.36 | 1.21 | 1.83 | 2.31 | 4.14 |

| Group | N (N female) | Skewness | Excess kurtosis | Mean (SD) | Percentiles 0 | 25 | 50 | 75 | 100 |
|---|---|---|---|---|---|---|---|---|---|
| Age 30 to 54 | 26 (16) | -0.44 | -0.94 | 1.56 (0.92) | 0.00 | 1.00 | 1.70 | 2.28 | 2.87 |
| Age 55 to 87 | 24 (20) | -1.04 | 0.57 | 1.86 (0.78) | 0.00 | 1.54 | 2.07 | 2.31 | 2.96 |
| **Adaptive tapping – Variability (CV of ITI) of continuation tapping (isochronous condition)** | | | | | | | | | |
| All | 105 (72) | 2.39 | 8.42 | 0.057 (0.020) | 0.031 | 0.044 | 0.052 | 0.064 | 0.166 |
| Age 18 to 21 | 26 (17) | 1.12 | 1.03 | 0.058 (0.015) | 0.035 | 0.047 | 0.054 | 0.065 | 0.099 |
| Age 22 to 29 | 29 (19) | 1.70 | 3.10 | 0.059 (0.019) | 0.037 | 0.046 | 0.054 | 0.065 | 0.122 |
| Age 30 to 54 | 26 (16) | 2.07 | 4.52 | 0.062 (0.029) | 0.033 | 0.044 | 0.050 | 0.069 | 0.166 |
| Age 55 to 87 | 24 (20) | 0.22 | -0.28 | 0.049 (0.010) | 0.031 | 0.043 | 0.050 | 0.055 | 0.072 |
| **Adaptive tapping – Adaptation index (acceleration trials)** | | | | | | | | | |
| All | 105 (72) | 2.18 | 8.00 | 1.442 (0.597) | 0.403 | 1.120 | 1.355 | 1.624 | 4.547 |
| Age 18 to 21 | 26 (17) | 0.17 | -0.46 | 1.432 (0.417) | 0.767 | 1.107 | 1.427 | 1.689 | 2.404 |
| Age 22 to 29 | 29 (19) | 2.05 | 5.37 | 1.407 (0.655) | 0.403 | 1.132 | 1.236 | 1.502 | 3.880 |
| Age 30 to 54 | 26 (16) | 2.47 | 7.69 | 1.495 (0.770) | 0.580 | 1.063 | 1.335 | 1.631 | 4.547 |
| Age 55 to 87 | 24 (20) | 1.20 | 1.55 | 1.439 (0.501) | 0.784 | 1.143 | 1.375 | 1.534 | 2.889 |
| **Adaptive tapping – Adaptation index (deceleration trials)** | | | | | | | | | |
| All * | 105 (72) | -0.36 | 5.15 | 1.263 (0.522) | -1.169 | 1.009 | 1.256 | 1.565 | 3.334 |
| Age 18 to 21 | 26 (17) | 0.54 | 0.42 | 1.390 (0.338) | 0.826 | 1.187 | 1.367 | 1.561 | 2.287 |
| Age 22 to 29 | 29 (19) | -2.18 | 6.33 | 1.186 (0.599) | -1.169 | 1.060 | 1.268 | 1.490 | 1.910 |
| Age 30 to 54 | 26 (16) | 0.92 | 1.40 | 1.347 (0.660) | 0.309 | 1.006 | 1.209 | 1.682 | 3.334 |
| Age 55 to 87 | 24 (20) | 0.34 | -0.89 | 1.128 (0.381) | 0.536 | 0.809 | 1.098 | 1.500 | 1.936 |

*Note.* Additional details for the Adaptive tapping task are presented in Tables S60-S66.

* Significant age regression ($p < .05$).

***Table S10:*** *Norms for Beat Tracking Index*

| | | | | | Percentiles | | | | |
|---|---|---|---|---|---|---|---|---|---|
| **Group** | **N (N female)** | **Skewness** | **Excess kurtosis** | **Mean (SD)** | **0** | **25** | **50** | **75** | **100** |
| | | | **BTI – Beat Tracking Index** | | | | | | |
| All | 101 (69) | -0.56 | 0.19 | -0.001 (0.801) | -2.190 | -0.501 | 0.090 | 0.612 | 1.406 |
| Age 18 to 21 | 24 (17) | -0.79 | 0.94 | 0.040 (0.766) | -2.146 | -0.336 | 0.141 | 0.511 | 1.193 |
| Age 22 to 29 | 28 (18) | -0.29 | -0.68 | -0.174 (0.985) | -2.190 | -0.775 | -0.145 | 0.718 | 1.276 |
| Age 30 to 54 | 24 (14) | -0.71 | 0.29 | 0.129 (0.798) | -1.805 | -0.198 | 0.193 | 0.624 | 1.406 |
| Age 55 to 87 | 25 (20) | 0.06 | -0.32 | 0.028 (0.594) | -1.154 | -0.252 | -0.004 | 0.460 | 1.358 |

*Note.* Additional details for the Beat Tracking Index are presented in Table S67.
