## Supplementary material for "Mobile version of the Battery for the Assessment of Auditory Sensorimotor and Timing Abilities (BAASTA): Implementation and adult norms": Annex 2 - full norm tables

##### Annex 2: Norm full tables

##### Table of Contents

Full tables by variable ... 5

**Table S11:** Duration discrimination – Threshold (% of IOI) – DurDisc_threshold ... 5

**Table S12:** Anisochrony detection with tones – Threshold (% of IOI) – Anisochrony_tones_threshold ... 8

**Table S13:** Anisochrony detection with music – Threshold (% of IOI) – Anisochrony_music_threshold ... 11

**Table S14:** Beat Alignment Test – Sensitivity index (d') for all trials – BAT_all_dprime ... 14

**Table S15:** Beat Alignment Test – Sensitivity index (d') for fast tempo trials – BAT_fast_dprime ... 17

**Table S16:** Beat Alignment Test – Sensitivity index (d') for medium tempo trials – BAT_med_dprime ... 20

**Table S17:** Beat Alignment Test – Sensitivity index (d') for slow tempo trials – BAT_slow_dprime ... 23

**Table S18:** Beat Alignment Test – Hit rate for all trials – BAT_all_hits ... 26

**Table S19:** Beat Alignment Test – Hit rate for fast tempo trials – BAT_fast_hits ... 29

**Table S20:** Beat Alignment Test – Hit rate for medium tempo trials – BAT_med_hits ... 32

**Table S21:** Beat Alignment Test – Hit rate for slow tempo trials – BAT_slow_hits ... 35

**Table S22:** Beat Alignment Test – False alarm rate for all trials trials – BAT_all_fa ... 38

**Table S23:** Beat Alignment Test – False alarm rate for fast tempo trials – BAT_fast_fa ... 41

**Table S24:** Beat Alignment Test – False alarm rate for medium tempo trials – BAT_med_fa44

**Table S25:** Beat Alignment Test – False alarm rate for slow tempo trials – BAT_slow_fa ... 47

**Table S26:** Unpaced tapping – Rate (ITI in ms) of spontaneous tapping, initial trial – Unpaced_spontaneous_right_mean_iti ... 50

**Table S27:** Unpaced tapping – Rate (ITI in ms) of spontaneous tapping, final trial – Unpaced_spontaneous_final_right_mean_iti ... 53

**Table S28:** Unpaced tapping – Rate (ITI in ms) of fast tapping – Unpaced_fast_mean_iti ... 56

**Table S29:** Unpaced tapping – Rate (ITI in ms) of slow tapping – Unpaced_slow_mean_iti. 59

### Full tables by variable

***Table S11:*** *Duration discrimination – Threshold (% of IOI) – DurDisc_threshold*

| | | | | | | |
|---|---|---|---|---|---|---|
| **Task** | Duration discrimination | | | | | |
| **Outcome measure** | Threshold (% of IOI) | | | | | |
| **Variable name** | DurDisc_threshold | | | | | |
| **Age and gender effects** | | | | | | |
| Age regression (slope) | | 0.078 | | | | |
| Age regression (p) | | 0.237 | | | | |
| Age group (p) | | 0.738 | | | | |
| Gender (p) | | 0.479 | | | | |
| **Group** | | All | Age 18 to 21 | Age 22 to 29 | Age 30 to 54 | Age 55 to 87 |
| N (N female) | | 101 (70) | 25 (16) | 29 (19) | 25 (16) | 22 (19) |
| **Normality** | | | | | | |
| Skewness | | 0.89 | 1.21 | 0.56 | 0.95 | -0.11 |
| Excess kurtosis | | 0.71 | 2.81 | -0.05 | -0.14 | -1.06 |
| **Scores** | | | | | | |
| Mean | | 29.1 | 26.4 | 29.2 | 30.9 | 29.9 |
| SD | | 12.7 | 9.7 | 11.5 | 17.0 | 12.0 |
| Percentiles | | | | | | |
| | 0 | 7.6 | 10.3 | 11.8 | 8.5 | 7.6 |
| | 1 | 8.5 | 11.2 | 12.4 | 9.7 | 8.7 |
| | 2 | 10.3 | 12.1 | 13.0 | 10.8 | 9.8 |
| | 3 | 11.8 | 12.9 | 13.7 | 11.9 | 10.9 |
| | 4 | 12.9 | 13.8 | 14.2 | 13.1 | 12.0 |
| | 5 | 13.3 | 14.0 | 14.5 | 13.3 | 13.0 |
| | 6 | 13.5 | 14.0 | 14.9 | 13.4 | 13.5 |
| | 7 | 13.9 | 14.1 | 15.3 | 13.4 | 13.9 |
| | 8 | 14.0 | 14.1 | 15.4 | 13.5 | 14.4 |
| | 9 | 14.1 | 14.8 | 15.4 | 13.9 | 14.9 |
| | 10 | 15.1 | 15.9 | 15.4 | 14.5 | 15.2 |
| | 11 | 15.4 | 16.9 | 15.4 | 15.2 | 15.5 |
| | 12 | 15.4 | 17.9 | 15.5 | 15.8 | 15.8 |
| | 13 | 15.8 | 18.4 | 15.6 | 16.2 | 16.0 |
| | 14 | 16.0 | 18.4 | 15.8 | 16.3 | 16.3 |
| | 15 | 16.1 | 18.5 | 15.8 | 16.5 | 16.6 |
| | 16 | 16.4 | 18.5 | 15.9 | 16.7 | 17.0 |
| | 17 | 16.8 | 18.5 | 16.0 | 16.8 | 17.4 |
| | 18 | 17.6 | 18.7 | 16.2 | 17.0 | 17.8 |
| | 19 | 18.2 | 18.9 | 17.0 | 17.2 | 18.2 |
| | 20 | 18.4 | 19.2 | 17.9 | 17.4 | 18.4 |

| | | | | | |
|---|---|---|---|---|---|
| 21 | 18.4 | 19.3 | 18.8 | 17.6 | 18.7 |
| 22 | 18.5 | 19.4 | 19.4 | 17.8 | 18.9 |
| 23 | 18.7 | 19.5 | 19.8 | 18.0 | 19.1 |
| 24 | 19.2 | 19.6 | 20.2 | 18.2 | 19.4 |
| 25 | 19.3 | 19.7 | 20.7 | 18.4 | 19.8 |
| 26 | 19.3 | 20.0 | 21.0 | 18.5 | 20.2 |
| 27 | 19.7 | 20.3 | 21.3 | 18.6 | 20.7 |
| 28 | 20.7 | 20.6 | 21.5 | 18.6 | 21.1 |
| 29 | 20.9 | 20.9 | 21.7 | 18.7 | 21.4 |
| 30 | 21.2 | 21.3 | 21.8 | 19.2 | 21.4 |
| 31 | 21.3 | 21.8 | 21.9 | 19.8 | 21.5 |
| 32 | 21.7 | 22.2 | 21.9 | 20.4 | 21.6 |
| 33 | 21.7 | 22.7 | 22.2 | 21.0 | 21.7 |
| 34 | 22.0 | 23.2 | 22.6 | 21.4 | 22.4 |
| 35 | 22.3 | 23.8 | 22.9 | 21.6 | 23.4 |
| 36 | 22.9 | 24.3 | 23.3 | 21.9 | 24.4 |
| 37 | 23.1 | 24.8 | 23.8 | 22.2 | 25.4 |
| 38 | 25.1 | 25.1 | 24.4 | 22.7 | 26.3 |
| 39 | 25.1 | 25.1 | 24.9 | 23.5 | 26.5 |
| 40 | 25.2 | 25.1 | 25.3 | 24.3 | 26.6 |
| 41 | 25.6 | 25.1 | 25.5 | 25.0 | 26.7 |
| 42 | 25.7 | 25.2 | 25.8 | 25.6 | 26.8 |
| 43 | 25.7 | 25.3 | 26.2 | 25.6 | 27.1 |
| 44 | 26.0 | 25.5 | 27.0 | 25.6 | 28.1 |
| 45 | 26.0 | 25.6 | 27.8 | 25.7 | 29.2 |
| 46 | 26.1 | 25.8 | 28.6 | 25.7 | 30.2 |
| 47 | 26.3 | 25.8 | 29.1 | 26.0 | 31.3 |
| 48 | 26.4 | 25.9 | 29.3 | 26.2 | 32.0 |
| 49 | 26.7 | 25.9 | 29.5 | 26.5 | 32.2 |
| 50 | 26.9 | 26.0 | 29.8 | 26.7 | 32.4 |
| 51 | 27.2 | 26.0 | 29.9 | 26.9 | 32.5 |
| 52 | 27.4 | 26.1 | 30.1 | 27.0 | 32.7 |
| 53 | 27.8 | 26.1 | 30.2 | 27.2 | 33.0 |
| 54 | 28.9 | 26.1 | 30.4 | 27.4 | 33.5 |
| 55 | 28.9 | 26.2 | 30.7 | 27.7 | 33.9 |
| 56 | 29.6 | 26.2 | 31.0 | 28.1 | 34.4 |
| 57 | 29.8 | 26.3 | 31.3 | 28.4 | 34.8 |
| 58 | 30.2 | 26.3 | 31.5 | 28.8 | 34.9 |
| 59 | 30.2 | 26.5 | 31.6 | 29.0 | 34.9 |
| 60 | 30.3 | 26.7 | 31.8 | 29.2 | 34.9 |
| 61 | 30.4 | 26.9 | 32.0 | 29.4 | 35.0 |
| 62 | 30.8 | 27.1 | 32.2 | 29.5 | 35.0 |
| 63 | 31.3 | 27.2 | 32.5 | 29.7 | 35.1 |
| 64 | 32.0 | 27.4 | 32.7 | 29.8 | 35.2 |

| | | | | | |
|---|---|---|---|---|---|
| 65 | 32.0 | 27.5 | 33.1 | 30.0 | 35.3 |
| 66 | 32.6 | 27.7 | 33.7 | 30.1 | 35.3 |
| 67 | 32.7 | 28.0 | 34.2 | 30.2 | 35.6 |
| 68 | 32.7 | 28.5 | 34.7 | 30.4 | 36.3 |
| 69 | 32.7 | 29.1 | 34.7 | 30.5 | 37.0 |
| 70 | 33.6 | 29.7 | 34.7 | 30.6 | 37.6 |
| 71 | 34.7 | 30.2 | 34.7 | 31.3 | 38.3 |
| 72 | 34.7 | 30.3 | 34.9 | 34.2 | 38.6 |
| 73 | 34.9 | 30.3 | 35.1 | 37.2 | 38.7 |
| 74 | 35.0 | 30.3 | 35.3 | 40.1 | 38.8 |
| 75 | 35.4 | 30.4 | 35.6 | 43.1 | 38.9 |
| 76 | 35.6 | 30.9 | 35.8 | 43.2 | 39.0 |
| 77 | 36.4 | 31.4 | 36.0 | 43.3 | 39.2 |
| 78 | 36.4 | 32.0 | 36.3 | 43.4 | 39.3 |
| 79 | 38.3 | 32.5 | 36.6 | 43.5 | 39.4 |
| 80 | 38.5 | 32.6 | 37.2 | 44.5 | 39.5 |
| 81 | 38.6 | 32.6 | 37.7 | 45.7 | 39.7 |
| 82 | 38.7 | 32.7 | 38.2 | 46.9 | 39.9 |
| 83 | 39.1 | 32.7 | 38.4 | 48.2 | 40.1 |
| 84 | 39.7 | 32.8 | 38.5 | 48.6 | 40.3 |
| 85 | 40.6 | 33.1 | 38.6 | 48.6 | 40.5 |
| 86 | 41.6 | 33.3 | 38.9 | 48.6 | 41.0 |
| 87 | 43.1 | 33.5 | 39.7 | 48.6 | 42.3 |
| 88 | 43.5 | 33.9 | 40.6 | 50.5 | 43.6 |
| 89 | 46.8 | 34.6 | 41.4 | 54.4 | 44.9 |
| 90 | 47.4 | 35.3 | 42.8 | 58.3 | 46.2 |
| 91 | 48.6 | 36.0 | 44.4 | 62.2 | 47.0 |
| 92 | 48.6 | 36.6 | 46.0 | 64.9 | 47.4 |
| 93 | 48.6 | 37.1 | 47.4 | 65.1 | 47.7 |
| 94 | 48.6 | 37.6 | 47.7 | 65.4 | 48.1 |
| 95 | 48.6 | 38.1 | 48.1 | 65.6 | 48.5 |
| 96 | 58.1 | 39.3 | 48.4 | 65.9 | 48.6 |
| 97 | 59.3 | 44.0 | 50.3 | 66.3 | 48.6 |
| 98 | 64.8 | 48.7 | 53.3 | 66.7 | 48.6 |
| 99 | 65.8 | 53.4 | 56.3 | 67.2 | 48.6 |
| 100 | 67.6 | 58.1 | 59.3 | 67.6 | 48.6 |

***Table S12:*** *Anisochrony detection with tones – Threshold (% of IOI) – Anisochrony_tones_threshold*

| | | | | | | |
|---|---|---|---|---|---|---|
| **Task** | | Anisochrony detection with tones | | | | |
| **Outcome measure** | | Threshold (% of IOI) | | | | |
| **Variable name** | | Anisochrony_tones_threshold | | | | |
| **Age and gender effects** | | | | | | |
| Age regression (slope) | | 0.066 | | | | |
| Age regression (p) | | 0.015 | | | | |
| Age group (p) | | 0.018 | | | | |
| Gender (p) | | 0.577 | | | | |
| **Group** | | All | Age 18 to 21 | Age 22 to 29 | Age 30 to 54 | Age 55 to 87 |
| N (N female) | | 100 (69) | 26 (18) | 26 (16) | 23 (15) | 25 (20) |
| **Normality** | | | | | | |
| Skewness | | 0.41 | 0.47 | 0.41 | 0.37 | -0.01 |
| Excess kurtosis | | -0.69 | -0.74 | -0.85 | -0.88 | -0.87 |
| **Scores** | | | | | | |
| Mean | | 12.4 | 12.2 | 10.9 | 11.4 | 15.2 |
| SD | | 5.1 | 5.5 | 4.3 | 4.3 | 5.3 |
| Percentiles | | | | | | |
| | 0 | 3.8 | 3.8 | 4.7 | 5.3 | 4.4 |
| | 1 | 4.4 | 4.0 | 4.8 | 5.5 | 5.3 |
| | 2 | 4.7 | 4.3 | 4.9 | 5.6 | 6.1 |
| | 3 | 4.7 | 4.5 | 4.9 | 5.8 | 6.9 |
| | 4 | 5.0 | 4.7 | 5.0 | 5.9 | 7.7 |
| | 5 | 5.3 | 4.9 | 5.1 | 6.0 | 8.1 |
| | 6 | 5.5 | 5.1 | 5.2 | 6.1 | 8.4 |
| | 7 | 5.5 | 5.3 | 5.3 | 6.2 | 8.7 |
| | 8 | 5.5 | 5.5 | 5.5 | 6.3 | 9.0 |
| | 9 | 6.0 | 5.8 | 5.5 | 6.4 | 9.2 |
| | 10 | 6.3 | 6.1 | 5.5 | 6.4 | 9.4 |
| | 11 | 6.5 | 6.4 | 5.5 | 6.4 | 9.6 |
| | 12 | 6.5 | 6.7 | 5.5 | 6.4 | 9.8 |
| | 13 | 6.7 | 6.9 | 5.7 | 6.5 | 9.9 |
| | 14 | 6.9 | 7.2 | 6.0 | 6.5 | 9.9 |
| | 15 | 7.2 | 7.4 | 6.3 | 6.6 | 9.9 |
| | 16 | 7.3 | 7.6 | 6.5 | 6.7 | 10.0 |
| | 17 | 7.4 | 7.7 | 6.7 | 6.8 | 10.1 |
| | 18 | 7.6 | 7.7 | 6.9 | 6.9 | 10.2 |
| | 19 | 7.8 | 7.8 | 7.0 | 7.0 | 10.4 |
| | 20 | 7.8 | 7.8 | 7.2 | 7.1 | 10.6 |
| | 21 | 8.0 | 7.9 | 7.2 | 7.2 | 10.8 |
| | 22 | 8.1 | 7.9 | 7.3 | 7.3 | 10.9 |

| | | | | | |
|---|---|---|---|---|---|
| 23 | 8.2 | 7.9 | 7.3 | 7.5 | 11.1 |
| 24 | 8.3 | 8.0 | 7.3 | 7.8 | 11.2 |
| 25 | 8.4 | 8.1 | 7.5 | 8.1 | 11.3 |
| 26 | 8.5 | 8.2 | 7.8 | 8.3 | 11.4 |
| 27 | 8.6 | 8.4 | 8.0 | 8.6 | 11.5 |
| 28 | 9.0 | 8.5 | 8.2 | 8.7 | 11.6 |
| 29 | 9.1 | 8.5 | 8.2 | 8.8 | 11.7 |
| 30 | 9.2 | 8.5 | 8.2 | 8.9 | 11.9 |
| 31 | 9.3 | 8.5 | 8.2 | 9.0 | 12.1 |
| 32 | 9.5 | 8.5 | 8.2 | 9.1 | 12.4 |
| 33 | 9.7 | 8.7 | 8.2 | 9.2 | 12.6 |
| 34 | 9.8 | 8.9 | 8.3 | 9.4 | 12.7 |
| 35 | 9.8 | 9.1 | 8.3 | 9.6 | 12.7 |
| 36 | 9.8 | 9.3 | 8.3 | 9.7 | 12.7 |
| 37 | 9.9 | 9.3 | 8.6 | 9.8 | 12.7 |
| 38 | 10.0 | 9.3 | 8.9 | 9.9 | 12.8 |
| 39 | 10.1 | 9.3 | 9.3 | 10.0 | 13.1 |
| 40 | 10.3 | 9.4 | 9.6 | 10.1 | 13.3 |
| 41 | 10.4 | 9.5 | 9.6 | 10.2 | 13.5 |
| 42 | 10.4 | 9.6 | 9.7 | 10.3 | 13.7 |
| 43 | 10.6 | 9.7 | 9.7 | 10.3 | 13.7 |
| 44 | 10.7 | 9.8 | 9.8 | 10.4 | 13.7 |
| 45 | 11.1 | 10.0 | 9.8 | 10.4 | 13.7 |
| 46 | 11.3 | 10.2 | 9.8 | 10.4 | 13.7 |
| 47 | 11.3 | 10.4 | 9.8 | 10.4 | 13.9 |
| 48 | 11.4 | 10.6 | 9.8 | 10.4 | 14.1 |
| 49 | 11.6 | 10.9 | 10.0 | 10.5 | 14.2 |
| 50 | 11.6 | 11.1 | 10.1 | 10.5 | 14.4 |
| 51 | 11.7 | 11.3 | 10.2 | 10.8 | 14.7 |
| 52 | 11.8 | 11.6 | 10.4 | 11.1 | 15.0 |
| 53 | 11.9 | 11.6 | 10.6 | 11.4 | 15.2 |
| 54 | 12.0 | 11.6 | 10.8 | 11.7 | 15.5 |
| 55 | 12.1 | 11.6 | 11.1 | 11.9 | 15.7 |
| 56 | 12.3 | 11.6 | 11.3 | 11.9 | 15.9 |
| 57 | 12.5 | 11.8 | 11.3 | 12.0 | 16.0 |
| 58 | 12.7 | 12.0 | 11.3 | 12.0 | 16.2 |
| 59 | 12.9 | 12.2 | 11.3 | 12.0 | 16.3 |
| 60 | 13.2 | 12.4 | 11.3 | 12.1 | 16.5 |
| 61 | 13.5 | 12.8 | 11.4 | 12.1 | 16.7 |
| 62 | 13.7 | 13.2 | 11.5 | 12.2 | 16.9 |
| 63 | 13.7 | 13.7 | 11.5 | 12.2 | 17.2 |
| 64 | 13.8 | 14.1 | 11.6 | 12.3 | 17.7 |
| 65 | 14.2 | 14.2 | 11.7 | 12.5 | 18.2 |
| 66 | 14.4 | 14.4 | 11.8 | 12.7 | 18.7 |

| | | | | | |
|---|---|---|---|---|---|
| 67 | 14.5 | 14.6 | 11.9 | 12.8 | 19.0 |
| 68 | 15.0 | 14.8 | 12.0 | 13.0 | 19.0 |
| 69 | 15.2 | 15.2 | 12.3 | 13.2 | 19.0 |
| 70 | 15.4 | 15.6 | 12.7 | 13.3 | 19.0 |
| 71 | 15.7 | 16.0 | 13.0 | 13.5 | 19.0 |
| 72 | 15.9 | 16.4 | 13.3 | 13.6 | 19.0 |
| 73 | 16.1 | 16.4 | 13.6 | 13.8 | 19.0 |
| 74 | 16.2 | 16.4 | 13.9 | 14.2 | 19.0 |
| 75 | 16.3 | 16.4 | 14.1 | 14.5 | 19.0 |
| 76 | 16.4 | 16.4 | 14.4 | 14.8 | 19.0 |
| 77 | 16.4 | 16.7 | 14.6 | 15.2 | 19.1 |
| 78 | 16.4 | 17.0 | 14.8 | 15.4 | 19.1 |
| 79 | 16.5 | 17.3 | 15.0 | 15.5 | 19.2 |
| 80 | 17.1 | 17.5 | 15.2 | 15.6 | 19.5 |
| 81 | 17.4 | 17.7 | 15.5 | 15.7 | 20.0 |
| 82 | 17.6 | 17.8 | 15.7 | 15.8 | 20.4 |
| 83 | 18.1 | 18.0 | 16.0 | 15.9 | 20.8 |
| 84 | 18.4 | 18.1 | 16.2 | 15.9 | 21.1 |
| 85 | 18.8 | 18.2 | 16.3 | 16.0 | 21.2 |
| 86 | 18.9 | 18.2 | 16.3 | 16.0 | 21.4 |
| 87 | 18.9 | 18.2 | 16.3 | 16.2 | 21.5 |
| 88 | 19.0 | 18.3 | 16.4 | 16.5 | 21.6 |
| 89 | 19.0 | 18.4 | 16.4 | 16.8 | 21.8 |
| 90 | 19.0 | 18.6 | 16.4 | 17.1 | 21.9 |
| 91 | 19.2 | 18.7 | 16.4 | 17.4 | 22.1 |
| 92 | 19.6 | 18.9 | 16.4 | 17.7 | 22.3 |
| 93 | 19.9 | 20.0 | 17.0 | 18.1 | 22.7 |
| 94 | 21.0 | 21.0 | 17.6 | 18.4 | 23.0 |
| 95 | 21.6 | 22.1 | 18.2 | 18.8 | 23.4 |
| 96 | 22.2 | 23.2 | 18.8 | 19.0 | 23.7 |
| 97 | 23.2 | 23.2 | 19.0 | 19.2 | 23.8 |
| 98 | 23.2 | 23.2 | 19.2 | 19.4 | 23.9 |
| 99 | 23.7 | 23.2 | 19.4 | 19.6 | 23.9 |
| 100 | 24.0 | 23.2 | 19.6 | 19.8 | 24.0 |

***Table S13:*** *Anisochrony detection with music – Threshold (% of IOI) – Anisochrony_music_threshold*

| | | | | | | |
|---|---|---|---|---|---|---|
| **Task** | Anisochrony detection with music | | | | | |
| **Outcome measure** | Threshold (% of IOI) | | | | | |
| **Variable name** | Anisochrony_music_threshold | | | | | |
| **Age and gender effects** | | | | | | |
| Age regression (slope) | | 0.073 | | | | |
| Age regression (p) | | 0.017 | | | | |
| Age group (p) | | 0.095 | | | | |
| Gender (p) | | 0.570 | | | | |
| **Group** | | All | Age 18 to 21 | Age 22 to 29 | Age 30 to 54 | Age 55 to 87 |
| N (N female) | | 98 (68) | 26 (18) | 25 (17) | 24 (14) | 23 (19) |
| **Normality** | | | | | | |
| Skewness | | 0.95 | 1.17 | 1.25 | 0.82 | 0.74 |
| Excess kurtosis | | 0.11 | 0.36 | 1.25 | -0.58 | 0.07 |
| **Scores** | | | | | | |
| Mean | | 12.0 | 11.0 | 11.0 | 11.7 | 14.6 |
| SD | | 7.1 | 7.6 | 5.4 | 7.8 | 7.0 |
| Percentiles | | | | | | |
| | 0 | 3.2 | 3.6 | 4.1 | 3.2 | 4.7 |
| | 1 | 3.6 | 3.6 | 4.2 | 3.4 | 4.8 |
| | 2 | 3.7 | 3.7 | 4.4 | 3.6 | 4.9 |
| | 3 | 4.0 | 3.7 | 4.5 | 3.8 | 5.0 |
| | 4 | 4.0 | 3.7 | 4.7 | 3.9 | 5.1 |
| | 5 | 4.0 | 3.8 | 4.9 | 4.0 | 5.3 |
| | 6 | 4.3 | 3.9 | 5.1 | 4.1 | 5.7 |
| | 7 | 4.3 | 3.9 | 5.2 | 4.2 | 6.0 |
| | 8 | 4.4 | 4.0 | 5.4 | 4.3 | 6.3 |
| | 9 | 4.5 | 4.1 | 5.6 | 4.3 | 6.6 |
| | 10 | 4.5 | 4.2 | 5.7 | 4.4 | 6.9 |
| | 11 | 4.5 | 4.2 | 5.9 | 4.4 | 7.1 |
| | 12 | 4.6 | 4.3 | 6.0 | 4.4 | 7.3 |
| | 13 | 4.7 | 4.4 | 6.2 | 4.4 | 7.6 |
| | 14 | 4.8 | 4.5 | 6.3 | 4.4 | 7.8 |
| | 15 | 4.9 | 4.5 | 6.3 | 4.4 | 8.1 |
| | 16 | 5.1 | 4.6 | 6.4 | 4.5 | 8.4 |
| | 17 | 5.1 | 4.6 | 6.6 | 4.5 | 8.7 |
| | 18 | 5.3 | 4.7 | 6.8 | 4.5 | 9.0 |
| | 19 | 5.5 | 4.7 | 7.1 | 4.5 | 9.2 |
| | 20 | 5.5 | 4.8 | 7.3 | 4.5 | 9.3 |
| | 21 | 5.7 | 4.8 | 7.5 | 4.5 | 9.4 |
| | 22 | 6.2 | 4.9 | 7.6 | 4.6 | 9.5 |

| | | | | | |
|---|---|---|---|---|---|
| 23 | 6.5 | 5.0 | 7.6 | 4.8 | 9.6 |
| 24 | 6.5 | 5.1 | 7.7 | 5.0 | 9.8 |
| 25 | 6.5 | 5.1 | 7.7 | 5.2 | 9.9 |
| 26 | 6.6 | 5.1 | 7.8 | 5.5 | 10.0 |
| 27 | 6.7 | 5.1 | 7.8 | 5.5 | 10.1 |
| 28 | 6.9 | 5.1 | 7.9 | 5.5 | 10.2 |
| 29 | 7.2 | 5.4 | 7.9 | 5.5 | 10.3 |
| 30 | 7.5 | 5.8 | 8.0 | 5.5 | 10.4 |
| 31 | 7.7 | 6.1 | 8.2 | 5.6 | 10.5 |
| 32 | 7.7 | 6.5 | 8.3 | 5.9 | 10.6 |
| 33 | 7.8 | 6.5 | 8.4 | 6.1 | 10.7 |
| 34 | 7.8 | 6.6 | 8.5 | 6.3 | 10.7 |
| 35 | 7.9 | 6.6 | 8.6 | 6.6 | 10.8 |
| 36 | 8.0 | 6.7 | 8.6 | 6.9 | 10.9 |
| 37 | 8.2 | 6.7 | 8.7 | 7.3 | 11.0 |
| 38 | 8.4 | 6.8 | 8.8 | 7.6 | 11.2 |
| 39 | 8.7 | 6.9 | 8.9 | 7.9 | 11.4 |
| 40 | 8.8 | 6.9 | 8.9 | 8.0 | 11.5 |
| 41 | 9.0 | 7.0 | 9.0 | 8.1 | 11.7 |
| 42 | 9.1 | 7.0 | 9.1 | 8.1 | 11.9 |
| 43 | 9.3 | 7.1 | 9.1 | 8.2 | 12.1 |
| 44 | 9.5 | 7.1 | 9.2 | 8.3 | 12.3 |
| 45 | 9.7 | 7.3 | 9.3 | 8.4 | 12.5 |
| 46 | 9.9 | 7.5 | 9.4 | 8.5 | 12.7 |
| 47 | 10.1 | 7.6 | 9.5 | 8.7 | 12.8 |
| 48 | 10.2 | 7.8 | 9.5 | 8.9 | 13.0 |
| 49 | 10.4 | 7.8 | 9.6 | 9.2 | 13.2 |
| 50 | 10.6 | 7.8 | 9.7 | 9.5 | 13.4 |
| 51 | 10.7 | 7.8 | 9.8 | 9.9 | 13.4 |
| 52 | 10.9 | 7.8 | 9.9 | 10.2 | 13.4 |
| 53 | 11.0 | 8.6 | 10.0 | 10.4 | 13.4 |
| 54 | 11.1 | 9.4 | 10.0 | 10.6 | 13.5 |
| 55 | 11.1 | 10.2 | 10.2 | 10.7 | 13.5 |
| 56 | 11.2 | 11.0 | 10.3 | 10.9 | 13.6 |
| 57 | 11.4 | 11.1 | 10.4 | 11.0 | 13.7 |
| 58 | 11.6 | 11.1 | 10.5 | 11.0 | 13.8 |
| 59 | 11.7 | 11.1 | 10.7 | 11.1 | 13.8 |
| 60 | 11.8 | 11.1 | 10.9 | 11.1 | 14.7 |
| 61 | 11.8 | 11.3 | 11.0 | 11.2 | 15.6 |
| 62 | 11.9 | 11.5 | 11.2 | 11.3 | 16.6 |
| 63 | 12.0 | 11.6 | 11.3 | 11.5 | 17.5 |
| 64 | 12.3 | 11.8 | 11.4 | 11.6 | 18.1 |
| 65 | 12.4 | 11.8 | 11.5 | 11.7 | 18.1 |
| 66 | 12.6 | 11.8 | 11.5 | 12.3 | 18.1 |

| | | | | | |
|---|---|---|---|---|---|
| 67 | 13.4 | 11.9 | 11.6 | 13.1 | 18.1 |
| 68 | 13.5 | 11.9 | 11.7 | 13.8 | 18.1 |
| 69 | 13.8 | 12.0 | 11.8 | 14.6 | 18.3 |
| 70 | 14.1 | 12.2 | 11.9 | 15.0 | 18.4 |
| 71 | 14.3 | 12.3 | 12.0 | 15.0 | 18.5 |
| 72 | 14.9 | 12.4 | 12.1 | 15.0 | 18.6 |
| 73 | 15.0 | 12.9 | 12.1 | 15.0 | 18.8 |
| 74 | 15.4 | 13.4 | 12.2 | 15.1 | 18.9 |
| 75 | 15.9 | 13.9 | 12.3 | 15.9 | 19.1 |
| 76 | 16.2 | 14.4 | 12.7 | 16.7 | 19.2 |
| 77 | 16.3 | 14.8 | 13.2 | 17.6 | 19.4 |
| 78 | 17.4 | 15.3 | 13.6 | 18.4 | 19.5 |
| 79 | 18.1 | 15.8 | 14.1 | 18.9 | 19.7 |
| 80 | 18.1 | 16.2 | 14.4 | 19.2 | 19.9 |
| 81 | 18.4 | 16.3 | 14.7 | 19.5 | 20.0 |
| 82 | 18.7 | 16.3 | 15.0 | 19.8 | 20.2 |
| 83 | 19.1 | 16.3 | 15.4 | 20.3 | 20.3 |
| 84 | 19.7 | 16.4 | 15.6 | 20.8 | 20.4 |
| 85 | 20.1 | 17.6 | 15.7 | 21.4 | 20.5 |
| 86 | 20.4 | 18.9 | 15.9 | 22.0 | 20.6 |
| 87 | 20.9 | 20.1 | 16.0 | 22.5 | 20.8 |
| 88 | 21.4 | 21.4 | 16.3 | 22.7 | 20.9 |
| 89 | 21.8 | 22.2 | 16.8 | 22.9 | 21.1 |
| 90 | 22.7 | 23.0 | 17.2 | 23.0 | 21.3 |
| 91 | 23.0 | 23.8 | 17.6 | 23.2 | 21.5 |
| 92 | 23.6 | 24.6 | 18.3 | 24.0 | 23.0 |
| 93 | 24.9 | 25.7 | 19.5 | 25.0 | 24.5 |
| 94 | 26.4 | 26.7 | 20.7 | 26.0 | 26.0 |
| 95 | 27.7 | 27.8 | 22.0 | 27.0 | 27.5 |
| 96 | 27.9 | 28.9 | 23.1 | 27.7 | 28.6 |
| 97 | 28.2 | 28.9 | 23.9 | 27.7 | 29.5 |
| 98 | 28.9 | 29.0 | 24.6 | 27.8 | 30.3 |
| 99 | 29.3 | 29.1 | 25.4 | 27.8 | 31.2 |
| 100 | 32.0 | 29.2 | 26.1 | 27.9 | 32.0 |

***Table S14:*** *Beat Alignment Test – Sensitivity index (d') for all trials – BAT_all_dprime*

| | | | | | | |
|---|---|---|---|---|---|---|
| **Task** | | Beat Alignment Test | | | | |
| **Outcome measure** | | Sensitivity index (d') for all trials | | | | |
| **Variable name** | | BAT_all_dprime | | | | |
| **Age and gender effects** | | | | | | |
| Age regression (slope) | | -0.010 | | | | |
| Age regression (p) | | 0.059 | | | | |
| Age group (p) | | 0.223 | | | | |
| Gender (p) | | 0.503 | | | | |
| **Group** | | All | Age 18 to 21 | Age 22 to 29 | Age 30 to 54 | Age 55 to 87 |
| N (N female) | | 108 (74) | 27 (18) | 29 (19) | 26 (16) | 26 (21) |
| **Normality** | | | | | | |
| Skewness | | -0.23 | -0.86 | 0.16 | -0.58 | 0.22 |
| Excess kurtosis | | -0.40 | 0.41 | -1.03 | 0.06 | -0.39 |
| **Scores** | | | | | | |
| Mean | | 2.76 | 3.02 | 2.83 | 2.69 | 2.51 |
| SD | | 1.03 | 1.06 | 1.10 | 1.07 | 0.87 |
| Percentiles | | | | | | |
| | 0 | 0.05 | 0.42 | 0.89 | 0.05 | 1.06 |
| | 1 | 0.43 | 0.46 | 0.94 | 0.23 | 1.07 |
| | 2 | 0.60 | 0.50 | 0.99 | 0.40 | 1.08 |
| | 3 | 0.78 | 0.54 | 1.04 | 0.58 | 1.09 |
| | 4 | 0.94 | 0.60 | 1.09 | 0.76 | 1.10 |
| | 5 | 1.07 | 0.78 | 1.12 | 0.88 | 1.10 |
| | 6 | 1.08 | 0.95 | 1.16 | 1.00 | 1.11 |
| | 7 | 1.11 | 1.13 | 1.20 | 1.12 | 1.11 |
| | 8 | 1.16 | 1.33 | 1.30 | 1.24 | 1.11 |
| | 9 | 1.23 | 1.57 | 1.41 | 1.26 | 1.24 |
| | 10 | 1.25 | 1.80 | 1.53 | 1.27 | 1.37 |
| | 11 | 1.29 | 2.04 | 1.63 | 1.29 | 1.50 |
| | 12 | 1.53 | 2.18 | 1.70 | 1.31 | 1.63 |
| | 13 | 1.60 | 2.20 | 1.77 | 1.37 | 1.66 |
| | 14 | 1.63 | 2.23 | 1.84 | 1.44 | 1.69 |
| | 15 | 1.74 | 2.25 | 1.88 | 1.50 | 1.71 |
| | 16 | 1.76 | 2.31 | 1.91 | 1.57 | 1.74 |
| | 17 | 1.82 | 2.38 | 1.94 | 1.72 | 1.74 |
| | 18 | 1.89 | 2.46 | 1.98 | 1.87 | 1.75 |
| | 19 | 2.01 | 2.53 | 2.01 | 2.01 | 1.75 |
| | 20 | 2.09 | 2.56 | 2.05 | 2.16 | 1.76 |
| | 21 | 2.10 | 2.58 | 2.08 | 2.17 | 1.77 |
| | 22 | 2.12 | 2.60 | 2.10 | 2.19 | 1.78 |
| | 23 | 2.14 | 2.62 | 2.11 | 2.20 | 1.80 |
| | 24 | 2.14 | 2.62 | 2.12 | 2.21 | 1.81 |

| | | | | | |
|---|---|---|---|---|---|
| 25 | 2.15 | 2.62 | 2.13 | 2.22 | 1.88 |
| 26 | 2.16 | 2.63 | 2.13 | 2.23 | 1.95 |
| 27 | 2.17 | 2.63 | 2.14 | 2.24 | 2.01 |
| 28 | 2.21 | 2.65 | 2.14 | 2.25 | 2.08 |
| 29 | 2.24 | 2.66 | 2.15 | 2.26 | 2.09 |
| 30 | 2.25 | 2.67 | 2.18 | 2.28 | 2.09 |
| 31 | 2.27 | 2.69 | 2.21 | 2.30 | 2.09 |
| 32 | 2.33 | 2.69 | 2.24 | 2.31 | 2.10 |
| 33 | 2.38 | 2.70 | 2.28 | 2.33 | 2.11 |
| 34 | 2.40 | 2.71 | 2.32 | 2.34 | 2.12 |
| 35 | 2.44 | 2.71 | 2.36 | 2.36 | 2.13 |
| 36 | 2.47 | 2.73 | 2.39 | 2.38 | 2.14 |
| 37 | 2.48 | 2.74 | 2.40 | 2.44 | 2.14 |
| 38 | 2.52 | 2.75 | 2.41 | 2.50 | 2.15 |
| 39 | 2.54 | 2.76 | 2.41 | 2.57 | 2.15 |
| 40 | 2.55 | 2.76 | 2.43 | 2.63 | 2.16 |
| 41 | 2.61 | 2.76 | 2.44 | 2.65 | 2.24 |
| 42 | 2.62 | 2.76 | 2.46 | 2.67 | 2.32 |
| 43 | 2.62 | 2.79 | 2.47 | 2.69 | 2.40 |
| 44 | 2.62 | 2.83 | 2.47 | 2.71 | 2.48 |
| 45 | 2.63 | 2.87 | 2.47 | 2.72 | 2.50 |
| 46 | 2.63 | 2.91 | 2.47 | 2.73 | 2.51 |
| 47 | 2.65 | 2.92 | 2.48 | 2.75 | 2.52 |
| 48 | 2.69 | 2.92 | 2.50 | 2.76 | 2.54 |
| 49 | 2.70 | 2.93 | 2.52 | 2.80 | 2.56 |
| 50 | 2.71 | 2.93 | 2.54 | 2.84 | 2.58 |
| 51 | 2.71 | 3.03 | 2.56 | 2.89 | 2.60 |
| 52 | 2.74 | 3.13 | 2.58 | 2.93 | 2.62 |
| 53 | 2.76 | 3.23 | 2.61 | 2.95 | 2.62 |
| 54 | 2.76 | 3.32 | 2.62 | 2.96 | 2.62 |
| 55 | 2.90 | 3.34 | 2.62 | 2.98 | 2.63 |
| 56 | 2.92 | 3.37 | 2.62 | 3.00 | 2.63 |
| 57 | 2.93 | 3.39 | 2.62 | 3.01 | 2.64 |
| 58 | 2.93 | 3.41 | 2.69 | 3.02 | 2.66 |
| 59 | 2.93 | 3.41 | 2.78 | 3.04 | 2.67 |
| 60 | 2.94 | 3.41 | 2.86 | 3.05 | 2.69 |
| 61 | 3.01 | 3.41 | 2.94 | 3.09 | 2.69 |
| 62 | 3.06 | 3.41 | 3.00 | 3.14 | 2.70 |
| 63 | 3.10 | 3.42 | 3.06 | 3.18 | 2.70 |
| 64 | 3.14 | 3.43 | 3.12 | 3.22 | 2.71 |
| 65 | 3.14 | 3.43 | 3.14 | 3.22 | 2.76 |
| 66 | 3.19 | 3.50 | 3.14 | 3.22 | 2.82 |
| 67 | 3.22 | 3.60 | 3.14 | 3.22 | 2.87 |
| 68 | 3.22 | 3.70 | 3.17 | 3.22 | 2.93 |

| | | | | | |
|---|---|---|---|---|---|
| 69 | 3.30 | 3.79 | 3.36 | 3.25 | 2.93 |
| 70 | 3.31 | 3.82 | 3.55 | 3.27 | 2.93 |
| 71 | 3.31 | 3.82 | 3.74 | 3.29 | 2.93 |
| 72 | 3.32 | 3.82 | 3.82 | 3.31 | 2.93 |
| 73 | 3.41 | 3.82 | 3.82 | 3.31 | 2.96 |
| 74 | 3.42 | 3.82 | 3.82 | 3.31 | 3.00 |
| 75 | 3.46 | 3.82 | 3.82 | 3.31 | 3.03 |
| 76 | 3.52 | 3.82 | 3.82 | 3.31 | 3.07 |
| 77 | 3.52 | 3.82 | 3.82 | 3.37 | 3.09 |
| 78 | 3.66 | 3.82 | 3.82 | 3.42 | 3.11 |
| 79 | 3.82 | 3.82 | 3.89 | 3.47 | 3.12 |
| 80 | 3.82 | 3.82 | 4.06 | 3.52 | 3.14 |
| 81 | 3.82 | 3.83 | 4.23 | 3.52 | 3.16 |
| 82 | 3.82 | 3.89 | 4.40 | 3.52 | 3.18 |
| 83 | 3.82 | 3.95 | 4.42 | 3.52 | 3.20 |
| 84 | 3.82 | 4.00 | 4.42 | 3.52 | 3.21 |
| 85 | 3.82 | 4.04 | 4.42 | 3.52 | 3.24 |
| 86 | 4.04 | 4.04 | 4.42 | 3.52 | 3.26 |
| 87 | 4.04 | 4.04 | 4.42 | 3.52 | 3.29 |
| 88 | 4.04 | 4.04 | 4.42 | 3.52 | 3.31 |
| 89 | 4.13 | 4.09 | 4.42 | 3.65 | 3.44 |
| 90 | 4.42 | 4.19 | 4.42 | 3.78 | 3.57 |
| 91 | 4.42 | 4.29 | 4.42 | 3.91 | 3.69 |
| 92 | 4.42 | 4.39 | 4.42 | 4.04 | 3.82 |
| 93 | 4.42 | 4.42 | 4.42 | 4.13 | 3.87 |
| 94 | 4.42 | 4.42 | 4.42 | 4.23 | 3.93 |
| 95 | 4.42 | 4.42 | 4.42 | 4.33 | 3.98 |
| 96 | 4.42 | 4.42 | 4.42 | 4.42 | 4.04 |
| 97 | 4.42 | 4.42 | 4.42 | 4.42 | 4.13 |
| 98 | 4.42 | 4.42 | 4.42 | 4.42 | 4.23 |
| 99 | 4.42 | 4.42 | 4.42 | 4.42 | 4.33 |
| 100 | 4.42 | 4.42 | 4.42 | 4.42 | 4.42 |

***Table S15:*** *Beat Alignment Test – Sensitivity index (d') for fast tempo trials – BAT_fast_dprime*

| | | | | | | |
|---|---|---|---|---|---|---|
| **Task** | | Beat Alignment Test | | | | |
| **Outcome measure** | | Sensitivity index (d') for fast tempo trials | | | | |
| **Variable name** | | BAT_fast_dprime | | | | |
| **Age and gender effects** | | | | | | |
| Age regression (slope) | | -0.010 | | | | |
| Age regression (p) | | 0.015 | | | | |
| Age group (p) | | 0.137 | | | | |
| Gender (p) | | 1.000 | | | | |
| **Group** | | All | Age 18 to 21 | Age 22 to 29 | Age 30 to 54 | Age 55 to 87 |
| N (N female) | | 108 (74) | 27 (18) | 29 (19) | 26 (16) | 26 (21) |
| **Normality** | | | | | | |
| Skewness | | -0.24 | -1.16 | -0.13 | -0.20 | 0.24 |
| Excess kurtosis | | -0.44 | 1.34 | -0.91 | -0.09 | -0.25 |
| **Scores** | | | | | | |
| Mean | | 2.22 | 2.46 | 2.31 | 2.12 | 1.98 |
| SD | | 0.87 | 0.80 | 0.96 | 0.83 | 0.82 |
| Percentiles | | | | | | |
| | 0 | 0.29 | 0.29 | 0.29 | 0.29 | 0.59 |
| | 1 | 0.29 | 0.34 | 0.38 | 0.34 | 0.59 |
| | 2 | 0.32 | 0.39 | 0.46 | 0.38 | 0.60 |
| | 3 | 0.47 | 0.43 | 0.54 | 0.43 | 0.61 |
| | 4 | 0.50 | 0.50 | 0.67 | 0.47 | 0.62 |
| | 5 | 0.59 | 0.72 | 0.87 | 0.62 | 0.64 |
| | 6 | 0.60 | 0.94 | 1.07 | 0.77 | 0.67 |
| | 7 | 0.67 | 1.16 | 1.28 | 0.92 | 0.70 |
| | 8 | 0.86 | 1.35 | 1.31 | 1.06 | 0.73 |
| | 9 | 0.99 | 1.49 | 1.31 | 1.13 | 0.79 |
| | 10 | 1.05 | 1.62 | 1.31 | 1.19 | 0.85 |
| | 11 | 1.25 | 1.75 | 1.32 | 1.25 | 0.91 |
| | 12 | 1.31 | 1.85 | 1.35 | 1.31 | 0.97 |
| | 13 | 1.31 | 1.90 | 1.38 | 1.35 | 0.98 |
| | 14 | 1.31 | 1.96 | 1.41 | 1.39 | 0.99 |
| | 15 | 1.42 | 2.02 | 1.42 | 1.43 | 1.00 |
| | 16 | 1.42 | 2.04 | 1.42 | 1.48 | 1.01 |
| | 17 | 1.43 | 2.05 | 1.42 | 1.51 | 1.11 |
| | 18 | 1.51 | 2.06 | 1.42 | 1.55 | 1.21 |
| | 19 | 1.60 | 2.06 | 1.47 | 1.59 | 1.31 |
| | 20 | 1.61 | 2.06 | 1.52 | 1.62 | 1.42 |
| | 21 | 1.62 | 2.06 | 1.58 | 1.63 | 1.47 |
| | 22 | 1.63 | 2.06 | 1.60 | 1.63 | 1.52 |
| | 23 | 1.64 | 2.06 | 1.60 | 1.63 | 1.57 |
| | 24 | 1.64 | 2.13 | 1.60 | 1.64 | 1.62 |

| | | | | | |
|---|---|---|---|---|---|
| 25 | 1.74 | 2.19 | 1.60 | 1.67 | 1.66 |
| 26 | 1.77 | 2.26 | 1.61 | 1.70 | 1.70 |
| 27 | 1.77 | 2.32 | 1.62 | 1.74 | 1.73 |
| 28 | 1.77 | 2.32 | 1.63 | 1.77 | 1.77 |
| 29 | 1.77 | 2.32 | 1.64 | 1.77 | 1.77 |
| 30 | 1.82 | 2.32 | 1.64 | 1.77 | 1.77 |
| 31 | 1.82 | 2.33 | 1.64 | 1.77 | 1.77 |
| 32 | 1.82 | 2.34 | 1.64 | 1.77 | 1.77 |
| 33 | 1.85 | 2.36 | 1.67 | 1.78 | 1.78 |
| 34 | 1.91 | 2.37 | 1.70 | 1.79 | 1.79 |
| 35 | 1.97 | 2.38 | 1.74 | 1.81 | 1.81 |
| 36 | 2.04 | 2.38 | 1.79 | 1.82 | 1.82 |
| 37 | 2.04 | 2.38 | 1.87 | 1.88 | 1.82 |
| 38 | 2.04 | 2.38 | 1.94 | 1.93 | 1.82 |
| 39 | 2.06 | 2.38 | 2.02 | 1.99 | 1.82 |
| 40 | 2.06 | 2.38 | 2.08 | 2.04 | 1.82 |
| 41 | 2.06 | 2.38 | 2.13 | 2.05 | 1.84 |
| 42 | 2.06 | 2.38 | 2.17 | 2.05 | 1.87 |
| 43 | 2.06 | 2.42 | 2.22 | 2.06 | 1.89 |
| 44 | 2.06 | 2.47 | 2.22 | 2.06 | 1.91 |
| 45 | 2.06 | 2.51 | 2.22 | 2.06 | 1.91 |
| 46 | 2.06 | 2.56 | 2.22 | 2.06 | 1.91 |
| 47 | 2.11 | 2.57 | 2.23 | 2.06 | 1.91 |
| 48 | 2.22 | 2.57 | 2.26 | 2.06 | 1.91 |
| 49 | 2.22 | 2.57 | 2.29 | 2.06 | 1.95 |
| 50 | 2.27 | 2.57 | 2.32 | 2.06 | 1.98 |
| 51 | 2.32 | 2.57 | 2.32 | 2.06 | 2.01 |
| 52 | 2.32 | 2.57 | 2.32 | 2.06 | 2.04 |
| 53 | 2.32 | 2.57 | 2.32 | 2.13 | 2.05 |
| 54 | 2.32 | 2.57 | 2.32 | 2.19 | 2.05 |
| 55 | 2.32 | 2.57 | 2.32 | 2.26 | 2.06 |
| 56 | 2.38 | 2.57 | 2.32 | 2.32 | 2.06 |
| 57 | 2.38 | 2.57 | 2.32 | 2.34 | 2.06 |
| 58 | 2.38 | 2.59 | 2.34 | 2.35 | 2.06 |
| 59 | 2.38 | 2.64 | 2.36 | 2.37 | 2.06 |
| 60 | 2.38 | 2.70 | 2.37 | 2.38 | 2.06 |
| 61 | 2.38 | 2.76 | 2.40 | 2.38 | 2.06 |
| 62 | 2.38 | 2.79 | 2.45 | 2.38 | 2.06 |
| 63 | 2.38 | 2.79 | 2.50 | 2.38 | 2.06 |
| 64 | 2.38 | 2.79 | 2.56 | 2.38 | 2.06 |
| 65 | 2.49 | 2.79 | 2.61 | 2.38 | 2.06 |
| 66 | 2.57 | 2.79 | 2.68 | 2.38 | 2.06 |
| 67 | 2.57 | 2.79 | 2.74 | 2.38 | 2.06 |
| 68 | 2.57 | 2.79 | 2.80 | 2.38 | 2.06 |

| | | | | | |
|---|---|---|---|---|---|
| 69 | 2.57 | 2.79 | 2.88 | 2.38 | 2.10 |
| 70 | 2.57 | 2.85 | 2.96 | 2.38 | 2.14 |
| 71 | 2.78 | 2.92 | 3.04 | 2.38 | 2.18 |
| 72 | 2.79 | 2.99 | 3.15 | 2.38 | 2.22 |
| 73 | 2.79 | 3.07 | 3.28 | 2.43 | 2.26 |
| 74 | 2.79 | 3.07 | 3.41 | 2.48 | 2.30 |
| 75 | 2.79 | 3.07 | 3.54 | 2.52 | 2.34 |
| 76 | 2.79 | 3.07 | 3.54 | 2.57 | 2.38 |
| 77 | 2.79 | 3.07 | 3.54 | 2.62 | 2.38 |
| 78 | 2.92 | 3.07 | 3.54 | 2.68 | 2.38 |
| 79 | 3.07 | 3.07 | 3.54 | 2.73 | 2.38 |
| 80 | 3.07 | 3.07 | 3.54 | 2.79 | 2.38 |
| 81 | 3.07 | 3.07 | 3.54 | 2.79 | 2.49 |
| 82 | 3.07 | 3.07 | 3.54 | 2.79 | 2.59 |
| 83 | 3.07 | 3.07 | 3.54 | 2.79 | 2.69 |
| 84 | 3.07 | 3.07 | 3.54 | 2.79 | 2.79 |
| 85 | 3.51 | 3.07 | 3.54 | 2.86 | 2.79 |
| 86 | 3.54 | 3.07 | 3.54 | 2.93 | 2.79 |
| 87 | 3.54 | 3.07 | 3.54 | 3.00 | 2.79 |
| 88 | 3.54 | 3.07 | 3.54 | 3.07 | 2.79 |
| 89 | 3.54 | 3.14 | 3.54 | 3.19 | 2.98 |
| 90 | 3.54 | 3.26 | 3.54 | 3.31 | 3.16 |
| 91 | 3.54 | 3.38 | 3.54 | 3.42 | 3.35 |
| 92 | 3.54 | 3.50 | 3.54 | 3.54 | 3.54 |
| 93 | 3.54 | 3.54 | 3.54 | 3.54 | 3.54 |
| 94 | 3.54 | 3.54 | 3.54 | 3.54 | 3.54 |
| 95 | 3.54 | 3.54 | 3.54 | 3.54 | 3.54 |
| 96 | 3.54 | 3.54 | 3.54 | 3.54 | 3.54 |
| 97 | 3.54 | 3.54 | 3.54 | 3.54 | 3.54 |
| 98 | 3.54 | 3.54 | 3.54 | 3.54 | 3.54 |
| 99 | 3.54 | 3.54 | 3.54 | 3.54 | 3.54 |
| 100 | 3.54 | 3.54 | 3.54 | 3.54 | 3.54 |

***Table S16:*** *Beat Alignment Test – Sensitivity index (d') for medium tempo trials – BAT_med_dprime*

| **Task** | Beat Alignment Test | | | | |
|---|---|---|---|---|---|
| **Outcome measure** | Sensitivity index (d') for medium tempo trials | | | | |
| **Variable name** | BAT_med_dprime | | | | |
| **Age and gender effects** | | | | | |
| Age regression (slope) | -0.005 | | | | |
| Age regression (p) | 0.191 | | | | |
| Age group (p) | 0.310 | | | | |
| Gender (p) | 0.964 | | | | |
| **Group** | All | Age 18 to 21 | Age 22 to 29 | Age 30 to 54 | Age 55 to 87 |
| N (N female) | 108 (74) | 27 (18) | 29 (19) | 26 (16) | 26 (21) |
| **Normality** | | | | | |
| Skewness | -0.72 | -1.65 | -0.17 | -0.36 | 0.11 |
| Excess kurtosis | 0.68 | 2.47 | -0.89 | -0.22 | -0.88 |
| **Scores** | | | | | |
| Mean | 2.43 | 2.61 | 2.44 | 2.30 | 2.36 |
| SD | 0.87 | 1.04 | 0.84 | 0.93 | 0.63 |
| Percentiles | | | | | |
| 0 | -0.62 | -0.62 | 0.59 | 0.00 | 1.31 |
| 1 | 0.01 | -0.42 | 0.75 | 0.23 | 1.35 |
| 2 | 0.21 | -0.22 | 0.91 | 0.45 | 1.39 |
| 3 | 0.65 | -0.02 | 1.07 | 0.68 | 1.43 |
| 4 | 0.88 | 0.18 | 1.20 | 0.91 | 1.47 |
| 5 | 1.00 | 0.37 | 1.29 | 0.97 | 1.47 |
| 6 | 1.17 | 0.56 | 1.37 | 1.04 | 1.47 |
| 7 | 1.24 | 0.74 | 1.46 | 1.10 | 1.47 |
| 8 | 1.31 | 0.97 | 1.47 | 1.17 | 1.47 |
| 9 | 1.41 | 1.27 | 1.47 | 1.20 | 1.51 |
| 10 | 1.47 | 1.57 | 1.47 | 1.24 | 1.55 |
| 11 | 1.47 | 1.88 | 1.47 | 1.28 | 1.58 |
| 12 | 1.47 | 2.06 | 1.47 | 1.31 | 1.62 |
| 13 | 1.47 | 2.11 | 1.47 | 1.35 | 1.62 |
| 14 | 1.48 | 2.15 | 1.47 | 1.39 | 1.62 |
| 15 | 1.60 | 2.20 | 1.50 | 1.43 | 1.62 |
| 16 | 1.60 | 2.23 | 1.53 | 1.48 | 1.62 |
| 17 | 1.62 | 2.26 | 1.57 | 1.51 | 1.62 |
| 18 | 1.62 | 2.29 | 1.60 | 1.54 | 1.62 |
| 19 | 1.63 | 2.32 | 1.61 | 1.57 | 1.62 |
| 20 | 1.69 | 2.32 | 1.62 | 1.60 | 1.62 |
| 21 | 1.77 | 2.32 | 1.63 | 1.64 | 1.70 |
| 22 | 1.80 | 2.32 | 1.66 | 1.68 | 1.77 |

| | | | | | |
|---|---|---|---|---|---|
| 23 | 1.88 | 2.32 | 1.69 | 1.73 | 1.84 |
| 24 | 1.91 | 2.32 | 1.73 | 1.77 | 1.91 |
| 25 | 1.91 | 2.32 | 1.77 | 1.78 | 1.92 |
| 26 | 1.91 | 2.32 | 1.81 | 1.79 | 1.92 |
| 27 | 1.92 | 2.33 | 1.85 | 1.81 | 1.92 |
| 28 | 2.04 | 2.34 | 1.89 | 1.82 | 1.92 |
| 29 | 2.04 | 2.36 | 1.93 | 1.84 | 1.95 |
| 30 | 2.04 | 2.37 | 1.96 | 1.87 | 1.98 |
| 31 | 2.06 | 2.38 | 2.00 | 1.89 | 2.01 |
| 32 | 2.06 | 2.38 | 2.04 | 1.91 | 2.04 |
| 33 | 2.11 | 2.38 | 2.05 | 1.91 | 2.05 |
| 34 | 2.22 | 2.38 | 2.05 | 1.91 | 2.05 |
| 35 | 2.22 | 2.40 | 2.06 | 1.91 | 2.06 |
| 36 | 2.27 | 2.45 | 2.07 | 1.91 | 2.06 |
| 37 | 2.32 | 2.50 | 2.12 | 1.95 | 2.10 |
| 38 | 2.32 | 2.55 | 2.16 | 1.99 | 2.14 |
| 39 | 2.32 | 2.57 | 2.20 | 2.03 | 2.18 |
| 40 | 2.32 | 2.57 | 2.24 | 2.06 | 2.22 |
| 41 | 2.32 | 2.57 | 2.27 | 2.13 | 2.24 |
| 42 | 2.32 | 2.57 | 2.30 | 2.19 | 2.27 |
| 43 | 2.32 | 2.61 | 2.32 | 2.26 | 2.30 |
| 44 | 2.38 | 2.67 | 2.32 | 2.32 | 2.32 |
| 45 | 2.38 | 2.72 | 2.32 | 2.32 | 2.34 |
| 46 | 2.38 | 2.78 | 2.32 | 2.32 | 2.35 |
| 47 | 2.38 | 2.79 | 2.33 | 2.32 | 2.37 |
| 48 | 2.38 | 2.79 | 2.35 | 2.32 | 2.38 |
| 49 | 2.38 | 2.79 | 2.37 | 2.34 | 2.38 |
| 50 | 2.38 | 2.79 | 2.38 | 2.35 | 2.38 |
| 51 | 2.38 | 2.79 | 2.44 | 2.37 | 2.38 |
| 52 | 2.50 | 2.79 | 2.49 | 2.38 | 2.38 |
| 53 | 2.57 | 2.79 | 2.54 | 2.38 | 2.38 |
| 54 | 2.57 | 2.80 | 2.57 | 2.38 | 2.38 |
| 55 | 2.57 | 2.87 | 2.57 | 2.38 | 2.38 |
| 56 | 2.57 | 2.95 | 2.57 | 2.38 | 2.38 |
| 57 | 2.57 | 3.02 | 2.57 | 2.38 | 2.43 |
| 58 | 2.57 | 3.07 | 2.57 | 2.38 | 2.48 |
| 59 | 2.60 | 3.07 | 2.57 | 2.38 | 2.52 |
| 60 | 2.79 | 3.07 | 2.57 | 2.38 | 2.57 |
| 61 | 2.79 | 3.07 | 2.59 | 2.43 | 2.57 |
| 62 | 2.79 | 3.07 | 2.65 | 2.48 | 2.57 |
| 63 | 2.79 | 3.07 | 2.71 | 2.52 | 2.57 |
| 64 | 2.79 | 3.07 | 2.77 | 2.57 | 2.57 |
| 65 | 2.79 | 3.07 | 2.79 | 2.62 | 2.62 |
| 66 | 2.79 | 3.07 | 2.79 | 2.68 | 2.68 |

| | | | | | |
|---|---|---|---|---|---|
| 67 | 2.79 | 3.07 | 2.79 | 2.73 | 2.73 |
| 68 | 2.79 | 3.07 | 2.80 | 2.79 | 2.79 |
| 69 | 2.79 | 3.07 | 2.88 | 2.79 | 2.79 |
| 70 | 2.79 | 3.17 | 2.96 | 2.79 | 2.79 |
| 71 | 3.06 | 3.29 | 3.04 | 2.79 | 2.79 |
| 72 | 3.07 | 3.41 | 3.07 | 2.79 | 2.79 |
| 73 | 3.07 | 3.53 | 3.07 | 2.79 | 2.79 |
| 74 | 3.07 | 3.54 | 3.07 | 2.79 | 2.79 |
| 75 | 3.07 | 3.54 | 3.07 | 2.79 | 2.79 |
| 76 | 3.07 | 3.54 | 3.20 | 2.79 | 2.79 |
| 77 | 3.07 | 3.54 | 3.33 | 2.98 | 2.79 |
| 78 | 3.07 | 3.54 | 3.46 | 3.16 | 2.79 |
| 79 | 3.32 | 3.54 | 3.54 | 3.35 | 2.79 |
| 80 | 3.54 | 3.54 | 3.54 | 3.54 | 2.79 |
| 81 | 3.54 | 3.54 | 3.54 | 3.54 | 2.86 |
| 82 | 3.54 | 3.54 | 3.54 | 3.54 | 2.93 |
| 83 | 3.54 | 3.54 | 3.54 | 3.54 | 3.00 |
| 84 | 3.54 | 3.54 | 3.54 | 3.54 | 3.07 |
| 85 | 3.54 | 3.54 | 3.54 | 3.54 | 3.07 |
| 86 | 3.54 | 3.54 | 3.54 | 3.54 | 3.07 |
| 87 | 3.54 | 3.54 | 3.54 | 3.54 | 3.07 |
| 88 | 3.54 | 3.54 | 3.54 | 3.54 | 3.07 |
| 89 | 3.54 | 3.54 | 3.54 | 3.54 | 3.07 |
| 90 | 3.54 | 3.54 | 3.54 | 3.54 | 3.07 |
| 91 | 3.54 | 3.54 | 3.54 | 3.54 | 3.07 |
| 92 | 3.54 | 3.54 | 3.54 | 3.54 | 3.07 |
| 93 | 3.54 | 3.54 | 3.54 | 3.54 | 3.19 |
| 94 | 3.54 | 3.54 | 3.54 | 3.54 | 3.31 |
| 95 | 3.54 | 3.54 | 3.54 | 3.54 | 3.42 |
| 96 | 3.54 | 3.54 | 3.54 | 3.54 | 3.54 |
| 97 | 3.54 | 3.54 | 3.54 | 3.54 | 3.54 |
| 98 | 3.54 | 3.54 | 3.54 | 3.54 | 3.54 |
| 99 | 3.54 | 3.54 | 3.54 | 3.54 | 3.54 |
| 100 | 3.54 | 3.54 | 3.54 | 3.54 | 3.54 |

***Table S17:*** *Beat Alignment Test – Sensitivity index (d') for slow tempo trials – BAT_slow_dprime*

| | | | | | | |
|---|---|---|---|---|---|---|
| **Task** | | Beat Alignment Test | | | | |
| **Outcome measure** | | Sensitivity index (d') for slow tempo trials | | | | |
| **Variable name** | | BAT_slow_dprime | | | | |
| **Age and gender effects** | | | | | | |
| Age regression (slope) | | -0.005 | | | | |
| Age regression (p) | | 0.109 | | | | |
| Age group (p) | | 0.351 | | | | |
| Gender (p) | | 0.152 | | | | |
| **Group** | | All | Age 18 to 21 | Age 22 to 29 | Age 30 to 54 | Age 55 to 87 |
| N (N female) | | 108 (74) | 27 (18) | 29 (19) | 26 (16) | 26 (21) |
| **Normality** | | | | | | |
| Skewness | | -0.83 | -1.00 | -0.83 | -1.05 | -0.20 |
| Excess kurtosis | | -0.18 | -0.03 | 0.17 | 0.05 | -1.33 |
| **Scores** | | | | | | |
| Mean | | 2.65 | 2.83 | 2.73 | 2.62 | 2.42 |
| SD | | 0.91 | 0.86 | 0.83 | 1.08 | 0.86 |
| Percentiles | | | | | | |
| | 0 | -0.15 | 0.59 | 0.44 | -0.15 | 1.02 |
| | 1 | 0.45 | 0.78 | 0.68 | 0.04 | 1.04 |
| | 2 | 0.59 | 0.96 | 0.92 | 0.22 | 1.06 |
| | 3 | 0.68 | 1.15 | 1.17 | 0.40 | 1.08 |
| | 4 | 1.02 | 1.32 | 1.32 | 0.59 | 1.09 |
| | 5 | 1.05 | 1.36 | 1.37 | 0.70 | 1.15 |
| | 6 | 1.12 | 1.40 | 1.42 | 0.80 | 1.20 |
| | 7 | 1.20 | 1.44 | 1.46 | 0.91 | 1.26 |
| | 8 | 1.27 | 1.49 | 1.58 | 1.02 | 1.31 |
| | 9 | 1.31 | 1.57 | 1.70 | 1.06 | 1.32 |
| | 10 | 1.31 | 1.65 | 1.82 | 1.09 | 1.32 |
| | 11 | 1.32 | 1.73 | 1.91 | 1.13 | 1.32 |
| | 12 | 1.32 | 1.78 | 1.92 | 1.17 | 1.32 |
| | 13 | 1.46 | 1.79 | 1.92 | 1.18 | 1.32 |
| | 14 | 1.47 | 1.80 | 1.92 | 1.20 | 1.32 |
| | 15 | 1.47 | 1.82 | 1.94 | 1.21 | 1.32 |
| | 16 | 1.49 | 1.86 | 1.98 | 1.23 | 1.32 |
| | 17 | 1.66 | 1.91 | 2.01 | 1.36 | 1.36 |
| | 18 | 1.77 | 1.97 | 2.04 | 1.50 | 1.39 |
| | 19 | 1.79 | 2.03 | 2.04 | 1.63 | 1.43 |
| | 20 | 1.86 | 2.08 | 2.04 | 1.77 | 1.47 |
| | 21 | 1.92 | 2.12 | 2.04 | 1.84 | 1.47 |
| | 22 | 1.98 | 2.17 | 2.04 | 1.92 | 1.47 |
| | 23 | 2.04 | 2.21 | 2.05 | 1.99 | 1.47 |
| | 24 | 2.04 | 2.22 | 2.06 | 2.06 | 1.48 |

| | | | | | |
|---|---|---|---|---|---|
| 25 | 2.06 | 2.22 | 2.06 | 2.10 | 1.52 |
| 26 | 2.06 | 2.22 | 2.15 | 2.14 | 1.56 |
| 27 | 2.06 | 2.22 | 2.24 | 2.18 | 1.60 |
| 28 | 2.06 | 2.32 | 2.33 | 2.22 | 1.64 |
| 29 | 2.07 | 2.41 | 2.38 | 2.26 | 1.74 |
| 30 | 2.22 | 2.50 | 2.38 | 2.30 | 1.85 |
| 31 | 2.22 | 2.58 | 2.38 | 2.34 | 1.96 |
| 32 | 2.26 | 2.64 | 2.38 | 2.38 | 2.06 |
| 33 | 2.38 | 2.70 | 2.43 | 2.43 | 2.06 |
| 34 | 2.38 | 2.75 | 2.48 | 2.48 | 2.06 |
| 35 | 2.47 | 2.79 | 2.53 | 2.52 | 2.06 |
| 36 | 2.57 | 2.79 | 2.57 | 2.57 | 2.06 |
| 37 | 2.57 | 2.79 | 2.57 | 2.62 | 2.06 |
| 38 | 2.57 | 2.79 | 2.57 | 2.68 | 2.06 |
| 39 | 2.57 | 2.83 | 2.57 | 2.73 | 2.06 |
| 40 | 2.57 | 2.90 | 2.61 | 2.79 | 2.06 |
| 41 | 2.76 | 2.98 | 2.68 | 2.79 | 2.19 |
| 42 | 2.79 | 3.05 | 2.74 | 2.79 | 2.32 |
| 43 | 2.79 | 3.07 | 2.79 | 2.79 | 2.44 |
| 44 | 2.79 | 3.07 | 2.79 | 2.79 | 2.57 |
| 45 | 2.79 | 3.07 | 2.79 | 2.86 | 2.57 |
| 46 | 2.79 | 3.07 | 2.79 | 2.93 | 2.57 |
| 47 | 2.79 | 3.07 | 2.79 | 3.00 | 2.57 |
| 48 | 2.79 | 3.07 | 2.79 | 3.07 | 2.57 |
| 49 | 2.79 | 3.07 | 2.79 | 3.07 | 2.62 |
| 50 | 2.79 | 3.07 | 2.79 | 3.07 | 2.68 |
| 51 | 2.79 | 3.07 | 2.79 | 3.07 | 2.73 |
| 52 | 2.79 | 3.07 | 2.79 | 3.07 | 2.79 |
| 53 | 2.79 | 3.07 | 2.79 | 3.07 | 2.79 |
| 54 | 3.01 | 3.09 | 2.79 | 3.07 | 2.79 |
| 55 | 3.07 | 3.21 | 2.79 | 3.07 | 2.79 |
| 56 | 3.07 | 3.33 | 2.79 | 3.07 | 2.79 |
| 57 | 3.07 | 3.45 | 2.79 | 3.07 | 2.79 |
| 58 | 3.07 | 3.54 | 2.86 | 3.07 | 2.79 |
| 59 | 3.07 | 3.54 | 2.94 | 3.07 | 2.79 |
| 60 | 3.07 | 3.54 | 3.02 | 3.07 | 2.79 |
| 61 | 3.07 | 3.54 | 3.11 | 3.19 | 2.79 |
| 62 | 3.07 | 3.54 | 3.24 | 3.31 | 2.79 |
| 63 | 3.07 | 3.54 | 3.37 | 3.42 | 2.79 |
| 64 | 3.07 | 3.54 | 3.50 | 3.54 | 2.79 |
| 65 | 3.33 | 3.54 | 3.54 | 3.54 | 2.79 |
| 66 | 3.54 | 3.54 | 3.54 | 3.54 | 2.79 |
| 67 | 3.54 | 3.54 | 3.54 | 3.54 | 2.79 |
| 68 | 3.54 | 3.54 | 3.54 | 3.54 | 2.79 |

| | | | | | |
|---|---|---|---|---|---|
| 69 | 3.54 | 3.54 | 3.54 | 3.54 | 2.86 |
| 70 | 3.54 | 3.54 | 3.54 | 3.54 | 2.93 |
| 71 | 3.54 | 3.54 | 3.54 | 3.54 | 3.00 |
| 72 | 3.54 | 3.54 | 3.54 | 3.54 | 3.07 |
| 73 | 3.54 | 3.54 | 3.54 | 3.54 | 3.07 |
| 74 | 3.54 | 3.54 | 3.54 | 3.54 | 3.07 |
| 75 | 3.54 | 3.54 | 3.54 | 3.54 | 3.07 |
| 76 | 3.54 | 3.54 | 3.54 | 3.54 | 3.07 |
| 77 | 3.54 | 3.54 | 3.54 | 3.54 | 3.07 |
| 78 | 3.54 | 3.54 | 3.54 | 3.54 | 3.07 |
| 79 | 3.54 | 3.54 | 3.54 | 3.54 | 3.07 |
| 80 | 3.54 | 3.54 | 3.54 | 3.54 | 3.07 |
| 81 | 3.54 | 3.54 | 3.54 | 3.54 | 3.19 |
| 82 | 3.54 | 3.54 | 3.54 | 3.54 | 3.31 |
| 83 | 3.54 | 3.54 | 3.54 | 3.54 | 3.42 |
| 84 | 3.54 | 3.54 | 3.54 | 3.54 | 3.54 |
| 85 | 3.54 | 3.54 | 3.54 | 3.54 | 3.54 |
| 86 | 3.54 | 3.54 | 3.54 | 3.54 | 3.54 |
| 87 | 3.54 | 3.54 | 3.54 | 3.54 | 3.54 |
| 88 | 3.54 | 3.54 | 3.54 | 3.54 | 3.54 |
| 89 | 3.54 | 3.54 | 3.54 | 3.54 | 3.54 |
| 90 | 3.54 | 3.54 | 3.54 | 3.54 | 3.54 |
| 91 | 3.54 | 3.54 | 3.54 | 3.54 | 3.54 |
| 92 | 3.54 | 3.54 | 3.54 | 3.54 | 3.54 |
| 93 | 3.54 | 3.54 | 3.54 | 3.54 | 3.54 |
| 94 | 3.54 | 3.54 | 3.54 | 3.54 | 3.54 |
| 95 | 3.54 | 3.54 | 3.54 | 3.54 | 3.54 |
| 96 | 3.54 | 3.54 | 3.54 | 3.54 | 3.54 |
| 97 | 3.54 | 3.54 | 3.54 | 3.54 | 3.54 |
| 98 | 3.54 | 3.54 | 3.54 | 3.54 | 3.54 |
| 99 | 3.54 | 3.54 | 3.54 | 3.54 | 3.54 |
| 100 | 3.54 | 3.54 | 3.54 | 3.54 | 3.54 |

***Table S18:*** *Beat Alignment Test – Hit rate for all trials – BAT_all_hits*

| | | | | | |
|---|---|---|---|---|---|
| **Task** | Beat Alignment Test | | | | |
| **Outcome measure** | Hit rate for all trials | | | | |
| **Variable name** | BAT_all_hits | | | | |
| **Age and gender effects** | | | | | |
| Age regression (slope) | -0.157 | | | | |
| Age regression (p) | 0.020 | | | | |
| Age group (p) | 0.222 | | | | |
| Gender (p) | 0.390 | | | | |
| **Group** | All | Age 18 to 21 | Age 22 to 29 | Age 30 to 54 | Age 55 to 87 |
| N (N female) | 108 (74) | 27 (18) | 29 (19) | 26 (16) | 26 (21) |
| **Normality** | | | | | |
| Skewness | -0.89 | -1.14 | -0.70 | -0.58 | -0.76 |
| Excess kurtosis | -0.19 | 0.12 | -0.95 | -1.20 | -0.22 |
| **Scores** | | | | | |
| Mean | 82.909 | 86.265 | 86.710 | 81.010 | 77.083 |
| SD | 17.013 | 14.915 | 13.756 | 19.124 | 19.021 |
| Percentiles | | | | | |
| 0 | 31.250 | 50.000 | 60.417 | 47.917 | 31.250 |
| 1 | 42.104 | 51.625 | 60.417 | 47.917 | 33.854 |
| 2 | 47.917 | 53.250 | 60.417 | 47.917 | 36.458 |
| 3 | 47.917 | 54.875 | 60.417 | 47.917 | 39.063 |
| 4 | 48.500 | 56.250 | 60.917 | 47.917 | 41.667 |
| 5 | 50.000 | 56.250 | 62.083 | 48.438 | 43.229 |
| 6 | 50.000 | 56.250 | 63.250 | 48.958 | 44.792 |
| 7 | 52.042 | 56.250 | 64.417 | 49.479 | 46.354 |
| 8 | 54.167 | 57.083 | 65.083 | 50.000 | 47.917 |
| 9 | 55.479 | 59.792 | 65.667 | 51.042 | 48.438 |
| 10 | 56.250 | 62.500 | 66.250 | 52.083 | 48.958 |
| 11 | 57.854 | 65.208 | 66.667 | 53.125 | 49.479 |
| 12 | 60.083 | 66.917 | 66.667 | 54.167 | 50.000 |
| 13 | 60.417 | 67.458 | 66.667 | 54.167 | 52.083 |
| 14 | 60.417 | 68.000 | 66.667 | 54.167 | 54.167 |
| 15 | 62.604 | 68.542 | 67.083 | 54.167 | 56.250 |
| 16 | 64.833 | 69.750 | 67.667 | 54.167 | 58.333 |
| 17 | 66.667 | 71.375 | 68.250 | 55.729 | 60.417 |
| 18 | 66.667 | 73.000 | 68.833 | 57.292 | 62.500 |
| 19 | 66.667 | 74.625 | 69.417 | 58.854 | 64.583 |
| 20 | 66.667 | 75.417 | 70.000 | 60.417 | 66.667 |
| 21 | 67.646 | 75.958 | 70.583 | 60.938 | 67.188 |
| 22 | 68.750 | 76.500 | 72.167 | 61.458 | 67.708 |
| 23 | 68.750 | 77.042 | 74.500 | 61.979 | 68.229 |
| 24 | 68.750 | 78.083 | 76.833 | 62.500 | 68.750 |

| | | | | | |
|---|---|---|---|---|---|
| 25 | 70.313 | 79.167 | 79.167 | 63.542 | 68.750 |
| 26 | 70.833 | 80.250 | 80.333 | 64.583 | 68.750 |
| 27 | 70.833 | 81.333 | 81.500 | 65.625 | 68.750 |
| 28 | 74.833 | 82.417 | 82.667 | 66.667 | 68.750 |
| 29 | 75.063 | 83.500 | 83.333 | 68.750 | 69.271 |
| 30 | 77.083 | 84.583 | 83.333 | 70.833 | 69.792 |
| 31 | 77.083 | 85.542 | 83.333 | 72.917 | 70.313 |
| 32 | 77.083 | 86.083 | 83.333 | 75.000 | 70.833 |
| 33 | 77.729 | 86.625 | 83.333 | 75.521 | 70.833 |
| 34 | 79.167 | 87.167 | 83.333 | 76.042 | 70.833 |
| 35 | 79.167 | 87.500 | 83.333 | 76.563 | 70.833 |
| 36 | 80.250 | 87.500 | 83.500 | 77.083 | 70.833 |
| 37 | 81.250 | 87.500 | 84.083 | 77.604 | 72.396 |
| 38 | 82.625 | 87.500 | 84.667 | 78.125 | 73.958 |
| 39 | 83.333 | 87.792 | 85.250 | 78.646 | 75.521 |
| 40 | 83.333 | 88.333 | 85.417 | 79.167 | 77.083 |
| 41 | 83.333 | 88.875 | 85.417 | 80.729 | 77.083 |
| 42 | 85.292 | 89.417 | 85.417 | 82.292 | 77.083 |
| 43 | 85.417 | 89.583 | 85.583 | 83.854 | 77.083 |
| 44 | 85.417 | 89.583 | 86.750 | 85.417 | 77.083 |
| 45 | 85.417 | 89.583 | 87.917 | 85.417 | 77.604 |
| 46 | 85.417 | 89.583 | 89.083 | 85.417 | 78.125 |
| 47 | 86.021 | 90.042 | 89.917 | 85.417 | 78.646 |
| 48 | 87.500 | 90.583 | 90.500 | 85.417 | 79.167 |
| 49 | 87.500 | 91.125 | 91.083 | 85.938 | 79.688 |
| 50 | 87.500 | 91.667 | 91.667 | 86.458 | 80.208 |
| 51 | 87.500 | 92.208 | 92.250 | 86.979 | 80.729 |
| 52 | 87.500 | 92.750 | 92.833 | 87.500 | 81.250 |
| 53 | 88.979 | 93.292 | 93.417 | 89.063 | 81.771 |
| 54 | 89.583 | 93.750 | 94.000 | 90.625 | 82.292 |
| 55 | 89.583 | 93.750 | 94.583 | 92.188 | 82.813 |
| 56 | 91.500 | 93.750 | 95.167 | 93.750 | 83.333 |
| 57 | 91.667 | 93.750 | 95.750 | 93.750 | 83.854 |
| 58 | 93.750 | 93.917 | 95.833 | 93.750 | 84.375 |
| 59 | 93.750 | 94.458 | 95.833 | 93.750 | 84.896 |
| 60 | 93.750 | 95.000 | 95.833 | 93.750 | 85.417 |
| 61 | 93.750 | 95.542 | 95.833 | 93.750 | 85.938 |
| 62 | 93.750 | 95.833 | 95.833 | 93.750 | 86.458 |
| 63 | 94.604 | 95.833 | 95.833 | 93.750 | 86.979 |
| 64 | 95.833 | 95.833 | 95.833 | 93.750 | 87.500 |
| 65 | 95.833 | 95.833 | 95.833 | 94.271 | 87.500 |
| 66 | 95.833 | 95.833 | 95.833 | 94.792 | 87.500 |
| 67 | 95.833 | 95.833 | 95.833 | 95.313 | 87.500 |
| 68 | 95.833 | 95.833 | 95.917 | 95.833 | 87.500 |

| | | | | | |
|---|---|---|---|---|---|
| 69 | 95.833 | 95.833 | 96.500 | 96.354 | 87.500 |
| 70 | 95.833 | 96.250 | 97.083 | 96.875 | 87.500 |
| 71 | 95.833 | 96.792 | 97.667 | 97.396 | 87.500 |
| 72 | 95.833 | 97.333 | 98.250 | 97.917 | 87.500 |
| 73 | 96.063 | 97.875 | 98.833 | 97.917 | 89.583 |
| 74 | 97.917 | 97.917 | 99.417 | 97.917 | 91.667 |
| 75 | 97.917 | 97.917 | 100.000 | 97.917 | 93.750 |
| 76 | 97.917 | 97.917 | 100.000 | 97.917 | 95.833 |
| 77 | 97.917 | 97.917 | 100.000 | 98.438 | 95.833 |
| 78 | 97.917 | 97.917 | 100.000 | 98.958 | 95.833 |
| 79 | 97.917 | 97.917 | 100.000 | 99.479 | 95.833 |
| 80 | 97.917 | 97.917 | 100.000 | 100.000 | 95.833 |
| 81 | 97.917 | 97.917 | 100.000 | 100.000 | 95.833 |
| 82 | 99.458 | 97.917 | 100.000 | 100.000 | 95.833 |
| 83 | 100.000 | 97.917 | 100.000 | 100.000 | 95.833 |
| 84 | 100.000 | 97.917 | 100.000 | 100.000 | 95.833 |
| 85 | 100.000 | 98.125 | 100.000 | 100.000 | 96.354 |
| 86 | 100.000 | 98.667 | 100.000 | 100.000 | 96.875 |
| 87 | 100.000 | 99.208 | 100.000 | 100.000 | 97.396 |
| 88 | 100.000 | 99.750 | 100.000 | 100.000 | 97.917 |
| 89 | 100.000 | 100.000 | 100.000 | 100.000 | 97.917 |
| 90 | 100.000 | 100.000 | 100.000 | 100.000 | 97.917 |
| 91 | 100.000 | 100.000 | 100.000 | 100.000 | 97.917 |
| 92 | 100.000 | 100.000 | 100.000 | 100.000 | 97.917 |
| 93 | 100.000 | 100.000 | 100.000 | 100.000 | 98.438 |
| 94 | 100.000 | 100.000 | 100.000 | 100.000 | 98.958 |
| 95 | 100.000 | 100.000 | 100.000 | 100.000 | 99.479 |
| 96 | 100.000 | 100.000 | 100.000 | 100.000 | 100.000 |
| 97 | 100.000 | 100.000 | 100.000 | 100.000 | 100.000 |
| 98 | 100.000 | 100.000 | 100.000 | 100.000 | 100.000 |
| 99 | 100.000 | 100.000 | 100.000 | 100.000 | 100.000 |
| 100 | 100.000 | 100.000 | 100.000 | 100.000 | 100.000 |

***Table S19:*** *Beat Alignment Test – Hit rate for fast tempo trials – BAT_fast_hits*

| | | | | | |
|---|---|---|---|---|---|
| **Task** | Beat Alignment Test | | | | |
| **Outcome measure** | Hit rate for fast tempo trials | | | | |
| **Variable name** | BAT_fast_hits | | | | |
| **Age and gender effects** | | | | | |
| Age regression (slope) | -0.223 | | | | |
| Age regression (p) | 0.010 | | | | |
| Age group (p) | 0.361 | | | | |
| Gender (p) | 1.000 | | | | |
| **Group** | All | Age 18 to 21 | Age 22 to 29 | Age 30 to 54 | Age 55 to 87 |
| N (N female) | 108 (74) | 27 (18) | 29 (19) | 26 (16) | 26 (21) |
| **Normality** | | | | | |
| Skewness | -0.88 | -1.07 | -0.96 | -0.54 | -0.50 |
| Excess kurtosis | -0.09 | 0.42 | -0.20 | -1.07 | -0.54 |
| **Scores** | | | | | |
| Mean | 80.729 | 84.954 | 85.776 | 78.125 | 73.317 |
| SD | 19.204 | 14.938 | 15.571 | 21.232 | 22.608 |
| Percentiles | | | | | |
| 0 | 18.750 | 43.750 | 50.000 | 37.500 | 18.750 |
| 1 | 37.500 | 48.625 | 50.000 | 39.063 | 23.438 |
| 2 | 38.375 | 53.500 | 50.000 | 40.625 | 28.125 |
| 3 | 43.750 | 58.375 | 50.000 | 42.188 | 32.813 |
| 4 | 43.750 | 62.500 | 51.500 | 43.750 | 37.500 |
| 5 | 43.750 | 62.500 | 55.000 | 43.750 | 39.063 |
| 6 | 46.375 | 62.500 | 58.500 | 43.750 | 40.625 |
| 7 | 50.000 | 62.500 | 62.000 | 43.750 | 42.188 |
| 8 | 50.000 | 62.500 | 62.500 | 43.750 | 43.750 |
| 9 | 50.000 | 62.500 | 62.500 | 45.313 | 45.313 |
| 10 | 50.000 | 62.500 | 62.500 | 46.875 | 46.875 |
| 11 | 50.000 | 62.500 | 63.000 | 48.438 | 48.438 |
| 12 | 50.000 | 62.500 | 64.750 | 50.000 | 50.000 |
| 13 | 55.688 | 62.500 | 66.500 | 50.000 | 50.000 |
| 14 | 56.250 | 62.500 | 68.250 | 50.000 | 50.000 |
| 15 | 62.500 | 62.500 | 68.750 | 50.000 | 50.000 |
| 16 | 62.500 | 64.500 | 68.750 | 50.000 | 50.000 |
| 17 | 62.500 | 67.750 | 68.750 | 50.000 | 51.563 |
| 18 | 62.500 | 71.000 | 69.000 | 50.000 | 53.125 |
| 19 | 62.500 | 74.250 | 70.750 | 50.000 | 54.688 |
| 20 | 62.500 | 76.250 | 72.500 | 50.000 | 56.250 |
| 21 | 62.500 | 77.875 | 74.250 | 53.125 | 56.250 |
| 22 | 62.500 | 79.500 | 75.000 | 56.250 | 56.250 |
| 23 | 62.500 | 81.125 | 75.000 | 59.375 | 56.250 |
| 24 | 62.500 | 81.250 | 75.000 | 62.500 | 56.250 |

| | | | | | |
|---|---|---|---|---|---|
| 25 | 67.188 | 81.250 | 75.000 | 64.063 | 57.813 |
| 26 | 68.750 | 81.250 | 76.750 | 65.625 | 59.375 |
| 27 | 68.750 | 81.250 | 78.500 | 67.188 | 60.938 |
| 28 | 68.750 | 81.250 | 80.250 | 68.750 | 62.500 |
| 29 | 75.000 | 81.250 | 81.250 | 70.313 | 62.500 |
| 30 | 75.000 | 81.250 | 81.250 | 71.875 | 62.500 |
| 31 | 75.000 | 81.250 | 81.250 | 73.438 | 62.500 |
| 32 | 75.000 | 81.250 | 81.250 | 75.000 | 62.500 |
| 33 | 75.000 | 81.250 | 81.250 | 75.000 | 62.500 |
| 34 | 75.000 | 81.250 | 81.250 | 75.000 | 62.500 |
| 35 | 75.000 | 81.250 | 81.250 | 75.000 | 62.500 |
| 36 | 78.250 | 81.250 | 81.750 | 75.000 | 62.500 |
| 37 | 81.250 | 81.250 | 83.500 | 75.000 | 62.500 |
| 38 | 81.250 | 81.250 | 85.250 | 75.000 | 62.500 |
| 39 | 81.250 | 82.125 | 87.000 | 75.000 | 62.500 |
| 40 | 81.250 | 83.750 | 87.500 | 75.000 | 62.500 |
| 41 | 81.250 | 85.375 | 87.500 | 75.000 | 64.063 |
| 42 | 81.250 | 87.000 | 87.500 | 75.000 | 65.625 |
| 43 | 81.250 | 87.500 | 87.750 | 75.000 | 67.188 |
| 44 | 81.250 | 87.500 | 89.500 | 75.000 | 68.750 |
| 45 | 81.250 | 87.500 | 91.250 | 76.563 | 70.313 |
| 46 | 81.250 | 87.500 | 93.000 | 78.125 | 71.875 |
| 47 | 81.250 | 87.500 | 93.750 | 79.688 | 73.438 |
| 48 | 83.500 | 87.500 | 93.750 | 81.250 | 75.000 |
| 49 | 87.500 | 87.500 | 93.750 | 81.250 | 76.563 |
| 50 | 87.500 | 87.500 | 93.750 | 81.250 | 78.125 |
| 51 | 87.500 | 89.125 | 93.750 | 81.250 | 79.688 |
| 52 | 87.500 | 90.750 | 93.750 | 81.250 | 81.250 |
| 53 | 87.500 | 92.375 | 93.750 | 82.813 | 81.250 |
| 54 | 87.500 | 93.750 | 93.750 | 84.375 | 81.250 |
| 55 | 87.500 | 93.750 | 93.750 | 85.938 | 81.250 |
| 56 | 87.500 | 93.750 | 93.750 | 87.500 | 81.250 |
| 57 | 87.500 | 93.750 | 93.750 | 87.500 | 81.250 |
| 58 | 93.750 | 93.750 | 93.750 | 87.500 | 81.250 |
| 59 | 93.750 | 93.750 | 93.750 | 87.500 | 81.250 |
| 60 | 93.750 | 93.750 | 93.750 | 87.500 | 81.250 |
| 61 | 93.750 | 93.750 | 93.750 | 89.063 | 82.813 |
| 62 | 93.750 | 93.750 | 93.750 | 90.625 | 84.375 |
| 63 | 93.750 | 93.750 | 93.750 | 92.188 | 85.938 |
| 64 | 93.750 | 93.750 | 93.750 | 93.750 | 87.500 |
| 65 | 93.750 | 93.750 | 93.750 | 93.750 | 87.500 |
| 66 | 93.750 | 93.750 | 93.750 | 93.750 | 87.500 |
| 67 | 93.750 | 93.750 | 93.750 | 93.750 | 87.500 |
| 68 | 93.750 | 93.750 | 94.000 | 93.750 | 87.500 |

| | | | | | |
|---|---|---|---|---|---|
| 69 | 93.750 | 93.750 | 95.750 | 95.313 | 87.500 |
| 70 | 93.750 | 93.750 | 97.500 | 96.875 | 87.500 |
| 71 | 93.750 | 93.750 | 99.250 | 98.438 | 87.500 |
| 72 | 93.750 | 93.750 | 100.000 | 100.000 | 87.500 |
| 73 | 94.438 | 93.750 | 100.000 | 100.000 | 89.063 |
| 74 | 100.000 | 93.750 | 100.000 | 100.000 | 90.625 |
| 75 | 100.000 | 93.750 | 100.000 | 100.000 | 92.188 |
| 76 | 100.000 | 93.750 | 100.000 | 100.000 | 93.750 |
| 77 | 100.000 | 93.875 | 100.000 | 100.000 | 95.313 |
| 78 | 100.000 | 95.500 | 100.000 | 100.000 | 96.875 |
| 79 | 100.000 | 97.125 | 100.000 | 100.000 | 98.438 |
| 80 | 100.000 | 98.750 | 100.000 | 100.000 | 100.000 |
| 81 | 100.000 | 100.000 | 100.000 | 100.000 | 100.000 |
| 82 | 100.000 | 100.000 | 100.000 | 100.000 | 100.000 |
| 83 | 100.000 | 100.000 | 100.000 | 100.000 | 100.000 |
| 84 | 100.000 | 100.000 | 100.000 | 100.000 | 100.000 |
| 85 | 100.000 | 100.000 | 100.000 | 100.000 | 100.000 |
| 86 | 100.000 | 100.000 | 100.000 | 100.000 | 100.000 |
| 87 | 100.000 | 100.000 | 100.000 | 100.000 | 100.000 |
| 88 | 100.000 | 100.000 | 100.000 | 100.000 | 100.000 |
| 89 | 100.000 | 100.000 | 100.000 | 100.000 | 100.000 |
| 90 | 100.000 | 100.000 | 100.000 | 100.000 | 100.000 |
| 91 | 100.000 | 100.000 | 100.000 | 100.000 | 100.000 |
| 92 | 100.000 | 100.000 | 100.000 | 100.000 | 100.000 |
| 93 | 100.000 | 100.000 | 100.000 | 100.000 | 100.000 |
| 94 | 100.000 | 100.000 | 100.000 | 100.000 | 100.000 |
| 95 | 100.000 | 100.000 | 100.000 | 100.000 | 100.000 |
| 96 | 100.000 | 100.000 | 100.000 | 100.000 | 100.000 |
| 97 | 100.000 | 100.000 | 100.000 | 100.000 | 100.000 |
| 98 | 100.000 | 100.000 | 100.000 | 100.000 | 100.000 |
| 99 | 100.000 | 100.000 | 100.000 | 100.000 | 100.000 |
| 100 | 100.000 | 100.000 | 100.000 | 100.000 | 100.000 |

***Table S20:*** *Beat Alignment Test – Hit rate for medium tempo trials – BAT_med_hits*

| | | | | | |
|---|---|---|---|---|---|
| **Task** | Beat Alignment Test | | | | |
| **Outcome measure** | Hit rate for medium tempo trials | | | | |
| **Variable name** | BAT_med_hits | | | | |
| **Age and gender effects** | | | | | |
| Age regression (slope) | -0.128 | | | | |
| Age regression (p) | 0.066 | | | | |
| Age group (p) | 0.581 | | | | |
| Gender (p) | 0.035 | | | | |
| **Group** | All | Age 18 to 21 | Age 22 to 29 | Age 30 to 54 | Age 55 to 87 |
| N (N female) | 108 (74) | 27 (18) | 29 (19) | 26 (16) | 26 (21) |
| **Normality** | | | | | |
| Skewness | -0.93 | -1.79 | -0.68 | -0.55 | -0.58 |
| Excess kurtosis | -0.07 | 2.43 | -0.88 | -1.08 | -0.74 |
| **Scores** | | | | | |
| Mean | 82.118 | 85.185 | 84.267 | 80.288 | 78.365 |
| SD | 18.435 | 19.469 | 16.076 | 20.209 | 18.134 |
| Percentiles | | | | | |
| 0 | 25.000 | 25.000 | 50.000 | 37.500 | 43.750 |
| 1 | 37.938 | 29.875 | 51.750 | 40.625 | 43.750 |
| 2 | 43.750 | 34.750 | 53.500 | 43.750 | 43.750 |
| 3 | 43.750 | 39.625 | 55.250 | 46.875 | 43.750 |
| 4 | 43.750 | 43.750 | 56.250 | 50.000 | 43.750 |
| 5 | 43.750 | 43.750 | 56.250 | 50.000 | 43.750 |
| 6 | 46.375 | 43.750 | 56.250 | 50.000 | 43.750 |
| 7 | 50.000 | 43.750 | 56.250 | 50.000 | 43.750 |
| 8 | 50.000 | 45.750 | 57.750 | 50.000 | 43.750 |
| 9 | 53.938 | 52.250 | 59.500 | 51.563 | 46.875 |
| 10 | 56.250 | 58.750 | 61.250 | 53.125 | 50.000 |
| 11 | 56.250 | 65.250 | 62.500 | 54.688 | 53.125 |
| 12 | 56.250 | 69.500 | 62.500 | 56.250 | 56.250 |
| 13 | 56.250 | 71.125 | 62.500 | 56.250 | 57.813 |
| 14 | 56.250 | 72.750 | 62.500 | 56.250 | 59.375 |
| 15 | 62.500 | 74.375 | 63.750 | 56.250 | 60.938 |
| 16 | 62.500 | 75.000 | 65.500 | 56.250 | 62.500 |
| 17 | 62.500 | 75.000 | 67.250 | 56.250 | 62.500 |
| 18 | 62.500 | 75.000 | 68.750 | 56.250 | 62.500 |
| 19 | 62.500 | 75.000 | 68.750 | 56.250 | 62.500 |
| 20 | 65.000 | 76.250 | 68.750 | 56.250 | 62.500 |
| 21 | 68.750 | 77.875 | 68.750 | 57.813 | 64.063 |
| 22 | 68.750 | 79.500 | 68.750 | 59.375 | 65.625 |
| 23 | 68.750 | 81.125 | 68.750 | 60.938 | 67.188 |
| 24 | 68.750 | 81.250 | 68.750 | 62.500 | 68.750 |

| | | | | | |
|---|---|---|---|---|---|
| 25 | 68.750 | 81.250 | 68.750 | 62.500 | 68.750 |
| 26 | 68.750 | 81.250 | 70.500 | 62.500 | 68.750 |
| 27 | 74.313 | 81.375 | 72.250 | 62.500 | 68.750 |
| 28 | 75.000 | 83.000 | 74.000 | 62.500 | 68.750 |
| 29 | 75.000 | 84.625 | 75.750 | 65.625 | 68.750 |
| 30 | 75.000 | 86.250 | 77.500 | 68.750 | 68.750 |
| 31 | 75.000 | 87.500 | 79.250 | 71.875 | 68.750 |
| 32 | 75.000 | 87.500 | 81.000 | 75.000 | 68.750 |
| 33 | 75.000 | 87.500 | 81.250 | 75.000 | 70.313 |
| 34 | 75.000 | 87.500 | 81.250 | 75.000 | 71.875 |
| 35 | 77.813 | 87.500 | 81.250 | 75.000 | 73.438 |
| 36 | 81.250 | 87.500 | 81.250 | 75.000 | 75.000 |
| 37 | 81.250 | 87.500 | 81.250 | 75.000 | 75.000 |
| 38 | 81.250 | 87.500 | 81.250 | 75.000 | 75.000 |
| 39 | 81.250 | 87.500 | 81.250 | 75.000 | 75.000 |
| 40 | 81.250 | 87.500 | 82.500 | 75.000 | 75.000 |
| 41 | 81.250 | 87.500 | 84.250 | 76.563 | 75.000 |
| 42 | 81.250 | 87.500 | 86.000 | 78.125 | 75.000 |
| 43 | 81.313 | 88.625 | 87.500 | 79.688 | 75.000 |
| 44 | 87.500 | 90.250 | 87.500 | 81.250 | 75.000 |
| 45 | 87.500 | 91.875 | 87.500 | 81.250 | 76.563 |
| 46 | 87.500 | 93.500 | 87.500 | 81.250 | 78.125 |
| 47 | 87.500 | 93.750 | 87.500 | 81.250 | 79.688 |
| 48 | 87.500 | 93.750 | 87.500 | 81.250 | 81.250 |
| 49 | 87.500 | 93.750 | 87.500 | 82.813 | 81.250 |
| 50 | 87.500 | 93.750 | 87.500 | 84.375 | 81.250 |
| 51 | 87.500 | 93.750 | 89.250 | 85.938 | 81.250 |
| 52 | 87.500 | 93.750 | 91.000 | 87.500 | 81.250 |
| 53 | 87.500 | 93.750 | 92.750 | 89.063 | 82.813 |
| 54 | 92.375 | 93.750 | 93.750 | 90.625 | 84.375 |
| 55 | 93.750 | 93.750 | 93.750 | 92.188 | 85.938 |
| 56 | 93.750 | 93.750 | 93.750 | 93.750 | 87.500 |
| 57 | 93.750 | 93.750 | 93.750 | 93.750 | 87.500 |
| 58 | 93.750 | 93.750 | 93.750 | 93.750 | 87.500 |
| 59 | 93.750 | 93.750 | 93.750 | 93.750 | 87.500 |
| 60 | 93.750 | 93.750 | 93.750 | 93.750 | 87.500 |
| 61 | 93.750 | 93.750 | 93.750 | 93.750 | 87.500 |
| 62 | 93.750 | 93.750 | 93.750 | 93.750 | 87.500 |
| 63 | 93.750 | 93.750 | 93.750 | 93.750 | 87.500 |
| 64 | 93.750 | 93.750 | 93.750 | 93.750 | 87.500 |
| 65 | 93.750 | 93.750 | 93.750 | 95.313 | 89.063 |
| 66 | 93.750 | 93.750 | 93.750 | 96.875 | 90.625 |
| 67 | 93.750 | 93.750 | 93.750 | 98.438 | 92.188 |
| 68 | 93.750 | 93.750 | 94.000 | 100.000 | 93.750 |

| | | | | | |
|---|---|---|---|---|---|
| 69 | 93.750 | 93.750 | 95.750 | 100.000 | 93.750 |
| 70 | 93.750 | 95.000 | 97.500 | 100.000 | 93.750 |
| 71 | 93.750 | 96.625 | 99.250 | 100.000 | 93.750 |
| 72 | 94.000 | 98.250 | 100.000 | 100.000 | 93.750 |
| 73 | 100.000 | 99.875 | 100.000 | 100.000 | 93.750 |
| 74 | 100.000 | 100.000 | 100.000 | 100.000 | 93.750 |
| 75 | 100.000 | 100.000 | 100.000 | 100.000 | 93.750 |
| 76 | 100.000 | 100.000 | 100.000 | 100.000 | 93.750 |
| 77 | 100.000 | 100.000 | 100.000 | 100.000 | 93.750 |
| 78 | 100.000 | 100.000 | 100.000 | 100.000 | 93.750 |
| 79 | 100.000 | 100.000 | 100.000 | 100.000 | 93.750 |
| 80 | 100.000 | 100.000 | 100.000 | 100.000 | 93.750 |
| 81 | 100.000 | 100.000 | 100.000 | 100.000 | 93.750 |
| 82 | 100.000 | 100.000 | 100.000 | 100.000 | 93.750 |
| 83 | 100.000 | 100.000 | 100.000 | 100.000 | 93.750 |
| 84 | 100.000 | 100.000 | 100.000 | 100.000 | 93.750 |
| 85 | 100.000 | 100.000 | 100.000 | 100.000 | 95.313 |
| 86 | 100.000 | 100.000 | 100.000 | 100.000 | 96.875 |
| 87 | 100.000 | 100.000 | 100.000 | 100.000 | 98.438 |
| 88 | 100.000 | 100.000 | 100.000 | 100.000 | 100.000 |
| 89 | 100.000 | 100.000 | 100.000 | 100.000 | 100.000 |
| 90 | 100.000 | 100.000 | 100.000 | 100.000 | 100.000 |
| 91 | 100.000 | 100.000 | 100.000 | 100.000 | 100.000 |
| 92 | 100.000 | 100.000 | 100.000 | 100.000 | 100.000 |
| 93 | 100.000 | 100.000 | 100.000 | 100.000 | 100.000 |
| 94 | 100.000 | 100.000 | 100.000 | 100.000 | 100.000 |
| 95 | 100.000 | 100.000 | 100.000 | 100.000 | 100.000 |
| 96 | 100.000 | 100.000 | 100.000 | 100.000 | 100.000 |
| 97 | 100.000 | 100.000 | 100.000 | 100.000 | 100.000 |
| 98 | 100.000 | 100.000 | 100.000 | 100.000 | 100.000 |
| 99 | 100.000 | 100.000 | 100.000 | 100.000 | 100.000 |
| 100 | 100.000 | 100.000 | 100.000 | 100.000 | 100.000 |

***Table S21:*** *Beat Alignment Test – Hit rate for slow tempo trials – BAT_slow_hits*

| | | | | | |
|---|---|---|---|---|---|
| **Task** | Beat Alignment Test | | | | |
| **Outcome measure** | Hit rate for slow tempo trials | | | | |
| **Variable name** | BAT_slow_hits | | | | |
| **Age and gender effects** | | | | | |
| Age regression (slope) | 0.000 | | | | |
| Age regression (p) | 1.000 | | | | |
| Age group (p) | 1.000 | | | | |
| Gender (p) | 1.000 | | | | |
| **Group** | All | Age 18 to 21 | Age 22 to 29 | Age 30 to 54 | Age 55 to 87 |
| N (N female) | 108 (74) | 27 (18) | 29 (19) | 26 (16) | 26 (21) |
| **Normality** | | | | | |
| Skewness | -1.24 | -1.12 | -1.32 | -0.92 | -0.97 |
| Excess kurtosis | 0.59 | -0.04 | 0.56 | -0.67 | -0.22 |
| **Scores** | | | | | |
| Mean | 85.880 | 88.657 | 90.086 | 84.615 | 79.567 |
| SD | 18.033 | 15.115 | 13.927 | 19.140 | 22.330 |
| Percentiles | | | | | |
| 0 | 31.250 | 50.000 | 56.250 | 43.750 | 31.250 |
| 1 | 31.688 | 53.250 | 56.250 | 45.313 | 31.250 |
| 2 | 38.375 | 56.500 | 56.250 | 46.875 | 31.250 |
| 3 | 45.063 | 59.750 | 56.250 | 48.438 | 31.250 |
| 4 | 50.000 | 62.500 | 57.000 | 50.000 | 31.250 |
| 5 | 50.000 | 62.500 | 58.750 | 50.000 | 32.813 |
| 6 | 50.000 | 62.500 | 60.500 | 50.000 | 34.375 |
| 7 | 53.063 | 62.500 | 62.250 | 50.000 | 35.938 |
| 8 | 56.250 | 63.000 | 64.000 | 50.000 | 37.500 |
| 9 | 56.250 | 64.625 | 65.750 | 51.563 | 40.625 |
| 10 | 60.625 | 66.250 | 67.500 | 53.125 | 43.750 |
| 11 | 62.500 | 67.875 | 69.250 | 54.688 | 46.875 |
| 12 | 62.500 | 68.750 | 71.000 | 56.250 | 50.000 |
| 13 | 62.500 | 68.750 | 72.750 | 57.813 | 53.125 |
| 14 | 62.500 | 68.750 | 74.500 | 59.375 | 56.250 |
| 15 | 62.500 | 68.750 | 76.250 | 60.938 | 59.375 |
| 16 | 62.500 | 68.750 | 78.000 | 62.500 | 62.500 |
| 17 | 63.688 | 68.750 | 79.750 | 62.500 | 62.500 |
| 18 | 68.750 | 68.750 | 81.250 | 62.500 | 62.500 |
| 19 | 68.750 | 68.750 | 81.250 | 62.500 | 62.500 |
| 20 | 68.750 | 71.250 | 81.250 | 62.500 | 62.500 |
| 21 | 68.750 | 74.500 | 81.250 | 64.063 | 62.500 |
| 22 | 68.750 | 77.750 | 82.250 | 65.625 | 62.500 |
| 23 | 72.563 | 81.000 | 84.000 | 67.188 | 62.500 |
| 24 | 75.000 | 81.250 | 85.750 | 68.750 | 62.500 |

| | | | | | |
|---|---|---|---|---|---|
| 25 | 75.000 | 81.250 | 87.500 | 70.313 | 64.063 |
| 26 | 80.125 | 81.250 | 87.500 | 71.875 | 65.625 |
| 27 | 81.250 | 81.375 | 87.500 | 73.438 | 67.188 |
| 28 | 81.250 | 83.000 | 87.500 | 75.000 | 68.750 |
| 29 | 81.250 | 84.625 | 87.500 | 76.563 | 70.313 |
| 30 | 81.250 | 86.250 | 87.500 | 78.125 | 71.875 |
| 31 | 81.250 | 87.500 | 87.500 | 79.688 | 73.438 |
| 32 | 81.250 | 87.500 | 87.500 | 81.250 | 75.000 |
| 33 | 83.188 | 87.500 | 87.500 | 82.813 | 76.563 |
| 34 | 87.500 | 87.500 | 87.500 | 84.375 | 78.125 |
| 35 | 87.500 | 88.125 | 87.500 | 85.938 | 79.688 |
| 36 | 87.500 | 89.750 | 88.000 | 87.500 | 81.250 |
| 37 | 87.500 | 91.375 | 89.750 | 87.500 | 81.250 |
| 38 | 87.500 | 93.000 | 91.500 | 87.500 | 81.250 |
| 39 | 87.500 | 93.750 | 93.250 | 87.500 | 81.250 |
| 40 | 87.500 | 93.750 | 93.750 | 87.500 | 81.250 |
| 41 | 87.500 | 93.750 | 93.750 | 89.063 | 81.250 |
| 42 | 87.500 | 93.750 | 93.750 | 90.625 | 81.250 |
| 43 | 87.563 | 93.750 | 93.750 | 92.188 | 81.250 |
| 44 | 93.750 | 93.750 | 93.750 | 93.750 | 81.250 |
| 45 | 93.750 | 93.750 | 93.750 | 93.750 | 82.813 |
| 46 | 93.750 | 93.750 | 93.750 | 93.750 | 84.375 |
| 47 | 93.750 | 93.750 | 94.750 | 93.750 | 85.938 |
| 48 | 93.750 | 93.750 | 96.500 | 93.750 | 87.500 |
| 49 | 93.750 | 93.750 | 98.250 | 93.750 | 87.500 |
| 50 | 93.750 | 93.750 | 100.000 | 93.750 | 87.500 |
| 51 | 93.750 | 95.375 | 100.000 | 93.750 | 87.500 |
| 52 | 93.750 | 97.000 | 100.000 | 93.750 | 87.500 |
| 53 | 93.750 | 98.625 | 100.000 | 93.750 | 87.500 |
| 54 | 93.750 | 100.000 | 100.000 | 93.750 | 87.500 |
| 55 | 93.750 | 100.000 | 100.000 | 93.750 | 87.500 |
| 56 | 93.750 | 100.000 | 100.000 | 93.750 | 87.500 |
| 57 | 99.938 | 100.000 | 100.000 | 95.313 | 89.063 |
| 58 | 100.000 | 100.000 | 100.000 | 96.875 | 90.625 |
| 59 | 100.000 | 100.000 | 100.000 | 98.438 | 92.188 |
| 60 | 100.000 | 100.000 | 100.000 | 100.000 | 93.750 |
| 61 | 100.000 | 100.000 | 100.000 | 100.000 | 93.750 |
| 62 | 100.000 | 100.000 | 100.000 | 100.000 | 93.750 |
| 63 | 100.000 | 100.000 | 100.000 | 100.000 | 93.750 |
| 64 | 100.000 | 100.000 | 100.000 | 100.000 | 93.750 |
| 65 | 100.000 | 100.000 | 100.000 | 100.000 | 93.750 |
| 66 | 100.000 | 100.000 | 100.000 | 100.000 | 93.750 |
| 67 | 100.000 | 100.000 | 100.000 | 100.000 | 93.750 |
| 68 | 100.000 | 100.000 | 100.000 | 100.000 | 93.750 |

| | | | | | |
|---|---|---|---|---|---|
| 69 | 100.000 | 100.000 | 100.000 | 100.000 | 95.313 |
| 70 | 100.000 | 100.000 | 100.000 | 100.000 | 96.875 |
| 71 | 100.000 | 100.000 | 100.000 | 100.000 | 98.438 |
| 72 | 100.000 | 100.000 | 100.000 | 100.000 | 100.000 |
| 73 | 100.000 | 100.000 | 100.000 | 100.000 | 100.000 |
| 74 | 100.000 | 100.000 | 100.000 | 100.000 | 100.000 |
| 75 | 100.000 | 100.000 | 100.000 | 100.000 | 100.000 |
| 76 | 100.000 | 100.000 | 100.000 | 100.000 | 100.000 |
| 77 | 100.000 | 100.000 | 100.000 | 100.000 | 100.000 |
| 78 | 100.000 | 100.000 | 100.000 | 100.000 | 100.000 |
| 79 | 100.000 | 100.000 | 100.000 | 100.000 | 100.000 |
| 80 | 100.000 | 100.000 | 100.000 | 100.000 | 100.000 |
| 81 | 100.000 | 100.000 | 100.000 | 100.000 | 100.000 |
| 82 | 100.000 | 100.000 | 100.000 | 100.000 | 100.000 |
| 83 | 100.000 | 100.000 | 100.000 | 100.000 | 100.000 |
| 84 | 100.000 | 100.000 | 100.000 | 100.000 | 100.000 |
| 85 | 100.000 | 100.000 | 100.000 | 100.000 | 100.000 |
| 86 | 100.000 | 100.000 | 100.000 | 100.000 | 100.000 |
| 87 | 100.000 | 100.000 | 100.000 | 100.000 | 100.000 |
| 88 | 100.000 | 100.000 | 100.000 | 100.000 | 100.000 |
| 89 | 100.000 | 100.000 | 100.000 | 100.000 | 100.000 |
| 90 | 100.000 | 100.000 | 100.000 | 100.000 | 100.000 |
| 91 | 100.000 | 100.000 | 100.000 | 100.000 | 100.000 |
| 92 | 100.000 | 100.000 | 100.000 | 100.000 | 100.000 |
| 93 | 100.000 | 100.000 | 100.000 | 100.000 | 100.000 |
| 94 | 100.000 | 100.000 | 100.000 | 100.000 | 100.000 |
| 95 | 100.000 | 100.000 | 100.000 | 100.000 | 100.000 |
| 96 | 100.000 | 100.000 | 100.000 | 100.000 | 100.000 |
| 97 | 100.000 | 100.000 | 100.000 | 100.000 | 100.000 |
| 98 | 100.000 | 100.000 | 100.000 | 100.000 | 100.000 |
| 99 | 100.000 | 100.000 | 100.000 | 100.000 | 100.000 |
| 100 | 100.000 | 100.000 | 100.000 | 100.000 | 100.000 |

***Table S22:*** *Beat Alignment Test – False alarm rate for all trials trials – BAT_all_fa*

| | | | | | |
|---|---|---|---|---|---|
| **Task** | Beat Alignment Test | | | | |
| **Outcome measure** | False alarm rate for all trials trials | | | | |
| **Variable name** | BAT_all_fa | | | | |
| **Age and gender effects** | | | | | |
| Age regression (slope) | 0.000 | | | | |
| Age regression (p) | 1.000 | | | | |
| Age group (p) | 1.000 | | | | |
| Gender (p) | 1.000 | | | | |
| **Group** | All | Age 18 to 21 | Age 22 to 29 | Age 30 to 54 | Age 55 to 87 |
| N (N female) | 108 (74) | 27 (18) | 29 (19) | 26 (16) | 26 (21) |
| **Normality** | | | | | |
| Skewness | 1.71 | 2.11 | 1.66 | 1.53 | 1.24 |
| Excess kurtosis | 2.49 | 3.69 | 2.18 | 1.84 | 0.17 |
| **Scores** | | | | | |
| Mean | 7.870 | 5.556 | 9.483 | 8.974 | 7.372 |
| SD | 10.977 | 9.245 | 12.594 | 12.001 | 9.740 |
| Percentiles | | | | | |
| 0 | 0.000 | 0.000 | 0.000 | 0.000 | 0.000 |
| 1 | 0.000 | 0.000 | 0.000 | 0.000 | 0.000 |
| 2 | 0.000 | 0.000 | 0.000 | 0.000 | 0.000 |
| 3 | 0.000 | 0.000 | 0.000 | 0.000 | 0.000 |
| 4 | 0.000 | 0.000 | 0.000 | 0.000 | 0.000 |
| 5 | 0.000 | 0.000 | 0.000 | 0.000 | 0.000 |
| 6 | 0.000 | 0.000 | 0.000 | 0.000 | 0.000 |
| 7 | 0.000 | 0.000 | 0.000 | 0.000 | 0.000 |
| 8 | 0.000 | 0.000 | 0.000 | 0.000 | 0.000 |
| 9 | 0.000 | 0.000 | 0.000 | 0.000 | 0.000 |
| 10 | 0.000 | 0.000 | 0.000 | 0.000 | 0.000 |
| 11 | 0.000 | 0.000 | 0.000 | 0.000 | 0.000 |
| 12 | 0.000 | 0.000 | 0.000 | 0.000 | 0.000 |
| 13 | 0.000 | 0.000 | 0.000 | 0.000 | 0.000 |
| 14 | 0.000 | 0.000 | 0.000 | 0.000 | 0.000 |
| 15 | 0.000 | 0.000 | 0.000 | 0.000 | 0.000 |
| 16 | 0.000 | 0.000 | 0.000 | 0.000 | 0.000 |
| 17 | 0.000 | 0.000 | 0.000 | 0.000 | 0.000 |
| 18 | 0.000 | 0.000 | 0.000 | 0.000 | 0.000 |
| 19 | 0.000 | 0.000 | 0.000 | 0.000 | 0.000 |
| 20 | 0.000 | 0.000 | 0.000 | 0.000 | 0.000 |
| 21 | 0.000 | 0.000 | 0.000 | 0.000 | 0.000 |
| 22 | 0.000 | 0.000 | 0.000 | 0.000 | 0.000 |
| 23 | 0.000 | 0.000 | 0.000 | 0.000 | 0.000 |
| 24 | 0.000 | 0.000 | 0.000 | 0.000 | 0.000 |

| | | | | | |
|---|---|---|---|---|---|
| 25 | 0.000 | 0.000 | 0.000 | 0.000 | 0.000 |
| 26 | 0.000 | 0.000 | 0.000 | 0.000 | 0.000 |
| 27 | 0.000 | 0.000 | 0.000 | 0.000 | 0.000 |
| 28 | 0.000 | 0.000 | 0.000 | 0.000 | 0.000 |
| 29 | 0.000 | 0.000 | 0.000 | 0.000 | 0.000 |
| 30 | 0.000 | 0.000 | 0.000 | 0.000 | 0.000 |
| 31 | 0.000 | 0.000 | 0.000 | 0.000 | 0.000 |
| 32 | 0.000 | 0.000 | 0.000 | 0.000 | 0.000 |
| 33 | 0.000 | 0.000 | 1.000 | 0.000 | 0.000 |
| 34 | 0.000 | 0.000 | 2.167 | 0.000 | 0.000 |
| 35 | 0.000 | 0.000 | 3.333 | 0.000 | 0.000 |
| 36 | 0.000 | 0.000 | 4.167 | 0.000 | 0.000 |
| 37 | 0.000 | 0.000 | 4.167 | 0.000 | 0.000 |
| 38 | 0.000 | 0.000 | 4.167 | 0.000 | 0.000 |
| 39 | 0.000 | 0.000 | 4.167 | 0.000 | 0.000 |
| 40 | 0.000 | 0.000 | 4.167 | 0.000 | 0.000 |
| 41 | 0.000 | 0.000 | 4.167 | 0.000 | 1.042 |
| 42 | 0.000 | 0.000 | 4.167 | 0.000 | 2.083 |
| 43 | 0.042 | 0.000 | 4.167 | 0.000 | 3.125 |
| 44 | 4.167 | 0.000 | 4.167 | 0.000 | 4.167 |
| 45 | 4.167 | 0.000 | 4.167 | 1.042 | 4.167 |
| 46 | 4.167 | 0.000 | 4.167 | 2.083 | 4.167 |
| 47 | 4.167 | 0.000 | 4.167 | 3.125 | 4.167 |
| 48 | 4.167 | 0.000 | 4.167 | 4.167 | 4.167 |
| 49 | 4.167 | 0.000 | 4.167 | 5.208 | 4.167 |
| 50 | 4.167 | 0.000 | 4.167 | 6.250 | 4.167 |
| 51 | 4.167 | 1.083 | 4.167 | 7.292 | 4.167 |
| 52 | 4.167 | 2.167 | 4.167 | 8.333 | 4.167 |
| 53 | 4.167 | 3.250 | 4.167 | 8.333 | 4.167 |
| 54 | 4.167 | 4.167 | 4.167 | 8.333 | 4.167 |
| 55 | 4.167 | 4.167 | 4.167 | 8.333 | 4.167 |
| 56 | 4.167 | 4.167 | 4.167 | 8.333 | 4.167 |
| 57 | 4.167 | 4.167 | 4.167 | 8.333 | 4.167 |
| 58 | 4.167 | 4.167 | 5.167 | 8.333 | 4.167 |
| 59 | 4.167 | 4.167 | 6.333 | 8.333 | 4.167 |
| 60 | 4.167 | 4.167 | 7.500 | 8.333 | 4.167 |
| 61 | 4.167 | 4.167 | 8.333 | 8.333 | 4.167 |
| 62 | 5.583 | 4.167 | 8.333 | 8.333 | 4.167 |
| 63 | 8.333 | 4.167 | 8.333 | 8.333 | 4.167 |
| 64 | 8.333 | 4.167 | 8.333 | 8.333 | 4.167 |
| 65 | 8.333 | 4.167 | 8.333 | 8.333 | 5.208 |
| 66 | 8.333 | 4.167 | 8.333 | 8.333 | 6.250 |
| 67 | 8.333 | 4.167 | 8.333 | 8.333 | 7.292 |
| 68 | 8.333 | 4.167 | 8.333 | 8.333 | 8.333 |

| | | | | | |
|---|---|---|---|---|---|
| 69 | 8.333 | 4.167 | 8.333 | 9.375 | 8.333 |
| 70 | 8.333 | 4.167 | 8.333 | 10.417 | 8.333 |
| 71 | 8.333 | 4.167 | 8.333 | 11.458 | 8.333 |
| 72 | 8.333 | 4.167 | 9.000 | 12.500 | 8.333 |
| 73 | 8.333 | 4.167 | 10.167 | 12.500 | 9.375 |
| 74 | 9.083 | 5.167 | 11.333 | 12.500 | 10.417 |
| 75 | 12.500 | 6.250 | 12.500 | 12.500 | 11.458 |
| 76 | 12.500 | 7.333 | 13.667 | 12.500 | 12.500 |
| 77 | 12.500 | 8.333 | 14.833 | 13.542 | 12.500 |
| 78 | 12.500 | 8.333 | 16.000 | 14.583 | 12.500 |
| 79 | 12.500 | 8.333 | 17.167 | 15.625 | 12.500 |
| 80 | 12.500 | 8.333 | 18.333 | 16.667 | 12.500 |
| 81 | 15.292 | 8.583 | 19.500 | 16.667 | 13.542 |
| 82 | 16.667 | 9.667 | 20.667 | 16.667 | 14.583 |
| 83 | 16.667 | 10.750 | 20.833 | 16.667 | 15.625 |
| 84 | 16.667 | 11.833 | 20.833 | 16.667 | 16.667 |
| 85 | 16.667 | 12.500 | 20.833 | 18.750 | 18.750 |
| 86 | 20.833 | 12.500 | 21.167 | 20.833 | 20.833 |
| 87 | 21.208 | 12.500 | 22.333 | 22.917 | 22.917 |
| 88 | 25.000 | 12.500 | 23.500 | 25.000 | 25.000 |
| 89 | 25.000 | 13.083 | 24.667 | 25.000 | 25.000 |
| 90 | 25.000 | 14.167 | 26.667 | 25.000 | 25.000 |
| 91 | 25.000 | 15.250 | 29.000 | 25.000 | 25.000 |
| 92 | 26.833 | 16.333 | 31.333 | 25.000 | 25.000 |
| 93 | 29.167 | 19.667 | 33.333 | 27.083 | 26.042 |
| 94 | 31.583 | 24.000 | 33.333 | 29.167 | 27.083 |
| 95 | 33.333 | 28.333 | 33.333 | 31.250 | 28.125 |
| 96 | 33.333 | 32.667 | 33.333 | 33.333 | 29.167 |
| 97 | 33.333 | 33.333 | 36.000 | 36.458 | 29.167 |
| 98 | 33.333 | 33.333 | 40.667 | 39.583 | 29.167 |
| 99 | 44.958 | 33.333 | 45.333 | 42.708 | 29.167 |
| 100 | 50.000 | 33.333 | 50.000 | 45.833 | 29.167 |

***Table S23:*** *Beat Alignment Test – False alarm rate for fast tempo trials – BAT_fast_fa*

| | | | | | |
|---|---|---|---|---|---|
| **Task** | Beat Alignment Test | | | | |
| **Outcome measure** | False alarm rate for fast tempo trials | | | | |
| **Variable name** | BAT_fast_fa | | | | |
| **Age and gender effects** | | | | | |
| Age regression (slope) | 0.000 | | | | |
| Age regression (p) | 1.000 | | | | |
| Age group (p) | 1.000 | | | | |
| Gender (p) | 1.000 | | | | |
| **Group** | All | Age 18 to 21 | Age 22 to 29 | Age 30 to 54 | Age 55 to 87 |
| N (N female) | 108 (74) | 27 (18) | 29 (19) | 26 (16) | 26 (21) |
| **Normality** | | | | | |
| Skewness | 1.31 | 1.92 | 0.95 | 0.83 | 1.41 |
| Excess kurtosis | 0.91 | 3.68 | -0.34 | -0.75 | 1.03 |
| **Scores** | | | | | |
| Mean | 10.880 | 7.407 | 12.500 | 11.538 | 12.019 |
| SD | 14.907 | 12.139 | 15.309 | 14.108 | 17.847 |
| Percentiles | | | | | |
| 0 | 0.000 | 0.000 | 0.000 | 0.000 | 0.000 |
| 1 | 0.000 | 0.000 | 0.000 | 0.000 | 0.000 |
| 2 | 0.000 | 0.000 | 0.000 | 0.000 | 0.000 |
| 3 | 0.000 | 0.000 | 0.000 | 0.000 | 0.000 |
| 4 | 0.000 | 0.000 | 0.000 | 0.000 | 0.000 |
| 5 | 0.000 | 0.000 | 0.000 | 0.000 | 0.000 |
| 6 | 0.000 | 0.000 | 0.000 | 0.000 | 0.000 |
| 7 | 0.000 | 0.000 | 0.000 | 0.000 | 0.000 |
| 8 | 0.000 | 0.000 | 0.000 | 0.000 | 0.000 |
| 9 | 0.000 | 0.000 | 0.000 | 0.000 | 0.000 |
| 10 | 0.000 | 0.000 | 0.000 | 0.000 | 0.000 |
| 11 | 0.000 | 0.000 | 0.000 | 0.000 | 0.000 |
| 12 | 0.000 | 0.000 | 0.000 | 0.000 | 0.000 |
| 13 | 0.000 | 0.000 | 0.000 | 0.000 | 0.000 |
| 14 | 0.000 | 0.000 | 0.000 | 0.000 | 0.000 |
| 15 | 0.000 | 0.000 | 0.000 | 0.000 | 0.000 |
| 16 | 0.000 | 0.000 | 0.000 | 0.000 | 0.000 |
| 17 | 0.000 | 0.000 | 0.000 | 0.000 | 0.000 |
| 18 | 0.000 | 0.000 | 0.000 | 0.000 | 0.000 |
| 19 | 0.000 | 0.000 | 0.000 | 0.000 | 0.000 |
| 20 | 0.000 | 0.000 | 0.000 | 0.000 | 0.000 |
| 21 | 0.000 | 0.000 | 0.000 | 0.000 | 0.000 |
| 22 | 0.000 | 0.000 | 0.000 | 0.000 | 0.000 |
| 23 | 0.000 | 0.000 | 0.000 | 0.000 | 0.000 |
| 24 | 0.000 | 0.000 | 0.000 | 0.000 | 0.000 |

| | | | | | |
|---|---|---|---|---|---|
| 25 | 0.000 | 0.000 | 0.000 | 0.000 | 0.000 |
| 26 | 0.000 | 0.000 | 0.000 | 0.000 | 0.000 |
| 27 | 0.000 | 0.000 | 0.000 | 0.000 | 0.000 |
| 28 | 0.000 | 0.000 | 0.000 | 0.000 | 0.000 |
| 29 | 0.000 | 0.000 | 0.000 | 0.000 | 0.000 |
| 30 | 0.000 | 0.000 | 0.000 | 0.000 | 0.000 |
| 31 | 0.000 | 0.000 | 0.000 | 0.000 | 0.000 |
| 32 | 0.000 | 0.000 | 0.000 | 0.000 | 0.000 |
| 33 | 0.000 | 0.000 | 0.000 | 0.000 | 0.000 |
| 34 | 0.000 | 0.000 | 0.000 | 0.000 | 0.000 |
| 35 | 0.000 | 0.000 | 0.000 | 0.000 | 0.000 |
| 36 | 0.000 | 0.000 | 0.000 | 0.000 | 0.000 |
| 37 | 0.000 | 0.000 | 0.000 | 0.000 | 0.000 |
| 38 | 0.000 | 0.000 | 0.000 | 0.000 | 0.000 |
| 39 | 0.000 | 0.000 | 0.000 | 0.000 | 0.000 |
| 40 | 0.000 | 0.000 | 0.000 | 0.000 | 0.000 |
| 41 | 0.000 | 0.000 | 0.000 | 0.000 | 0.000 |
| 42 | 0.000 | 0.000 | 0.000 | 0.000 | 0.000 |
| 43 | 0.000 | 0.000 | 0.000 | 0.000 | 0.000 |
| 44 | 0.000 | 0.000 | 0.000 | 0.000 | 0.000 |
| 45 | 0.000 | 0.000 | 0.000 | 0.000 | 0.000 |
| 46 | 0.000 | 0.000 | 0.000 | 0.000 | 0.000 |
| 47 | 0.000 | 0.000 | 2.000 | 0.000 | 0.000 |
| 48 | 0.000 | 0.000 | 5.500 | 0.000 | 0.000 |
| 49 | 0.000 | 0.000 | 9.000 | 3.125 | 0.000 |
| 50 | 0.000 | 0.000 | 12.500 | 6.250 | 0.000 |
| 51 | 0.000 | 0.000 | 12.500 | 9.375 | 0.000 |
| 52 | 0.000 | 0.000 | 12.500 | 12.500 | 0.000 |
| 53 | 0.000 | 0.000 | 12.500 | 12.500 | 0.000 |
| 54 | 0.000 | 0.000 | 12.500 | 12.500 | 0.000 |
| 55 | 10.625 | 0.000 | 12.500 | 12.500 | 0.000 |
| 56 | 12.500 | 0.000 | 12.500 | 12.500 | 0.000 |
| 57 | 12.500 | 0.000 | 12.500 | 12.500 | 3.125 |
| 58 | 12.500 | 0.000 | 12.500 | 12.500 | 6.250 |
| 59 | 12.500 | 0.000 | 12.500 | 12.500 | 9.375 |
| 60 | 12.500 | 0.000 | 12.500 | 12.500 | 12.500 |
| 61 | 12.500 | 0.000 | 12.500 | 12.500 | 12.500 |
| 62 | 12.500 | 1.500 | 12.500 | 12.500 | 12.500 |
| 63 | 12.500 | 4.750 | 12.500 | 12.500 | 12.500 |
| 64 | 12.500 | 8.000 | 12.500 | 12.500 | 12.500 |
| 65 | 12.500 | 11.250 | 12.500 | 12.500 | 12.500 |
| 66 | 12.500 | 12.500 | 12.500 | 12.500 | 12.500 |
| 67 | 12.500 | 12.500 | 12.500 | 12.500 | 12.500 |
| 68 | 12.500 | 12.500 | 12.500 | 12.500 | 12.500 |

| | | | | | |
|---|---|---|---|---|---|
| 69 | 12.500 | 12.500 | 12.500 | 12.500 | 12.500 |
| 70 | 12.500 | 12.500 | 12.500 | 12.500 | 12.500 |
| 71 | 12.500 | 12.500 | 12.500 | 12.500 | 12.500 |
| 72 | 12.500 | 12.500 | 14.500 | 12.500 | 12.500 |
| 73 | 12.500 | 12.500 | 18.000 | 15.625 | 15.625 |
| 74 | 12.500 | 12.500 | 21.500 | 18.750 | 18.750 |
| 75 | 12.500 | 12.500 | 25.000 | 21.875 | 21.875 |
| 76 | 16.500 | 12.500 | 25.000 | 25.000 | 25.000 |
| 77 | 25.000 | 12.500 | 25.000 | 25.000 | 25.000 |
| 78 | 25.000 | 12.500 | 25.000 | 25.000 | 25.000 |
| 79 | 25.000 | 12.500 | 25.000 | 25.000 | 25.000 |
| 80 | 25.000 | 12.500 | 25.000 | 25.000 | 25.000 |
| 81 | 25.000 | 12.500 | 25.000 | 25.000 | 25.000 |
| 82 | 25.000 | 12.500 | 25.000 | 25.000 | 25.000 |
| 83 | 25.000 | 12.500 | 28.000 | 25.000 | 25.000 |
| 84 | 25.000 | 12.500 | 31.500 | 25.000 | 25.000 |
| 85 | 25.000 | 13.750 | 35.000 | 28.125 | 28.125 |
| 86 | 25.000 | 17.000 | 37.500 | 31.250 | 31.250 |
| 87 | 26.125 | 20.250 | 37.500 | 34.375 | 34.375 |
| 88 | 37.500 | 23.500 | 37.500 | 37.500 | 37.500 |
| 89 | 37.500 | 25.000 | 37.500 | 37.500 | 37.500 |
| 90 | 37.500 | 25.000 | 37.500 | 37.500 | 37.500 |
| 91 | 37.500 | 25.000 | 37.500 | 37.500 | 37.500 |
| 92 | 37.500 | 25.000 | 37.500 | 37.500 | 37.500 |
| 93 | 37.500 | 25.000 | 37.500 | 37.500 | 40.625 |
| 94 | 37.500 | 25.000 | 37.500 | 37.500 | 43.750 |
| 95 | 37.500 | 25.000 | 37.500 | 37.500 | 46.875 |
| 96 | 37.500 | 25.000 | 37.500 | 37.500 | 50.000 |
| 97 | 47.375 | 30.500 | 39.500 | 37.500 | 53.125 |
| 98 | 50.000 | 37.000 | 43.000 | 37.500 | 56.250 |
| 99 | 50.000 | 43.500 | 46.500 | 37.500 | 59.375 |
| 100 | 62.500 | 50.000 | 50.000 | 37.500 | 62.500 |

***Table S24:*** *Beat Alignment Test – False alarm rate for medium tempo trials – BAT_med_fa*

| | | | | | |
|---|---|---|---|---|---|
| **Task** | Beat Alignment Test | | | | |
| **Outcome measure** | False alarm rate for medium tempo trials | | | | |
| **Variable name** | BAT_med_fa | | | | |
| **Age and gender effects** | | | | | |
| Age regression (slope) | 0.000 | | | | |
| Age regression (p) | 1.000 | | | | |
| Age group (p) | 1.000 | | | | |
| Gender (p) | 1.000 | | | | |
| **Group** | All | Age 18 to 21 | Age 22 to 29 | Age 30 to 54 | Age 55 to 87 |
| N (N female) | 108 (74) | 27 (18) | 29 (19) | 26 (16) | 26 (21) |
| **Normality** | | | | | |
| Skewness | 2.28 | 2.58 | 2.14 | 1.61 | 1.32 |
| Excess kurtosis | 5.27 | 6.04 | 4.34 | 1.68 | 0.77 |
| **Scores** | | | | | |
| Mean | 6.944 | 5.556 | 8.621 | 9.135 | 4.327 |
| SD | 12.481 | 12.175 | 14.979 | 13.945 | 7.020 |
| Percentiles | | | | | |
| 0 | 0.000 | 0.000 | 0.000 | 0.000 | 0.000 |
| 1 | 0.000 | 0.000 | 0.000 | 0.000 | 0.000 |
| 2 | 0.000 | 0.000 | 0.000 | 0.000 | 0.000 |
| 3 | 0.000 | 0.000 | 0.000 | 0.000 | 0.000 |
| 4 | 0.000 | 0.000 | 0.000 | 0.000 | 0.000 |
| 5 | 0.000 | 0.000 | 0.000 | 0.000 | 0.000 |
| 6 | 0.000 | 0.000 | 0.000 | 0.000 | 0.000 |
| 7 | 0.000 | 0.000 | 0.000 | 0.000 | 0.000 |
| 8 | 0.000 | 0.000 | 0.000 | 0.000 | 0.000 |
| 9 | 0.000 | 0.000 | 0.000 | 0.000 | 0.000 |
| 10 | 0.000 | 0.000 | 0.000 | 0.000 | 0.000 |
| 11 | 0.000 | 0.000 | 0.000 | 0.000 | 0.000 |
| 12 | 0.000 | 0.000 | 0.000 | 0.000 | 0.000 |
| 13 | 0.000 | 0.000 | 0.000 | 0.000 | 0.000 |
| 14 | 0.000 | 0.000 | 0.000 | 0.000 | 0.000 |
| 15 | 0.000 | 0.000 | 0.000 | 0.000 | 0.000 |
| 16 | 0.000 | 0.000 | 0.000 | 0.000 | 0.000 |
| 17 | 0.000 | 0.000 | 0.000 | 0.000 | 0.000 |
| 18 | 0.000 | 0.000 | 0.000 | 0.000 | 0.000 |
| 19 | 0.000 | 0.000 | 0.000 | 0.000 | 0.000 |
| 20 | 0.000 | 0.000 | 0.000 | 0.000 | 0.000 |
| 21 | 0.000 | 0.000 | 0.000 | 0.000 | 0.000 |
| 22 | 0.000 | 0.000 | 0.000 | 0.000 | 0.000 |
| 23 | 0.000 | 0.000 | 0.000 | 0.000 | 0.000 |
| 24 | 0.000 | 0.000 | 0.000 | 0.000 | 0.000 |

| | | | | | |
|---|---|---|---|---|---|
| 25 | 0.000 | 0.000 | 0.000 | 0.000 | 0.000 |
| 26 | 0.000 | 0.000 | 0.000 | 0.000 | 0.000 |
| 27 | 0.000 | 0.000 | 0.000 | 0.000 | 0.000 |
| 28 | 0.000 | 0.000 | 0.000 | 0.000 | 0.000 |
| 29 | 0.000 | 0.000 | 0.000 | 0.000 | 0.000 |
| 30 | 0.000 | 0.000 | 0.000 | 0.000 | 0.000 |
| 31 | 0.000 | 0.000 | 0.000 | 0.000 | 0.000 |
| 32 | 0.000 | 0.000 | 0.000 | 0.000 | 0.000 |
| 33 | 0.000 | 0.000 | 0.000 | 0.000 | 0.000 |
| 34 | 0.000 | 0.000 | 0.000 | 0.000 | 0.000 |
| 35 | 0.000 | 0.000 | 0.000 | 0.000 | 0.000 |
| 36 | 0.000 | 0.000 | 0.000 | 0.000 | 0.000 |
| 37 | 0.000 | 0.000 | 0.000 | 0.000 | 0.000 |
| 38 | 0.000 | 0.000 | 0.000 | 0.000 | 0.000 |
| 39 | 0.000 | 0.000 | 0.000 | 0.000 | 0.000 |
| 40 | 0.000 | 0.000 | 0.000 | 0.000 | 0.000 |
| 41 | 0.000 | 0.000 | 0.000 | 0.000 | 0.000 |
| 42 | 0.000 | 0.000 | 0.000 | 0.000 | 0.000 |
| 43 | 0.000 | 0.000 | 0.000 | 0.000 | 0.000 |
| 44 | 0.000 | 0.000 | 0.000 | 0.000 | 0.000 |
| 45 | 0.000 | 0.000 | 0.000 | 0.000 | 0.000 |
| 46 | 0.000 | 0.000 | 0.000 | 0.000 | 0.000 |
| 47 | 0.000 | 0.000 | 0.000 | 0.000 | 0.000 |
| 48 | 0.000 | 0.000 | 0.000 | 0.000 | 0.000 |
| 49 | 0.000 | 0.000 | 0.000 | 0.000 | 0.000 |
| 50 | 0.000 | 0.000 | 0.000 | 0.000 | 0.000 |
| 51 | 0.000 | 0.000 | 0.000 | 0.000 | 0.000 |
| 52 | 0.000 | 0.000 | 0.000 | 0.000 | 0.000 |
| 53 | 0.000 | 0.000 | 0.000 | 0.000 | 0.000 |
| 54 | 0.000 | 0.000 | 0.000 | 0.000 | 0.000 |
| 55 | 0.000 | 0.000 | 0.000 | 0.000 | 0.000 |
| 56 | 0.000 | 0.000 | 0.000 | 0.000 | 0.000 |
| 57 | 0.000 | 0.000 | 0.000 | 3.125 | 0.000 |
| 58 | 0.000 | 0.000 | 0.000 | 6.250 | 0.000 |
| 59 | 0.000 | 0.000 | 0.000 | 9.375 | 0.000 |
| 60 | 0.000 | 0.000 | 0.000 | 12.500 | 0.000 |
| 61 | 0.000 | 0.000 | 1.000 | 12.500 | 0.000 |
| 62 | 0.000 | 0.000 | 4.500 | 12.500 | 0.000 |
| 63 | 0.000 | 0.000 | 8.000 | 12.500 | 0.000 |
| 64 | 0.000 | 0.000 | 11.500 | 12.500 | 0.000 |
| 65 | 0.000 | 0.000 | 12.500 | 12.500 | 0.000 |
| 66 | 7.750 | 0.000 | 12.500 | 12.500 | 0.000 |
| 67 | 12.500 | 0.000 | 12.500 | 12.500 | 0.000 |
| 68 | 12.500 | 0.000 | 12.500 | 12.500 | 0.000 |

| | | | | | |
|---|---|---|---|---|---|
| 69 | 12.500 | 0.000 | 12.500 | 12.500 | 3.125 |
| 70 | 12.500 | 0.000 | 12.500 | 12.500 | 6.250 |
| 71 | 12.500 | 0.000 | 12.500 | 12.500 | 9.375 |
| 72 | 12.500 | 0.000 | 12.500 | 12.500 | 12.500 |
| 73 | 12.500 | 0.000 | 12.500 | 12.500 | 12.500 |
| 74 | 12.500 | 3.000 | 12.500 | 12.500 | 12.500 |
| 75 | 12.500 | 6.250 | 12.500 | 12.500 | 12.500 |
| 76 | 12.500 | 9.500 | 12.500 | 12.500 | 12.500 |
| 77 | 12.500 | 12.500 | 12.500 | 12.500 | 12.500 |
| 78 | 12.500 | 12.500 | 12.500 | 12.500 | 12.500 |
| 79 | 12.500 | 12.500 | 12.500 | 12.500 | 12.500 |
| 80 | 12.500 | 12.500 | 12.500 | 12.500 | 12.500 |
| 81 | 12.500 | 12.500 | 12.500 | 12.500 | 12.500 |
| 82 | 12.500 | 12.500 | 12.500 | 12.500 | 12.500 |
| 83 | 12.500 | 12.500 | 12.500 | 12.500 | 12.500 |
| 84 | 12.500 | 12.500 | 12.500 | 12.500 | 12.500 |
| 85 | 12.500 | 12.500 | 12.500 | 15.625 | 12.500 |
| 86 | 12.500 | 12.500 | 13.500 | 18.750 | 12.500 |
| 87 | 12.500 | 12.500 | 17.000 | 21.875 | 12.500 |
| 88 | 12.500 | 12.500 | 20.500 | 25.000 | 12.500 |
| 89 | 12.500 | 12.500 | 24.000 | 28.125 | 12.500 |
| 90 | 16.250 | 12.500 | 27.500 | 31.250 | 12.500 |
| 91 | 25.000 | 12.500 | 31.000 | 34.375 | 12.500 |
| 92 | 25.000 | 12.500 | 34.500 | 37.500 | 12.500 |
| 93 | 31.375 | 17.000 | 37.500 | 37.500 | 12.500 |
| 94 | 37.500 | 23.500 | 37.500 | 37.500 | 12.500 |
| 95 | 37.500 | 30.000 | 37.500 | 37.500 | 12.500 |
| 96 | 37.500 | 36.500 | 37.500 | 37.500 | 12.500 |
| 97 | 37.500 | 40.250 | 41.500 | 40.625 | 15.625 |
| 98 | 48.250 | 43.500 | 48.500 | 43.750 | 18.750 |
| 99 | 50.000 | 46.750 | 55.500 | 46.875 | 21.875 |
| 100 | 62.500 | 50.000 | 62.500 | 50.000 | 25.000 |

***Table S25:*** *Beat Alignment Test – False alarm rate for slow tempo trials – BAT_slow_fa*

| **Task** | Beat Alignment Test | | | | |
|---|---|---|---|---|---|
| **Outcome measure** | False alarm rate for slow tempo trials | | | | |
| **Variable name** | BAT_slow_fa | | | | |
| **Age and gender effects** | | | | | |
| Age regression (slope) | 0.000 | | | | |
| Age regression (p) | 1.000 | | | | |
| Age group (p) | 1.000 | | | | |
| Gender (p) | 1.000 | | | | |
| **Group** | All | Age 18 to 21 | Age 22 to 29 | Age 30 to 54 | Age 55 to 87 |
| N (N female) | 108 (74) | 27 (18) | 29 (19) | 26 (16) | 26 (21) |
| **Normality** | | | | | |
| Skewness | 2.23 | 2.75 | 1.93 | 2.11 | 2.05 |
| Excess kurtosis | 4.49 | 7.94 | 3.20 | 3.42 | 3.44 |
| **Scores** | | | | | |
| Mean | 5.787 | 3.704 | 7.328 | 6.250 | 5.769 |
| SD | 11.391 | 8.360 | 12.729 | 13.346 | 10.742 |
| Percentiles | | | | | |
| 0 | 0.000 | 0.000 | 0.000 | 0.000 | 0.000 |
| 1 | 0.000 | 0.000 | 0.000 | 0.000 | 0.000 |
| 2 | 0.000 | 0.000 | 0.000 | 0.000 | 0.000 |
| 3 | 0.000 | 0.000 | 0.000 | 0.000 | 0.000 |
| 4 | 0.000 | 0.000 | 0.000 | 0.000 | 0.000 |
| 5 | 0.000 | 0.000 | 0.000 | 0.000 | 0.000 |
| 6 | 0.000 | 0.000 | 0.000 | 0.000 | 0.000 |
| 7 | 0.000 | 0.000 | 0.000 | 0.000 | 0.000 |
| 8 | 0.000 | 0.000 | 0.000 | 0.000 | 0.000 |
| 9 | 0.000 | 0.000 | 0.000 | 0.000 | 0.000 |
| 10 | 0.000 | 0.000 | 0.000 | 0.000 | 0.000 |
| 11 | 0.000 | 0.000 | 0.000 | 0.000 | 0.000 |
| 12 | 0.000 | 0.000 | 0.000 | 0.000 | 0.000 |
| 13 | 0.000 | 0.000 | 0.000 | 0.000 | 0.000 |
| 14 | 0.000 | 0.000 | 0.000 | 0.000 | 0.000 |
| 15 | 0.000 | 0.000 | 0.000 | 0.000 | 0.000 |
| 16 | 0.000 | 0.000 | 0.000 | 0.000 | 0.000 |
| 17 | 0.000 | 0.000 | 0.000 | 0.000 | 0.000 |
| 18 | 0.000 | 0.000 | 0.000 | 0.000 | 0.000 |
| 19 | 0.000 | 0.000 | 0.000 | 0.000 | 0.000 |
| 20 | 0.000 | 0.000 | 0.000 | 0.000 | 0.000 |
| 21 | 0.000 | 0.000 | 0.000 | 0.000 | 0.000 |
| 22 | 0.000 | 0.000 | 0.000 | 0.000 | 0.000 |
| 23 | 0.000 | 0.000 | 0.000 | 0.000 | 0.000 |
| 24 | 0.000 | 0.000 | 0.000 | 0.000 | 0.000 |

| | | | | | |
|---|---|---|---|---|---|
| 25 | 0.000 | 0.000 | 0.000 | 0.000 | 0.000 |
| 26 | 0.000 | 0.000 | 0.000 | 0.000 | 0.000 |
| 27 | 0.000 | 0.000 | 0.000 | 0.000 | 0.000 |
| 28 | 0.000 | 0.000 | 0.000 | 0.000 | 0.000 |
| 29 | 0.000 | 0.000 | 0.000 | 0.000 | 0.000 |
| 30 | 0.000 | 0.000 | 0.000 | 0.000 | 0.000 |
| 31 | 0.000 | 0.000 | 0.000 | 0.000 | 0.000 |
| 32 | 0.000 | 0.000 | 0.000 | 0.000 | 0.000 |
| 33 | 0.000 | 0.000 | 0.000 | 0.000 | 0.000 |
| 34 | 0.000 | 0.000 | 0.000 | 0.000 | 0.000 |
| 35 | 0.000 | 0.000 | 0.000 | 0.000 | 0.000 |
| 36 | 0.000 | 0.000 | 0.000 | 0.000 | 0.000 |
| 37 | 0.000 | 0.000 | 0.000 | 0.000 | 0.000 |
| 38 | 0.000 | 0.000 | 0.000 | 0.000 | 0.000 |
| 39 | 0.000 | 0.000 | 0.000 | 0.000 | 0.000 |
| 40 | 0.000 | 0.000 | 0.000 | 0.000 | 0.000 |
| 41 | 0.000 | 0.000 | 0.000 | 0.000 | 0.000 |
| 42 | 0.000 | 0.000 | 0.000 | 0.000 | 0.000 |
| 43 | 0.000 | 0.000 | 0.000 | 0.000 | 0.000 |
| 44 | 0.000 | 0.000 | 0.000 | 0.000 | 0.000 |
| 45 | 0.000 | 0.000 | 0.000 | 0.000 | 0.000 |
| 46 | 0.000 | 0.000 | 0.000 | 0.000 | 0.000 |
| 47 | 0.000 | 0.000 | 0.000 | 0.000 | 0.000 |
| 48 | 0.000 | 0.000 | 0.000 | 0.000 | 0.000 |
| 49 | 0.000 | 0.000 | 0.000 | 0.000 | 0.000 |
| 50 | 0.000 | 0.000 | 0.000 | 0.000 | 0.000 |
| 51 | 0.000 | 0.000 | 0.000 | 0.000 | 0.000 |
| 52 | 0.000 | 0.000 | 0.000 | 0.000 | 0.000 |
| 53 | 0.000 | 0.000 | 0.000 | 0.000 | 0.000 |
| 54 | 0.000 | 0.000 | 0.000 | 0.000 | 0.000 |
| 55 | 0.000 | 0.000 | 0.000 | 0.000 | 0.000 |
| 56 | 0.000 | 0.000 | 0.000 | 0.000 | 0.000 |
| 57 | 0.000 | 0.000 | 0.000 | 0.000 | 0.000 |
| 58 | 0.000 | 0.000 | 0.000 | 0.000 | 0.000 |
| 59 | 0.000 | 0.000 | 0.000 | 0.000 | 0.000 |
| 60 | 0.000 | 0.000 | 0.000 | 0.000 | 0.000 |
| 61 | 0.000 | 0.000 | 0.000 | 0.000 | 0.000 |
| 62 | 0.000 | 0.000 | 0.000 | 0.000 | 0.000 |
| 63 | 0.000 | 0.000 | 0.000 | 0.000 | 0.000 |
| 64 | 0.000 | 0.000 | 0.000 | 0.000 | 0.000 |
| 65 | 0.000 | 0.000 | 2.500 | 0.000 | 0.000 |
| 66 | 0.000 | 0.000 | 6.000 | 0.000 | 0.000 |
| 67 | 0.000 | 0.000 | 9.500 | 0.000 | 0.000 |
| 68 | 0.000 | 0.000 | 12.500 | 0.000 | 0.000 |

| | | | | | |
|---|---|---|---|---|---|
| 69 | 0.000 | 0.000 | 12.500 | 0.000 | 3.125 |
| 70 | 0.000 | 0.000 | 12.500 | 0.000 | 6.250 |
| 71 | 0.000 | 0.000 | 12.500 | 0.000 | 9.375 |
| 72 | 0.500 | 0.000 | 12.500 | 0.000 | 12.500 |
| 73 | 12.500 | 0.000 | 12.500 | 0.000 | 12.500 |
| 74 | 12.500 | 0.000 | 12.500 | 0.000 | 12.500 |
| 75 | 12.500 | 0.000 | 12.500 | 0.000 | 12.500 |
| 76 | 12.500 | 0.000 | 12.500 | 0.000 | 12.500 |
| 77 | 12.500 | 0.250 | 12.500 | 3.125 | 12.500 |
| 78 | 12.500 | 3.500 | 12.500 | 6.250 | 12.500 |
| 79 | 12.500 | 6.750 | 12.500 | 9.375 | 12.500 |
| 80 | 12.500 | 10.000 | 12.500 | 12.500 | 12.500 |
| 81 | 12.500 | 12.500 | 12.500 | 12.500 | 12.500 |
| 82 | 12.500 | 12.500 | 12.500 | 12.500 | 12.500 |
| 83 | 12.500 | 12.500 | 12.500 | 12.500 | 12.500 |
| 84 | 12.500 | 12.500 | 12.500 | 12.500 | 12.500 |
| 85 | 12.500 | 12.500 | 12.500 | 15.625 | 12.500 |
| 86 | 12.500 | 12.500 | 13.500 | 18.750 | 12.500 |
| 87 | 12.500 | 12.500 | 17.000 | 21.875 | 12.500 |
| 88 | 12.500 | 12.500 | 20.500 | 25.000 | 12.500 |
| 89 | 12.500 | 12.500 | 24.000 | 25.000 | 12.500 |
| 90 | 16.250 | 12.500 | 25.000 | 25.000 | 12.500 |
| 91 | 25.000 | 12.500 | 25.000 | 25.000 | 12.500 |
| 92 | 25.000 | 12.500 | 25.000 | 25.000 | 12.500 |
| 93 | 25.000 | 12.500 | 25.500 | 28.125 | 18.750 |
| 94 | 32.250 | 12.500 | 29.000 | 31.250 | 25.000 |
| 95 | 37.500 | 12.500 | 32.500 | 34.375 | 31.250 |
| 96 | 37.500 | 12.500 | 36.000 | 37.500 | 37.500 |
| 97 | 37.500 | 18.000 | 39.500 | 40.625 | 37.500 |
| 98 | 37.500 | 24.500 | 43.000 | 43.750 | 37.500 |
| 99 | 49.125 | 31.000 | 46.500 | 46.875 | 37.500 |
| 100 | 50.000 | 37.500 | 50.000 | 50.000 | 37.500 |

***Table S26:*** *Unpaced tapping – Rate (ITI in ms) of spontaneous tapping, initial trial – Unpaced_spontaneous_right_mean_iti*

| | | | | | |
|---|---|---|---|---|---|
| **Task** | Unpaced tapping | | | | |
| **Outcome measure** | Rate (ITI in ms) of spontaneous tapping, initial trial | | | | |
| **Variable name** | Unpaced_spontaneous_right_mean_iti | | | | |
| **Age and gender effects** | | | | | |
| Age regression (slope) | -0.517 | | | | |
| Age regression (p) | 0.578 | | | | |
| Age group (p) | 0.652 | | | | |
| Gender (p) | 0.102 | | | | |
| **Group** | All | Age 18 to 21 | Age 22 to 29 | Age 30 to 54 | Age 55 to 87 |
| N (N female) | 106 (72) | 26 (17) | 29 (19) | 26 (16) | 25 (20) |
| **Normality** | | | | | |
| Skewness | 3.22 | 0.51 | 0.78 | 2.76 | 0.38 |
| Excess kurtosis | 18.96 | -0.11 | 1.13 | 9.19 | -1.24 |
| **Scores** | | | | | |
| Mean | 685.3 | 721.1 | 661.1 | 718.9 | 641.0 |
| SD | 243.0 | 225.8 | 152.7 | 394.5 | 102.2 |
| Percentiles | | | | | |
| 0 | 233.9 | 336.3 | 386.4 | 233.9 | 488.8 |
| 1 | 338.8 | 367.4 | 401.2 | 273.7 | 496.3 |
| 2 | 387.0 | 398.4 | 416.1 | 313.5 | 503.8 |
| 3 | 394.1 | 429.5 | 431.0 | 353.3 | 511.3 |
| 4 | 404.3 | 460.5 | 446.0 | 393.0 | 518.8 |
| 5 | 426.6 | 463.3 | 461.2 | 394.7 | 520.9 |
| 6 | 445.8 | 466.0 | 476.3 | 396.4 | 521.9 |
| 7 | 462.8 | 468.7 | 491.5 | 398.1 | 522.9 |
| 8 | 467.6 | 471.4 | 494.2 | 399.8 | 524.0 |
| 9 | 469.8 | 474.0 | 494.7 | 405.5 | 527.2 |
| 10 | 473.4 | 476.6 | 495.3 | 411.1 | 531.4 |
| 11 | 478.9 | 479.2 | 497.0 | 416.7 | 535.6 |
| 12 | 482.9 | 481.8 | 501.7 | 422.3 | 539.8 |
| 13 | 484.9 | 482.2 | 506.4 | 433.5 | 543.5 |
| 14 | 487.9 | 482.7 | 511.1 | 444.7 | 546.7 |
| 15 | 492.5 | 483.2 | 515.2 | 455.9 | 549.8 |
| 16 | 495.3 | 483.6 | 519.1 | 467.1 | 552.9 |
| 17 | 509.9 | 495.4 | 523.0 | 467.4 | 555.4 |
| 18 | 519.2 | 507.2 | 526.3 | 467.8 | 556.4 |
| 19 | 524.1 | 519.0 | 526.7 | 468.1 | 557.5 |
| 20 | 526.3 | 530.8 | 527.2 | 468.5 | 558.6 |
| 21 | 527.9 | 530.9 | 527.6 | 470.2 | 559.5 |
| 22 | 530.8 | 531.0 | 531.1 | 471.9 | 559.7 |

| | | | | | |
|---|---|---|---|---|---|
| 23 | 532.8 | 531.1 | 537.1 | 473.6 | 559.9 |
| 24 | 543.3 | 531.2 | 543.0 | 475.4 | 560.1 |
| 25 | 549.4 | 536.1 | 549.0 | 477.9 | 560.2 |
| 26 | 551.9 | 540.9 | 558.8 | 480.5 | 561.2 |
| 27 | 555.4 | 545.7 | 568.6 | 483.1 | 562.2 |
| 28 | 557.4 | 550.6 | 578.4 | 485.6 | 563.2 |
| 29 | 559.8 | 551.9 | 585.7 | 504.3 | 564.2 |
| 30 | 560.4 | 553.3 | 589.6 | 523.1 | 564.5 |
| 31 | 562.6 | 554.6 | 593.6 | 541.8 | 564.8 |
| 32 | 565.0 | 556.0 | 597.5 | 560.5 | 565.1 |
| 33 | 566.1 | 558.6 | 598.5 | 566.2 | 565.3 |
| 34 | 571.4 | 561.2 | 598.9 | 571.9 | 566.7 |
| 35 | 577.7 | 563.9 | 599.4 | 577.6 | 568.6 |
| 36 | 582.5 | 566.5 | 602.0 | 583.4 | 570.6 |
| 37 | 583.9 | 593.2 | 609.9 | 588.0 | 572.5 |
| 38 | 596.6 | 620.0 | 617.8 | 592.7 | 574.1 |
| 39 | 599.7 | 646.8 | 625.6 | 597.3 | 575.5 |
| 40 | 602.0 | 673.6 | 628.3 | 602.0 | 576.9 |
| 41 | 603.5 | 675.0 | 628.9 | 606.7 | 578.2 |
| 42 | 608.3 | 676.4 | 629.4 | 611.4 | 581.1 |
| 43 | 613.4 | 677.8 | 630.2 | 616.1 | 586.8 |
| 44 | 622.0 | 679.3 | 632.5 | 620.8 | 592.6 |
| 45 | 627.0 | 679.4 | 634.7 | 622.3 | 598.4 |
| 46 | 628.5 | 679.5 | 636.9 | 623.8 | 603.4 |
| 47 | 630.5 | 679.6 | 639.0 | 625.2 | 604.5 |
| 48 | 634.0 | 679.8 | 641.0 | 626.7 | 605.6 |
| 49 | 639.9 | 682.2 | 643.1 | 627.9 | 606.8 |
| 50 | 643.8 | 684.6 | 645.1 | 629.1 | 607.9 |
| 51 | 649.7 | 687.0 | 653.0 | 630.3 | 608.9 |
| 52 | 665.2 | 689.4 | 660.8 | 631.4 | 609.9 |
| 53 | 673.4 | 708.2 | 668.6 | 648.3 | 611.0 |
| 54 | 676.4 | 727.0 | 673.6 | 665.2 | 612.0 |
| 55 | 678.8 | 745.7 | 674.9 | 682.0 | 618.2 |
| 56 | 679.7 | 764.5 | 676.1 | 698.9 | 625.5 |
| 57 | 681.6 | 768.9 | 677.4 | 709.6 | 632.7 |
| 58 | 687.9 | 773.2 | 678.6 | 720.3 | 640.0 |
| 59 | 689.4 | 777.6 | 679.9 | 731.0 | 644.2 |
| 60 | 698.9 | 782.0 | 681.1 | 741.7 | 646.8 |
| 61 | 715.0 | 783.6 | 682.5 | 742.8 | 649.4 |
| 62 | 727.7 | 785.2 | 684.3 | 743.9 | 652.1 |
| 63 | 731.8 | 786.8 | 686.2 | 745.0 | 662.2 |
| 64 | 737.5 | 788.4 | 688.1 | 746.2 | 680.0 |
| 65 | 742.0 | 800.1 | 693.7 | 747.4 | 697.7 |
| 66 | 743.7 | 811.8 | 700.9 | 748.6 | 715.5 |

| | | | | | |
|---|---|---|---|---|---|
| 67 | 745.5 | 823.4 | 708.1 | 749.9 | 727.6 |
| 68 | 747.6 | 835.1 | 715.5 | 751.1 | 728.5 |
| 69 | 750.4 | 843.3 | 723.5 | 761.3 | 729.4 |
| 70 | 756.0 | 851.5 | 731.6 | 771.4 | 730.3 |
| 71 | 762.8 | 859.6 | 739.6 | 781.6 | 731.2 |
| 72 | 764.5 | 867.8 | 743.4 | 791.8 | 732.5 |
| 73 | 766.7 | 870.6 | 744.0 | 808.7 | 733.8 |
| 74 | 774.3 | 873.5 | 744.5 | 825.6 | 735.1 |
| 75 | 777.8 | 876.4 | 745.1 | 842.5 | 736.5 |
| 76 | 781.2 | 879.2 | 751.5 | 859.4 | 739.7 |
| 77 | 786.0 | 880.8 | 757.9 | 860.1 | 742.9 |
| 78 | 788.2 | 882.4 | 764.3 | 860.8 | 746.1 |
| 79 | 791.4 | 884.0 | 769.0 | 861.5 | 749.3 |
| 80 | 791.8 | 885.5 | 771.5 | 862.1 | 752.0 |
| 81 | 792.7 | 902.3 | 774.1 | 864.5 | 754.7 |
| 82 | 802.3 | 919.1 | 776.6 | 866.9 | 757.4 |
| 83 | 835.8 | 935.8 | 780.5 | 869.3 | 760.0 |
| 84 | 843.5 | 952.6 | 784.6 | 871.7 | 761.5 |
| 85 | 860.1 | 963.0 | 788.7 | 877.9 | 762.3 |
| 86 | 863.8 | 973.4 | 791.7 | 884.2 | 763.2 |
| 87 | 869.1 | 983.7 | 791.9 | 890.4 | 764.0 |
| 88 | 874.7 | 994.1 | 792.1 | 896.7 | 766.1 |
| 89 | 882.1 | 995.4 | 792.4 | 907.2 | 769.4 |
| 90 | 891.1 | 996.8 | 793.7 | 917.8 | 772.6 |
| 91 | 919.9 | 998.1 | 795.4 | 928.3 | 775.9 |
| 92 | 947.1 | 999.5 | 797.2 | 938.9 | 778.8 |
| 93 | 956.0 | 1000.9 | 805.0 | 1009.6 | 780.8 |
| 94 | 983.2 | 1002.4 | 849.6 | 1080.3 | 782.9 |
| 95 | 998.1 | 1003.8 | 894.2 | 1151.0 | 784.9 |
| 96 | 1004.1 | 1005.3 | 938.7 | 1221.7 | 788.8 |
| 97 | 1094.0 | 1080.1 | 982.1 | 1501.7 | 801.4 |
| 98 | 1210.5 | 1154.9 | 1024.6 | 1781.6 | 814.1 |
| 99 | 1300.4 | 1229.8 | 1067.2 | 2061.6 | 826.8 |
| 100 | 2341.5 | 1304.6 | 1109.7 | 2341.5 | 839.5 |

***Table S27:*** *Unpaced tapping – Rate (ITI in ms) of spontaneous tapping, final trial – Unpaced_spontaneous_final_right_mean_iti*

| | | | | | | |
|---|---|---|---|---|---|---|
| **Task** | | Unpaced tapping | | | | |
| **Outcome measure** | | Rate (ITI in ms) of spontaneous tapping, final trial | | | | |
| **Variable name** | | Unpaced_spontaneous_final_right_mean_iti | | | | |
| **Age and gender effects** | | | | | | |
| Age regression (slope) | | 0.216 | | | | |
| Age regression (p) | | 0.648 | | | | |
| Age group (p) | | 0.945 | | | | |
| Gender (p) | | 0.940 | | | | |
| **Group** | | All | Age 18 to 21 | Age 22 to 29 | Age 30 to 54 | Age 55 to 87 |
| N (N female) | | 104 (72) | 26 (17) | 28 (19) | 25 (16) | 25 (20) |
| **Normality** | | | | | | |
| Skewness | | 4.86 | 0.62 | 0.49 | 4.10 | 0.56 |
| Excess kurtosis | | 35.75 | -0.42 | -0.23 | 16.56 | 2.73 |
| **Scores** | | | | | | |
| Mean | | 626.6 | 626.3 | 605.0 | 655.9 | 621.8 |
| SD | | 177.3 | 119.6 | 88.9 | 297.4 | 145.8 |
| Percentiles | | | | | | |
| | 0 | 250.6 | 459.0 | 450.3 | 401.7 | 250.6 |
| | 1 | 403.2 | 460.6 | 460.3 | 413.9 | 302.5 |
| | 2 | 450.4 | 462.2 | 470.4 | 426.1 | 354.5 |
| | 3 | 453.1 | 463.8 | 480.4 | 438.3 | 406.4 |
| | 4 | 459.7 | 465.4 | 487.5 | 450.5 | 458.4 |
| | 5 | 465.6 | 466.8 | 487.7 | 461.7 | 478.4 |
| | 6 | 467.7 | 468.2 | 487.8 | 472.7 | 492.0 |
| | 7 | 474.4 | 469.6 | 488.0 | 483.6 | 505.7 |
| | 8 | 487.6 | 471.0 | 491.0 | 494.6 | 519.4 |
| | 9 | 490.8 | 480.0 | 495.9 | 503.9 | 524.5 |
| | 10 | 500.7 | 488.9 | 500.8 | 512.4 | 525.3 |
| | 11 | 506.5 | 497.9 | 505.7 | 520.8 | 526.1 |
| | 12 | 508.4 | 506.9 | 509.0 | 529.3 | 526.9 |
| | 13 | 513.7 | 507.9 | 512.2 | 536.5 | 527.9 |
| | 14 | 518.3 | 508.9 | 515.4 | 542.5 | 529.1 |
| | 15 | 520.7 | 509.9 | 518.0 | 548.4 | 530.2 |
| | 16 | 523.1 | 510.9 | 518.2 | 554.4 | 531.3 |
| | 17 | 523.5 | 514.0 | 518.4 | 559.2 | 532.5 |
| | 18 | 524.7 | 517.0 | 518.6 | 561.9 | 533.9 |
| | 19 | 526.5 | 520.1 | 519.3 | 564.5 | 535.4 |
| | 20 | 530.2 | 523.1 | 520.5 | 567.1 | 536.8 |
| | 21 | 533.0 | 523.7 | 521.6 | 569.3 | 538.1 |
| | 22 | 536.4 | 524.3 | 522.8 | 569.5 | 539.3 |

| | | | | | |
|---|---|---|---|---|---|
| 23 | 541.2 | 524.9 | 530.7 | 569.8 | 540.4 |
| 24 | 549.2 | 525.4 | 540.4 | 570.0 | 541.5 |
| 25 | 556.7 | 536.3 | 550.1 | 570.3 | 542.6 |
| 26 | 559.0 | 547.2 | 559.2 | 570.3 | 544.8 |
| 27 | 559.7 | 558.1 | 559.5 | 570.3 | 547.0 |
| 28 | 560.3 | 569.0 | 559.9 | 570.3 | 549.2 |
| 29 | 567.6 | 571.1 | 560.2 | 570.3 | 551.4 |
| 30 | 569.0 | 573.2 | 561.2 | 570.3 | 553.3 |
| 31 | 569.2 | 575.2 | 563.5 | 570.4 | 555.3 |
| 32 | 570.2 | 577.3 | 565.7 | 570.4 | 557.2 |
| 33 | 570.3 | 578.5 | 567.9 | 570.5 | 559.2 |
| 34 | 570.5 | 579.6 | 569.2 | 570.8 | 562.6 |
| 35 | 571.6 | 580.8 | 570.0 | 571.2 | 566.8 |
| 36 | 572.3 | 581.9 | 570.8 | 571.6 | 571.1 |
| 37 | 573.7 | 582.1 | 571.6 | 572.0 | 575.3 |
| 38 | 575.1 | 582.3 | 574.9 | 572.3 | 577.7 |
| 39 | 577.3 | 582.4 | 578.4 | 572.7 | 578.3 |
| 40 | 577.9 | 582.6 | 581.8 | 573.0 | 578.8 |
| 41 | 580.3 | 583.0 | 584.7 | 573.3 | 579.4 |
| 42 | 582.1 | 583.4 | 586.3 | 573.7 | 580.2 |
| 43 | 583.0 | 583.7 | 587.9 | 573.9 | 581.6 |
| 44 | 584.2 | 584.1 | 589.5 | 574.2 | 582.9 |
| 45 | 584.7 | 584.5 | 590.4 | 574.5 | 584.3 |
| 46 | 585.5 | 584.8 | 590.7 | 575.2 | 586.6 |
| 47 | 586.1 | 585.2 | 590.9 | 578.1 | 593.4 |
| 48 | 588.3 | 585.6 | 591.2 | 580.9 | 600.2 |
| 49 | 590.7 | 588.1 | 592.8 | 583.8 | 607.0 |
| 50 | 593.4 | 590.5 | 594.6 | 586.7 | 613.9 |
| 51 | 596.7 | 593.0 | 596.3 | 589.5 | 614.5 |
| 52 | 598.1 | 595.5 | 598.0 | 592.3 | 615.1 |
| 53 | 600.7 | 600.9 | 599.3 | 595.1 | 615.7 |
| 54 | 603.6 | 606.3 | 600.5 | 597.9 | 616.3 |
| 55 | 608.9 | 611.7 | 601.7 | 599.6 | 619.8 |
| 56 | 612.6 | 617.1 | 603.7 | 601.0 | 623.9 |
| 57 | 613.6 | 618.8 | 606.6 | 602.4 | 628.1 |
| 58 | 615.2 | 620.4 | 609.5 | 603.9 | 632.2 |
| 59 | 616.2 | 622.1 | 612.4 | 605.5 | 634.8 |
| 60 | 617.0 | 623.8 | 613.6 | 607.2 | 636.6 |
| 61 | 622.6 | 630.2 | 614.3 | 608.8 | 638.4 |
| 62 | 624.0 | 636.7 | 615.0 | 610.5 | 640.3 |
| 63 | 626.9 | 643.2 | 615.7 | 613.9 | 641.9 |
| 64 | 632.4 | 649.6 | 618.0 | 619.1 | 643.2 |
| 65 | 633.5 | 649.7 | 620.2 | 624.3 | 644.5 |
| 66 | 641.0 | 649.8 | 622.5 | 629.4 | 645.9 |

| | | | | | |
|---|---|---|---|---|---|
| 67 | 642.8 | 649.8 | 624.3 | 633.7 | 646.9 |
| 68 | 646.8 | 649.9 | 625.2 | 636.0 | 647.2 |
| 69 | 648.2 | 658.9 | 626.1 | 638.4 | 647.6 |
| 70 | 648.4 | 667.9 | 627.0 | 640.8 | 647.9 |
| 71 | 649.7 | 676.8 | 632.5 | 643.0 | 650.3 |
| 72 | 651.2 | 685.8 | 640.7 | 644.3 | 662.5 |
| 73 | 658.1 | 693.8 | 648.9 | 645.6 | 674.8 |
| 74 | 663.3 | 701.8 | 657.2 | 646.9 | 687.1 |
| 75 | 677.2 | 709.8 | 658.2 | 648.2 | 699.4 |
| 76 | 680.3 | 717.8 | 658.6 | 655.1 | 703.0 |
| 77 | 690.0 | 719.0 | 659.1 | 662.0 | 706.6 |
| 78 | 700.4 | 720.2 | 662.0 | 668.9 | 710.3 |
| 79 | 703.1 | 721.4 | 673.6 | 675.8 | 713.9 |
| 80 | 705.8 | 722.7 | 685.3 | 677.2 | 715.8 |
| 81 | 710.9 | 728.4 | 696.9 | 677.5 | 717.4 |
| 82 | 716.0 | 734.2 | 702.8 | 677.7 | 719.0 |
| 83 | 717.9 | 740.0 | 703.2 | 678.0 | 720.5 |
| 84 | 718.6 | 745.7 | 703.6 | 684.5 | 721.3 |
| 85 | 720.2 | 751.0 | 704.0 | 694.1 | 721.6 |
| 86 | 721.9 | 756.3 | 705.0 | 703.6 | 722.0 |
| 87 | 722.6 | 761.5 | 706.1 | 713.2 | 722.4 |
| 88 | 736.3 | 766.8 | 707.2 | 722.5 | 725.1 |
| 89 | 745.2 | 780.4 | 708.5 | 731.6 | 730.3 |
| 90 | 752.8 | 794.0 | 711.5 | 740.7 | 735.4 |
| 91 | 763.9 | 807.6 | 714.5 | 749.8 | 740.6 |
| 92 | 772.0 | 821.2 | 717.4 | 757.3 | 752.6 |
| 93 | 775.7 | 828.2 | 725.5 | 761.6 | 778.1 |
| 94 | 807.9 | 835.1 | 740.9 | 765.8 | 803.6 |
| 95 | 820.3 | 842.0 | 756.3 | 770.1 | 829.2 |
| 96 | 845.6 | 849.0 | 771.7 | 823.7 | 858.8 |
| 97 | 850.3 | 860.9 | 783.6 | 1124.1 | 908.7 |
| 98 | 894.0 | 872.9 | 794.0 | 1424.5 | 958.7 |
| 99 | 1053.7 | 884.8 | 804.5 | 1725.0 | 1008.6 |
| 100 | 2025.4 | 896.7 | 814.9 | 2025.4 | 1058.6 |

***Table S28:*** *Unpaced tapping – Rate (ITI in ms) of fast tapping – Unpaced_fast_mean_iti*

| | | | | | | |
|---|---|---|---|---|---|---|
| **Task** | | Unpaced tapping | | | | |
| **Outcome measure** | | Rate (ITI in ms) of fast tapping | | | | |
| **Variable name** | | Unpaced_fast_mean_iti | | | | |
| **Age and gender effects** | | | | | | |
| Age regression (slope) | | 0.849 | | | | |
| Age regression (p) | | 0.122 | | | | |
| Age group (p) | | 0.030 | | | | |
| Gender (p) | | 0.477 | | | | |
| **Group** | | All | Age 18 to 21 | Age 22 to 29 | Age 30 to 54 | Age 55 to 87 |
| N (N female) | | 107 (73) | 27 (18) | 29 (19) | 26 (16) | 25 (20) |
| **Normality** | | | | | | |
| Skewness | | 0.96 | 0.44 | 1.63 | 1.01 | 0.16 |
| Excess kurtosis | | 1.03 | -0.83 | 3.36 | -0.27 | -1.13 |
| **Scores** | | | | | | |
| Mean | | 333.9 | 345.3 | 334.3 | 276.6 | 380.7 |
| SD | | 134.5 | 132.7 | 153.5 | 105.0 | 126.0 |
| Percentiles | | | | | | |
| | 0 | 161.0 | 176.2 | 168.8 | 161.0 | 195.5 |
| | 1 | 169.1 | 176.5 | 171.2 | 164.4 | 195.7 |
| | 2 | 174.9 | 176.8 | 173.6 | 167.9 | 195.9 |
| | 3 | 176.4 | 177.1 | 176.0 | 171.3 | 196.1 |
| | 4 | 177.4 | 177.6 | 178.4 | 174.7 | 196.3 |
| | 5 | 178.5 | 178.8 | 180.7 | 176.3 | 201.1 |
| | 6 | 181.5 | 180.0 | 183.0 | 177.9 | 206.9 |
| | 7 | 183.6 | 181.2 | 185.4 | 179.5 | 212.6 |
| | 8 | 186.8 | 182.7 | 186.2 | 181.1 | 218.4 |
| | 9 | 188.0 | 184.7 | 186.9 | 182.9 | 222.9 |
| | 10 | 188.7 | 186.8 | 187.5 | 184.6 | 226.7 |
| | 11 | 189.7 | 188.8 | 189.2 | 186.3 | 230.5 |
| | 12 | 191.2 | 190.5 | 193.8 | 188.0 | 234.3 |
| | 13 | 194.0 | 191.7 | 198.3 | 188.3 | 236.3 |
| | 14 | 195.4 | 193.0 | 202.8 | 188.6 | 236.4 |
| | 15 | 196.3 | 194.2 | 205.8 | 188.9 | 236.6 |
| | 16 | 196.7 | 199.9 | 208.3 | 189.3 | 236.8 |
| | 17 | 201.1 | 208.3 | 210.7 | 189.9 | 237.3 |
| | 18 | 204.2 | 216.7 | 213.0 | 190.5 | 238.7 |
| | 19 | 205.8 | 225.2 | 213.9 | 191.1 | 240.1 |
| | 20 | 210.0 | 228.5 | 214.7 | 191.7 | 241.5 |
| | 21 | 213.7 | 230.4 | 215.6 | 192.9 | 243.7 |
| | 22 | 216.2 | 232.2 | 217.2 | 194.2 | 250.3 |
| | 23 | 217.9 | 234.0 | 219.1 | 195.5 | 256.8 |
| | 24 | 221.4 | 236.3 | 221.1 | 196.8 | 263.4 |

| | | | | | |
|---|---|---|---|---|---|
| 25 | 223.0 | 238.6 | 223.1 | 197.8 | 269.9 |
| 26 | 223.4 | 240.9 | 223.3 | 198.9 | 279.3 |
| 27 | 225.8 | 243.1 | 223.4 | 200.0 | 288.6 |
| 28 | 227.1 | 244.3 | 223.5 | 201.1 | 298.0 |
| 29 | 232.4 | 245.5 | 224.0 | 202.1 | 307.3 |
| 30 | 235.8 | 246.7 | 225.0 | 203.1 | 312.0 |
| 31 | 236.8 | 248.9 | 226.0 | 204.2 | 315.7 |
| 32 | 238.1 | 254.2 | 227.0 | 205.2 | 319.3 |
| 33 | 242.5 | 259.5 | 231.4 | 206.2 | 323.0 |
| 34 | 243.1 | 264.8 | 236.4 | 207.2 | 326.6 |
| 35 | 245.2 | 268.3 | 241.4 | 208.3 | 330.1 |
| 36 | 247.7 | 268.9 | 247.0 | 209.3 | 333.6 |
| 37 | 248.3 | 269.5 | 254.2 | 211.1 | 337.1 |
| 38 | 250.5 | 270.1 | 261.4 | 212.9 | 339.7 |
| 39 | 259.5 | 272.3 | 268.5 | 214.7 | 341.4 |
| 40 | 268.8 | 275.8 | 271.5 | 216.5 | 343.0 |
| 41 | 270.2 | 279.2 | 272.8 | 218.1 | 344.7 |
| 42 | 270.5 | 282.7 | 274.2 | 219.7 | 347.1 |
| 43 | 273.3 | 285.0 | 275.8 | 221.3 | 351.1 |
| 44 | 280.7 | 286.7 | 279.8 | 222.9 | 355.0 |
| 45 | 287.7 | 288.5 | 283.7 | 226.7 | 359.0 |
| 46 | 290.2 | 290.2 | 287.7 | 230.5 | 362.3 |
| 47 | 292.2 | 294.9 | 290.5 | 234.4 | 362.5 |
| 48 | 293.5 | 300.1 | 292.4 | 238.2 | 362.7 |
| 49 | 293.7 | 305.3 | 294.3 | 240.7 | 362.8 |
| 50 | 296.2 | 310.5 | 296.2 | 243.2 | 363.0 |
| 51 | 296.5 | 316.8 | 296.2 | 245.7 | 365.2 |
| 52 | 302.3 | 323.1 | 296.2 | 248.2 | 367.3 |
| 53 | 309.2 | 329.4 | 296.2 | 248.3 | 369.5 |
| 54 | 310.8 | 336.2 | 296.8 | 248.4 | 371.6 |
| 55 | 315.4 | 345.5 | 298.3 | 248.6 | 378.0 |
| 56 | 326.4 | 354.8 | 299.7 | 248.7 | 385.1 |
| 57 | 332.2 | 364.0 | 301.1 | 250.3 | 392.3 |
| 58 | 336.8 | 371.6 | 308.3 | 252.0 | 399.5 |
| 59 | 342.6 | 375.4 | 316.4 | 253.6 | 405.2 |
| 60 | 347.4 | 379.3 | 324.5 | 255.2 | 410.2 |
| 61 | 357.6 | 383.1 | 333.3 | 264.5 | 415.3 |
| 62 | 362.8 | 390.4 | 344.1 | 273.9 | 420.3 |
| 63 | 367.5 | 401.7 | 354.9 | 283.3 | 425.5 |
| 64 | 370.2 | 412.9 | 365.6 | 292.6 | 430.8 |
| 65 | 371.8 | 424.2 | 372.3 | 292.9 | 436.1 |
| 66 | 384.6 | 431.1 | 377.4 | 293.1 | 441.4 |
| 67 | 387.1 | 435.4 | 382.5 | 293.3 | 445.9 |
| 68 | 401.3 | 439.6 | 387.4 | 293.6 | 448.5 |

| | | | | | |
|---|---|---|---|---|---|
| 69 | 404.8 | 443.8 | 391.5 | 293.6 | 451.2 |
| 70 | 423.5 | 445.6 | 395.5 | 293.7 | 453.9 |
| 71 | 426.4 | 446.5 | 399.6 | 293.7 | 456.5 |
| 72 | 427.1 | 447.5 | 405.3 | 293.7 | 459.0 |
| 73 | 428.4 | 448.5 | 412.3 | 298.2 | 461.4 |
| 74 | 431.8 | 450.6 | 419.4 | 302.6 | 463.9 |
| 75 | 440.4 | 452.9 | 426.4 | 307.1 | 466.3 |
| 76 | 444.9 | 455.1 | 426.4 | 311.6 | 473.9 |
| 77 | 447.2 | 457.3 | 426.4 | 320.8 | 481.5 |
| 78 | 453.3 | 458.8 | 426.5 | 330.0 | 489.1 |
| 79 | 455.6 | 460.3 | 426.7 | 339.2 | 496.6 |
| 80 | 455.8 | 461.8 | 427.2 | 348.4 | 504.2 |
| 81 | 456.1 | 463.6 | 427.7 | 370.3 | 511.6 |
| 82 | 457.1 | 466.4 | 428.3 | 392.2 | 519.1 |
| 83 | 462.9 | 469.1 | 439.9 | 414.1 | 526.6 |
| 84 | 466.6 | 471.9 | 453.3 | 436.0 | 532.1 |
| 85 | 473.8 | 477.6 | 466.8 | 440.9 | 536.6 |
| 86 | 478.1 | 488.2 | 477.3 | 445.8 | 541.0 |
| 87 | 489.5 | 498.8 | 480.3 | 450.6 | 545.5 |
| 88 | 499.5 | 509.4 | 483.3 | 455.5 | 549.2 |
| 89 | 507.3 | 516.1 | 486.3 | 455.5 | 552.3 |
| 90 | 516.8 | 519.4 | 490.4 | 455.6 | 555.4 |
| 91 | 523.5 | 522.7 | 495.1 | 455.6 | 558.5 |
| 92 | 527.8 | 526.0 | 499.7 | 455.7 | 561.6 |
| 93 | 528.9 | 527.3 | 506.8 | 455.7 | 564.8 |
| 94 | 541.0 | 527.7 | 528.6 | 455.8 | 567.9 |
| 95 | 556.7 | 528.1 | 550.5 | 455.8 | 571.0 |
| 96 | 570.5 | 528.5 | 572.4 | 455.9 | 574.8 |
| 97 | 580.3 | 555.7 | 628.8 | 472.0 | 581.9 |
| 98 | 600.8 | 587.9 | 711.3 | 488.2 | 589.1 |
| 99 | 649.1 | 620.0 | 793.7 | 504.3 | 596.2 |
| 100 | 876.1 | 652.1 | 876.1 | 520.4 | 603.3 |

***Table S29:*** *Unpaced tapping – Rate (ITI in ms) of slow tapping – Unpaced_slow_mean_iti*

| | | | | | | |
|---|---|---|---|---|---|---|
| **Task** | | Unpaced tapping | | | | |
| **Outcome measure** | | Rate (ITI in ms) of slow tapping | | | | |
| **Variable name** | | Unpaced_slow_mean_iti | | | | |
| **Age and gender effects** | | | | | | |
| Age regression (slope) | | -1.195 | | | | |
| Age regression (p) | | 0.510 | | | | |
| Age group (p) | | 0.073 | | | | |
| Gender (p) | | 0.395 | | | | |
| **Group** | | All | Age 18 to 21 | Age 22 to 29 | Age 30 to 54 | Age 55 to 87 |
| N (N female) | | 103 (71) | 25 (16) | 28 (19) | 25 (16) | 25 (20) |
| **Normality** | | | | | | |
| Skewness | | 1.18 | 0.86 | 2.14 | 0.63 | 1.42 |
| Excess kurtosis | | 0.66 | -0.09 | 4.73 | -0.99 | 1.58 |
| **Scores** | | | | | | |
| Mean | | 1230.1 | 1325.9 | 1107.3 | 1351.7 | 1150.3 |
| SD | | 465.8 | 394.6 | 440.5 | 545.3 | 448.8 |
| Percentiles | | | | | | |
| | 0 | 564.3 | 835.7 | 687.6 | 599.9 | 564.3 |
| | 1 | 601.3 | 854.2 | 689.1 | 640.0 | 589.8 |
| | 2 | 671.2 | 872.7 | 690.7 | 680.2 | 615.3 |
| | 3 | 687.9 | 891.2 | 692.3 | 720.3 | 640.8 |
| | 4 | 696.1 | 909.7 | 700.4 | 760.4 | 666.2 |
| | 5 | 729.6 | 917.1 | 724.1 | 776.6 | 681.9 |
| | 6 | 750.6 | 922.4 | 747.9 | 788.1 | 695.6 |
| | 7 | 769.1 | 927.6 | 771.7 | 799.5 | 709.3 |
| | 8 | 782.0 | 932.9 | 782.0 | 811.0 | 723.0 |
| | 9 | 787.2 | 937.8 | 783.1 | 816.1 | 730.9 |
| | 10 | 795.9 | 942.4 | 784.2 | 818.0 | 735.9 |
| | 11 | 798.0 | 947.1 | 785.3 | 820.0 | 740.8 |
| | 12 | 805.5 | 951.7 | 788.1 | 821.9 | 745.8 |
| | 13 | 815.3 | 955.4 | 791.2 | 830.1 | 754.0 |
| | 14 | 817.1 | 958.1 | 794.2 | 844.4 | 765.4 |
| | 15 | 818.7 | 960.9 | 797.0 | 858.8 | 776.7 |
| | 16 | 821.7 | 963.6 | 798.6 | 873.1 | 788.1 |
| | 17 | 827.2 | 966.5 | 800.2 | 884.6 | 797.4 |
| | 18 | 847.1 | 969.6 | 801.8 | 890.3 | 802.5 |
| | 19 | 873.3 | 972.7 | 804.6 | 896.1 | 807.5 |
| | 20 | 887.5 | 975.8 | 808.6 | 901.8 | 812.6 |
| | 21 | 899.7 | 979.6 | 812.7 | 907.3 | 821.6 |
| | 22 | 907.3 | 986.7 | 816.8 | 911.7 | 850.0 |
| | 23 | 910.3 | 993.8 | 818.4 | 916.0 | 878.4 |
| | 24 | 916.2 | 1000.9 | 819.3 | 920.4 | 906.9 |

| | | | | | |
|---|---|---|---|---|---|
| 25 | 922.4 | 1008.0 | 820.2 | 924.8 | 935.3 |
| 26 | 929.9 | 1012.4 | 822.0 | 931.5 | 939.6 |
| 27 | 935.0 | 1016.8 | 834.6 | 938.2 | 943.9 |
| 28 | 939.1 | 1021.1 | 847.1 | 944.9 | 948.3 |
| 29 | 948.3 | 1025.5 | 859.6 | 951.6 | 952.6 |
| 30 | 953.1 | 1028.6 | 870.2 | 964.1 | 954.0 |
| 31 | 953.8 | 1031.5 | 877.6 | 977.8 | 954.9 |
| 32 | 955.9 | 1034.4 | 884.9 | 991.5 | 955.7 |
| 33 | 956.9 | 1037.2 | 892.3 | 1005.2 | 956.6 |
| 34 | 958.6 | 1039.8 | 897.2 | 1018.4 | 956.9 |
| 35 | 961.8 | 1042.1 | 900.8 | 1031.4 | 956.9 |
| 36 | 964.7 | 1044.5 | 904.5 | 1044.4 | 956.9 |
| 37 | 975.0 | 1046.8 | 908.1 | 1057.3 | 956.9 |
| 38 | 980.2 | 1058.0 | 911.3 | 1064.7 | 959.8 |
| 39 | 1002.0 | 1077.9 | 914.5 | 1066.6 | 965.5 |
| 40 | 1009.4 | 1097.9 | 917.7 | 1068.5 | 971.3 |
| 41 | 1015.0 | 1117.9 | 921.6 | 1070.4 | 977.0 |
| 42 | 1024.6 | 1139.2 | 927.6 | 1072.8 | 983.6 |
| 43 | 1031.8 | 1163.1 | 933.5 | 1076.2 | 992.1 |
| 44 | 1037.5 | 1187.0 | 939.5 | 1079.6 | 1000.6 |
| 45 | 1038.9 | 1211.0 | 944.8 | 1083.0 | 1009.1 |
| 46 | 1047.3 | 1231.5 | 949.4 | 1090.7 | 1017.1 |
| 47 | 1054.1 | 1234.8 | 954.1 | 1119.8 | 1022.6 |
| 48 | 1057.5 | 1238.1 | 958.7 | 1149.0 | 1028.1 |
| 49 | 1063.7 | 1241.4 | 960.2 | 1178.1 | 1033.5 |
| 50 | 1069.4 | 1244.8 | 961.1 | 1207.3 | 1039.0 |
| 51 | 1071.9 | 1267.9 | 962.1 | 1208.0 | 1043.5 |
| 52 | 1086.7 | 1291.1 | 965.7 | 1208.7 | 1048.0 |
| 53 | 1108.5 | 1314.2 | 984.5 | 1209.4 | 1052.4 |
| 54 | 1117.7 | 1337.4 | 1003.4 | 1210.2 | 1056.9 |
| 55 | 1119.8 | 1341.7 | 1022.2 | 1217.8 | 1060.0 |
| 56 | 1124.3 | 1342.3 | 1035.3 | 1226.8 | 1062.8 |
| 57 | 1134.1 | 1342.9 | 1041.2 | 1235.8 | 1065.6 |
| 58 | 1157.8 | 1343.5 | 1047.1 | 1244.8 | 1068.5 |
| 59 | 1187.2 | 1344.6 | 1053.0 | 1258.6 | 1077.1 |
| 60 | 1190.1 | 1346.0 | 1065.2 | 1274.7 | 1088.6 |
| 61 | 1199.7 | 1347.4 | 1079.6 | 1290.8 | 1100.2 |
| 62 | 1208.0 | 1348.8 | 1094.1 | 1306.9 | 1111.7 |
| 63 | 1215.6 | 1353.1 | 1108.1 | 1327.3 | 1126.0 |
| 64 | 1234.8 | 1360.5 | 1111.2 | 1352.0 | 1143.0 |
| 65 | 1245.7 | 1367.9 | 1114.3 | 1376.7 | 1159.9 |
| 66 | 1250.5 | 1375.2 | 1117.4 | 1401.5 | 1176.9 |
| 67 | 1263.1 | 1386.5 | 1119.8 | 1433.3 | 1188.9 |
| 68 | 1277.9 | 1405.5 | 1120.9 | 1479.3 | 1191.2 |

| | | | | | |
|---|---|---|---|---|---|
| 69 | 1293.7 | 1424.5 | 1121.9 | 1525.4 | 1193.4 |
| 70 | 1325.5 | 1443.6 | 1123.0 | 1571.4 | 1195.7 |
| 71 | 1342.3 | 1460.0 | 1128.3 | 1615.8 | 1200.7 |
| 72 | 1346.2 | 1463.4 | 1136.1 | 1651.9 | 1219.6 |
| 73 | 1354.8 | 1466.8 | 1143.9 | 1687.9 | 1238.5 |
| 74 | 1364.8 | 1470.2 | 1151.7 | 1724.0 | 1257.4 |
| 75 | 1374.5 | 1473.6 | 1160.9 | 1760.0 | 1276.3 |
| 76 | 1399.8 | 1533.8 | 1170.3 | 1838.3 | 1296.6 |
| 77 | 1430.6 | 1594.0 | 1179.7 | 1916.6 | 1317.0 |
| 78 | 1451.5 | 1654.2 | 1191.1 | 1994.9 | 1337.3 |
| 79 | 1467.7 | 1714.4 | 1209.9 | 2073.2 | 1357.7 |
| 80 | 1555.3 | 1725.9 | 1228.6 | 2092.7 | 1362.6 |
| 81 | 1634.3 | 1727.6 | 1247.3 | 2100.5 | 1364.5 |
| 82 | 1665.1 | 1729.4 | 1259.7 | 2108.2 | 1366.4 |
| 83 | 1707.3 | 1731.1 | 1266.3 | 2116.0 | 1368.2 |
| 84 | 1729.4 | 1752.0 | 1272.9 | 2118.9 | 1380.4 |
| 85 | 1751.5 | 1782.3 | 1279.4 | 2119.3 | 1397.9 |
| 86 | 1830.8 | 1812.7 | 1361.8 | 2119.8 | 1415.3 |
| 87 | 1895.7 | 1843.1 | 1461.3 | 2120.3 | 1432.7 |
| 88 | 1924.3 | 1864.4 | 1560.9 | 2122.5 | 1500.9 |
| 89 | 1935.7 | 1876.5 | 1650.1 | 2126.3 | 1620.0 |
| 90 | 2002.6 | 1888.7 | 1656.7 | 2130.2 | 1739.0 |
| 91 | 2074.1 | 1900.8 | 1663.4 | 2134.0 | 1858.1 |
| 92 | 2113.4 | 1910.5 | 1670.1 | 2145.1 | 1956.0 |
| 93 | 2120.3 | 1915.4 | 1712.0 | 2170.6 | 2011.4 |
| 94 | 2134.6 | 1920.2 | 1805.0 | 2196.1 | 2066.8 |
| 95 | 2165.3 | 1925.1 | 1898.1 | 2221.6 | 2122.3 |
| 96 | 2236.9 | 1945.5 | 1991.2 | 2249.8 | 2178.2 |
| 97 | 2332.9 | 2043.8 | 2149.8 | 2291.5 | 2236.9 |
| 98 | 2410.1 | 2142.1 | 2336.0 | 2333.3 | 2295.6 |
| 99 | 2416.6 | 2240.3 | 2522.2 | 2375.0 | 2354.3 |
| 100 | 2708.4 | 2338.6 | 2708.4 | 2416.7 | 2413.1 |

***Table S30:*** *Unpaced tapping – Variability (CV of ITI) of spontaneous tapping, initial trial – Unpaced_spontaneous_right_CV_iti*

| | | | | | |
|---|---|---|---|---|---|
| **Task** | Unpaced tapping | | | | |
| **Outcome measure** | Variability (CV of ITI) of spontaneous tapping, initial trial | | | | |
| **Variable name** | Unpaced_spontaneous_right_CV_iti | | | | |
| **Age and gender effects** | | | | | |
| Age regression (slope) | 0.000 | | | | |
| Age regression (p) | 0.043 | | | | |
| Age group (p) | 0.380 | | | | |
| Gender (p) | 0.792 | | | | |
| **Group** | All | Age 18 to 21 | Age 22 to 29 | Age 30 to 54 | Age 55 to 87 |
| N (N female) | 106 (72) | 26 (17) | 29 (19) | 26 (16) | 25 (20) |
| **Normality** | | | | | |
| Skewness | 5.59 | 4.17 | 4.32 | 0.57 | 2.68 |
| Excess kurtosis | 34.90 | 17.00 | 18.88 | -0.61 | 8.97 |
| **Scores** | | | | | |
| Mean | 0.057 | 0.066 | 0.062 | 0.051 | 0.049 |
| SD | 0.039 | 0.059 | 0.046 | 0.015 | 0.016 |
| Percentiles | | | | | |
| 0 | 0.027 | 0.036 | 0.035 | 0.031 | 0.027 |
| 1 | 0.031 | 0.036 | 0.035 | 0.031 | 0.028 |
| 2 | 0.032 | 0.036 | 0.035 | 0.032 | 0.030 |
| 3 | 0.033 | 0.036 | 0.035 | 0.032 | 0.031 |
| 4 | 0.034 | 0.036 | 0.035 | 0.032 | 0.033 |
| 5 | 0.034 | 0.036 | 0.036 | 0.033 | 0.034 |
| 6 | 0.035 | 0.036 | 0.036 | 0.033 | 0.035 |
| 7 | 0.036 | 0.037 | 0.036 | 0.033 | 0.036 |
| 8 | 0.036 | 0.037 | 0.036 | 0.034 | 0.037 |
| 9 | 0.036 | 0.037 | 0.036 | 0.034 | 0.037 |
| 10 | 0.036 | 0.037 | 0.037 | 0.034 | 0.038 |
| 11 | 0.037 | 0.037 | 0.037 | 0.034 | 0.038 |
| 12 | 0.037 | 0.037 | 0.037 | 0.034 | 0.038 |
| 13 | 0.037 | 0.038 | 0.037 | 0.035 | 0.039 |
| 14 | 0.037 | 0.038 | 0.037 | 0.036 | 0.039 |
| 15 | 0.038 | 0.039 | 0.037 | 0.037 | 0.039 |
| 16 | 0.038 | 0.040 | 0.038 | 0.038 | 0.039 |
| 17 | 0.038 | 0.040 | 0.038 | 0.038 | 0.040 |
| 18 | 0.039 | 0.041 | 0.038 | 0.038 | 0.040 |
| 19 | 0.039 | 0.042 | 0.039 | 0.039 | 0.040 |
| 20 | 0.040 | 0.042 | 0.040 | 0.039 | 0.040 |
| 21 | 0.040 | 0.042 | 0.041 | 0.039 | 0.041 |
| 22 | 0.041 | 0.043 | 0.042 | 0.039 | 0.041 |

| | | | | | |
|---|---|---|---|---|---|
| 23 | 0.041 | 0.043 | 0.042 | 0.040 | 0.041 |
| 24 | 0.041 | 0.043 | 0.043 | 0.040 | 0.042 |
| 25 | 0.041 | 0.043 | 0.044 | 0.040 | 0.042 |
| 26 | 0.042 | 0.043 | 0.044 | 0.040 | 0.042 |
| 27 | 0.042 | 0.043 | 0.045 | 0.041 | 0.042 |
| 28 | 0.042 | 0.044 | 0.045 | 0.041 | 0.042 |
| 29 | 0.043 | 0.044 | 0.046 | 0.041 | 0.043 |
| 30 | 0.043 | 0.044 | 0.046 | 0.041 | 0.043 |
| 31 | 0.043 | 0.044 | 0.046 | 0.041 | 0.043 |
| 32 | 0.043 | 0.044 | 0.046 | 0.041 | 0.043 |
| 33 | 0.043 | 0.045 | 0.046 | 0.041 | 0.043 |
| 34 | 0.043 | 0.045 | 0.047 | 0.041 | 0.043 |
| 35 | 0.044 | 0.046 | 0.047 | 0.041 | 0.043 |
| 36 | 0.044 | 0.047 | 0.047 | 0.041 | 0.043 |
| 37 | 0.045 | 0.047 | 0.047 | 0.042 | 0.043 |
| 38 | 0.045 | 0.047 | 0.047 | 0.042 | 0.043 |
| 39 | 0.046 | 0.047 | 0.048 | 0.043 | 0.043 |
| 40 | 0.046 | 0.048 | 0.048 | 0.043 | 0.043 |
| 41 | 0.047 | 0.048 | 0.049 | 0.044 | 0.043 |
| 42 | 0.047 | 0.048 | 0.050 | 0.044 | 0.043 |
| 43 | 0.047 | 0.049 | 0.050 | 0.045 | 0.044 |
| 44 | 0.048 | 0.049 | 0.051 | 0.045 | 0.044 |
| 45 | 0.048 | 0.050 | 0.052 | 0.046 | 0.045 |
| 46 | 0.048 | 0.051 | 0.052 | 0.047 | 0.045 |
| 47 | 0.049 | 0.052 | 0.053 | 0.048 | 0.045 |
| 48 | 0.049 | 0.053 | 0.053 | 0.048 | 0.046 |
| 49 | 0.049 | 0.053 | 0.054 | 0.048 | 0.046 |
| 50 | 0.050 | 0.053 | 0.054 | 0.049 | 0.047 |
| 51 | 0.050 | 0.053 | 0.054 | 0.049 | 0.047 |
| 52 | 0.050 | 0.053 | 0.054 | 0.049 | 0.047 |
| 53 | 0.050 | 0.054 | 0.054 | 0.050 | 0.048 |
| 54 | 0.052 | 0.054 | 0.054 | 0.050 | 0.048 |
| 55 | 0.052 | 0.054 | 0.055 | 0.051 | 0.048 |
| 56 | 0.053 | 0.054 | 0.055 | 0.052 | 0.048 |
| 57 | 0.053 | 0.054 | 0.055 | 0.053 | 0.049 |
| 58 | 0.053 | 0.055 | 0.055 | 0.053 | 0.049 |
| 59 | 0.054 | 0.055 | 0.055 | 0.054 | 0.049 |
| 60 | 0.054 | 0.055 | 0.055 | 0.055 | 0.049 |
| 61 | 0.054 | 0.056 | 0.055 | 0.055 | 0.050 |
| 62 | 0.054 | 0.056 | 0.055 | 0.055 | 0.050 |
| 63 | 0.055 | 0.056 | 0.055 | 0.055 | 0.050 |
| 64 | 0.055 | 0.057 | 0.055 | 0.055 | 0.050 |
| 65 | 0.055 | 0.057 | 0.055 | 0.056 | 0.050 |
| 66 | 0.055 | 0.058 | 0.055 | 0.056 | 0.050 |

| | | | | | |
|---|---|---|---|---|---|
| 67 | 0.055 | 0.059 | 0.055 | 0.057 | 0.050 |
| 68 | 0.055 | 0.060 | 0.055 | 0.058 | 0.050 |
| 69 | 0.055 | 0.061 | 0.056 | 0.058 | 0.050 |
| 70 | 0.055 | 0.061 | 0.056 | 0.059 | 0.050 |
| 71 | 0.056 | 0.061 | 0.057 | 0.060 | 0.051 |
| 72 | 0.057 | 0.062 | 0.057 | 0.060 | 0.051 |
| 73 | 0.057 | 0.062 | 0.058 | 0.060 | 0.051 |
| 74 | 0.059 | 0.062 | 0.059 | 0.060 | 0.052 |
| 75 | 0.060 | 0.062 | 0.059 | 0.061 | 0.052 |
| 76 | 0.060 | 0.063 | 0.060 | 0.061 | 0.053 |
| 77 | 0.061 | 0.063 | 0.061 | 0.062 | 0.053 |
| 78 | 0.061 | 0.064 | 0.062 | 0.062 | 0.054 |
| 79 | 0.062 | 0.064 | 0.063 | 0.063 | 0.054 |
| 80 | 0.063 | 0.065 | 0.063 | 0.064 | 0.054 |
| 81 | 0.063 | 0.066 | 0.064 | 0.064 | 0.054 |
| 82 | 0.064 | 0.066 | 0.064 | 0.065 | 0.054 |
| 83 | 0.064 | 0.067 | 0.067 | 0.065 | 0.055 |
| 84 | 0.065 | 0.067 | 0.071 | 0.065 | 0.055 |
| 85 | 0.065 | 0.070 | 0.074 | 0.066 | 0.055 |
| 86 | 0.067 | 0.073 | 0.077 | 0.068 | 0.055 |
| 87 | 0.068 | 0.076 | 0.080 | 0.069 | 0.055 |
| 88 | 0.072 | 0.079 | 0.082 | 0.071 | 0.055 |
| 89 | 0.075 | 0.079 | 0.084 | 0.071 | 0.057 |
| 90 | 0.077 | 0.079 | 0.085 | 0.072 | 0.058 |
| 91 | 0.078 | 0.079 | 0.085 | 0.073 | 0.060 |
| 92 | 0.079 | 0.079 | 0.085 | 0.073 | 0.061 |
| 93 | 0.082 | 0.088 | 0.085 | 0.074 | 0.063 |
| 94 | 0.085 | 0.098 | 0.085 | 0.075 | 0.064 |
| 95 | 0.085 | 0.108 | 0.085 | 0.076 | 0.065 |
| 96 | 0.085 | 0.118 | 0.085 | 0.078 | 0.068 |
| 97 | 0.109 | 0.174 | 0.118 | 0.079 | 0.080 |
| 98 | 0.118 | 0.231 | 0.174 | 0.081 | 0.091 |
| 99 | 0.280 | 0.287 | 0.231 | 0.083 | 0.102 |
| 100 | 0.343 | 0.343 | 0.288 | 0.085 | 0.114 |

***Table S31:*** *Unpaced tapping – Variability (CV of ITI) of spontaneous tapping, final trial – Unpaced_spontaneous_final_right_CV_iti*

| | | | | | |
|---|---|---|---|---|---|
| **Task** | Unpaced tapping | | | | |
| **Outcome measure** | Variability (CV of ITI) of spontaneous tapping, final trial | | | | |
| **Variable name** | Unpaced_spontaneous_final_right_CV_iti | | | | |
| **Age and gender effects** | | | | | |
| Age regression (slope) | 0.000 | | | | |
| Age regression (p) | 0.283 | | | | |
| Age group (p) | 0.515 | | | | |
| Gender (p) | 0.200 | | | | |
| **Group** | All | Age 18 to 21 | Age 22 to 29 | Age 30 to 54 | Age 55 to 87 |
| N (N female) | 104 (72) | 26 (17) | 28 (19) | 25 (16) | 25 (20) |
| **Normality** | | | | | |
| Skewness | 2.26 | 1.91 | 2.24 | 2.24 | 1.17 |
| Excess kurtosis | 6.60 | 4.03 | 5.52 | 6.56 | 1.20 |
| **Scores** | | | | | |
| Mean | 0.052 | 0.054 | 0.055 | 0.049 | 0.049 |
| SD | 0.018 | 0.020 | 0.022 | 0.018 | 0.012 |
| Percentiles | | | | | |
| 0 | 0.029 | 0.031 | 0.032 | 0.029 | 0.032 |
| 1 | 0.030 | 0.032 | 0.032 | 0.029 | 0.033 |
| 2 | 0.031 | 0.033 | 0.033 | 0.029 | 0.034 |
| 3 | 0.032 | 0.033 | 0.033 | 0.030 | 0.034 |
| 4 | 0.032 | 0.034 | 0.034 | 0.030 | 0.035 |
| 5 | 0.032 | 0.035 | 0.034 | 0.030 | 0.036 |
| 6 | 0.033 | 0.036 | 0.035 | 0.031 | 0.036 |
| 7 | 0.034 | 0.037 | 0.036 | 0.031 | 0.037 |
| 8 | 0.035 | 0.038 | 0.037 | 0.032 | 0.037 |
| 9 | 0.035 | 0.038 | 0.037 | 0.033 | 0.038 |
| 10 | 0.035 | 0.038 | 0.038 | 0.033 | 0.038 |
| 11 | 0.036 | 0.038 | 0.038 | 0.034 | 0.038 |
| 12 | 0.037 | 0.038 | 0.038 | 0.034 | 0.038 |
| 13 | 0.037 | 0.039 | 0.039 | 0.035 | 0.039 |
| 14 | 0.038 | 0.039 | 0.039 | 0.035 | 0.039 |
| 15 | 0.038 | 0.039 | 0.039 | 0.035 | 0.039 |
| 16 | 0.038 | 0.040 | 0.039 | 0.035 | 0.040 |
| 17 | 0.038 | 0.040 | 0.040 | 0.035 | 0.040 |
| 18 | 0.039 | 0.041 | 0.040 | 0.035 | 0.040 |
| 19 | 0.039 | 0.041 | 0.040 | 0.035 | 0.040 |
| 20 | 0.040 | 0.042 | 0.041 | 0.036 | 0.040 |
| 21 | 0.040 | 0.042 | 0.042 | 0.036 | 0.041 |
| 22 | 0.040 | 0.042 | 0.043 | 0.036 | 0.041 |

| | | | | | |
|---|---|---|---|---|---|
| 23 | 0.041 | 0.042 | 0.043 | 0.037 | 0.041 |
| 24 | 0.042 | 0.042 | 0.043 | 0.037 | 0.041 |
| 25 | 0.042 | 0.042 | 0.044 | 0.037 | 0.042 |
| 26 | 0.042 | 0.042 | 0.044 | 0.038 | 0.042 |
| 27 | 0.042 | 0.043 | 0.044 | 0.039 | 0.042 |
| 28 | 0.042 | 0.043 | 0.045 | 0.040 | 0.042 |
| 29 | 0.042 | 0.043 | 0.045 | 0.041 | 0.042 |
| 30 | 0.043 | 0.043 | 0.046 | 0.041 | 0.042 |
| 31 | 0.043 | 0.044 | 0.046 | 0.041 | 0.042 |
| 32 | 0.043 | 0.044 | 0.046 | 0.042 | 0.042 |
| 33 | 0.043 | 0.044 | 0.046 | 0.042 | 0.043 |
| 34 | 0.044 | 0.045 | 0.046 | 0.042 | 0.043 |
| 35 | 0.044 | 0.045 | 0.046 | 0.042 | 0.043 |
| 36 | 0.044 | 0.045 | 0.046 | 0.042 | 0.043 |
| 37 | 0.044 | 0.045 | 0.046 | 0.042 | 0.043 |
| 38 | 0.044 | 0.046 | 0.047 | 0.042 | 0.043 |
| 39 | 0.045 | 0.046 | 0.047 | 0.043 | 0.043 |
| 40 | 0.045 | 0.046 | 0.048 | 0.043 | 0.044 |
| 41 | 0.045 | 0.046 | 0.048 | 0.043 | 0.044 |
| 42 | 0.045 | 0.046 | 0.048 | 0.044 | 0.044 |
| 43 | 0.045 | 0.047 | 0.049 | 0.044 | 0.044 |
| 44 | 0.046 | 0.047 | 0.049 | 0.044 | 0.044 |
| 45 | 0.046 | 0.047 | 0.050 | 0.045 | 0.044 |
| 46 | 0.046 | 0.047 | 0.050 | 0.045 | 0.044 |
| 47 | 0.047 | 0.048 | 0.050 | 0.045 | 0.044 |
| 48 | 0.047 | 0.048 | 0.050 | 0.045 | 0.045 |
| 49 | 0.048 | 0.049 | 0.050 | 0.045 | 0.045 |
| 50 | 0.048 | 0.049 | 0.050 | 0.045 | 0.045 |
| 51 | 0.049 | 0.050 | 0.050 | 0.046 | 0.045 |
| 52 | 0.049 | 0.051 | 0.050 | 0.046 | 0.045 |
| 53 | 0.049 | 0.051 | 0.050 | 0.047 | 0.045 |
| 54 | 0.049 | 0.051 | 0.050 | 0.048 | 0.045 |
| 55 | 0.050 | 0.051 | 0.051 | 0.048 | 0.046 |
| 56 | 0.050 | 0.051 | 0.051 | 0.048 | 0.047 |
| 57 | 0.051 | 0.051 | 0.051 | 0.048 | 0.048 |
| 58 | 0.051 | 0.051 | 0.052 | 0.049 | 0.049 |
| 59 | 0.051 | 0.052 | 0.052 | 0.049 | 0.049 |
| 60 | 0.051 | 0.052 | 0.052 | 0.050 | 0.049 |
| 61 | 0.051 | 0.052 | 0.053 | 0.050 | 0.049 |
| 62 | 0.052 | 0.053 | 0.053 | 0.051 | 0.049 |
| 63 | 0.052 | 0.053 | 0.053 | 0.051 | 0.050 |
| 64 | 0.052 | 0.054 | 0.054 | 0.051 | 0.050 |
| 65 | 0.052 | 0.054 | 0.055 | 0.051 | 0.051 |
| 66 | 0.053 | 0.054 | 0.055 | 0.051 | 0.051 |

| | | | | | |
|---|---|---|---|---|---|
| 67 | 0.053 | 0.054 | 0.056 | 0.051 | 0.052 |
| 68 | 0.053 | 0.054 | 0.056 | 0.052 | 0.052 |
| 69 | 0.054 | 0.055 | 0.057 | 0.053 | 0.052 |
| 70 | 0.054 | 0.056 | 0.057 | 0.054 | 0.052 |
| 71 | 0.054 | 0.056 | 0.058 | 0.054 | 0.052 |
| 72 | 0.056 | 0.057 | 0.058 | 0.054 | 0.052 |
| 73 | 0.056 | 0.058 | 0.058 | 0.055 | 0.053 |
| 74 | 0.057 | 0.058 | 0.058 | 0.055 | 0.053 |
| 75 | 0.058 | 0.058 | 0.059 | 0.055 | 0.053 |
| 76 | 0.058 | 0.059 | 0.059 | 0.056 | 0.053 |
| 77 | 0.059 | 0.059 | 0.059 | 0.057 | 0.053 |
| 78 | 0.059 | 0.060 | 0.059 | 0.058 | 0.053 |
| 79 | 0.059 | 0.060 | 0.059 | 0.058 | 0.053 |
| 80 | 0.060 | 0.061 | 0.060 | 0.059 | 0.055 |
| 81 | 0.060 | 0.062 | 0.060 | 0.059 | 0.056 |
| 82 | 0.061 | 0.064 | 0.061 | 0.059 | 0.058 |
| 83 | 0.061 | 0.066 | 0.062 | 0.060 | 0.060 |
| 84 | 0.061 | 0.068 | 0.064 | 0.060 | 0.061 |
| 85 | 0.062 | 0.069 | 0.065 | 0.060 | 0.061 |
| 86 | 0.064 | 0.070 | 0.068 | 0.060 | 0.061 |
| 87 | 0.065 | 0.071 | 0.071 | 0.061 | 0.061 |
| 88 | 0.067 | 0.072 | 0.074 | 0.061 | 0.061 |
| 89 | 0.068 | 0.074 | 0.076 | 0.061 | 0.062 |
| 90 | 0.068 | 0.076 | 0.076 | 0.062 | 0.063 |
| 91 | 0.071 | 0.078 | 0.076 | 0.062 | 0.064 |
| 92 | 0.075 | 0.080 | 0.077 | 0.063 | 0.065 |
| 93 | 0.077 | 0.082 | 0.079 | 0.064 | 0.066 |
| 94 | 0.079 | 0.084 | 0.086 | 0.066 | 0.067 |
| 95 | 0.082 | 0.087 | 0.093 | 0.067 | 0.068 |
| 96 | 0.088 | 0.089 | 0.100 | 0.070 | 0.069 |
| 97 | 0.101 | 0.098 | 0.108 | 0.082 | 0.072 |
| 98 | 0.116 | 0.106 | 0.117 | 0.094 | 0.076 |
| 99 | 0.122 | 0.114 | 0.127 | 0.105 | 0.079 |
| 100 | 0.136 | 0.123 | 0.136 | 0.117 | 0.082 |

***Table S32:*** *Unpaced tapping – Variability (CV of ITI) of fast tapping – Unpaced_fast_CV_iti*

| | | | | | |
|---|---|---|---|---|---|
| **Task** | Unpaced tapping | | | | |
| **Outcome measure** | Variability (CV of ITI) of fast tapping | | | | |
| **Variable name** | Unpaced_fast_CV_iti | | | | |
| **Age and gender effects** | | | | | |
| Age regression (slope) | 0.000 | | | | |
| Age regression (p) | 0.042 | | | | |
| Age group (p) | 0.059 | | | | |
| Gender (p) | 0.090 | | | | |
| **Group** | All | Age 18 to 21 | Age 22 to 29 | Age 30 to 54 | Age 55 to 87 |
| N (N female) | 107 (73) | 27 (18) | 29 (19) | 26 (16) | 25 (20) |
| **Normality** | | | | | |
| Skewness | 2.67 | 2.56 | 1.87 | 2.46 | 1.37 |
| Excess kurtosis | 8.64 | 5.98 | 3.90 | 6.15 | 1.80 |
| **Scores** | | | | | |
| Mean | 0.061 | 0.067 | 0.065 | 0.056 | 0.053 |
| SD | 0.029 | 0.039 | 0.025 | 0.028 | 0.019 |
| Percentiles | | | | | |
| 0 | 0.027 | 0.036 | 0.036 | 0.032 | 0.027 |
| 1 | 0.031 | 0.036 | 0.037 | 0.032 | 0.028 |
| 2 | 0.032 | 0.037 | 0.038 | 0.033 | 0.029 |
| 3 | 0.033 | 0.038 | 0.039 | 0.033 | 0.030 |
| 4 | 0.034 | 0.039 | 0.040 | 0.033 | 0.031 |
| 5 | 0.036 | 0.039 | 0.041 | 0.033 | 0.033 |
| 6 | 0.036 | 0.039 | 0.041 | 0.033 | 0.034 |
| 7 | 0.037 | 0.039 | 0.042 | 0.033 | 0.035 |
| 8 | 0.037 | 0.040 | 0.043 | 0.033 | 0.037 |
| 9 | 0.038 | 0.040 | 0.043 | 0.034 | 0.038 |
| 10 | 0.039 | 0.041 | 0.044 | 0.035 | 0.038 |
| 11 | 0.039 | 0.041 | 0.044 | 0.036 | 0.039 |
| 12 | 0.040 | 0.042 | 0.044 | 0.037 | 0.040 |
| 13 | 0.040 | 0.042 | 0.045 | 0.037 | 0.040 |
| 14 | 0.040 | 0.042 | 0.045 | 0.037 | 0.041 |
| 15 | 0.041 | 0.042 | 0.045 | 0.037 | 0.041 |
| 16 | 0.042 | 0.043 | 0.045 | 0.037 | 0.041 |
| 17 | 0.042 | 0.043 | 0.046 | 0.038 | 0.041 |
| 18 | 0.042 | 0.044 | 0.046 | 0.038 | 0.042 |
| 19 | 0.042 | 0.045 | 0.046 | 0.039 | 0.042 |
| 20 | 0.042 | 0.045 | 0.047 | 0.039 | 0.042 |
| 21 | 0.042 | 0.045 | 0.047 | 0.039 | 0.042 |
| 22 | 0.043 | 0.045 | 0.048 | 0.039 | 0.042 |
| 23 | 0.044 | 0.046 | 0.048 | 0.039 | 0.042 |
| 24 | 0.044 | 0.046 | 0.048 | 0.040 | 0.042 |

| | | | | | |
|---|---|---|---|---|---|
| 25 | 0.044 | 0.046 | 0.048 | 0.041 | 0.042 |
| 26 | 0.044 | 0.046 | 0.049 | 0.042 | 0.042 |
| 27 | 0.045 | 0.047 | 0.050 | 0.043 | 0.042 |
| 28 | 0.045 | 0.047 | 0.050 | 0.044 | 0.042 |
| 29 | 0.045 | 0.047 | 0.051 | 0.044 | 0.042 |
| 30 | 0.046 | 0.047 | 0.052 | 0.045 | 0.042 |
| 31 | 0.046 | 0.047 | 0.052 | 0.045 | 0.043 |
| 32 | 0.046 | 0.048 | 0.053 | 0.046 | 0.043 |
| 33 | 0.046 | 0.048 | 0.053 | 0.046 | 0.043 |
| 34 | 0.047 | 0.049 | 0.053 | 0.046 | 0.043 |
| 35 | 0.047 | 0.049 | 0.053 | 0.046 | 0.043 |
| 36 | 0.047 | 0.050 | 0.054 | 0.046 | 0.044 |
| 37 | 0.048 | 0.051 | 0.055 | 0.047 | 0.044 |
| 38 | 0.048 | 0.051 | 0.057 | 0.047 | 0.044 |
| 39 | 0.048 | 0.052 | 0.058 | 0.047 | 0.044 |
| 40 | 0.048 | 0.052 | 0.059 | 0.048 | 0.044 |
| 41 | 0.049 | 0.052 | 0.059 | 0.048 | 0.044 |
| 42 | 0.049 | 0.052 | 0.059 | 0.048 | 0.044 |
| 43 | 0.049 | 0.053 | 0.059 | 0.048 | 0.044 |
| 44 | 0.049 | 0.053 | 0.059 | 0.048 | 0.044 |
| 45 | 0.050 | 0.054 | 0.060 | 0.048 | 0.045 |
| 46 | 0.050 | 0.055 | 0.060 | 0.048 | 0.045 |
| 47 | 0.051 | 0.055 | 0.060 | 0.048 | 0.045 |
| 48 | 0.051 | 0.056 | 0.060 | 0.048 | 0.046 |
| 49 | 0.051 | 0.057 | 0.060 | 0.049 | 0.046 |
| 50 | 0.052 | 0.058 | 0.060 | 0.049 | 0.047 |
| 51 | 0.052 | 0.059 | 0.061 | 0.049 | 0.047 |
| 52 | 0.053 | 0.059 | 0.061 | 0.049 | 0.047 |
| 53 | 0.053 | 0.059 | 0.062 | 0.049 | 0.047 |
| 54 | 0.054 | 0.060 | 0.062 | 0.049 | 0.047 |
| 55 | 0.055 | 0.060 | 0.062 | 0.049 | 0.047 |
| 56 | 0.056 | 0.060 | 0.062 | 0.049 | 0.048 |
| 57 | 0.058 | 0.060 | 0.062 | 0.049 | 0.048 |
| 58 | 0.058 | 0.060 | 0.062 | 0.050 | 0.049 |
| 59 | 0.059 | 0.060 | 0.062 | 0.050 | 0.049 |
| 60 | 0.060 | 0.061 | 0.063 | 0.050 | 0.050 |
| 61 | 0.060 | 0.061 | 0.063 | 0.050 | 0.050 |
| 62 | 0.060 | 0.061 | 0.063 | 0.050 | 0.051 |
| 63 | 0.060 | 0.062 | 0.063 | 0.050 | 0.051 |
| 64 | 0.061 | 0.062 | 0.063 | 0.051 | 0.052 |
| 65 | 0.061 | 0.062 | 0.063 | 0.051 | 0.053 |
| 66 | 0.062 | 0.063 | 0.064 | 0.051 | 0.053 |
| 67 | 0.062 | 0.063 | 0.064 | 0.052 | 0.054 |
| 68 | 0.062 | 0.063 | 0.065 | 0.052 | 0.056 |

| | | | | | |
|---|---|---|---|---|---|
| 69 | 0.062 | 0.063 | 0.066 | 0.053 | 0.058 |
| 70 | 0.063 | 0.064 | 0.066 | 0.054 | 0.060 |
| 71 | 0.063 | 0.064 | 0.067 | 0.055 | 0.062 |
| 72 | 0.063 | 0.065 | 0.067 | 0.056 | 0.062 |
| 73 | 0.064 | 0.065 | 0.067 | 0.056 | 0.062 |
| 74 | 0.065 | 0.066 | 0.067 | 0.057 | 0.062 |
| 75 | 0.065 | 0.068 | 0.067 | 0.057 | 0.062 |
| 76 | 0.066 | 0.069 | 0.068 | 0.057 | 0.063 |
| 77 | 0.066 | 0.070 | 0.069 | 0.058 | 0.064 |
| 78 | 0.067 | 0.070 | 0.070 | 0.059 | 0.064 |
| 79 | 0.069 | 0.070 | 0.071 | 0.060 | 0.065 |
| 80 | 0.070 | 0.070 | 0.073 | 0.061 | 0.066 |
| 81 | 0.070 | 0.070 | 0.074 | 0.062 | 0.067 |
| 82 | 0.070 | 0.071 | 0.076 | 0.064 | 0.068 |
| 83 | 0.073 | 0.072 | 0.077 | 0.065 | 0.069 |
| 84 | 0.073 | 0.073 | 0.078 | 0.066 | 0.070 |
| 85 | 0.074 | 0.074 | 0.079 | 0.068 | 0.071 |
| 86 | 0.076 | 0.076 | 0.080 | 0.069 | 0.072 |
| 87 | 0.077 | 0.078 | 0.085 | 0.071 | 0.072 |
| 88 | 0.079 | 0.079 | 0.089 | 0.073 | 0.073 |
| 89 | 0.083 | 0.084 | 0.093 | 0.078 | 0.074 |
| 90 | 0.090 | 0.093 | 0.096 | 0.083 | 0.075 |
| 91 | 0.094 | 0.101 | 0.099 | 0.089 | 0.076 |
| 92 | 0.100 | 0.109 | 0.102 | 0.094 | 0.077 |
| 93 | 0.108 | 0.123 | 0.105 | 0.099 | 0.080 |
| 94 | 0.111 | 0.139 | 0.107 | 0.103 | 0.083 |
| 95 | 0.111 | 0.155 | 0.108 | 0.107 | 0.085 |
| 96 | 0.112 | 0.170 | 0.110 | 0.111 | 0.089 |
| 97 | 0.145 | 0.181 | 0.117 | 0.124 | 0.094 |
| 98 | 0.161 | 0.190 | 0.129 | 0.137 | 0.100 |
| 99 | 0.172 | 0.200 | 0.141 | 0.149 | 0.106 |
| 100 | 0.209 | 0.209 | 0.152 | 0.162 | 0.111 |

***Table S33:*** *Unpaced tapping – Variability (CV of ITI) of slow tapping – Unpaced_slow_CV_iti*

| | | | | | |
|---|---|---|---|---|---|
| **Task** | Unpaced tapping | | | | |
| **Outcome measure** | Variability (CV of ITI) of slow tapping | | | | |
| **Variable name** | Unpaced_slow_CV_iti | | | | |
| **Age and gender effects** | | | | | |
| Age regression (slope) | 0.000 | | | | |
| Age regression (p) | 0.584 | | | | |
| Age group (p) | 0.377 | | | | |
| Gender (p) | 0.410 | | | | |
| **Group** | All | Age 18 to 21 | Age 22 to 29 | Age 30 to 54 | Age 55 to 87 |
| N (N female) | 103 (71) | 25 (16) | 28 (19) | 25 (16) | 25 (20) |
| **Normality** | | | | | |
| Skewness | 7.46 | 4.23 | 3.06 | 1.66 | 0.71 |
| Excess kurtosis | 64.15 | 17.18 | 10.60 | 3.37 | 0.03 |
| **Scores** | | | | | |
| Mean | 0.064 | 0.085 | 0.056 | 0.055 | 0.060 |
| SD | 0.061 | 0.115 | 0.031 | 0.026 | 0.019 |
| Percentiles | | | | | |
| 0 | 0.022 | 0.022 | 0.029 | 0.023 | 0.032 |
| 1 | 0.023 | 0.025 | 0.029 | 0.023 | 0.033 |
| 2 | 0.024 | 0.028 | 0.030 | 0.023 | 0.034 |
| 3 | 0.028 | 0.031 | 0.030 | 0.023 | 0.035 |
| 4 | 0.029 | 0.034 | 0.030 | 0.024 | 0.037 |
| 5 | 0.030 | 0.035 | 0.030 | 0.024 | 0.037 |
| 6 | 0.030 | 0.035 | 0.030 | 0.025 | 0.037 |
| 7 | 0.032 | 0.035 | 0.030 | 0.026 | 0.037 |
| 8 | 0.032 | 0.036 | 0.032 | 0.027 | 0.038 |
| 9 | 0.034 | 0.037 | 0.034 | 0.028 | 0.038 |
| 10 | 0.035 | 0.037 | 0.036 | 0.029 | 0.038 |
| 11 | 0.035 | 0.038 | 0.038 | 0.030 | 0.038 |
| 12 | 0.036 | 0.039 | 0.039 | 0.031 | 0.038 |
| 13 | 0.037 | 0.040 | 0.040 | 0.032 | 0.039 |
| 14 | 0.037 | 0.041 | 0.040 | 0.033 | 0.040 |
| 15 | 0.038 | 0.042 | 0.041 | 0.033 | 0.041 |
| 16 | 0.038 | 0.043 | 0.041 | 0.034 | 0.042 |
| 17 | 0.039 | 0.044 | 0.041 | 0.034 | 0.043 |
| 18 | 0.040 | 0.044 | 0.041 | 0.035 | 0.044 |
| 19 | 0.041 | 0.044 | 0.041 | 0.035 | 0.044 |
| 20 | 0.041 | 0.044 | 0.041 | 0.035 | 0.045 |
| 21 | 0.041 | 0.044 | 0.041 | 0.035 | 0.045 |
| 22 | 0.041 | 0.045 | 0.041 | 0.036 | 0.046 |
| 23 | 0.042 | 0.045 | 0.041 | 0.036 | 0.046 |
| 24 | 0.042 | 0.045 | 0.041 | 0.036 | 0.047 |

| | | | | | |
|---|---|---|---|---|---|
| 25 | 0.043 | 0.045 | 0.041 | 0.037 | 0.047 |
| 26 | 0.044 | 0.046 | 0.041 | 0.038 | 0.047 |
| 27 | 0.044 | 0.046 | 0.041 | 0.039 | 0.047 |
| 28 | 0.044 | 0.046 | 0.042 | 0.041 | 0.048 |
| 29 | 0.045 | 0.046 | 0.042 | 0.042 | 0.048 |
| 30 | 0.045 | 0.046 | 0.042 | 0.043 | 0.048 |
| 31 | 0.045 | 0.046 | 0.042 | 0.043 | 0.049 |
| 32 | 0.045 | 0.046 | 0.043 | 0.044 | 0.049 |
| 33 | 0.046 | 0.046 | 0.043 | 0.044 | 0.050 |
| 34 | 0.046 | 0.047 | 0.044 | 0.045 | 0.050 |
| 35 | 0.046 | 0.048 | 0.044 | 0.047 | 0.051 |
| 36 | 0.047 | 0.049 | 0.045 | 0.048 | 0.051 |
| 37 | 0.048 | 0.050 | 0.045 | 0.050 | 0.051 |
| 38 | 0.048 | 0.051 | 0.045 | 0.050 | 0.051 |
| 39 | 0.048 | 0.051 | 0.045 | 0.051 | 0.051 |
| 40 | 0.048 | 0.051 | 0.045 | 0.051 | 0.051 |
| 41 | 0.050 | 0.051 | 0.045 | 0.051 | 0.051 |
| 42 | 0.050 | 0.051 | 0.045 | 0.051 | 0.051 |
| 43 | 0.051 | 0.051 | 0.046 | 0.052 | 0.051 |
| 44 | 0.051 | 0.052 | 0.046 | 0.052 | 0.051 |
| 45 | 0.051 | 0.052 | 0.046 | 0.052 | 0.052 |
| 46 | 0.051 | 0.052 | 0.047 | 0.053 | 0.052 |
| 47 | 0.051 | 0.053 | 0.047 | 0.053 | 0.053 |
| 48 | 0.052 | 0.053 | 0.048 | 0.053 | 0.054 |
| 49 | 0.052 | 0.053 | 0.048 | 0.053 | 0.056 |
| 50 | 0.052 | 0.053 | 0.048 | 0.053 | 0.057 |
| 51 | 0.052 | 0.054 | 0.048 | 0.053 | 0.057 |
| 52 | 0.053 | 0.054 | 0.048 | 0.053 | 0.058 |
| 53 | 0.053 | 0.054 | 0.048 | 0.053 | 0.058 |
| 54 | 0.053 | 0.055 | 0.048 | 0.053 | 0.058 |
| 55 | 0.053 | 0.055 | 0.048 | 0.053 | 0.059 |
| 56 | 0.053 | 0.055 | 0.048 | 0.053 | 0.060 |
| 57 | 0.053 | 0.055 | 0.049 | 0.054 | 0.061 |
| 58 | 0.054 | 0.056 | 0.050 | 0.054 | 0.062 |
| 59 | 0.054 | 0.057 | 0.051 | 0.054 | 0.063 |
| 60 | 0.054 | 0.059 | 0.052 | 0.054 | 0.063 |
| 61 | 0.055 | 0.062 | 0.052 | 0.054 | 0.063 |
| 62 | 0.056 | 0.064 | 0.052 | 0.054 | 0.064 |
| 63 | 0.056 | 0.066 | 0.052 | 0.054 | 0.064 |
| 64 | 0.057 | 0.066 | 0.052 | 0.055 | 0.064 |
| 65 | 0.059 | 0.067 | 0.053 | 0.055 | 0.065 |
| 66 | 0.060 | 0.068 | 0.053 | 0.055 | 0.065 |
| 67 | 0.063 | 0.069 | 0.053 | 0.056 | 0.066 |
| 68 | 0.064 | 0.070 | 0.053 | 0.057 | 0.067 |

| | | | | | |
|---|---|---|---|---|---|
| 69 | 0.064 | 0.072 | 0.053 | 0.058 | 0.068 |
| 70 | 0.065 | 0.073 | 0.053 | 0.059 | 0.069 |
| 71 | 0.065 | 0.075 | 0.053 | 0.060 | 0.070 |
| 72 | 0.066 | 0.075 | 0.054 | 0.061 | 0.070 |
| 73 | 0.066 | 0.075 | 0.054 | 0.062 | 0.070 |
| 74 | 0.067 | 0.076 | 0.054 | 0.064 | 0.070 |
| 75 | 0.068 | 0.076 | 0.056 | 0.065 | 0.070 |
| 76 | 0.069 | 0.076 | 0.059 | 0.065 | 0.071 |
| 77 | 0.070 | 0.076 | 0.062 | 0.065 | 0.073 |
| 78 | 0.070 | 0.076 | 0.064 | 0.066 | 0.074 |
| 79 | 0.071 | 0.076 | 0.065 | 0.066 | 0.076 |
| 80 | 0.073 | 0.077 | 0.066 | 0.067 | 0.077 |
| 81 | 0.074 | 0.078 | 0.066 | 0.067 | 0.077 |
| 82 | 0.075 | 0.079 | 0.068 | 0.068 | 0.078 |
| 83 | 0.076 | 0.080 | 0.070 | 0.069 | 0.079 |
| 84 | 0.076 | 0.087 | 0.072 | 0.069 | 0.079 |
| 85 | 0.078 | 0.097 | 0.074 | 0.069 | 0.079 |
| 86 | 0.079 | 0.107 | 0.075 | 0.069 | 0.079 |
| 87 | 0.079 | 0.116 | 0.076 | 0.070 | 0.079 |
| 88 | 0.080 | 0.122 | 0.078 | 0.070 | 0.080 |
| 89 | 0.085 | 0.124 | 0.079 | 0.071 | 0.082 |
| 90 | 0.089 | 0.126 | 0.082 | 0.071 | 0.083 |
| 91 | 0.090 | 0.128 | 0.086 | 0.072 | 0.085 |
| 92 | 0.092 | 0.130 | 0.089 | 0.075 | 0.086 |
| 93 | 0.107 | 0.133 | 0.091 | 0.084 | 0.087 |
| 94 | 0.110 | 0.136 | 0.091 | 0.093 | 0.088 |
| 95 | 0.120 | 0.138 | 0.092 | 0.102 | 0.089 |
| 96 | 0.129 | 0.160 | 0.092 | 0.111 | 0.090 |
| 97 | 0.139 | 0.275 | 0.111 | 0.118 | 0.095 |
| 98 | 0.140 | 0.390 | 0.137 | 0.126 | 0.100 |
| 99 | 0.188 | 0.505 | 0.163 | 0.133 | 0.105 |
| 100 | 0.620 | 0.620 | 0.189 | 0.140 | 0.109 |

***Table S34:*** *Paced tapping with tones – Variability (CV of ITI), fast tempo – Paced_metro_450_mean_CV_iti*

| | | | | | | |
|---|---|---|---|---|---|---|
| **Task** | Paced tapping with tones | | | | | |
| **Outcome measure** | Variability (CV of ITI), fast tempo | | | | | |
| **Variable name** | Paced_metro_450_mean_CV_iti | | | | | |
| **Age and gender effects** | | | | | | |
| Age regression (slope) | | 0.000 | | | | |
| Age regression (p) | | 0.160 | | | | |
| Age group (p) | | 0.326 | | | | |
| Gender (p) | | 0.534 | | | | |
| **Group** | | All | Age 18 to 21 | Age 22 to 29 | Age 30 to 54 | Age 55 to 87 |
| N (N female) | | 107 (73) | 27 (18) | 29 (19) | 26 (16) | 25 (20) |
| **Normality** | | | | | | |
| Skewness | | 1.00 | 1.63 | 0.93 | 0.43 | 0.78 |
| Excess kurtosis | | 1.41 | 3.31 | 0.57 | -0.88 | 0.88 |
| **Scores** | | | | | | |
| Mean | | 0.053 | 0.056 | 0.053 | 0.052 | 0.049 |
| SD | | 0.015 | 0.015 | 0.014 | 0.015 | 0.013 |
| Percentiles | | | | | | |
| | 0 | 0.029 | 0.038 | 0.031 | 0.029 | 0.032 |
| | 1 | 0.031 | 0.038 | 0.032 | 0.030 | 0.032 |
| | 2 | 0.032 | 0.038 | 0.034 | 0.031 | 0.032 |
| | 3 | 0.033 | 0.038 | 0.035 | 0.032 | 0.032 |
| | 4 | 0.033 | 0.039 | 0.036 | 0.034 | 0.033 |
| | 5 | 0.034 | 0.039 | 0.036 | 0.034 | 0.033 |
| | 6 | 0.034 | 0.040 | 0.037 | 0.034 | 0.033 |
| | 7 | 0.035 | 0.040 | 0.037 | 0.035 | 0.033 |
| | 8 | 0.035 | 0.041 | 0.037 | 0.035 | 0.033 |
| | 9 | 0.036 | 0.041 | 0.037 | 0.035 | 0.033 |
| | 10 | 0.036 | 0.041 | 0.037 | 0.035 | 0.033 |
| | 11 | 0.037 | 0.041 | 0.037 | 0.035 | 0.033 |
| | 12 | 0.037 | 0.041 | 0.038 | 0.035 | 0.033 |
| | 13 | 0.037 | 0.042 | 0.039 | 0.036 | 0.034 |
| | 14 | 0.038 | 0.042 | 0.039 | 0.036 | 0.034 |
| | 15 | 0.038 | 0.043 | 0.040 | 0.037 | 0.035 |
| | 16 | 0.038 | 0.043 | 0.040 | 0.038 | 0.036 |
| | 17 | 0.038 | 0.043 | 0.041 | 0.038 | 0.036 |
| | 18 | 0.039 | 0.043 | 0.041 | 0.038 | 0.036 |
| | 19 | 0.039 | 0.044 | 0.041 | 0.038 | 0.036 |
| | 20 | 0.039 | 0.044 | 0.042 | 0.038 | 0.036 |
| | 21 | 0.040 | 0.044 | 0.042 | 0.038 | 0.036 |
| | 22 | 0.040 | 0.044 | 0.043 | 0.038 | 0.036 |

| | | | | | |
|---|---|---|---|---|---|
| 23 | 0.041 | 0.045 | 0.044 | 0.038 | 0.037 |
| 24 | 0.041 | 0.045 | 0.045 | 0.038 | 0.037 |
| 25 | 0.042 | 0.046 | 0.046 | 0.038 | 0.037 |
| 26 | 0.042 | 0.046 | 0.046 | 0.038 | 0.037 |
| 27 | 0.042 | 0.047 | 0.047 | 0.038 | 0.038 |
| 28 | 0.043 | 0.047 | 0.047 | 0.039 | 0.038 |
| 29 | 0.044 | 0.047 | 0.047 | 0.039 | 0.039 |
| 30 | 0.044 | 0.047 | 0.047 | 0.039 | 0.040 |
| 31 | 0.045 | 0.047 | 0.047 | 0.040 | 0.040 |
| 32 | 0.045 | 0.048 | 0.047 | 0.040 | 0.041 |
| 33 | 0.046 | 0.048 | 0.047 | 0.041 | 0.042 |
| 34 | 0.046 | 0.048 | 0.047 | 0.042 | 0.042 |
| 35 | 0.046 | 0.048 | 0.047 | 0.043 | 0.043 |
| 36 | 0.047 | 0.049 | 0.048 | 0.044 | 0.043 |
| 37 | 0.047 | 0.049 | 0.048 | 0.044 | 0.043 |
| 38 | 0.047 | 0.049 | 0.048 | 0.044 | 0.044 |
| 39 | 0.047 | 0.050 | 0.048 | 0.045 | 0.044 |
| 40 | 0.048 | 0.050 | 0.048 | 0.045 | 0.045 |
| 41 | 0.048 | 0.050 | 0.048 | 0.045 | 0.045 |
| 42 | 0.048 | 0.050 | 0.048 | 0.046 | 0.046 |
| 43 | 0.048 | 0.051 | 0.048 | 0.046 | 0.046 |
| 44 | 0.048 | 0.051 | 0.049 | 0.046 | 0.047 |
| 45 | 0.049 | 0.051 | 0.049 | 0.047 | 0.048 |
| 46 | 0.049 | 0.051 | 0.050 | 0.047 | 0.048 |
| 47 | 0.049 | 0.052 | 0.050 | 0.048 | 0.048 |
| 48 | 0.050 | 0.053 | 0.051 | 0.049 | 0.048 |
| 49 | 0.051 | 0.053 | 0.052 | 0.049 | 0.049 |
| 50 | 0.051 | 0.054 | 0.052 | 0.049 | 0.049 |
| 51 | 0.051 | 0.055 | 0.052 | 0.049 | 0.049 |
| 52 | 0.051 | 0.055 | 0.052 | 0.049 | 0.050 |
| 53 | 0.051 | 0.056 | 0.052 | 0.049 | 0.050 |
| 54 | 0.052 | 0.056 | 0.053 | 0.050 | 0.051 |
| 55 | 0.052 | 0.057 | 0.053 | 0.050 | 0.051 |
| 56 | 0.052 | 0.057 | 0.054 | 0.051 | 0.051 |
| 57 | 0.053 | 0.058 | 0.054 | 0.051 | 0.051 |
| 58 | 0.054 | 0.059 | 0.054 | 0.051 | 0.051 |
| 59 | 0.054 | 0.059 | 0.054 | 0.051 | 0.051 |
| 60 | 0.054 | 0.059 | 0.054 | 0.052 | 0.052 |
| 61 | 0.054 | 0.059 | 0.054 | 0.053 | 0.052 |
| 62 | 0.055 | 0.059 | 0.054 | 0.055 | 0.052 |
| 63 | 0.056 | 0.059 | 0.054 | 0.056 | 0.052 |
| 64 | 0.056 | 0.059 | 0.054 | 0.058 | 0.053 |
| 65 | 0.057 | 0.060 | 0.055 | 0.059 | 0.053 |
| 66 | 0.057 | 0.060 | 0.055 | 0.060 | 0.054 |

| | | | | | |
|---|---|---|---|---|---|
| 67 | 0.057 | 0.060 | 0.055 | 0.061 | 0.054 |
| 68 | 0.058 | 0.060 | 0.055 | 0.062 | 0.055 |
| 69 | 0.058 | 0.060 | 0.055 | 0.063 | 0.055 |
| 70 | 0.059 | 0.060 | 0.056 | 0.064 | 0.056 |
| 71 | 0.059 | 0.060 | 0.056 | 0.064 | 0.057 |
| 72 | 0.060 | 0.060 | 0.057 | 0.065 | 0.057 |
| 73 | 0.060 | 0.061 | 0.058 | 0.065 | 0.057 |
| 74 | 0.060 | 0.061 | 0.059 | 0.065 | 0.057 |
| 75 | 0.060 | 0.061 | 0.060 | 0.065 | 0.057 |
| 76 | 0.061 | 0.061 | 0.060 | 0.065 | 0.057 |
| 77 | 0.061 | 0.061 | 0.060 | 0.066 | 0.057 |
| 78 | 0.061 | 0.062 | 0.060 | 0.066 | 0.057 |
| 79 | 0.061 | 0.062 | 0.061 | 0.066 | 0.057 |
| 80 | 0.062 | 0.062 | 0.061 | 0.067 | 0.057 |
| 81 | 0.062 | 0.062 | 0.061 | 0.067 | 0.058 |
| 82 | 0.064 | 0.063 | 0.061 | 0.068 | 0.058 |
| 83 | 0.065 | 0.064 | 0.062 | 0.069 | 0.058 |
| 84 | 0.065 | 0.065 | 0.063 | 0.070 | 0.058 |
| 85 | 0.066 | 0.066 | 0.065 | 0.070 | 0.059 |
| 86 | 0.066 | 0.066 | 0.066 | 0.070 | 0.059 |
| 87 | 0.067 | 0.066 | 0.068 | 0.070 | 0.059 |
| 88 | 0.068 | 0.067 | 0.070 | 0.070 | 0.060 |
| 89 | 0.070 | 0.068 | 0.072 | 0.070 | 0.060 |
| 90 | 0.070 | 0.070 | 0.073 | 0.071 | 0.060 |
| 91 | 0.071 | 0.072 | 0.075 | 0.071 | 0.060 |
| 92 | 0.072 | 0.073 | 0.077 | 0.071 | 0.061 |
| 93 | 0.073 | 0.076 | 0.079 | 0.071 | 0.062 |
| 94 | 0.077 | 0.079 | 0.081 | 0.071 | 0.062 |
| 95 | 0.083 | 0.082 | 0.083 | 0.072 | 0.063 |
| 96 | 0.085 | 0.085 | 0.084 | 0.072 | 0.065 |
| 97 | 0.086 | 0.091 | 0.086 | 0.076 | 0.070 |
| 98 | 0.086 | 0.097 | 0.088 | 0.079 | 0.075 |
| 99 | 0.091 | 0.103 | 0.089 | 0.083 | 0.080 |
| 100 | 0.109 | 0.109 | 0.091 | 0.086 | 0.085 |

***Table S35:*** *Paced tapping with tones – Variability (CV of ITI), medium tempo – Paced_metro_600_mean_CV_iti*

| | | | | | | |
|---|---|---|---|---|---|---|
| **Task** | Paced tapping with tones | | | | | |
| **Outcome measure** | Variability (CV of ITI), medium tempo | | | | | |
| **Variable name** | Paced_metro_600_mean_CV_iti | | | | | |
| **Age and gender effects** | | | | | | |
| Age regression (slope) | | 0.000 | | | | |
| Age regression (p) | | 0.223 | | | | |
| Age group (p) | | 0.475 | | | | |
| Gender (p) | | 0.990 | | | | |
| **Group** | | All | Age 18 to 21 | Age 22 to 29 | Age 30 to 54 | Age 55 to 87 |
| N (N female) | | 108 (74) | 27 (18) | 29 (19) | 26 (16) | 26 (21) |
| **Normality** | | | | | | |
| Skewness | | 0.62 | 0.90 | 0.62 | 0.63 | 0.51 |
| Excess kurtosis | | -0.01 | 1.14 | 0.14 | -0.64 | -0.85 |
| **Scores** | | | | | | |
| Mean | | 0.046 | 0.048 | 0.046 | 0.047 | 0.044 |
| SD | | 0.011 | 0.009 | 0.012 | 0.012 | 0.010 |
| Percentiles | | | | | | |
| | 0 | 0.024 | 0.036 | 0.024 | 0.029 | 0.030 |
| | 1 | 0.029 | 0.037 | 0.026 | 0.030 | 0.030 |
| | 2 | 0.030 | 0.037 | 0.027 | 0.030 | 0.031 |
| | 3 | 0.030 | 0.037 | 0.029 | 0.031 | 0.031 |
| | 4 | 0.032 | 0.037 | 0.030 | 0.031 | 0.032 |
| | 5 | 0.032 | 0.037 | 0.031 | 0.032 | 0.032 |
| | 6 | 0.032 | 0.037 | 0.031 | 0.032 | 0.032 |
| | 7 | 0.032 | 0.037 | 0.032 | 0.032 | 0.032 |
| | 8 | 0.033 | 0.037 | 0.032 | 0.033 | 0.032 |
| | 9 | 0.033 | 0.037 | 0.033 | 0.034 | 0.032 |
| | 10 | 0.034 | 0.037 | 0.033 | 0.034 | 0.033 |
| | 11 | 0.034 | 0.037 | 0.034 | 0.035 | 0.033 |
| | 12 | 0.035 | 0.037 | 0.034 | 0.036 | 0.033 |
| | 13 | 0.035 | 0.038 | 0.034 | 0.036 | 0.033 |
| | 14 | 0.036 | 0.038 | 0.034 | 0.036 | 0.033 |
| | 15 | 0.036 | 0.038 | 0.035 | 0.036 | 0.033 |
| | 16 | 0.036 | 0.039 | 0.035 | 0.036 | 0.034 |
| | 17 | 0.036 | 0.039 | 0.035 | 0.037 | 0.034 |
| | 18 | 0.036 | 0.040 | 0.036 | 0.037 | 0.034 |
| | 19 | 0.037 | 0.040 | 0.036 | 0.038 | 0.035 |
| | 20 | 0.037 | 0.040 | 0.036 | 0.038 | 0.035 |
| | 21 | 0.037 | 0.040 | 0.036 | 0.038 | 0.035 |
| | 22 | 0.037 | 0.041 | 0.037 | 0.038 | 0.035 |

| | | | | | |
|---|---|---|---|---|---|
| 23 | 0.037 | 0.041 | 0.037 | 0.039 | 0.036 |
| 24 | 0.037 | 0.041 | 0.037 | 0.039 | 0.036 |
| 25 | 0.038 | 0.041 | 0.037 | 0.039 | 0.036 |
| 26 | 0.038 | 0.041 | 0.037 | 0.039 | 0.036 |
| 27 | 0.038 | 0.041 | 0.037 | 0.039 | 0.036 |
| 28 | 0.039 | 0.041 | 0.037 | 0.039 | 0.036 |
| 29 | 0.039 | 0.042 | 0.037 | 0.039 | 0.036 |
| 30 | 0.039 | 0.042 | 0.038 | 0.040 | 0.037 |
| 31 | 0.039 | 0.042 | 0.038 | 0.041 | 0.037 |
| 32 | 0.039 | 0.042 | 0.038 | 0.041 | 0.038 |
| 33 | 0.039 | 0.043 | 0.039 | 0.041 | 0.038 |
| 34 | 0.040 | 0.043 | 0.039 | 0.041 | 0.038 |
| 35 | 0.041 | 0.043 | 0.039 | 0.041 | 0.038 |
| 36 | 0.041 | 0.043 | 0.039 | 0.041 | 0.038 |
| 37 | 0.041 | 0.044 | 0.039 | 0.041 | 0.038 |
| 38 | 0.041 | 0.044 | 0.039 | 0.042 | 0.038 |
| 39 | 0.041 | 0.044 | 0.039 | 0.042 | 0.038 |
| 40 | 0.042 | 0.045 | 0.039 | 0.042 | 0.038 |
| 41 | 0.042 | 0.046 | 0.040 | 0.042 | 0.039 |
| 42 | 0.042 | 0.046 | 0.041 | 0.042 | 0.040 |
| 43 | 0.042 | 0.046 | 0.042 | 0.043 | 0.040 |
| 44 | 0.043 | 0.047 | 0.042 | 0.043 | 0.041 |
| 45 | 0.043 | 0.047 | 0.043 | 0.043 | 0.041 |
| 46 | 0.043 | 0.047 | 0.043 | 0.043 | 0.041 |
| 47 | 0.043 | 0.047 | 0.044 | 0.043 | 0.042 |
| 48 | 0.043 | 0.047 | 0.044 | 0.043 | 0.042 |
| 49 | 0.044 | 0.047 | 0.044 | 0.043 | 0.042 |
| 50 | 0.044 | 0.047 | 0.045 | 0.043 | 0.042 |
| 51 | 0.044 | 0.047 | 0.045 | 0.043 | 0.042 |
| 52 | 0.045 | 0.048 | 0.045 | 0.043 | 0.042 |
| 53 | 0.045 | 0.048 | 0.045 | 0.044 | 0.042 |
| 54 | 0.046 | 0.048 | 0.046 | 0.044 | 0.042 |
| 55 | 0.046 | 0.049 | 0.046 | 0.044 | 0.043 |
| 56 | 0.046 | 0.050 | 0.046 | 0.044 | 0.043 |
| 57 | 0.047 | 0.050 | 0.046 | 0.045 | 0.043 |
| 58 | 0.047 | 0.051 | 0.046 | 0.046 | 0.043 |
| 59 | 0.047 | 0.051 | 0.046 | 0.046 | 0.044 |
| 60 | 0.048 | 0.052 | 0.046 | 0.047 | 0.044 |
| 61 | 0.049 | 0.052 | 0.046 | 0.048 | 0.045 |
| 62 | 0.049 | 0.052 | 0.047 | 0.048 | 0.046 |
| 63 | 0.050 | 0.052 | 0.048 | 0.049 | 0.047 |
| 64 | 0.050 | 0.052 | 0.049 | 0.049 | 0.048 |
| 65 | 0.050 | 0.052 | 0.051 | 0.049 | 0.048 |
| 66 | 0.051 | 0.052 | 0.052 | 0.049 | 0.049 |

| | | | | | |
|---|---|---|---|---|---|
| 67 | 0.051 | 0.052 | 0.054 | 0.050 | 0.049 |
| 68 | 0.052 | 0.053 | 0.055 | 0.050 | 0.050 |
| 69 | 0.052 | 0.053 | 0.055 | 0.051 | 0.050 |
| 70 | 0.053 | 0.053 | 0.056 | 0.052 | 0.050 |
| 71 | 0.053 | 0.053 | 0.056 | 0.053 | 0.050 |
| 72 | 0.053 | 0.053 | 0.056 | 0.055 | 0.051 |
| 73 | 0.053 | 0.053 | 0.056 | 0.055 | 0.051 |
| 74 | 0.055 | 0.053 | 0.056 | 0.055 | 0.051 |
| 75 | 0.055 | 0.053 | 0.056 | 0.055 | 0.051 |
| 76 | 0.055 | 0.053 | 0.056 | 0.055 | 0.052 |
| 77 | 0.055 | 0.053 | 0.056 | 0.056 | 0.052 |
| 78 | 0.055 | 0.054 | 0.056 | 0.056 | 0.052 |
| 79 | 0.056 | 0.054 | 0.056 | 0.057 | 0.053 |
| 80 | 0.056 | 0.054 | 0.056 | 0.057 | 0.053 |
| 81 | 0.056 | 0.055 | 0.056 | 0.058 | 0.054 |
| 82 | 0.056 | 0.055 | 0.056 | 0.059 | 0.054 |
| 83 | 0.057 | 0.055 | 0.056 | 0.059 | 0.055 |
| 84 | 0.057 | 0.055 | 0.056 | 0.060 | 0.056 |
| 85 | 0.058 | 0.055 | 0.057 | 0.062 | 0.056 |
| 86 | 0.058 | 0.056 | 0.057 | 0.064 | 0.057 |
| 87 | 0.058 | 0.057 | 0.057 | 0.066 | 0.057 |
| 88 | 0.059 | 0.058 | 0.057 | 0.068 | 0.058 |
| 89 | 0.059 | 0.058 | 0.058 | 0.068 | 0.058 |
| 90 | 0.060 | 0.059 | 0.059 | 0.068 | 0.058 |
| 91 | 0.061 | 0.059 | 0.060 | 0.068 | 0.059 |
| 92 | 0.062 | 0.059 | 0.061 | 0.068 | 0.059 |
| 93 | 0.064 | 0.059 | 0.062 | 0.069 | 0.060 |
| 94 | 0.066 | 0.060 | 0.063 | 0.069 | 0.060 |
| 95 | 0.067 | 0.060 | 0.064 | 0.070 | 0.061 |
| 96 | 0.068 | 0.060 | 0.065 | 0.071 | 0.062 |
| 97 | 0.070 | 0.064 | 0.068 | 0.071 | 0.063 |
| 98 | 0.071 | 0.068 | 0.072 | 0.071 | 0.064 |
| 99 | 0.076 | 0.072 | 0.076 | 0.071 | 0.065 |
| 100 | 0.080 | 0.076 | 0.080 | 0.071 | 0.066 |

***Table S36:*** *Paced tapping with tones – Variability (CV of ITI), slow tempo – Paced_metro_750_mean_CV_iti*

| | | | | | |
|---|---|---|---|---|---|
| **Task** | Paced tapping with tones | | | | |
| **Outcome measure** | Variability (CV of ITI), slow tempo | | | | |
| **Variable name** | Paced_metro_750_mean_CV_iti | | | | |
| **Age and gender effects** | | | | | |
| Age regression (slope) | 0.000 | | | | |
| Age regression (p) | 0.377 | | | | |
| Age group (p) | 0.895 | | | | |
| Gender (p) | 0.828 | | | | |
| **Group** | All | Age 18 to 21 | Age 22 to 29 | Age 30 to 54 | Age 55 to 87 |
| N (N female) | 107 (73) | 27 (18) | 29 (19) | 26 (16) | 25 (20) |
| **Normality** | | | | | |
| Skewness | 0.51 | 1.02 | 0.25 | 0.27 | 0.48 |
| Excess kurtosis | -0.28 | 0.82 | -0.76 | -1.21 | -0.62 |
| **Scores** | | | | | |
| Mean | 0.050 | 0.052 | 0.051 | 0.050 | 0.049 |
| SD | 0.013 | 0.013 | 0.012 | 0.013 | 0.014 |
| Percentiles | | | | | |
| 0 | 0.027 | 0.033 | 0.034 | 0.031 | 0.027 |
| 1 | 0.031 | 0.033 | 0.034 | 0.032 | 0.028 |
| 2 | 0.031 | 0.034 | 0.034 | 0.032 | 0.029 |
| 3 | 0.032 | 0.034 | 0.034 | 0.032 | 0.030 |
| 4 | 0.032 | 0.034 | 0.034 | 0.032 | 0.031 |
| 5 | 0.033 | 0.035 | 0.034 | 0.032 | 0.031 |
| 6 | 0.033 | 0.036 | 0.034 | 0.033 | 0.031 |
| 7 | 0.034 | 0.037 | 0.034 | 0.033 | 0.032 |
| 8 | 0.034 | 0.037 | 0.035 | 0.033 | 0.032 |
| 9 | 0.034 | 0.038 | 0.035 | 0.034 | 0.032 |
| 10 | 0.034 | 0.038 | 0.036 | 0.035 | 0.033 |
| 11 | 0.035 | 0.038 | 0.036 | 0.036 | 0.034 |
| 12 | 0.037 | 0.039 | 0.036 | 0.037 | 0.034 |
| 13 | 0.037 | 0.040 | 0.037 | 0.037 | 0.035 |
| 14 | 0.037 | 0.041 | 0.037 | 0.037 | 0.035 |
| 15 | 0.037 | 0.042 | 0.037 | 0.037 | 0.036 |
| 16 | 0.037 | 0.042 | 0.038 | 0.037 | 0.037 |
| 17 | 0.038 | 0.043 | 0.038 | 0.038 | 0.037 |
| 18 | 0.038 | 0.043 | 0.038 | 0.038 | 0.037 |
| 19 | 0.038 | 0.043 | 0.038 | 0.038 | 0.037 |
| 20 | 0.039 | 0.043 | 0.039 | 0.039 | 0.037 |
| 21 | 0.039 | 0.043 | 0.039 | 0.039 | 0.038 |
| 22 | 0.039 | 0.043 | 0.039 | 0.039 | 0.038 |

| | | | | | |
|---|---|---|---|---|---|
| 23 | 0.039 | 0.044 | 0.039 | 0.040 | 0.038 |
| 24 | 0.039 | 0.044 | 0.039 | 0.040 | 0.038 |
| 25 | 0.039 | 0.044 | 0.039 | 0.040 | 0.038 |
| 26 | 0.040 | 0.045 | 0.040 | 0.040 | 0.038 |
| 27 | 0.040 | 0.045 | 0.040 | 0.040 | 0.038 |
| 28 | 0.041 | 0.045 | 0.041 | 0.040 | 0.038 |
| 29 | 0.041 | 0.045 | 0.041 | 0.041 | 0.039 |
| 30 | 0.042 | 0.045 | 0.042 | 0.041 | 0.039 |
| 31 | 0.042 | 0.045 | 0.042 | 0.041 | 0.039 |
| 32 | 0.043 | 0.045 | 0.043 | 0.041 | 0.039 |
| 33 | 0.043 | 0.046 | 0.044 | 0.042 | 0.039 |
| 34 | 0.044 | 0.046 | 0.045 | 0.043 | 0.039 |
| 35 | 0.044 | 0.046 | 0.046 | 0.043 | 0.039 |
| 36 | 0.044 | 0.046 | 0.047 | 0.044 | 0.039 |
| 37 | 0.045 | 0.046 | 0.048 | 0.044 | 0.039 |
| 38 | 0.045 | 0.046 | 0.048 | 0.044 | 0.040 |
| 39 | 0.045 | 0.046 | 0.049 | 0.044 | 0.040 |
| 40 | 0.046 | 0.046 | 0.049 | 0.044 | 0.041 |
| 41 | 0.046 | 0.046 | 0.049 | 0.044 | 0.041 |
| 42 | 0.047 | 0.046 | 0.049 | 0.044 | 0.042 |
| 43 | 0.047 | 0.047 | 0.049 | 0.045 | 0.044 |
| 44 | 0.047 | 0.047 | 0.049 | 0.045 | 0.045 |
| 45 | 0.047 | 0.047 | 0.050 | 0.046 | 0.047 |
| 46 | 0.048 | 0.047 | 0.050 | 0.047 | 0.049 |
| 47 | 0.049 | 0.047 | 0.050 | 0.048 | 0.049 |
| 48 | 0.049 | 0.047 | 0.050 | 0.049 | 0.049 |
| 49 | 0.049 | 0.047 | 0.050 | 0.049 | 0.049 |
| 50 | 0.049 | 0.047 | 0.050 | 0.049 | 0.049 |
| 51 | 0.049 | 0.047 | 0.050 | 0.049 | 0.049 |
| 52 | 0.049 | 0.047 | 0.050 | 0.049 | 0.049 |
| 53 | 0.050 | 0.047 | 0.051 | 0.050 | 0.049 |
| 54 | 0.050 | 0.048 | 0.051 | 0.050 | 0.049 |
| 55 | 0.050 | 0.048 | 0.051 | 0.050 | 0.050 |
| 56 | 0.050 | 0.048 | 0.052 | 0.050 | 0.050 |
| 57 | 0.050 | 0.049 | 0.052 | 0.050 | 0.050 |
| 58 | 0.050 | 0.049 | 0.053 | 0.050 | 0.050 |
| 59 | 0.051 | 0.051 | 0.053 | 0.050 | 0.051 |
| 60 | 0.051 | 0.052 | 0.054 | 0.050 | 0.051 |
| 61 | 0.052 | 0.053 | 0.055 | 0.050 | 0.052 |
| 62 | 0.053 | 0.054 | 0.055 | 0.050 | 0.052 |
| 63 | 0.054 | 0.054 | 0.055 | 0.050 | 0.053 |
| 64 | 0.054 | 0.054 | 0.056 | 0.050 | 0.053 |
| 65 | 0.054 | 0.054 | 0.056 | 0.052 | 0.053 |
| 66 | 0.056 | 0.055 | 0.056 | 0.054 | 0.053 |

| | | | | | |
|---|---|---|---|---|---|
| 67 | 0.056 | 0.055 | 0.056 | 0.056 | 0.053 |
| 68 | 0.056 | 0.056 | 0.056 | 0.058 | 0.055 |
| 69 | 0.057 | 0.056 | 0.057 | 0.059 | 0.056 |
| 70 | 0.058 | 0.056 | 0.057 | 0.060 | 0.057 |
| 71 | 0.058 | 0.057 | 0.058 | 0.061 | 0.058 |
| 72 | 0.058 | 0.057 | 0.058 | 0.061 | 0.058 |
| 73 | 0.059 | 0.057 | 0.058 | 0.062 | 0.059 |
| 74 | 0.059 | 0.057 | 0.059 | 0.062 | 0.059 |
| 75 | 0.059 | 0.058 | 0.059 | 0.063 | 0.059 |
| 76 | 0.060 | 0.058 | 0.059 | 0.063 | 0.059 |
| 77 | 0.060 | 0.059 | 0.059 | 0.063 | 0.059 |
| 78 | 0.061 | 0.060 | 0.060 | 0.063 | 0.059 |
| 79 | 0.061 | 0.061 | 0.060 | 0.063 | 0.059 |
| 80 | 0.063 | 0.063 | 0.060 | 0.063 | 0.060 |
| 81 | 0.063 | 0.064 | 0.060 | 0.064 | 0.060 |
| 82 | 0.064 | 0.064 | 0.060 | 0.064 | 0.061 |
| 83 | 0.064 | 0.064 | 0.061 | 0.064 | 0.061 |
| 84 | 0.064 | 0.064 | 0.062 | 0.064 | 0.062 |
| 85 | 0.064 | 0.065 | 0.063 | 0.065 | 0.062 |
| 86 | 0.064 | 0.066 | 0.064 | 0.065 | 0.063 |
| 87 | 0.065 | 0.067 | 0.064 | 0.065 | 0.064 |
| 88 | 0.066 | 0.067 | 0.064 | 0.065 | 0.064 |
| 89 | 0.068 | 0.068 | 0.065 | 0.066 | 0.066 |
| 90 | 0.068 | 0.068 | 0.065 | 0.067 | 0.067 |
| 91 | 0.068 | 0.068 | 0.066 | 0.069 | 0.069 |
| 92 | 0.069 | 0.068 | 0.067 | 0.070 | 0.070 |
| 93 | 0.070 | 0.070 | 0.067 | 0.070 | 0.072 |
| 94 | 0.070 | 0.072 | 0.068 | 0.070 | 0.074 |
| 95 | 0.071 | 0.075 | 0.068 | 0.070 | 0.075 |
| 96 | 0.076 | 0.077 | 0.068 | 0.070 | 0.077 |
| 97 | 0.077 | 0.080 | 0.070 | 0.071 | 0.078 |
| 98 | 0.078 | 0.083 | 0.072 | 0.071 | 0.078 |
| 99 | 0.079 | 0.086 | 0.075 | 0.071 | 0.079 |
| 100 | 0.089 | 0.089 | 0.077 | 0.072 | 0.079 |

***Table S37:*** *Paced tapping with tones – Consistency (logit of vector length), fast tempo – Paced_metro_450_vecLenLogit*

| | | | | | | |
|---|---|---|---|---|---|---|
| **Task** | | Paced tapping with tones | | | | |
| **Outcome measure** | | Consistency (logit of vector length), fast tempo | | | | |
| **Variable name** | | Paced_metro_450_vecLenLogit | | | | |
| **Age and gender effects** | | | | | | |
| Age regression (slope) | | 0.008 | | | | |
| Age regression (p) | | 0.022 | | | | |
| Age group (p) | | 0.036 | | | | |
| Gender (p) | | 0.855 | | | | |
| **Group** | | All | Age 18 to 21 | Age 22 to 29 | Age 30 to 54 | Age 55 to 87 |
| N (N female) | | 107 (73) | 27 (18) | 29 (19) | 26 (16) | 25 (20) |
| **Normality** | | | | | | |
| Skewness | | -1.66 | -2.52 | -1.00 | -1.29 | -0.13 |
| Excess kurtosis | | 3.82 | 8.30 | 1.16 | 0.69 | -0.12 |
| **Scores** | | | | | | |
| Mean | | 2.77 | 2.58 | 2.60 | 2.82 | 3.11 |
| SD | | 0.86 | 0.87 | 0.93 | 1.01 | 0.43 |
| Percentiles | | | | | | |
| | 0 | -0.98 | -0.98 | -0.25 | 0.25 | 2.09 |
| | 1 | -0.22 | -0.31 | 0.14 | 0.35 | 2.21 |
| | 2 | 0.29 | 0.37 | 0.53 | 0.45 | 2.33 |
| | 3 | 0.66 | 1.05 | 0.93 | 0.55 | 2.45 |
| | 4 | 0.80 | 1.63 | 1.20 | 0.65 | 2.57 |
| | 5 | 1.27 | 1.71 | 1.30 | 0.66 | 2.59 |
| | 6 | 1.54 | 1.79 | 1.41 | 0.67 | 2.60 |
| | 7 | 1.58 | 1.87 | 1.52 | 0.68 | 2.61 |
| | 8 | 1.65 | 1.94 | 1.54 | 0.69 | 2.62 |
| | 9 | 1.69 | 1.97 | 1.54 | 0.94 | 2.63 |
| | 10 | 1.73 | 2.01 | 1.54 | 1.19 | 2.64 |
| | 11 | 1.87 | 2.04 | 1.55 | 1.44 | 2.65 |
| | 12 | 1.93 | 2.06 | 1.60 | 1.69 | 2.67 |
| | 13 | 1.94 | 2.06 | 1.64 | 1.75 | 2.68 |
| | 14 | 2.04 | 2.07 | 1.68 | 1.81 | 2.68 |
| | 15 | 2.07 | 2.07 | 1.71 | 1.87 | 2.69 |
| | 16 | 2.08 | 2.09 | 1.72 | 1.94 | 2.69 |
| | 17 | 2.09 | 2.13 | 1.73 | 2.01 | 2.70 |
| | 18 | 2.13 | 2.17 | 1.75 | 2.09 | 2.70 |
| | 19 | 2.22 | 2.21 | 1.81 | 2.17 | 2.71 |
| | 20 | 2.23 | 2.22 | 1.86 | 2.25 | 2.71 |
| | 21 | 2.25 | 2.23 | 1.91 | 2.33 | 2.72 |
| | 22 | 2.27 | 2.24 | 1.96 | 2.41 | 2.73 |

| | | | | | |
|---|---|---|---|---|---|
| 23 | 2.30 | 2.25 | 2.00 | 2.50 | 2.75 |
| 24 | 2.33 | 2.26 | 2.04 | 2.58 | 2.76 |
| 25 | 2.38 | 2.28 | 2.08 | 2.60 | 2.78 |
| 26 | 2.43 | 2.29 | 2.09 | 2.63 | 2.80 |
| 27 | 2.47 | 2.30 | 2.10 | 2.65 | 2.83 |
| 28 | 2.49 | 2.32 | 2.12 | 2.68 | 2.85 |
| 29 | 2.56 | 2.34 | 2.13 | 2.69 | 2.87 |
| 30 | 2.59 | 2.35 | 2.16 | 2.69 | 2.88 |
| 31 | 2.59 | 2.37 | 2.19 | 2.70 | 2.88 |
| 32 | 2.62 | 2.37 | 2.22 | 2.71 | 2.88 |
| 33 | 2.67 | 2.38 | 2.24 | 2.72 | 2.88 |
| 34 | 2.68 | 2.38 | 2.27 | 2.74 | 2.89 |
| 35 | 2.70 | 2.40 | 2.29 | 2.75 | 2.90 |
| 36 | 2.71 | 2.43 | 2.32 | 2.77 | 2.92 |
| 37 | 2.72 | 2.45 | 2.36 | 2.81 | 2.93 |
| 38 | 2.74 | 2.48 | 2.41 | 2.84 | 2.95 |
| 39 | 2.75 | 2.51 | 2.45 | 2.88 | 3.00 |
| 40 | 2.77 | 2.53 | 2.46 | 2.92 | 3.04 |
| 41 | 2.77 | 2.56 | 2.46 | 2.97 | 3.08 |
| 42 | 2.81 | 2.58 | 2.47 | 3.02 | 3.11 |
| 43 | 2.84 | 2.61 | 2.48 | 3.07 | 3.11 |
| 44 | 2.86 | 2.65 | 2.58 | 3.13 | 3.11 |
| 45 | 2.87 | 2.69 | 2.69 | 3.13 | 3.11 |
| 46 | 2.88 | 2.73 | 2.79 | 3.13 | 3.11 |
| 47 | 2.88 | 2.74 | 2.84 | 3.14 | 3.13 |
| 48 | 2.92 | 2.74 | 2.84 | 3.14 | 3.14 |
| 49 | 2.93 | 2.74 | 2.84 | 3.15 | 3.15 |
| 50 | 2.94 | 2.75 | 2.85 | 3.16 | 3.16 |
| 51 | 3.00 | 2.75 | 2.89 | 3.17 | 3.16 |
| 52 | 3.05 | 2.76 | 2.93 | 3.18 | 3.17 |
| 53 | 3.05 | 2.76 | 2.97 | 3.18 | 3.17 |
| 54 | 3.07 | 2.77 | 3.00 | 3.18 | 3.17 |
| 55 | 3.08 | 2.80 | 3.02 | 3.18 | 3.18 |
| 56 | 3.10 | 2.82 | 3.03 | 3.19 | 3.18 |
| 57 | 3.10 | 2.85 | 3.05 | 3.22 | 3.18 |
| 58 | 3.11 | 2.86 | 3.06 | 3.26 | 3.18 |
| 59 | 3.12 | 2.87 | 3.07 | 3.30 | 3.19 |
| 60 | 3.14 | 2.87 | 3.09 | 3.34 | 3.19 |
| 61 | 3.15 | 2.87 | 3.10 | 3.36 | 3.19 |
| 62 | 3.15 | 2.88 | 3.12 | 3.39 | 3.19 |
| 63 | 3.16 | 2.90 | 3.13 | 3.41 | 3.20 |
| 64 | 3.16 | 2.91 | 3.15 | 3.44 | 3.23 |
| 65 | 3.17 | 2.93 | 3.15 | 3.45 | 3.26 |
| 66 | 3.17 | 2.96 | 3.16 | 3.46 | 3.29 |

| | | | | | |
|---|---|---|---|---|---|
| 67 | 3.18 | 2.98 | 3.16 | 3.47 | 3.32 |
| 68 | 3.19 | 3.01 | 3.16 | 3.49 | 3.34 |
| 69 | 3.19 | 3.04 | 3.17 | 3.50 | 3.36 |
| 70 | 3.19 | 3.05 | 3.17 | 3.50 | 3.37 |
| 71 | 3.22 | 3.06 | 3.17 | 3.51 | 3.39 |
| 72 | 3.23 | 3.06 | 3.18 | 3.52 | 3.39 |
| 73 | 3.27 | 3.06 | 3.19 | 3.52 | 3.39 |
| 74 | 3.32 | 3.07 | 3.21 | 3.52 | 3.39 |
| 75 | 3.36 | 3.07 | 3.22 | 3.52 | 3.39 |
| 76 | 3.39 | 3.07 | 3.22 | 3.52 | 3.40 |
| 77 | 3.41 | 3.08 | 3.22 | 3.52 | 3.41 |
| 78 | 3.43 | 3.08 | 3.23 | 3.52 | 3.42 |
| 79 | 3.44 | 3.09 | 3.23 | 3.53 | 3.43 |
| 80 | 3.44 | 3.09 | 3.23 | 3.53 | 3.44 |
| 81 | 3.45 | 3.10 | 3.23 | 3.55 | 3.44 |
| 82 | 3.46 | 3.12 | 3.24 | 3.57 | 3.44 |
| 83 | 3.49 | 3.13 | 3.29 | 3.59 | 3.44 |
| 84 | 3.49 | 3.14 | 3.35 | 3.61 | 3.45 |
| 85 | 3.50 | 3.18 | 3.41 | 3.61 | 3.46 |
| 86 | 3.52 | 3.25 | 3.46 | 3.61 | 3.48 |
| 87 | 3.52 | 3.32 | 3.50 | 3.61 | 3.49 |
| 88 | 3.52 | 3.39 | 3.53 | 3.61 | 3.51 |
| 89 | 3.55 | 3.43 | 3.57 | 3.65 | 3.53 |
| 90 | 3.59 | 3.44 | 3.59 | 3.68 | 3.56 |
| 91 | 3.61 | 3.45 | 3.61 | 3.71 | 3.59 |
| 92 | 3.61 | 3.45 | 3.62 | 3.74 | 3.62 |
| 93 | 3.63 | 3.46 | 3.64 | 3.75 | 3.68 |
| 94 | 3.70 | 3.47 | 3.68 | 3.76 | 3.74 |
| 95 | 3.76 | 3.48 | 3.72 | 3.77 | 3.80 |
| 96 | 3.78 | 3.49 | 3.76 | 3.78 | 3.86 |
| 97 | 3.84 | 3.49 | 3.81 | 3.83 | 3.88 |
| 98 | 3.93 | 3.50 | 3.87 | 3.88 | 3.90 |
| 99 | 3.97 | 3.51 | 3.94 | 3.93 | 3.92 |
| 100 | 4.00 | 3.52 | 4.00 | 3.97 | 3.94 |

***Table S38:*** *Paced tapping with tones – Consistency (logit of vector length), medium tempo – Paced_metro_600_vecLenLogit*

| | | | | | | |
|---|---|---|---|---|---|---|
| **Task** | | Paced tapping with tones | | | | |
| **Outcome measure** | | Consistency (logit of vector length), medium tempo | | | | |
| **Variable name** | | Paced_metro_600_vecLenLogit | | | | |
| **Age and gender effects** | | | | | | |
| Age regression (slope) | | 0.006 | | | | |
| Age regression (p) | | 0.072 | | | | |
| Age group (p) | | 0.403 | | | | |
| Gender (p) | | 0.150 | | | | |
| **Group** | | All | Age 18 to 21 | Age 22 to 29 | Age 30 to 54 | Age 55 to 87 |
| N (N female) | | 108 (74) | 27 (18) | 29 (19) | 26 (16) | 26 (21) |
| **Normality** | | | | | | |
| Skewness | | -1.23 | -1.23 | -0.96 | -0.96 | -1.73 |
| Excess kurtosis | | 1.76 | 1.36 | 0.93 | 0.49 | 4.79 |
| **Scores** | | | | | | |
| Mean | | 2.97 | 2.93 | 2.98 | 2.78 | 3.19 |
| SD | | 0.78 | 0.68 | 0.73 | 0.97 | 0.72 |
| Percentiles | | | | | | |
| | 0 | 0.10 | 1.01 | 0.84 | 0.10 | 0.59 |
| | 1 | 0.61 | 1.08 | 1.05 | 0.37 | 1.08 |
| | 2 | 0.87 | 1.16 | 1.26 | 0.65 | 1.57 |
| | 3 | 1.05 | 1.23 | 1.46 | 0.92 | 2.06 |
| | 4 | 1.22 | 1.32 | 1.63 | 1.20 | 2.56 |
| | 5 | 1.34 | 1.48 | 1.73 | 1.26 | 2.56 |
| | 6 | 1.48 | 1.65 | 1.84 | 1.32 | 2.56 |
| | 7 | 1.56 | 1.81 | 1.95 | 1.38 | 2.57 |
| | 8 | 1.61 | 1.95 | 1.99 | 1.44 | 2.57 |
| | 9 | 1.82 | 2.05 | 2.01 | 1.47 | 2.57 |
| | 10 | 1.95 | 2.14 | 2.04 | 1.49 | 2.57 |
| | 11 | 2.00 | 2.23 | 2.09 | 1.51 | 2.58 |
| | 12 | 2.04 | 2.30 | 2.21 | 1.54 | 2.58 |
| | 13 | 2.06 | 2.36 | 2.32 | 1.56 | 2.59 |
| | 14 | 2.27 | 2.41 | 2.43 | 1.59 | 2.59 |
| | 15 | 2.46 | 2.47 | 2.47 | 1.61 | 2.60 |
| | 16 | 2.49 | 2.53 | 2.49 | 1.64 | 2.60 |
| | 17 | 2.52 | 2.58 | 2.50 | 1.73 | 2.65 |
| | 18 | 2.54 | 2.64 | 2.52 | 1.82 | 2.69 |
| | 19 | 2.56 | 2.70 | 2.53 | 1.92 | 2.73 |
| | 20 | 2.57 | 2.72 | 2.55 | 2.01 | 2.77 |
| | 21 | 2.57 | 2.72 | 2.56 | 2.02 | 2.80 |
| | 22 | 2.59 | 2.72 | 2.57 | 2.03 | 2.83 |

| | | | | | |
|---|---|---|---|---|---|
| 23 | 2.61 | 2.72 | 2.58 | 2.03 | 2.86 |
| 24 | 2.62 | 2.73 | 2.60 | 2.04 | 2.88 |
| 25 | 2.70 | 2.74 | 2.61 | 2.17 | 2.91 |
| 26 | 2.72 | 2.75 | 2.61 | 2.29 | 2.93 |
| 27 | 2.75 | 2.75 | 2.62 | 2.41 | 2.95 |
| 28 | 2.76 | 2.76 | 2.63 | 2.54 | 2.97 |
| 29 | 2.77 | 2.76 | 2.65 | 2.60 | 2.97 |
| 30 | 2.79 | 2.76 | 2.70 | 2.66 | 2.97 |
| 31 | 2.80 | 2.77 | 2.75 | 2.72 | 2.97 |
| 32 | 2.83 | 2.79 | 2.79 | 2.79 | 2.97 |
| 33 | 2.86 | 2.82 | 2.82 | 2.80 | 2.99 |
| 34 | 2.87 | 2.84 | 2.84 | 2.81 | 3.01 |
| 35 | 2.88 | 2.86 | 2.86 | 2.82 | 3.03 |
| 36 | 2.88 | 2.86 | 2.89 | 2.82 | 3.05 |
| 37 | 2.89 | 2.87 | 2.90 | 2.83 | 3.07 |
| 38 | 2.93 | 2.87 | 2.92 | 2.84 | 3.08 |
| 39 | 2.95 | 2.87 | 2.94 | 2.85 | 3.09 |
| 40 | 2.96 | 2.88 | 2.95 | 2.86 | 3.10 |
| 41 | 2.96 | 2.88 | 2.96 | 2.89 | 3.11 |
| 42 | 2.97 | 2.89 | 2.96 | 2.91 | 3.13 |
| 43 | 2.97 | 2.90 | 2.97 | 2.94 | 3.14 |
| 44 | 2.97 | 2.92 | 2.97 | 2.96 | 3.16 |
| 45 | 2.98 | 2.94 | 2.98 | 2.97 | 3.17 |
| 46 | 3.00 | 2.95 | 2.98 | 2.98 | 3.18 |
| 47 | 3.02 | 2.96 | 2.99 | 2.99 | 3.20 |
| 48 | 3.03 | 2.96 | 3.01 | 3.00 | 3.21 |
| 49 | 3.04 | 2.96 | 3.03 | 3.00 | 3.25 |
| 50 | 3.05 | 2.96 | 3.05 | 3.01 | 3.29 |
| 51 | 3.06 | 2.98 | 3.05 | 3.01 | 3.32 |
| 52 | 3.07 | 3.00 | 3.06 | 3.01 | 3.36 |
| 53 | 3.09 | 3.01 | 3.07 | 3.03 | 3.37 |
| 54 | 3.13 | 3.02 | 3.09 | 3.04 | 3.38 |
| 55 | 3.14 | 3.03 | 3.11 | 3.05 | 3.40 |
| 56 | 3.15 | 3.03 | 3.14 | 3.07 | 3.41 |
| 57 | 3.17 | 3.03 | 3.16 | 3.09 | 3.41 |
| 58 | 3.21 | 3.05 | 3.18 | 3.10 | 3.41 |
| 59 | 3.22 | 3.07 | 3.20 | 3.12 | 3.41 |
| 60 | 3.24 | 3.10 | 3.22 | 3.14 | 3.42 |
| 61 | 3.26 | 3.13 | 3.23 | 3.16 | 3.44 |
| 62 | 3.27 | 3.16 | 3.24 | 3.18 | 3.47 |
| 63 | 3.29 | 3.20 | 3.25 | 3.20 | 3.50 |
| 64 | 3.32 | 3.24 | 3.26 | 3.22 | 3.53 |
| 65 | 3.34 | 3.28 | 3.27 | 3.23 | 3.55 |
| 66 | 3.36 | 3.31 | 3.27 | 3.24 | 3.57 |

| | | | | | |
|---|---|---|---|---|---|
| 67 | 3.37 | 3.32 | 3.28 | 3.25 | 3.59 |
| 68 | 3.39 | 3.33 | 3.29 | 3.25 | 3.61 |
| 69 | 3.41 | 3.34 | 3.30 | 3.28 | 3.62 |
| 70 | 3.41 | 3.36 | 3.32 | 3.31 | 3.62 |
| 71 | 3.42 | 3.38 | 3.33 | 3.34 | 3.63 |
| 72 | 3.44 | 3.40 | 3.36 | 3.37 | 3.63 |
| 73 | 3.44 | 3.41 | 3.40 | 3.38 | 3.63 |
| 74 | 3.47 | 3.42 | 3.44 | 3.39 | 3.63 |
| 75 | 3.49 | 3.43 | 3.47 | 3.39 | 3.63 |
| 76 | 3.56 | 3.43 | 3.55 | 3.40 | 3.63 |
| 77 | 3.61 | 3.44 | 3.62 | 3.41 | 3.64 |
| 78 | 3.62 | 3.45 | 3.70 | 3.42 | 3.64 |
| 79 | 3.63 | 3.46 | 3.75 | 3.43 | 3.64 |
| 80 | 3.63 | 3.47 | 3.75 | 3.43 | 3.65 |
| 81 | 3.63 | 3.48 | 3.76 | 3.48 | 3.65 |
| 82 | 3.64 | 3.52 | 3.77 | 3.53 | 3.66 |
| 83 | 3.65 | 3.55 | 3.77 | 3.58 | 3.67 |
| 84 | 3.67 | 3.59 | 3.77 | 3.63 | 3.68 |
| 85 | 3.69 | 3.61 | 3.78 | 3.65 | 3.72 |
| 86 | 3.72 | 3.62 | 3.78 | 3.66 | 3.76 |
| 87 | 3.75 | 3.62 | 3.78 | 3.68 | 3.81 |
| 88 | 3.77 | 3.62 | 3.79 | 3.69 | 3.85 |
| 89 | 3.78 | 3.63 | 3.79 | 3.72 | 3.85 |
| 90 | 3.80 | 3.63 | 3.80 | 3.75 | 3.85 |
| 91 | 3.81 | 3.64 | 3.80 | 3.78 | 3.86 |
| 92 | 3.81 | 3.65 | 3.81 | 3.81 | 3.86 |
| 93 | 3.83 | 3.66 | 3.82 | 3.87 | 3.92 |
| 94 | 3.86 | 3.68 | 3.84 | 3.93 | 3.98 |
| 95 | 3.88 | 3.70 | 3.86 | 3.99 | 4.05 |
| 96 | 3.93 | 3.71 | 3.88 | 4.04 | 4.11 |
| 97 | 4.02 | 3.74 | 3.89 | 4.07 | 4.15 |
| 98 | 4.10 | 3.76 | 3.91 | 4.10 | 4.19 |
| 99 | 4.16 | 3.79 | 3.93 | 4.13 | 4.23 |
| 100 | 4.28 | 3.81 | 3.95 | 4.16 | 4.28 |

***Table S39:*** *Paced tapping with tones – Consistency (logit of vector length), slow tempo – Paced_metro_750_vecLenLogit*

| | | | | | | |
|---|---|---|---|---|---|---|
| **Task** | Paced tapping with tones | | | | | |
| **Outcome measure** | Consistency (logit of vector length), slow tempo | | | | | |
| **Variable name** | Paced_metro_750_vecLenLogit | | | | | |
| **Age and gender effects** | | | | | | |
| Age regression (slope) | | 0.001 | | | | |
| Age regression (p) | | 0.754 | | | | |
| Age group (p) | | 0.950 | | | | |
| Gender (p) | | 0.125 | | | | |
| **Group** | | All | Age 18 to 21 | Age 22 to 29 | Age 30 to 54 | Age 55 to 87 |
| N (N female) | | 107 (73) | 27 (18) | 29 (19) | 26 (16) | 25 (20) |
| **Normality** | | | | | | |
| Skewness | | -1.65 | -1.95 | -1.79 | -1.18 | -1.65 |
| Excess kurtosis | | 3.17 | 4.10 | 3.75 | 0.97 | 3.44 |
| **Scores** | | | | | | |
| Mean | | 2.94 | 2.99 | 2.98 | 2.84 | 2.94 |
| SD | | 0.95 | 0.89 | 0.84 | 1.00 | 1.10 |
| Percentiles | | | | | | |
| | 0 | -0.78 | 0.12 | 0.05 | 0.02 | -0.78 |
| | 1 | 0.02 | 0.21 | 0.42 | 0.26 | -0.31 |
| | 2 | 0.06 | 0.31 | 0.79 | 0.49 | 0.15 |
| | 3 | 0.19 | 0.40 | 1.15 | 0.72 | 0.62 |
| | 4 | 0.60 | 0.56 | 1.37 | 0.96 | 1.09 |
| | 5 | 1.02 | 1.06 | 1.39 | 1.05 | 1.25 |
| | 6 | 1.23 | 1.56 | 1.41 | 1.15 | 1.34 |
| | 7 | 1.35 | 2.07 | 1.43 | 1.24 | 1.43 |
| | 8 | 1.36 | 2.44 | 1.71 | 1.34 | 1.52 |
| | 9 | 1.40 | 2.51 | 2.03 | 1.35 | 1.60 |
| | 10 | 1.50 | 2.59 | 2.35 | 1.35 | 1.68 |
| | 11 | 1.76 | 2.66 | 2.58 | 1.36 | 1.76 |
| | 12 | 2.10 | 2.70 | 2.58 | 1.37 | 1.83 |
| | 13 | 2.25 | 2.71 | 2.59 | 1.57 | 1.95 |
| | 14 | 2.39 | 2.72 | 2.59 | 1.78 | 2.11 |
| | 15 | 2.46 | 2.72 | 2.60 | 1.98 | 2.27 |
| | 16 | 2.54 | 2.73 | 2.61 | 2.19 | 2.43 |
| | 17 | 2.58 | 2.73 | 2.61 | 2.21 | 2.55 |
| | 18 | 2.60 | 2.73 | 2.62 | 2.23 | 2.59 |
| | 19 | 2.62 | 2.73 | 2.62 | 2.25 | 2.63 |
| | 20 | 2.63 | 2.73 | 2.62 | 2.27 | 2.66 |
| | 21 | 2.69 | 2.74 | 2.62 | 2.32 | 2.70 |
| | 22 | 2.70 | 2.74 | 2.65 | 2.37 | 2.71 |

| | | | | | |
|---|---|---|---|---|---|
| 23 | 2.71 | 2.75 | 2.71 | 2.42 | 2.73 |
| 24 | 2.73 | 2.76 | 2.77 | 2.47 | 2.74 |
| 25 | 2.73 | 2.77 | 2.83 | 2.52 | 2.76 |
| 26 | 2.74 | 2.79 | 2.84 | 2.58 | 2.76 |
| 27 | 2.75 | 2.80 | 2.85 | 2.63 | 2.76 |
| 28 | 2.76 | 2.81 | 2.86 | 2.69 | 2.76 |
| 29 | 2.76 | 2.82 | 2.87 | 2.70 | 2.76 |
| 30 | 2.77 | 2.83 | 2.88 | 2.71 | 2.77 |
| 31 | 2.79 | 2.84 | 2.88 | 2.73 | 2.77 |
| 32 | 2.82 | 2.84 | 2.89 | 2.74 | 2.77 |
| 33 | 2.84 | 2.84 | 2.90 | 2.74 | 2.77 |
| 34 | 2.84 | 2.84 | 2.92 | 2.74 | 2.78 |
| 35 | 2.84 | 2.84 | 2.93 | 2.75 | 2.80 |
| 36 | 2.85 | 2.84 | 2.95 | 2.75 | 2.82 |
| 37 | 2.87 | 2.84 | 2.96 | 2.78 | 2.83 |
| 38 | 2.88 | 2.84 | 2.98 | 2.81 | 2.85 |
| 39 | 2.89 | 2.85 | 2.99 | 2.85 | 2.87 |
| 40 | 2.90 | 2.87 | 3.01 | 2.88 | 2.88 |
| 41 | 2.91 | 2.89 | 3.02 | 2.88 | 2.90 |
| 42 | 2.93 | 2.91 | 3.03 | 2.89 | 2.91 |
| 43 | 2.94 | 2.93 | 3.04 | 2.89 | 2.92 |
| 44 | 2.98 | 2.96 | 3.04 | 2.89 | 2.93 |
| 45 | 3.02 | 3.00 | 3.04 | 2.96 | 2.94 |
| 46 | 3.04 | 3.03 | 3.04 | 3.04 | 2.95 |
| 47 | 3.04 | 3.06 | 3.05 | 3.11 | 2.98 |
| 48 | 3.07 | 3.08 | 3.06 | 3.18 | 3.01 |
| 49 | 3.08 | 3.11 | 3.07 | 3.18 | 3.05 |
| 50 | 3.10 | 3.14 | 3.08 | 3.18 | 3.08 |
| 51 | 3.10 | 3.17 | 3.11 | 3.18 | 3.08 |
| 52 | 3.14 | 3.20 | 3.13 | 3.18 | 3.09 |
| 53 | 3.17 | 3.23 | 3.16 | 3.19 | 3.09 |
| 54 | 3.18 | 3.25 | 3.18 | 3.21 | 3.09 |
| 55 | 3.19 | 3.26 | 3.19 | 3.22 | 3.10 |
| 56 | 3.22 | 3.28 | 3.20 | 3.23 | 3.10 |
| 57 | 3.24 | 3.29 | 3.21 | 3.24 | 3.10 |
| 58 | 3.26 | 3.30 | 3.22 | 3.25 | 3.10 |
| 59 | 3.27 | 3.31 | 3.24 | 3.26 | 3.16 |
| 60 | 3.27 | 3.32 | 3.26 | 3.27 | 3.26 |
| 61 | 3.29 | 3.32 | 3.27 | 3.27 | 3.35 |
| 62 | 3.30 | 3.33 | 3.30 | 3.27 | 3.45 |
| 63 | 3.32 | 3.35 | 3.32 | 3.28 | 3.49 |
| 64 | 3.34 | 3.37 | 3.34 | 3.28 | 3.50 |
| 65 | 3.35 | 3.38 | 3.35 | 3.28 | 3.50 |
| 66 | 3.38 | 3.41 | 3.36 | 3.28 | 3.50 |

| | | | | | |
|---|---|---|---|---|---|
| 67 | 3.39 | 3.44 | 3.37 | 3.29 | 3.51 |
| 68 | 3.39 | 3.46 | 3.38 | 3.29 | 3.51 |
| 69 | 3.44 | 3.49 | 3.38 | 3.30 | 3.51 |
| 70 | 3.49 | 3.50 | 3.38 | 3.31 | 3.52 |
| 71 | 3.50 | 3.50 | 3.39 | 3.33 | 3.52 |
| 72 | 3.50 | 3.51 | 3.40 | 3.34 | 3.54 |
| 73 | 3.50 | 3.51 | 3.41 | 3.38 | 3.55 |
| 74 | 3.51 | 3.52 | 3.42 | 3.42 | 3.56 |
| 75 | 3.52 | 3.53 | 3.43 | 3.46 | 3.58 |
| 76 | 3.54 | 3.54 | 3.45 | 3.50 | 3.59 |
| 77 | 3.55 | 3.55 | 3.47 | 3.51 | 3.60 |
| 78 | 3.56 | 3.56 | 3.49 | 3.52 | 3.62 |
| 79 | 3.57 | 3.57 | 3.51 | 3.53 | 3.63 |
| 80 | 3.59 | 3.58 | 3.52 | 3.55 | 3.68 |
| 81 | 3.62 | 3.59 | 3.54 | 3.59 | 3.74 |
| 82 | 3.62 | 3.60 | 3.56 | 3.62 | 3.81 |
| 83 | 3.63 | 3.61 | 3.60 | 3.66 | 3.87 |
| 84 | 3.70 | 3.61 | 3.64 | 3.70 | 3.90 |
| 85 | 3.71 | 3.62 | 3.68 | 3.71 | 3.91 |
| 86 | 3.75 | 3.62 | 3.71 | 3.72 | 3.93 |
| 87 | 3.77 | 3.62 | 3.73 | 3.74 | 3.94 |
| 88 | 3.78 | 3.62 | 3.75 | 3.75 | 3.95 |
| 89 | 3.81 | 3.65 | 3.77 | 3.79 | 3.95 |
| 90 | 3.85 | 3.68 | 3.78 | 3.83 | 3.96 |
| 91 | 3.90 | 3.72 | 3.80 | 3.87 | 3.96 |
| 92 | 3.93 | 3.76 | 3.81 | 3.92 | 3.98 |
| 93 | 3.96 | 3.78 | 3.83 | 3.93 | 4.05 |
| 94 | 3.97 | 3.79 | 3.87 | 3.95 | 4.11 |
| 95 | 3.98 | 3.80 | 3.92 | 3.96 | 4.17 |
| 96 | 4.05 | 3.80 | 3.97 | 3.98 | 4.24 |
| 97 | 4.16 | 3.91 | 4.00 | 4.03 | 4.28 |
| 98 | 4.22 | 4.04 | 4.02 | 4.08 | 4.33 |
| 99 | 4.29 | 4.17 | 4.05 | 4.13 | 4.37 |
| 100 | 4.41 | 4.29 | 4.07 | 4.18 | 4.41 |

***Table S40:*** *Paced tapping with tones – Accuracy (vector direction %), fast tempo – Paced_metro_450_vecDirPct*

| | | | | | | |
|---|---|---|---|---|---|---|
| **Task** | | Paced tapping with tones | | | | |
| **Outcome measure** | | Accuracy (vector direction %), fast tempo | | | | |
| **Variable name** | | Paced_metro_450_vecDirPct | | | | |
| **Age and gender effects** | | | | | | |
| Age regression (slope) | | 0.049 | | | | |
| Age regression (p) | | 0.127 | | | | |
| Age group (p) | | 0.479 | | | | |
| Gender (p) | | 0.477 | | | | |
| **Group** | | All | Age 18 to 21 | Age 22 to 29 | Age 30 to 54 | Age 55 to 87 |
| N (N female) | | 97 (65) | 23 (15) | 25 (15) | 24 (15) | 25 (20) |
| **Normality** | | | | | | |
| Skewness | | -0.43 | -1.02 | -0.14 | 0.23 | -0.70 |
| Excess kurtosis | | -0.23 | 0.94 | -0.99 | -0.48 | -0.37 |
| **Scores** | | | | | | |
| Mean | | -7.1 | -7.9 | -7.5 | -7.4 | -5.8 |
| SD | | 5.9 | 5.9 | 6.1 | 5.0 | 6.4 |
| Percentiles | | | | | | |
| | 0 | -24.6 | -24.6 | -18.8 | -17.4 | -21.4 |
| | 1 | -21.5 | -23.0 | -18.3 | -16.8 | -20.2 |
| | 2 | -19.0 | -21.4 | -17.8 | -16.2 | -19.1 |
| | 3 | -17.5 | -19.8 | -17.4 | -15.7 | -18.0 |
| | 4 | -17.3 | -18.2 | -16.9 | -15.1 | -16.8 |
| | 5 | -16.9 | -17.1 | -16.8 | -14.5 | -16.1 |
| | 6 | -16.7 | -16.6 | -16.7 | -13.8 | -15.5 |
| | 7 | -16.6 | -16.1 | -16.7 | -13.1 | -14.9 |
| | 8 | -15.6 | -15.6 | -16.7 | -12.4 | -14.3 |
| | 9 | -15.0 | -15.1 | -16.4 | -11.9 | -14.0 |
| | 10 | -14.9 | -15.0 | -16.0 | -11.9 | -13.9 |
| | 11 | -14.6 | -14.8 | -15.6 | -11.8 | -13.8 |
| | 12 | -14.3 | -14.6 | -15.1 | -11.7 | -13.6 |
| | 13 | -14.2 | -14.4 | -14.8 | -11.7 | -13.5 |
| | 14 | -14.1 | -14.3 | -14.6 | -11.7 | -13.4 |
| | 15 | -13.9 | -14.3 | -14.4 | -11.6 | -13.3 |
| | 16 | -13.4 | -14.3 | -14.2 | -11.6 | -13.1 |
| | 17 | -13.0 | -14.3 | -13.9 | -11.6 | -12.8 |
| | 18 | -12.7 | -14.3 | -13.7 | -11.5 | -12.2 |
| | 19 | -12.2 | -13.5 | -13.4 | -11.5 | -11.5 |
| | 20 | -11.9 | -12.6 | -13.1 | -11.5 | -10.8 |
| | 21 | -11.7 | -11.8 | -12.8 | -11.5 | -10.3 |
| | 22 | -11.7 | -10.9 | -12.7 | -11.4 | -10.2 |

| | | | | | |
|---|---|---|---|---|---|
| 23 | -11.6 | -10.2 | -12.5 | -11.2 | -10.2 |
| 24 | -11.4 | -10.0 | -12.4 | -11.0 | -10.1 |
| 25 | -10.6 | -9.8 | -12.3 | -10.9 | -10.1 |
| 26 | -10.3 | -9.6 | -12.1 | -10.7 | -9.9 |
| 27 | -10.3 | -9.4 | -12.0 | -10.6 | -9.7 |
| 28 | -10.2 | -9.2 | -11.8 | -10.5 | -9.6 |
| 29 | -10.1 | -9.1 | -11.7 | -10.4 | -9.4 |
| 30 | -9.9 | -9.0 | -11.2 | -10.3 | -9.1 |
| 31 | -9.6 | -8.9 | -10.6 | -10.2 | -8.8 |
| 32 | -9.6 | -8.8 | -10.0 | -10.1 | -8.5 |
| 33 | -9.4 | -8.7 | -9.4 | -10.0 | -8.2 |
| 34 | -9.3 | -8.6 | -9.2 | -9.9 | -8.0 |
| 35 | -9.3 | -8.4 | -9.2 | -9.9 | -7.8 |
| 36 | -9.2 | -8.3 | -9.2 | -9.8 | -7.5 |
| 37 | -9.0 | -8.2 | -9.1 | -9.7 | -7.3 |
| 38 | -8.7 | -8.1 | -9.1 | -9.7 | -7.0 |
| 39 | -8.6 | -8.0 | -9.0 | -9.6 | -6.5 |
| 40 | -8.5 | -7.9 | -8.8 | -9.6 | -6.0 |
| 41 | -8.3 | -7.8 | -8.7 | -9.6 | -5.5 |
| 42 | -8.2 | -7.7 | -8.6 | -9.6 | -5.1 |
| 43 | -8.1 | -7.6 | -8.5 | -9.6 | -4.9 |
| 44 | -7.9 | -7.4 | -8.5 | -9.4 | -4.7 |
| 45 | -7.7 | -7.3 | -8.4 | -9.2 | -4.5 |
| 46 | -7.2 | -7.2 | -8.3 | -9.0 | -4.3 |
| 47 | -7.2 | -7.1 | -8.2 | -8.7 | -4.3 |
| 48 | -7.2 | -7.0 | -8.1 | -8.5 | -4.3 |
| 49 | -7.1 | -6.9 | -8.0 | -8.2 | -4.2 |
| 50 | -6.8 | -6.8 | -7.9 | -7.8 | -4.2 |
| 51 | -6.8 | -6.7 | -7.8 | -7.5 | -4.2 |
| 52 | -6.5 | -6.6 | -7.6 | -7.2 | -4.2 |
| 53 | -6.4 | -6.5 | -7.4 | -7.1 | -4.1 |
| 54 | -6.0 | -6.4 | -7.3 | -7.0 | -4.1 |
| 55 | -5.8 | -6.3 | -6.7 | -6.9 | -4.0 |
| 56 | -5.6 | -6.1 | -6.1 | -6.8 | -3.8 |
| 57 | -5.5 | -6.0 | -5.5 | -6.8 | -3.7 |
| 58 | -5.3 | -5.8 | -4.8 | -6.7 | -3.5 |
| 59 | -5.1 | -5.6 | -4.5 | -6.6 | -3.3 |
| 60 | -4.9 | -5.6 | -4.3 | -6.6 | -3.1 |
| 61 | -4.8 | -5.6 | -4.1 | -6.5 | -2.9 |
| 62 | -4.7 | -5.5 | -3.9 | -6.3 | -2.7 |
| 63 | -4.5 | -5.5 | -3.7 | -6.2 | -2.6 |
| 64 | -4.4 | -5.4 | -3.7 | -6.1 | -2.5 |
| 65 | -4.3 | -5.4 | -3.6 | -5.9 | -2.4 |
| 66 | -4.2 | -5.3 | -3.6 | -5.9 | -2.3 |

| | | | | | |
|---|---|---|---|---|---|
| 67 | -4.0 | -5.2 | -3.6 | -5.8 | -2.1 |
| 68 | -3.7 | -5.1 | -3.5 | -5.8 | -1.8 |
| 69 | -3.5 | -5.0 | -3.5 | -5.8 | -1.4 |
| 70 | -3.5 | -4.9 | -3.5 | -5.6 | -1.1 |
| 71 | -3.4 | -4.9 | -3.4 | -5.4 | -0.8 |
| 72 | -3.4 | -4.8 | -3.4 | -5.2 | -0.6 |
| 73 | -3.1 | -4.8 | -3.4 | -5.0 | -0.4 |
| 74 | -2.9 | -4.7 | -3.4 | -4.8 | -0.3 |
| 75 | -2.8 | -4.6 | -3.4 | -4.4 | -0.1 |
| 76 | -2.7 | -4.5 | -3.3 | -4.0 | 0.0 |
| 77 | -2.6 | -4.4 | -3.2 | -3.6 | 0.1 |
| 78 | -2.3 | -4.2 | -3.1 | -3.2 | 0.3 |
| 79 | -2.0 | -3.8 | -3.0 | -2.9 | 0.4 |
| 80 | -1.5 | -3.5 | -2.6 | -2.7 | 0.4 |
| 81 | -1.2 | -3.1 | -2.2 | -2.4 | 0.4 |
| 82 | -1.2 | -2.8 | -1.8 | -2.2 | 0.5 |
| 83 | -1.0 | -2.8 | -1.3 | -1.9 | 0.5 |
| 84 | -0.9 | -2.8 | -1.1 | -1.7 | 0.5 |
| 85 | -0.7 | -2.7 | -0.9 | -1.5 | 0.6 |
| 86 | -0.5 | -2.7 | -0.8 | -1.3 | 0.6 |
| 87 | -0.4 | -2.5 | -0.6 | -1.1 | 0.6 |
| 88 | -0.2 | -2.2 | -0.5 | -0.9 | 0.7 |
| 89 | 0.0 | -1.9 | -0.5 | -0.7 | 0.8 |
| 90 | 0.3 | -1.7 | -0.5 | -0.5 | 0.8 |
| 91 | 0.4 | -1.4 | -0.5 | -0.3 | 0.9 |
| 92 | 0.5 | -1.3 | -0.3 | -0.2 | 1.0 |
| 93 | 0.7 | -1.2 | 0.3 | -0.1 | 1.3 |
| 94 | 0.9 | -1.1 | 0.8 | 0.0 | 1.6 |
| 95 | 1.2 | -1.0 | 1.4 | 0.1 | 1.9 |
| 96 | 1.9 | -0.7 | 1.9 | 0.5 | 2.1 |
| 97 | 2.2 | -0.3 | 2.2 | 1.1 | 2.2 |
| 98 | 2.5 | 0.2 | 2.5 | 1.8 | 2.3 |
| 99 | 3.0 | 0.6 | 2.8 | 2.5 | 2.3 |
| 100 | 3.2 | 1.1 | 3.0 | 3.2 | 2.4 |

***Table S41:*** *Paced tapping with tones – Accuracy (vector direction %), medium tempo – Paced_metro_600_vecDirPct*

| | | | | | | |
|---|---|---|---|---|---|---|
| **Task** | | Paced tapping with tones | | | | |
| **Outcome measure** | | Accuracy (vector direction %), medium tempo | | | | |
| **Variable name** | | Paced_metro_600_vecDirPct | | | | |
| **Age and gender effects** | | | | | | |
| Age regression (slope) | | 0.023 | | | | |
| Age regression (p) | | 0.322 | | | | |
| Age group (p) | | 0.764 | | | | |
| Gender (p) | | 0.252 | | | | |
| **Group** | | All | Age 18 to 21 | Age 22 to 29 | Age 30 to 54 | Age 55 to 87 |
| N (N female) | | 106 (73) | 27 (18) | 29 (19) | 24 (15) | 26 (21) |
| **Normality** | | | | | | |
| Skewness | | -0.94 | -0.61 | -0.82 | -1.14 | -1.22 |
| Excess kurtosis | | 0.58 | -0.66 | -0.15 | 0.82 | 1.98 |
| **Scores** | | | | | | |
| Mean | | -6.5 | -7.3 | -6.4 | -6.4 | -6.1 |
| SD | | 5.6 | 5.3 | 5.7 | 5.4 | 6.1 |
| Percentiles | | | | | | |
| | 0 | -24.9 | -19.1 | -20.7 | -21.7 | -24.9 |
| | 1 | -21.6 | -18.2 | -19.5 | -20.3 | -23.3 |
| | 2 | -20.5 | -17.3 | -18.3 | -19.0 | -21.7 |
| | 3 | -19.0 | -16.3 | -17.1 | -17.6 | -20.1 |
| | 4 | -18.0 | -15.6 | -16.4 | -16.3 | -18.4 |
| | 5 | -16.4 | -15.6 | -16.4 | -15.5 | -16.4 |
| | 6 | -16.1 | -15.5 | -16.3 | -15.0 | -14.4 |
| | 7 | -15.7 | -15.5 | -16.3 | -14.6 | -12.4 |
| | 8 | -15.5 | -15.5 | -16.0 | -14.1 | -10.4 |
| | 9 | -15.4 | -15.4 | -15.6 | -13.7 | -10.4 |
| | 10 | -15.1 | -15.3 | -15.3 | -13.5 | -10.3 |
| | 11 | -14.8 | -15.3 | -14.7 | -13.2 | -10.2 |
| | 12 | -14.1 | -15.2 | -13.6 | -13.0 | -10.2 |
| | 13 | -13.1 | -15.0 | -12.5 | -12.7 | -10.2 |
| | 14 | -12.4 | -14.9 | -11.4 | -12.3 | -10.2 |
| | 15 | -11.6 | -14.7 | -11.1 | -11.8 | -10.2 |
| | 16 | -11.2 | -14.3 | -11.1 | -11.3 | -10.2 |
| | 17 | -11.1 | -13.6 | -11.1 | -10.8 | -9.9 |
| | 18 | -10.6 | -13.0 | -11.1 | -10.5 | -9.7 |
| | 19 | -10.4 | -12.4 | -10.9 | -10.3 | -9.4 |
| | 20 | -10.4 | -12.1 | -10.7 | -10.1 | -9.2 |
| | 21 | -10.2 | -11.9 | -10.5 | -9.9 | -9.1 |
| | 22 | -10.2 | -11.7 | -10.4 | -9.7 | -9.1 |

| | | | | | |
|---|---|---|---|---|---|
| 23 | -10.1 | -11.5 | -10.3 | -9.5 | -9.0 |
| 24 | -10.1 | -11.1 | -10.2 | -9.2 | -8.9 |
| 25 | -9.9 | -10.7 | -10.1 | -9.0 | -8.9 |
| 26 | -9.6 | -10.3 | -10.1 | -8.7 | -8.9 |
| 27 | -9.1 | -10.0 | -10.1 | -8.5 | -8.8 |
| 28 | -8.9 | -9.7 | -10.1 | -8.1 | -8.8 |
| 29 | -8.8 | -9.4 | -9.9 | -7.8 | -8.8 |
| 30 | -8.8 | -9.1 | -9.2 | -7.5 | -8.8 |
| 31 | -8.7 | -8.8 | -8.6 | -7.4 | -8.7 |
| 32 | -8.7 | -8.6 | -8.0 | -7.3 | -8.7 |
| 33 | -8.3 | -8.4 | -7.8 | -7.3 | -8.7 |
| 34 | -8.0 | -8.2 | -7.6 | -7.2 | -8.7 |
| 35 | -7.9 | -8.1 | -7.4 | -7.2 | -8.7 |
| 36 | -7.7 | -7.9 | -7.3 | -7.0 | -8.7 |
| 37 | -7.4 | -7.8 | -7.1 | -6.8 | -8.5 |
| 38 | -7.3 | -7.7 | -7.0 | -6.7 | -8.3 |
| 39 | -7.2 | -7.6 | -6.9 | -6.5 | -8.1 |
| 40 | -7.0 | -7.4 | -6.7 | -6.4 | -7.9 |
| 41 | -6.8 | -7.2 | -6.6 | -6.3 | -7.4 |
| 42 | -6.5 | -7.1 | -6.4 | -6.2 | -6.8 |
| 43 | -6.4 | -6.9 | -6.2 | -6.2 | -6.3 |
| 44 | -6.2 | -6.8 | -6.0 | -6.1 | -5.7 |
| 45 | -6.1 | -6.6 | -5.8 | -6.0 | -5.6 |
| 46 | -6.1 | -6.5 | -5.6 | -5.9 | -5.5 |
| 47 | -5.9 | -6.4 | -5.4 | -5.8 | -5.4 |
| 48 | -5.7 | -6.3 | -5.1 | -5.7 | -5.4 |
| 49 | -5.6 | -6.2 | -4.9 | -5.5 | -5.3 |
| 50 | -5.5 | -6.1 | -4.6 | -5.4 | -5.2 |
| 51 | -5.4 | -6.1 | -4.5 | -5.3 | -5.0 |
| 52 | -5.2 | -6.1 | -4.4 | -5.2 | -4.9 |
| 53 | -5.1 | -6.0 | -4.3 | -5.1 | -4.8 |
| 54 | -5.0 | -6.0 | -4.1 | -5.0 | -4.6 |
| 55 | -4.7 | -5.8 | -3.7 | -4.8 | -4.5 |
| 56 | -4.6 | -5.6 | -3.3 | -4.7 | -4.3 |
| 57 | -4.4 | -5.5 | -2.9 | -4.5 | -4.2 |
| 58 | -4.3 | -5.3 | -2.8 | -4.2 | -4.1 |
| 59 | -4.2 | -5.3 | -2.8 | -3.8 | -4.0 |
| 60 | -3.9 | -5.2 | -2.8 | -3.5 | -3.9 |
| 61 | -3.6 | -5.1 | -2.8 | -3.2 | -3.9 |
| 62 | -3.4 | -5.0 | -2.7 | -3.2 | -3.8 |
| 63 | -3.2 | -4.8 | -2.7 | -3.2 | -3.7 |
| 64 | -3.1 | -4.5 | -2.7 | -3.2 | -3.6 |
| 65 | -2.8 | -4.3 | -2.7 | -3.1 | -3.3 |
| 66 | -2.8 | -4.1 | -2.6 | -3.1 | -3.0 |

| | | | | | |
|---|---|---|---|---|---|
| 67 | -2.8 | -3.9 | -2.6 | -3.0 | -2.7 |
| 68 | -2.7 | -3.7 | -2.6 | -2.9 | -2.4 |
| 69 | -2.7 | -3.4 | -2.6 | -2.8 | -2.3 |
| 70 | -2.7 | -3.3 | -2.5 | -2.6 | -2.2 |
| 71 | -2.6 | -3.1 | -2.5 | -2.5 | -2.1 |
| 72 | -2.5 | -2.9 | -2.5 | -2.3 | -2.0 |
| 73 | -2.5 | -2.8 | -2.5 | -2.2 | -1.9 |
| 74 | -2.4 | -2.8 | -2.5 | -2.0 | -1.7 |
| 75 | -2.4 | -2.8 | -2.5 | -1.9 | -1.6 |
| 76 | -2.4 | -2.7 | -2.4 | -1.8 | -1.5 |
| 77 | -2.2 | -2.7 | -2.4 | -1.7 | -1.5 |
| 78 | -2.1 | -2.7 | -2.4 | -1.6 | -1.5 |
| 79 | -2.0 | -2.7 | -2.3 | -1.5 | -1.5 |
| 80 | -1.6 | -2.7 | -2.1 | -1.5 | -1.5 |
| 81 | -1.6 | -2.6 | -1.9 | -1.5 | -1.4 |
| 82 | -1.5 | -2.6 | -1.7 | -1.5 | -1.3 |
| 83 | -1.5 | -2.5 | -1.6 | -1.5 | -1.3 |
| 84 | -1.5 | -2.5 | -1.5 | -1.4 | -1.2 |
| 85 | -1.5 | -2.4 | -1.4 | -1.4 | -1.0 |
| 86 | -1.4 | -2.3 | -1.3 | -1.3 | -0.8 |
| 87 | -1.4 | -2.3 | -1.2 | -1.3 | -0.6 |
| 88 | -1.3 | -2.2 | -1.0 | -1.3 | -0.4 |
| 89 | -1.2 | -2.1 | -0.8 | -1.3 | -0.1 |
| 90 | -1.2 | -2.0 | -0.7 | -1.3 | 0.2 |
| 91 | -0.9 | -1.8 | -0.7 | -1.3 | 0.5 |
| 92 | -0.7 | -1.6 | -0.6 | -1.3 | 0.9 |
| 93 | -0.6 | -1.6 | -0.6 | -1.2 | 0.9 |
| 94 | -0.4 | -1.6 | -0.2 | -1.2 | 1.0 |
| 95 | 0.3 | -1.5 | 0.1 | -1.2 | 1.0 |
| 96 | 0.7 | -1.5 | 0.4 | -1.1 | 1.1 |
| 97 | 0.8 | -1.0 | 0.7 | -1.0 | 1.6 |
| 98 | 1.1 | -0.4 | 1.1 | -0.8 | 2.1 |
| 99 | 1.7 | 0.2 | 1.4 | -0.7 | 2.7 |
| 100 | 3.2 | 0.8 | 1.8 | -0.6 | 3.2 |

***Table S42:*** *Paced tapping with tones – Accuracy (vector direction %), slow tempo – Paced_metro_750_vecDirPct*

| | | | | | | |
|---|---|---|---|---|---|---|
| **Task** | Paced tapping with tones | | | | | |
| **Outcome measure** | Accuracy (vector direction %), slow tempo | | | | | |
| **Variable name** | Paced_metro_750_vecDirPct | | | | | |
| **Age and gender effects** | | | | | | |
| Age regression (slope) | | -0.025 | | | | |
| Age regression (p) | | 0.290 | | | | |
| Age group (p) | | 0.062 | | | | |
| Gender (p) | | 0.843 | | | | |
| **Group** | | All | Age 18 to 21 | Age 22 to 29 | Age 30 to 54 | Age 55 to 87 |
| N (N female) | | 104 (70) | 27 (18) | 28 (18) | 24 (14) | 25 (20) |
| **Normality** | | | | | | |
| Skewness | | -0.36 | -0.85 | 0.20 | -0.29 | -0.13 |
| Excess kurtosis | | 0.02 | -0.05 | -0.56 | -0.15 | -1.07 |
| **Scores** | | | | | | |
| Mean | | -5.3 | -6.3 | -3.8 | -3.9 | -7.0 |
| SD | | 4.7 | 5.1 | 3.8 | 5.1 | 4.3 |
| Percentiles | | | | | | |
| | 0 | -17.6 | -17.6 | -10.8 | -14.9 | -14.6 |
| | 1 | -17.4 | -17.6 | -10.5 | -14.5 | -14.5 |
| | 2 | -16.0 | -17.5 | -10.2 | -14.0 | -14.4 |
| | 3 | -14.9 | -17.5 | -9.9 | -13.6 | -14.4 |
| | 4 | -14.6 | -17.3 | -9.7 | -13.1 | -14.3 |
| | 5 | -14.1 | -17.0 | -9.4 | -12.8 | -13.9 |
| | 6 | -12.9 | -16.6 | -9.2 | -12.6 | -13.6 |
| | 7 | -12.6 | -16.3 | -9.0 | -12.4 | -13.2 |
| | 8 | -12.2 | -15.7 | -8.8 | -12.1 | -12.8 |
| | 9 | -11.8 | -14.4 | -8.6 | -11.9 | -12.6 |
| | 10 | -11.3 | -13.1 | -8.4 | -11.5 | -12.5 |
| | 11 | -11.1 | -11.8 | -8.3 | -11.1 | -12.4 |
| | 12 | -11.0 | -11.0 | -8.2 | -10.7 | -12.3 |
| | 13 | -10.8 | -11.0 | -8.2 | -10.3 | -12.2 |
| | 14 | -10.7 | -10.9 | -8.1 | -9.7 | -11.9 |
| | 15 | -10.4 | -10.8 | -8.0 | -9.1 | -11.7 |
| | 16 | -10.2 | -10.6 | -7.8 | -8.5 | -11.5 |
| | 17 | -9.8 | -10.4 | -7.5 | -7.9 | -11.3 |
| | 18 | -9.7 | -10.2 | -7.3 | -7.6 | -11.2 |
| | 19 | -9.3 | -10.0 | -7.1 | -7.4 | -11.2 |
| | 20 | -9.0 | -9.9 | -6.8 | -7.2 | -11.1 |
| | 21 | -8.7 | -9.8 | -6.6 | -7.0 | -11.1 |
| | 22 | -8.4 | -9.8 | -6.4 | -6.8 | -10.9 |

| | | | | | |
|---|---|---|---|---|---|
| 23 | -8.3 | -9.7 | -6.3 | -6.7 | -10.8 |
| 24 | -8.3 | -9.3 | -6.3 | -6.6 | -10.7 |
| 25 | -8.3 | -9.0 | -6.2 | -6.4 | -10.5 |
| 26 | -8.1 | -8.6 | -6.2 | -6.3 | -10.2 |
| 27 | -8.0 | -8.2 | -6.2 | -6.0 | -9.8 |
| 28 | -8.0 | -8.2 | -6.2 | -5.6 | -9.5 |
| 29 | -7.8 | -8.1 | -6.2 | -5.2 | -9.2 |
| 30 | -7.7 | -8.1 | -6.2 | -4.8 | -9.0 |
| 31 | -7.7 | -8.0 | -6.2 | -4.6 | -8.9 |
| 32 | -7.2 | -7.9 | -6.1 | -4.6 | -8.7 |
| 33 | -7.1 | -7.9 | -6.1 | -4.6 | -8.6 |
| 34 | -6.8 | -7.8 | -6.0 | -4.6 | -8.5 |
| 35 | -6.6 | -7.7 | -5.9 | -4.6 | -8.5 |
| 36 | -6.4 | -7.7 | -5.7 | -4.6 | -8.4 |
| 37 | -6.3 | -7.7 | -5.5 | -4.5 | -8.3 |
| 38 | -6.2 | -7.7 | -5.5 | -4.5 | -8.3 |
| 39 | -6.2 | -7.5 | -5.4 | -4.4 | -8.3 |
| 40 | -6.1 | -7.2 | -5.3 | -4.4 | -8.3 |
| 41 | -5.9 | -7.0 | -5.2 | -4.4 | -8.3 |
| 42 | -5.5 | -6.7 | -4.8 | -4.4 | -8.3 |
| 43 | -5.5 | -6.4 | -4.5 | -4.3 | -8.2 |
| 44 | -5.3 | -6.1 | -4.2 | -4.3 | -8.1 |
| 45 | -5.2 | -5.9 | -4.0 | -4.2 | -8.0 |
| 46 | -5.0 | -5.6 | -4.0 | -4.1 | -8.0 |
| 47 | -4.8 | -5.4 | -3.9 | -4.0 | -7.7 |
| 48 | -4.7 | -5.2 | -3.8 | -3.9 | -7.5 |
| 49 | -4.7 | -5.1 | -3.8 | -3.9 | -7.3 |
| 50 | -4.6 | -4.9 | -3.6 | -3.9 | -7.1 |
| 51 | -4.6 | -4.8 | -3.5 | -3.9 | -6.8 |
| 52 | -4.5 | -4.8 | -3.4 | -3.9 | -6.6 |
| 53 | -4.4 | -4.7 | -3.3 | -3.7 | -6.3 |
| 54 | -4.2 | -4.6 | -3.2 | -3.5 | -6.1 |
| 55 | -4.1 | -4.5 | -3.1 | -3.3 | -5.9 |
| 56 | -4.0 | -4.4 | -3.1 | -3.1 | -5.7 |
| 57 | -4.0 | -4.2 | -3.0 | -3.0 | -5.6 |
| 58 | -3.9 | -4.1 | -3.0 | -2.8 | -5.4 |
| 59 | -3.8 | -3.8 | -3.0 | -2.7 | -5.3 |
| 60 | -3.5 | -3.6 | -2.9 | -2.5 | -5.2 |
| 61 | -3.3 | -3.4 | -2.8 | -2.4 | -5.2 |
| 62 | -3.1 | -3.2 | -2.7 | -2.2 | -5.1 |
| 63 | -3.0 | -3.2 | -2.6 | -2.1 | -5.0 |
| 64 | -3.0 | -3.1 | -2.5 | -1.9 | -4.9 |
| 65 | -3.0 | -3.0 | -2.4 | -1.8 | -4.8 |
| 66 | -2.7 | -3.0 | -2.4 | -1.6 | -4.8 |

| | | | | | |
|---|---|---|---|---|---|
| 67 | -2.6 | -2.9 | -2.3 | -1.6 | -4.7 |
| 68 | -2.6 | -2.8 | -2.1 | -1.5 | -4.7 |
| 69 | -2.4 | -2.7 | -2.0 | -1.4 | -4.6 |
| 70 | -2.4 | -2.7 | -1.9 | -1.3 | -4.6 |
| 71 | -2.3 | -2.7 | -1.8 | -1.3 | -4.5 |
| 72 | -2.2 | -2.7 | -1.8 | -1.2 | -4.4 |
| 73 | -2.2 | -2.7 | -1.7 | -1.1 | -4.3 |
| 74 | -2.2 | -2.6 | -1.7 | -1.1 | -4.1 |
| 75 | -2.1 | -2.5 | -1.5 | -1.1 | -4.0 |
| 76 | -2.0 | -2.5 | -1.4 | -1.1 | -3.6 |
| 77 | -1.8 | -2.4 | -1.3 | -1.1 | -3.2 |
| 78 | -1.7 | -2.3 | -1.2 | -1.1 | -2.7 |
| 79 | -1.7 | -2.3 | -1.0 | -1.0 | -2.3 |
| 80 | -1.6 | -2.2 | -0.9 | -1.0 | -2.2 |
| 81 | -1.4 | -2.2 | -0.8 | -1.0 | -2.2 |
| 82 | -1.3 | -2.2 | -0.6 | -1.0 | -2.1 |
| 83 | -1.2 | -2.2 | -0.4 | -0.9 | -2.1 |
| 84 | -1.1 | -2.2 | -0.2 | -0.6 | -2.1 |
| 85 | -1.1 | -2.1 | 0.0 | -0.3 | -2.1 |
| 86 | -1.0 | -2.0 | 0.3 | 0.1 | -2.1 |
| 87 | -0.8 | -1.9 | 0.7 | 0.4 | -2.1 |
| 88 | -0.7 | -1.8 | 1.0 | 1.2 | -2.0 |
| 89 | -0.6 | -1.6 | 1.3 | 1.9 | -1.8 |
| 90 | -0.5 | -1.5 | 1.4 | 2.7 | -1.7 |
| 91 | -0.1 | -1.4 | 1.4 | 3.5 | -1.5 |
| 92 | 0.3 | -1.2 | 1.4 | 3.7 | -1.4 |
| 93 | 1.0 | -1.1 | 1.5 | 3.8 | -1.2 |
| 94 | 1.3 | -1.0 | 1.8 | 3.8 | -1.0 |
| 95 | 1.4 | -0.8 | 2.1 | 3.8 | -0.8 |
| 96 | 2.3 | -0.7 | 2.4 | 4.0 | -0.6 |
| 97 | 3.6 | -0.3 | 2.9 | 4.5 | -0.5 |
| 98 | 3.8 | 0.2 | 3.4 | 4.9 | -0.5 |
| 99 | 4.6 | 0.7 | 4.0 | 5.4 | -0.4 |
| 100 | 5.8 | 1.2 | 4.6 | 5.8 | -0.4 |

***Table S43:*** *Paced tapping with tones – Accuracy (vector direction degrees), fast tempo – Paced_metro_450_mean_vector_direction*

| | | | | | | |
|---|---|---|---|---|---|---|
| **Task** | | Paced tapping with tones | | | | |
| **Outcome measure** | | Accuracy (vector direction degrees), fast tempo | | | | |
| **Variable name** | | Paced_metro_450_mean_vector_direction | | | | |
| **Age and gender effects** | | | | | | |
| Age regression (slope) | | 0.274 | | | | |
| Age regression (p) | | 0.038 | | | | |
| Age group (p) | | 0.210 | | | | |
| Gender (p) | | 0.692 | | | | |
| **Group** | | All | Age 18 to 21 | Age 22 to 29 | Age 30 to 54 | Age 55 to 87 |
| N (N female) | | 105 (71) | 26 (17) | 28 (18) | 26 (16) | 25 (20) |
| **Normality** | | | | | | |
| Skewness | | -1.08 | -1.09 | -1.03 | -0.82 | -0.70 |
| Excess kurtosis | | 1.05 | 0.19 | 0.82 | 0.84 | -0.37 |
| **Scores** | | | | | | |
| Mean | | -31.5 | -37.3 | -35.6 | -31.8 | -20.7 |
| SD | | 29.1 | 31.9 | 33.2 | 25.2 | 23.1 |
| Percentiles | | | | | | |
| | 0 | -129.1 | -113.1 | -129.1 | -94.3 | -77.0 |
| | 1 | -112.8 | -111.1 | -121.9 | -93.9 | -72.9 |
| | 2 | -105.0 | -109.2 | -114.7 | -93.6 | -68.8 |
| | 3 | -101.5 | -107.2 | -107.6 | -93.3 | -64.7 |
| | 4 | -94.2 | -105.2 | -101.6 | -92.9 | -60.6 |
| | 5 | -93.8 | -102.4 | -98.2 | -85.3 | -58.1 |
| | 6 | -92.3 | -99.6 | -94.9 | -77.7 | -55.9 |
| | 7 | -89.9 | -96.8 | -91.6 | -70.1 | -53.7 |
| | 8 | -85.0 | -94.0 | -86.7 | -62.5 | -51.5 |
| | 9 | -73.6 | -92.7 | -80.6 | -60.3 | -50.5 |
| | 10 | -65.6 | -91.3 | -74.5 | -58.1 | -50.0 |
| | 11 | -62.4 | -90.0 | -68.4 | -55.9 | -49.6 |
| | 12 | -61.3 | -88.7 | -65.9 | -53.7 | -49.1 |
| | 13 | -60.2 | -82.1 | -64.0 | -51.0 | -48.7 |
| | 14 | -59.9 | -75.4 | -62.0 | -48.4 | -48.2 |
| | 15 | -56.6 | -68.8 | -60.4 | -45.7 | -47.7 |
| | 16 | -54.0 | -62.2 | -60.3 | -43.0 | -47.2 |
| | 17 | -53.7 | -60.2 | -60.1 | -42.8 | -46.1 |
| | 18 | -52.2 | -58.3 | -60.0 | -42.5 | -43.7 |
| | 19 | -51.4 | -56.3 | -59.1 | -42.3 | -41.4 |
| | 20 | -50.9 | -54.4 | -57.5 | -42.1 | -39.0 |
| | 21 | -50.6 | -53.7 | -55.8 | -42.0 | -37.0 |
| | 22 | -49.1 | -53.0 | -54.1 | -41.8 | -36.8 |

| | | | | | |
|---|---|---|---|---|---|
| 23 | -47.1 | -52.3 | -53.1 | -41.7 | -36.6 |
| 24 | -46.2 | -51.6 | -52.2 | -41.6 | -36.4 |
| 25 | -44.1 | -51.5 | -51.4 | -41.5 | -36.2 |
| 26 | -43.0 | -51.5 | -50.5 | -41.5 | -35.7 |
| 27 | -42.1 | -51.4 | -49.3 | -41.4 | -35.1 |
| 28 | -42.0 | -51.3 | -48.1 | -41.3 | -34.5 |
| 29 | -41.6 | -47.7 | -46.9 | -40.5 | -33.9 |
| 30 | -40.7 | -44.1 | -45.9 | -39.8 | -32.9 |
| 31 | -38.0 | -40.4 | -45.4 | -39.1 | -31.8 |
| 32 | -37.0 | -36.8 | -44.9 | -38.3 | -30.7 |
| 33 | -36.9 | -36.0 | -44.3 | -38.0 | -29.6 |
| 34 | -36.6 | -35.2 | -43.8 | -37.6 | -28.7 |
| 35 | -36.0 | -34.4 | -43.2 | -37.3 | -27.9 |
| 36 | -35.1 | -33.6 | -42.6 | -36.9 | -27.1 |
| 37 | -34.4 | -33.1 | -42.1 | -36.6 | -26.4 |
| 38 | -34.1 | -32.6 | -39.7 | -36.2 | -25.1 |
| 39 | -33.7 | -32.2 | -37.3 | -35.9 | -23.3 |
| 40 | -33.3 | -31.7 | -34.9 | -35.5 | -21.5 |
| 41 | -32.9 | -31.2 | -33.1 | -35.3 | -19.7 |
| 42 | -32.1 | -30.8 | -33.0 | -35.0 | -18.3 |
| 43 | -31.3 | -30.3 | -33.0 | -34.7 | -17.6 |
| 44 | -30.9 | -29.8 | -32.9 | -34.5 | -16.9 |
| 45 | -30.1 | -29.4 | -32.6 | -34.5 | -16.3 |
| 46 | -29.9 | -29.0 | -32.1 | -34.4 | -15.7 |
| 47 | -29.3 | -28.7 | -31.7 | -34.4 | -15.5 |
| 48 | -28.6 | -28.3 | -31.2 | -34.4 | -15.4 |
| 49 | -28.3 | -27.7 | -30.9 | -33.5 | -15.2 |
| 50 | -26.0 | -27.1 | -30.5 | -32.6 | -15.1 |
| 51 | -26.0 | -26.6 | -30.2 | -31.7 | -15.0 |
| 52 | -26.0 | -26.0 | -29.9 | -30.8 | -15.0 |
| 53 | -25.5 | -25.6 | -29.5 | -29.5 | -14.9 |
| 54 | -24.5 | -25.2 | -29.2 | -28.2 | -14.8 |
| 55 | -24.2 | -24.8 | -28.8 | -26.9 | -14.3 |
| 56 | -23.3 | -24.4 | -28.3 | -25.6 | -13.8 |
| 57 | -22.5 | -24.1 | -27.6 | -25.3 | -13.2 |
| 58 | -21.0 | -23.7 | -26.9 | -25.1 | -12.7 |
| 59 | -20.5 | -23.3 | -26.2 | -24.8 | -12.0 |
| 60 | -20.0 | -22.9 | -24.2 | -24.5 | -11.3 |
| 61 | -19.2 | -22.3 | -21.6 | -24.2 | -10.6 |
| 62 | -18.4 | -21.6 | -19.1 | -23.9 | -9.8 |
| 63 | -17.7 | -20.9 | -16.6 | -23.6 | -9.3 |
| 64 | -17.3 | -20.2 | -15.8 | -23.4 | -8.9 |
| 65 | -16.9 | -20.1 | -14.9 | -22.8 | -8.6 |
| 66 | -16.2 | -20.0 | -14.1 | -22.3 | -8.2 |

| | | | | | |
|---|---|---|---|---|---|
| 67 | -15.8 | -19.9 | -13.4 | -21.8 | -7.6 |
| 68 | -15.3 | -19.7 | -13.3 | -21.2 | -6.4 |
| 69 | -14.9 | -19.4 | -13.1 | -21.1 | -5.2 |
| 70 | -13.7 | -19.0 | -12.9 | -20.9 | -4.0 |
| 71 | -13.0 | -18.6 | -12.8 | -20.8 | -2.9 |
| 72 | -12.5 | -18.2 | -12.7 | -20.6 | -2.2 |
| 73 | -12.4 | -17.9 | -12.5 | -19.8 | -1.6 |
| 74 | -12.3 | -17.7 | -12.4 | -19.0 | -0.9 |
| 75 | -11.2 | -17.5 | -12.4 | -18.1 | -0.3 |
| 76 | -10.6 | -17.3 | -12.4 | -17.3 | 0.1 |
| 77 | -10.2 | -16.9 | -12.4 | -15.8 | 0.5 |
| 78 | -9.5 | -16.6 | -12.2 | -14.2 | 0.9 |
| 79 | -9.2 | -16.3 | -11.8 | -12.7 | 1.3 |
| 80 | -7.8 | -15.9 | -11.3 | -11.2 | 1.5 |
| 81 | -6.7 | -14.5 | -10.8 | -10.2 | 1.5 |
| 82 | -4.9 | -13.1 | -9.7 | -9.2 | 1.6 |
| 83 | -4.2 | -11.7 | -8.0 | -8.2 | 1.7 |
| 84 | -3.9 | -10.3 | -6.3 | -7.2 | 1.9 |
| 85 | -3.2 | -10.1 | -4.6 | -6.4 | 2.0 |
| 86 | -2.5 | -9.9 | -3.8 | -5.7 | 2.2 |
| 87 | -1.8 | -9.7 | -3.2 | -4.9 | 2.3 |
| 88 | -1.3 | -9.6 | -2.5 | -4.1 | 2.5 |
| 89 | -0.6 | -8.4 | -2.0 | -3.3 | 2.7 |
| 90 | 0.3 | -7.3 | -1.9 | -2.5 | 2.9 |
| 91 | 1.2 | -6.2 | -1.8 | -1.7 | 3.1 |
| 92 | 1.6 | -5.1 | -1.8 | -0.9 | 3.6 |
| 93 | 2.2 | -4.7 | -0.8 | -0.5 | 4.7 |
| 94 | 3.1 | -4.3 | 1.5 | -0.1 | 5.7 |
| 95 | 3.7 | -3.8 | 3.8 | 0.4 | 6.8 |
| 96 | 6.3 | -3.4 | 6.1 | 0.8 | 7.7 |
| 97 | 7.6 | -1.6 | 7.6 | 3.4 | 7.9 |
| 98 | 8.6 | 0.2 | 8.7 | 6.1 | 8.2 |
| 99 | 10.8 | 2.0 | 9.8 | 8.8 | 8.4 |
| 100 | 11.4 | 3.8 | 10.9 | 11.4 | 8.6 |

***Table S44:*** *Paced tapping with tones – Accuracy (vector direction degrees), medium tempo – Paced_metro_600_mean_vector_direction*

| | | | | | | |
|---|---|---|---|---|---|---|
| **Task** | | Paced tapping with tones | | | | |
| **Outcome measure** | | Accuracy (vector direction degrees), medium tempo | | | | |
| **Variable name** | | Paced_metro_600_mean_vector_direction | | | | |
| **Age and gender effects** | | | | | | |
| Age regression (slope) | | 0.083 | | | | |
| Age regression (p) | | 0.322 | | | | |
| Age group (p) | | 0.781 | | | | |
| Gender (p) | | 0.330 | | | | |
| **Group** | | All | Age 18 to 21 | Age 22 to 29 | Age 30 to 54 | Age 55 to 87 |
| N (N female) | | 107 (73) | 27 (18) | 29 (19) | 25 (15) | 26 (21) |
| **Normality** | | | | | | |
| Skewness | | -1.10 | -0.61 | -0.82 | -1.46 | -1.22 |
| Excess kurtosis | | 1.17 | -0.66 | -0.15 | 1.73 | 1.98 |
| **Scores** | | | | | | |
| Mean | | -24.3 | -26.3 | -22.9 | -26.2 | -21.9 |
| SD | | 21.2 | 19.0 | 20.5 | 24.4 | 22.0 |
| Percentiles | | | | | | |
| | 0 | -99.0 | -68.6 | -74.4 | -99.0 | -89.8 |
| | 1 | -89.1 | -65.4 | -70.1 | -94.0 | -83.9 |
| | 2 | -77.6 | -62.1 | -65.9 | -88.9 | -78.1 |
| | 3 | -73.4 | -58.9 | -61.6 | -83.9 | -72.3 |
| | 4 | -68.1 | -56.1 | -59.1 | -78.9 | -66.4 |
| | 5 | -64.2 | -56.0 | -58.9 | -73.8 | -59.2 |
| | 6 | -59.0 | -55.9 | -58.8 | -68.7 | -52.0 |
| | 7 | -57.9 | -55.8 | -58.7 | -63.6 | -44.8 |
| | 8 | -56.5 | -55.6 | -57.6 | -58.5 | -37.6 |
| | 9 | -55.9 | -55.4 | -56.3 | -55.7 | -37.3 |
| | 10 | -55.2 | -55.2 | -54.9 | -54.0 | -37.1 |
| | 11 | -54.3 | -54.9 | -52.9 | -52.3 | -36.8 |
| | 12 | -53.1 | -54.5 | -49.0 | -50.5 | -36.6 |
| | 13 | -50.4 | -54.0 | -45.1 | -49.2 | -36.6 |
| | 14 | -46.4 | -53.5 | -41.2 | -48.3 | -36.5 |
| | 15 | -44.3 | -52.9 | -40.0 | -47.4 | -36.5 |
| | 16 | -41.3 | -51.4 | -40.0 | -46.4 | -36.5 |
| | 17 | -40.1 | -49.1 | -40.0 | -45.2 | -35.7 |
| | 18 | -39.9 | -46.9 | -39.9 | -43.3 | -34.9 |
| | 19 | -38.1 | -44.6 | -39.2 | -41.5 | -34.0 |
| | 20 | -37.6 | -43.5 | -38.5 | -39.7 | -33.2 |
| | 21 | -37.3 | -42.8 | -37.9 | -38.0 | -32.9 |
| | 22 | -36.6 | -42.0 | -37.4 | -37.3 | -32.6 |

| | | | | | |
|---|---|---|---|---|---|
| 23 | -36.5 | -41.2 | -37.1 | -36.7 | -32.3 |
| 24 | -36.5 | -39.9 | -36.8 | -36.0 | -32.0 |
| 25 | -36.2 | -38.6 | -36.5 | -35.3 | -32.0 |
| 26 | -35.6 | -37.2 | -36.5 | -34.4 | -31.9 |
| 27 | -34.0 | -35.8 | -36.5 | -33.4 | -31.8 |
| 28 | -32.4 | -34.8 | -36.5 | -32.5 | -31.7 |
| 29 | -31.9 | -33.7 | -35.5 | -31.6 | -31.6 |
| 30 | -31.8 | -32.7 | -33.3 | -30.5 | -31.5 |
| 31 | -31.5 | -31.7 | -31.0 | -29.3 | -31.5 |
| 32 | -31.4 | -31.0 | -28.8 | -28.2 | -31.4 |
| 33 | -31.3 | -30.3 | -28.0 | -27.0 | -31.4 |
| 34 | -29.2 | -29.6 | -27.4 | -26.5 | -31.3 |
| 35 | -28.5 | -29.0 | -26.8 | -26.4 | -31.3 |
| 36 | -28.3 | -28.6 | -26.2 | -26.2 | -31.3 |
| 37 | -27.4 | -28.2 | -25.7 | -26.0 | -30.6 |
| 38 | -26.6 | -27.8 | -25.2 | -25.6 | -29.9 |
| 39 | -26.2 | -27.2 | -24.7 | -25.0 | -29.2 |
| 40 | -25.7 | -26.6 | -24.2 | -24.4 | -28.6 |
| 41 | -25.0 | -26.0 | -23.6 | -23.8 | -26.5 |
| 42 | -24.0 | -25.5 | -23.0 | -23.3 | -24.5 |
| 43 | -23.3 | -24.9 | -22.4 | -23.0 | -22.5 |
| 44 | -22.8 | -24.4 | -21.7 | -22.6 | -20.5 |
| 45 | -22.2 | -23.9 | -21.0 | -22.3 | -20.2 |
| 46 | -22.0 | -23.4 | -20.3 | -22.0 | -19.9 |
| 47 | -21.7 | -23.0 | -19.4 | -21.6 | -19.6 |
| 48 | -20.6 | -22.7 | -18.5 | -21.2 | -19.3 |
| 49 | -20.5 | -22.3 | -17.5 | -20.8 | -18.9 |
| 50 | -20.0 | -22.0 | -16.5 | -20.5 | -18.6 |
| 51 | -19.3 | -21.9 | -16.2 | -20.0 | -18.2 |
| 52 | -19.2 | -21.8 | -15.8 | -19.6 | -17.8 |
| 53 | -18.6 | -21.7 | -15.5 | -19.1 | -17.2 |
| 54 | -18.2 | -21.5 | -14.7 | -18.7 | -16.7 |
| 55 | -17.5 | -20.9 | -13.3 | -18.3 | -16.2 |
| 56 | -16.7 | -20.3 | -11.9 | -17.8 | -15.6 |
| 57 | -16.1 | -19.7 | -10.5 | -17.4 | -15.2 |
| 58 | -15.5 | -19.2 | -10.2 | -17.0 | -14.9 |
| 59 | -15.3 | -18.9 | -10.1 | -16.0 | -14.5 |
| 60 | -14.6 | -18.7 | -10.1 | -14.7 | -14.1 |
| 61 | -13.5 | -18.4 | -10.0 | -13.5 | -13.9 |
| 62 | -12.5 | -17.9 | -9.9 | -12.2 | -13.6 |
| 63 | -11.7 | -17.2 | -9.8 | -11.5 | -13.4 |
| 64 | -11.3 | -16.4 | -9.6 | -11.5 | -13.1 |
| 65 | -10.4 | -15.6 | -9.6 | -11.4 | -12.0 |
| 66 | -10.0 | -14.8 | -9.5 | -11.3 | -10.9 |

| | | | | | |
|---|---|---|---|---|---|
| 67 | -9.9 | -14.0 | -9.4 | -11.2 | -9.8 |
| 68 | -9.9 | -13.2 | -9.4 | -10.8 | -8.7 |
| 69 | -9.7 | -12.4 | -9.3 | -10.4 | -8.3 |
| 70 | -9.6 | -11.8 | -9.1 | -10.1 | -7.9 |
| 71 | -9.5 | -11.2 | -9.0 | -9.7 | -7.5 |
| 72 | -9.3 | -10.6 | -8.9 | -9.1 | -7.1 |
| 73 | -8.9 | -10.0 | -8.9 | -8.5 | -6.7 |
| 74 | -8.8 | -9.9 | -8.9 | -8.0 | -6.3 |
| 75 | -8.7 | -9.9 | -8.9 | -7.4 | -5.9 |
| 76 | -8.6 | -9.9 | -8.8 | -7.0 | -5.4 |
| 77 | -8.2 | -9.9 | -8.7 | -6.5 | -5.4 |
| 78 | -7.6 | -9.8 | -8.6 | -6.1 | -5.4 |
| 79 | -7.2 | -9.7 | -8.3 | -5.7 | -5.3 |
| 80 | -6.2 | -9.6 | -7.5 | -5.5 | -5.3 |
| 81 | -5.8 | -9.5 | -6.8 | -5.5 | -5.0 |
| 82 | -5.6 | -9.3 | -6.0 | -5.4 | -4.8 |
| 83 | -5.5 | -9.1 | -5.7 | -5.3 | -4.5 |
| 84 | -5.4 | -8.9 | -5.5 | -5.2 | -4.3 |
| 85 | -5.3 | -8.7 | -5.2 | -5.0 | -3.6 |
| 86 | -5.2 | -8.4 | -4.9 | -4.9 | -2.9 |
| 87 | -5.0 | -8.2 | -4.2 | -4.7 | -2.1 |
| 88 | -4.6 | -8.0 | -3.5 | -4.6 | -1.4 |
| 89 | -4.5 | -7.6 | -2.8 | -4.6 | -0.3 |
| 90 | -4.3 | -7.1 | -2.6 | -4.6 | 0.8 |
| 91 | -3.5 | -6.5 | -2.4 | -4.6 | 1.9 |
| 92 | -2.4 | -5.9 | -2.3 | -4.5 | 3.1 |
| 93 | -2.1 | -5.7 | -2.0 | -4.5 | 3.3 |
| 94 | -1.6 | -5.6 | -0.9 | -4.4 | 3.5 |
| 95 | 0.9 | -5.6 | 0.3 | -4.3 | 3.7 |
| 96 | 2.5 | -5.5 | 1.4 | -4.2 | 3.9 |
| 97 | 3.0 | -3.7 | 2.6 | -3.6 | 5.8 |
| 98 | 3.8 | -1.6 | 3.9 | -3.1 | 7.7 |
| 99 | 6.2 | 0.6 | 5.1 | -2.5 | 9.5 |
| 100 | 11.4 | 2.7 | 6.3 | -2.0 | 11.4 |

***Table S45:*** *Paced tapping with tones – Accuracy (vector direction degrees), slow tempo – Paced_metro_750_mean_vector_direction*

| **Task** | Paced tapping with tones | | | | |
|---|---|---|---|---|---|
| **Outcome measure** | Accuracy (vector direction degrees), slow tempo | | | | |
| **Variable name** | Paced_metro_750_mean_vector_direction | | | | |
| **Age and gender effects** | | | | | |
| Age regression (slope) | -0.092 | | | | |
| Age regression (p) | 0.295 | | | | |
| Age group (p) | 0.094 | | | | |
| Gender (p) | 0.928 | | | | |
| **Group** | All | Age 18 to 21 | Age 22 to 29 | Age 30 to 54 | Age 55 to 87 |
| N (N female) | 105 (71) | 27 (18) | 28 (18) | 25 (15) | 25 (20) |
| **Normality** | | | | | |
| Skewness | -1.47 | -0.85 | 0.20 | -2.00 | -0.13 |
| Excess kurtosis | 5.69 | -0.05 | -0.56 | 5.50 | -1.07 |
| **Scores** | | | | | |
| Mean | -19.9 | -22.8 | -13.7 | -18.6 | -25.1 |
| SD | 19.8 | 18.2 | 13.8 | 28.3 | 15.5 |
| Percentiles | | | | | |
| 0 | -122.7 | -63.5 | -38.9 | -122.7 | -52.6 |
| 1 | -63.5 | -63.3 | -37.8 | -106.2 | -52.3 |
| 2 | -62.3 | -63.1 | -36.8 | -89.6 | -52.0 |
| 3 | -57.3 | -62.8 | -35.7 | -73.1 | -51.7 |
| 4 | -53.6 | -62.5 | -34.8 | -56.5 | -51.4 |
| 5 | -52.4 | -61.2 | -33.9 | -52.3 | -50.2 |
| 6 | -50.2 | -59.9 | -33.1 | -50.6 | -48.8 |
| 7 | -46.4 | -58.7 | -32.2 | -48.9 | -47.4 |
| 8 | -45.2 | -56.4 | -31.5 | -47.3 | -46.1 |
| 9 | -43.8 | -51.7 | -31.0 | -46.1 | -45.4 |
| 10 | -42.2 | -47.0 | -30.4 | -45.3 | -45.1 |
| 11 | -40.4 | -42.4 | -29.8 | -44.4 | -44.7 |
| 12 | -39.9 | -39.7 | -29.6 | -43.5 | -44.4 |
| 13 | -39.4 | -39.4 | -29.4 | -42.4 | -43.8 |
| 14 | -38.8 | -39.1 | -29.2 | -41.0 | -43.0 |
| 15 | -38.2 | -38.8 | -28.9 | -39.5 | -42.2 |
| 16 | -37.4 | -38.3 | -28.0 | -38.1 | -41.3 |
| 17 | -36.3 | -37.5 | -27.1 | -36.4 | -40.7 |
| 18 | -35.3 | -36.8 | -26.2 | -34.1 | -40.5 |
| 19 | -34.9 | -36.1 | -25.4 | -31.8 | -40.3 |
| 20 | -33.2 | -35.7 | -24.6 | -29.5 | -40.1 |
| 21 | -32.0 | -35.4 | -23.8 | -27.5 | -39.9 |
| 22 | -30.9 | -35.1 | -23.1 | -26.8 | -39.4 |

| | | | | | |
|---|---|---|---|---|---|
| 23 | -29.9 | -34.8 | -22.8 | -26.1 | -38.9 |
| 24 | -29.8 | -33.6 | -22.6 | -25.3 | -38.4 |
| 25 | -29.7 | -32.3 | -22.5 | -24.6 | -37.9 |
| 26 | -29.7 | -30.9 | -22.3 | -24.2 | -36.7 |
| 27 | -29.0 | -29.7 | -22.3 | -23.7 | -35.4 |
| 28 | -28.9 | -29.5 | -22.3 | -23.3 | -34.2 |
| 29 | -28.6 | -29.3 | -22.3 | -22.8 | -33.0 |
| 30 | -27.8 | -29.1 | -22.3 | -21.5 | -32.4 |
| 31 | -27.6 | -28.9 | -22.2 | -20.1 | -31.9 |
| 32 | -27.1 | -28.6 | -22.1 | -18.7 | -31.4 |
| 33 | -25.6 | -28.3 | -22.0 | -17.2 | -31.0 |
| 34 | -25.1 | -28.0 | -21.6 | -16.7 | -30.7 |
| 35 | -24.3 | -27.9 | -21.1 | -16.7 | -30.4 |
| 36 | -23.4 | -27.8 | -20.5 | -16.7 | -30.2 |
| 37 | -22.8 | -27.7 | -19.9 | -16.7 | -30.0 |
| 38 | -22.5 | -27.7 | -19.6 | -16.6 | -29.8 |
| 39 | -22.3 | -27.1 | -19.4 | -16.4 | -29.8 |
| 40 | -22.1 | -26.1 | -19.1 | -16.2 | -29.8 |
| 41 | -21.8 | -25.1 | -18.6 | -16.0 | -29.8 |
| 42 | -20.5 | -24.1 | -17.4 | -15.8 | -29.7 |
| 43 | -19.9 | -23.1 | -16.3 | -15.8 | -29.5 |
| 44 | -19.5 | -22.1 | -15.2 | -15.7 | -29.2 |
| 45 | -19.0 | -21.1 | -14.5 | -15.6 | -29.0 |
| 46 | -18.2 | -20.1 | -14.3 | -15.5 | -28.6 |
| 47 | -17.7 | -19.4 | -14.1 | -15.2 | -27.8 |
| 48 | -17.1 | -18.8 | -13.9 | -14.8 | -27.0 |
| 49 | -16.8 | -18.2 | -13.5 | -14.5 | -26.2 |
| 50 | -16.7 | -17.6 | -13.1 | -14.1 | -25.4 |
| 51 | -16.7 | -17.4 | -12.7 | -14.1 | -24.5 |
| 52 | -16.3 | -17.2 | -12.4 | -14.0 | -23.6 |
| 53 | -15.8 | -17.0 | -12.0 | -14.0 | -22.7 |
| 54 | -15.5 | -16.7 | -11.6 | -13.9 | -21.8 |
| 55 | -14.9 | -16.2 | -11.3 | -13.3 | -21.2 |
| 56 | -14.6 | -15.8 | -11.0 | -12.6 | -20.7 |
| 57 | -14.4 | -15.3 | -11.0 | -11.9 | -20.1 |
| 58 | -14.1 | -14.7 | -10.9 | -11.2 | -19.6 |
| 59 | -13.9 | -13.8 | -10.8 | -10.6 | -19.2 |
| 60 | -13.3 | -13.0 | -10.5 | -10.0 | -18.9 |
| 61 | -12.1 | -12.1 | -10.2 | -9.4 | -18.6 |
| 62 | -11.4 | -11.6 | -9.8 | -8.9 | -18.3 |
| 63 | -11.0 | -11.4 | -9.4 | -8.3 | -18.0 |
| 64 | -10.9 | -11.2 | -9.1 | -7.7 | -17.7 |
| 65 | -10.8 | -11.0 | -8.8 | -7.2 | -17.5 |
| 66 | -10.1 | -10.7 | -8.5 | -6.6 | -17.2 |

| | | | | | |
|---|---|---|---|---|---|
| 67 | -9.6 | -10.4 | -8.1 | -6.1 | -16.9 |
| 68 | -9.5 | -10.1 | -7.7 | -5.7 | -16.8 |
| 69 | -8.8 | -9.8 | -7.2 | -5.4 | -16.6 |
| 70 | -8.6 | -9.7 | -6.8 | -5.1 | -16.4 |
| 71 | -8.3 | -9.7 | -6.5 | -4.8 | -16.2 |
| 72 | -8.1 | -9.6 | -6.3 | -4.6 | -15.8 |
| 73 | -7.9 | -9.5 | -6.2 | -4.4 | -15.4 |
| 74 | -7.8 | -9.3 | -6.0 | -4.2 | -14.9 |
| 75 | -7.5 | -9.1 | -5.6 | -4.0 | -14.5 |
| 76 | -7.4 | -8.8 | -5.1 | -3.9 | -13.0 |
| 77 | -6.6 | -8.6 | -4.7 | -3.9 | -11.4 |
| 78 | -6.2 | -8.4 | -4.3 | -3.8 | -9.9 |
| 79 | -6.1 | -8.2 | -3.8 | -3.8 | -8.3 |
| 80 | -5.8 | -8.1 | -3.3 | -3.8 | -8.0 |
| 81 | -5.1 | -7.9 | -2.7 | -3.7 | -7.8 |
| 82 | -4.7 | -7.9 | -2.1 | -3.6 | -7.7 |
| 83 | -4.3 | -7.9 | -1.4 | -3.6 | -7.5 |
| 84 | -4.2 | -7.9 | -0.7 | -2.8 | -7.5 |
| 85 | -3.9 | -7.7 | 0.1 | -1.6 | -7.5 |
| 86 | -3.7 | -7.2 | 1.2 | -0.5 | -7.5 |
| 87 | -3.0 | -6.8 | 2.5 | 0.7 | -7.4 |
| 88 | -2.5 | -6.3 | 3.7 | 2.7 | -7.2 |
| 89 | -2.3 | -5.9 | 4.8 | 5.6 | -6.6 |
| 90 | -1.7 | -5.4 | 4.9 | 8.5 | -6.0 |
| 91 | -0.4 | -4.9 | 4.9 | 11.4 | -5.5 |
| 92 | 0.9 | -4.4 | 5.0 | 13.4 | -4.9 |
| 93 | 3.4 | -4.0 | 5.5 | 13.5 | -4.2 |
| 94 | 4.7 | -3.5 | 6.5 | 13.6 | -3.4 |
| 95 | 5.0 | -3.0 | 7.5 | 13.7 | -2.7 |
| 96 | 8.2 | -2.6 | 8.5 | 14.1 | -2.1 |
| 97 | 12.8 | -1.0 | 10.3 | 15.8 | -1.9 |
| 98 | 13.8 | 0.8 | 12.4 | 17.5 | -1.8 |
| 99 | 16.4 | 2.5 | 14.5 | 19.2 | -1.6 |
| 100 | 21.0 | 4.3 | 16.6 | 21.0 | -1.4 |

***Table S46:*** *Paced tapping with music – Variability (CV of ITI), music 1 – Paced_music_badine_mean_CV_iti*

| | | | | | | |
|---|---|---|---|---|---|---|
| **Task** | | Paced tapping with music | | | | |
| **Outcome measure** | | Variability (CV of ITI), music 1 | | | | |
| **Variable name** | | Paced_music_badine_mean_CV_iti | | | | |
| **Age and gender effects** | | | | | | |
| Age regression (slope) | | 0.000 | | | | |
| Age regression (p) | | 0.777 | | | | |
| Age group (p) | | 0.811 | | | | |
| Gender (p) | | 0.832 | | | | |
| **Group** | | All | Age 18 to 21 | Age 22 to 29 | Age 30 to 54 | Age 55 to 87 |
| N (N female) | | 106 (72) | 27 (18) | 29 (19) | 24 (14) | 26 (21) |
| **Normality** | | | | | | |
| Skewness | | 3.34 | 0.69 | 0.19 | 2.78 | 1.38 |
| Excess kurtosis | | 17.42 | -0.75 | -1.01 | 7.30 | 2.03 |
| **Scores** | | | | | | |
| Mean | | 0.055 | 0.054 | 0.055 | 0.059 | 0.052 |
| SD | | 0.026 | 0.018 | 0.018 | 0.043 | 0.017 |
| Percentiles | | | | | | |
| | 0 | 0.028 | 0.028 | 0.028 | 0.030 | 0.029 |
| | 1 | 0.028 | 0.028 | 0.029 | 0.031 | 0.030 |
| | 2 | 0.029 | 0.028 | 0.029 | 0.031 | 0.030 |
| | 3 | 0.029 | 0.028 | 0.030 | 0.032 | 0.031 |
| | 4 | 0.030 | 0.029 | 0.030 | 0.032 | 0.032 |
| | 5 | 0.031 | 0.031 | 0.031 | 0.033 | 0.032 |
| | 6 | 0.031 | 0.033 | 0.031 | 0.033 | 0.032 |
| | 7 | 0.032 | 0.036 | 0.031 | 0.034 | 0.033 |
| | 8 | 0.033 | 0.037 | 0.032 | 0.034 | 0.033 |
| | 9 | 0.033 | 0.037 | 0.032 | 0.035 | 0.034 |
| | 10 | 0.033 | 0.037 | 0.033 | 0.035 | 0.036 |
| | 11 | 0.034 | 0.037 | 0.033 | 0.036 | 0.037 |
| | 12 | 0.034 | 0.038 | 0.033 | 0.036 | 0.039 |
| | 13 | 0.035 | 0.038 | 0.034 | 0.037 | 0.039 |
| | 14 | 0.036 | 0.039 | 0.034 | 0.037 | 0.039 |
| | 15 | 0.037 | 0.040 | 0.034 | 0.037 | 0.039 |
| | 16 | 0.037 | 0.040 | 0.034 | 0.037 | 0.039 |
| | 17 | 0.037 | 0.040 | 0.034 | 0.037 | 0.039 |
| | 18 | 0.037 | 0.040 | 0.034 | 0.037 | 0.039 |
| | 19 | 0.038 | 0.040 | 0.034 | 0.038 | 0.039 |
| | 20 | 0.038 | 0.040 | 0.035 | 0.038 | 0.039 |
| | 21 | 0.039 | 0.040 | 0.035 | 0.038 | 0.039 |

| | | | | | |
|---|---|---|---|---|---|
| 22 | 0.039 | 0.040 | 0.035 | 0.038 | 0.040 |
| 23 | 0.039 | 0.040 | 0.036 | 0.038 | 0.040 |
| 24 | 0.039 | 0.040 | 0.037 | 0.039 | 0.040 |
| 25 | 0.039 | 0.041 | 0.037 | 0.039 | 0.040 |
| 26 | 0.040 | 0.042 | 0.038 | 0.039 | 0.040 |
| 27 | 0.040 | 0.042 | 0.038 | 0.039 | 0.040 |
| 28 | 0.040 | 0.042 | 0.038 | 0.039 | 0.040 |
| 29 | 0.040 | 0.042 | 0.039 | 0.039 | 0.041 |
| 30 | 0.040 | 0.042 | 0.040 | 0.039 | 0.041 |
| 31 | 0.040 | 0.043 | 0.042 | 0.039 | 0.042 |
| 32 | 0.041 | 0.043 | 0.044 | 0.040 | 0.043 |
| 33 | 0.042 | 0.043 | 0.044 | 0.040 | 0.043 |
| 34 | 0.042 | 0.043 | 0.045 | 0.040 | 0.043 |
| 35 | 0.043 | 0.043 | 0.046 | 0.040 | 0.043 |
| 36 | 0.043 | 0.043 | 0.047 | 0.040 | 0.043 |
| 37 | 0.043 | 0.044 | 0.047 | 0.040 | 0.043 |
| 38 | 0.043 | 0.044 | 0.048 | 0.040 | 0.044 |
| 39 | 0.044 | 0.044 | 0.049 | 0.040 | 0.044 |
| 40 | 0.044 | 0.045 | 0.050 | 0.041 | 0.044 |
| 41 | 0.044 | 0.045 | 0.050 | 0.041 | 0.045 |
| 42 | 0.045 | 0.046 | 0.050 | 0.041 | 0.045 |
| 43 | 0.046 | 0.046 | 0.050 | 0.042 | 0.045 |
| 44 | 0.046 | 0.046 | 0.051 | 0.042 | 0.045 |
| 45 | 0.046 | 0.046 | 0.052 | 0.042 | 0.045 |
| 46 | 0.046 | 0.046 | 0.053 | 0.042 | 0.045 |
| 47 | 0.046 | 0.047 | 0.053 | 0.043 | 0.045 |
| 48 | 0.047 | 0.047 | 0.054 | 0.043 | 0.046 |
| 49 | 0.047 | 0.047 | 0.054 | 0.044 | 0.046 |
| 50 | 0.047 | 0.047 | 0.054 | 0.045 | 0.046 |
| 51 | 0.048 | 0.048 | 0.055 | 0.046 | 0.046 |
| 52 | 0.048 | 0.048 | 0.056 | 0.046 | 0.046 |
| 53 | 0.048 | 0.048 | 0.058 | 0.047 | 0.047 |
| 54 | 0.049 | 0.048 | 0.058 | 0.047 | 0.047 |
| 55 | 0.050 | 0.048 | 0.059 | 0.047 | 0.047 |
| 56 | 0.050 | 0.048 | 0.059 | 0.047 | 0.047 |
| 57 | 0.050 | 0.048 | 0.060 | 0.047 | 0.049 |
| 58 | 0.052 | 0.049 | 0.060 | 0.047 | 0.050 |
| 59 | 0.053 | 0.050 | 0.060 | 0.048 | 0.052 |
| 60 | 0.053 | 0.051 | 0.060 | 0.048 | 0.053 |
| 61 | 0.054 | 0.052 | 0.061 | 0.048 | 0.053 |
| 62 | 0.054 | 0.053 | 0.062 | 0.049 | 0.054 |
| 63 | 0.054 | 0.053 | 0.062 | 0.049 | 0.054 |
| 64 | 0.054 | 0.054 | 0.063 | 0.050 | 0.054 |

| | | | | | |
|---|---|---|---|---|---|
| 65 | 0.055 | 0.054 | 0.064 | 0.050 | 0.054 |
| 66 | 0.056 | 0.054 | 0.064 | 0.050 | 0.054 |
| 67 | 0.059 | 0.055 | 0.064 | 0.050 | 0.055 |
| 68 | 0.060 | 0.055 | 0.064 | 0.050 | 0.055 |
| 69 | 0.060 | 0.055 | 0.065 | 0.050 | 0.056 |
| 70 | 0.060 | 0.056 | 0.066 | 0.051 | 0.057 |
| 71 | 0.061 | 0.059 | 0.067 | 0.052 | 0.059 |
| 72 | 0.061 | 0.061 | 0.068 | 0.052 | 0.060 |
| 73 | 0.062 | 0.063 | 0.068 | 0.053 | 0.060 |
| 74 | 0.064 | 0.066 | 0.068 | 0.054 | 0.060 |
| 75 | 0.064 | 0.068 | 0.068 | 0.057 | 0.060 |
| 76 | 0.064 | 0.071 | 0.068 | 0.059 | 0.061 |
| 77 | 0.066 | 0.073 | 0.068 | 0.062 | 0.061 |
| 78 | 0.067 | 0.073 | 0.068 | 0.064 | 0.061 |
| 79 | 0.068 | 0.073 | 0.069 | 0.065 | 0.061 |
| 80 | 0.069 | 0.074 | 0.069 | 0.065 | 0.061 |
| 81 | 0.069 | 0.074 | 0.069 | 0.065 | 0.061 |
| 82 | 0.070 | 0.074 | 0.069 | 0.066 | 0.061 |
| 83 | 0.071 | 0.074 | 0.070 | 0.066 | 0.061 |
| 84 | 0.073 | 0.074 | 0.071 | 0.067 | 0.061 |
| 85 | 0.074 | 0.075 | 0.073 | 0.069 | 0.063 |
| 86 | 0.074 | 0.077 | 0.074 | 0.070 | 0.065 |
| 87 | 0.075 | 0.080 | 0.075 | 0.071 | 0.068 |
| 88 | 0.077 | 0.082 | 0.077 | 0.078 | 0.070 |
| 89 | 0.080 | 0.083 | 0.078 | 0.084 | 0.071 |
| 90 | 0.081 | 0.083 | 0.079 | 0.091 | 0.073 |
| 91 | 0.082 | 0.083 | 0.080 | 0.097 | 0.074 |
| 92 | 0.083 | 0.083 | 0.080 | 0.107 | 0.075 |
| 93 | 0.083 | 0.083 | 0.081 | 0.118 | 0.077 |
| 94 | 0.084 | 0.084 | 0.081 | 0.130 | 0.079 |
| 95 | 0.089 | 0.084 | 0.081 | 0.141 | 0.080 |
| 96 | 0.092 | 0.084 | 0.082 | 0.154 | 0.082 |
| 97 | 0.098 | 0.085 | 0.084 | 0.171 | 0.088 |
| 98 | 0.105 | 0.087 | 0.087 | 0.188 | 0.094 |
| 99 | 0.146 | 0.089 | 0.089 | 0.206 | 0.100 |
| 100 | 0.223 | 0.090 | 0.092 | 0.223 | 0.106 |

***Table S47:*** *Paced tapping with music – Variability (CV of ITI), music 2 – Paced_music_ross_mean_CV_iti*

| | | | | | | |
|---|---|---|---|---|---|---|
| **Task** | | Paced tapping with music | | | | |
| **Outcome measure** | | Variability (CV of ITI), music 2 | | | | |
| **Variable name** | | Paced_music_ross_mean_CV_iti | | | | |
| **Age and gender effects** | | | | | | |
| Age regression (slope) | | 0.000 | | | | |
| Age regression (p) | | 0.201 | | | | |
| Age group (p) | | 0.232 | | | | |
| Gender (p) | | 0.557 | | | | |
| **Group** | | All | Age 18 to 21 | Age 22 to 29 | Age 30 to 54 | Age 55 to 87 |
| N (N female) | | 102 (70) | 24 (17) | 28 (18) | 25 (15) | 25 (20) |
| **Normality** | | | | | | |
| Skewness | | 3.63 | 1.80 | 0.74 | 2.56 | 0.20 |
| Excess kurtosis | | 18.32 | 2.91 | -0.06 | 6.39 | -0.81 |
| **Scores** | | | | | | |
| Mean | | 0.057 | 0.060 | 0.057 | 0.063 | 0.049 |
| SD | | 0.031 | 0.026 | 0.018 | 0.052 | 0.011 |
| Percentiles | | | | | | |
| | 0 | 0.026 | 0.031 | 0.034 | 0.026 | 0.030 |
| | 1 | 0.027 | 0.032 | 0.034 | 0.026 | 0.030 |
| | 2 | 0.028 | 0.034 | 0.034 | 0.027 | 0.031 |
| | 3 | 0.030 | 0.035 | 0.035 | 0.027 | 0.031 |
| | 4 | 0.031 | 0.036 | 0.035 | 0.027 | 0.032 |
| | 5 | 0.031 | 0.037 | 0.036 | 0.028 | 0.033 |
| | 6 | 0.032 | 0.037 | 0.036 | 0.028 | 0.034 |
| | 7 | 0.032 | 0.037 | 0.037 | 0.028 | 0.034 |
| | 8 | 0.034 | 0.038 | 0.037 | 0.028 | 0.035 |
| | 9 | 0.034 | 0.038 | 0.037 | 0.028 | 0.036 |
| | 10 | 0.034 | 0.039 | 0.037 | 0.029 | 0.036 |
| | 11 | 0.035 | 0.040 | 0.038 | 0.030 | 0.036 |
| | 12 | 0.036 | 0.041 | 0.038 | 0.031 | 0.036 |
| | 13 | 0.036 | 0.042 | 0.038 | 0.031 | 0.036 |
| | 14 | 0.037 | 0.042 | 0.038 | 0.032 | 0.036 |
| | 15 | 0.037 | 0.042 | 0.038 | 0.032 | 0.036 |
| | 16 | 0.037 | 0.042 | 0.038 | 0.032 | 0.036 |
| | 17 | 0.037 | 0.042 | 0.038 | 0.032 | 0.037 |
| | 18 | 0.037 | 0.042 | 0.038 | 0.033 | 0.037 |
| | 19 | 0.038 | 0.042 | 0.039 | 0.033 | 0.037 |
| | 20 | 0.038 | 0.042 | 0.039 | 0.033 | 0.038 |
| | 21 | 0.038 | 0.042 | 0.039 | 0.034 | 0.038 |

| | | | | | |
|---|---|---|---|---|---|
| 22 | 0.038 | 0.042 | 0.039 | 0.034 | 0.038 |
| 23 | 0.039 | 0.043 | 0.040 | 0.034 | 0.038 |
| 24 | 0.039 | 0.044 | 0.042 | 0.034 | 0.038 |
| 25 | 0.039 | 0.044 | 0.043 | 0.034 | 0.039 |
| 26 | 0.040 | 0.045 | 0.044 | 0.035 | 0.039 |
| 27 | 0.041 | 0.045 | 0.045 | 0.035 | 0.040 |
| 28 | 0.042 | 0.045 | 0.045 | 0.036 | 0.041 |
| 29 | 0.042 | 0.045 | 0.045 | 0.037 | 0.041 |
| 30 | 0.042 | 0.045 | 0.045 | 0.037 | 0.042 |
| 31 | 0.042 | 0.046 | 0.047 | 0.037 | 0.042 |
| 32 | 0.042 | 0.046 | 0.048 | 0.037 | 0.042 |
| 33 | 0.043 | 0.047 | 0.049 | 0.037 | 0.042 |
| 34 | 0.044 | 0.048 | 0.049 | 0.037 | 0.042 |
| 35 | 0.044 | 0.048 | 0.050 | 0.038 | 0.043 |
| 36 | 0.045 | 0.048 | 0.050 | 0.038 | 0.043 |
| 37 | 0.045 | 0.048 | 0.050 | 0.039 | 0.044 |
| 38 | 0.045 | 0.049 | 0.050 | 0.039 | 0.044 |
| 39 | 0.046 | 0.049 | 0.050 | 0.040 | 0.045 |
| 40 | 0.047 | 0.049 | 0.050 | 0.040 | 0.046 |
| 41 | 0.047 | 0.049 | 0.050 | 0.040 | 0.047 |
| 42 | 0.048 | 0.050 | 0.050 | 0.041 | 0.047 |
| 43 | 0.049 | 0.050 | 0.051 | 0.041 | 0.048 |
| 44 | 0.049 | 0.051 | 0.051 | 0.041 | 0.048 |
| 45 | 0.049 | 0.052 | 0.052 | 0.042 | 0.048 |
| 46 | 0.050 | 0.053 | 0.052 | 0.042 | 0.049 |
| 47 | 0.050 | 0.053 | 0.053 | 0.043 | 0.049 |
| 48 | 0.050 | 0.054 | 0.054 | 0.043 | 0.050 |
| 49 | 0.050 | 0.055 | 0.054 | 0.044 | 0.050 |
| 50 | 0.051 | 0.055 | 0.054 | 0.044 | 0.050 |
| 51 | 0.051 | 0.056 | 0.054 | 0.045 | 0.051 |
| 52 | 0.051 | 0.056 | 0.054 | 0.045 | 0.051 |
| 53 | 0.052 | 0.056 | 0.055 | 0.046 | 0.051 |
| 54 | 0.052 | 0.056 | 0.056 | 0.046 | 0.051 |
| 55 | 0.052 | 0.057 | 0.056 | 0.047 | 0.051 |
| 56 | 0.053 | 0.057 | 0.057 | 0.047 | 0.051 |
| 57 | 0.053 | 0.057 | 0.057 | 0.048 | 0.051 |
| 58 | 0.054 | 0.057 | 0.058 | 0.048 | 0.051 |
| 59 | 0.054 | 0.057 | 0.058 | 0.049 | 0.051 |
| 60 | 0.054 | 0.057 | 0.058 | 0.050 | 0.051 |
| 61 | 0.054 | 0.057 | 0.058 | 0.051 | 0.052 |
| 62 | 0.055 | 0.058 | 0.059 | 0.052 | 0.052 |
| 63 | 0.056 | 0.058 | 0.059 | 0.052 | 0.052 |
| 64 | 0.057 | 0.058 | 0.060 | 0.053 | 0.052 |

| | | | | | |
|---|---|---|---|---|---|
| 65 | 0.057 | 0.058 | 0.061 | 0.053 | 0.052 |
| 66 | 0.057 | 0.060 | 0.063 | 0.053 | 0.052 |
| 67 | 0.058 | 0.062 | 0.063 | 0.053 | 0.053 |
| 68 | 0.058 | 0.064 | 0.064 | 0.054 | 0.053 |
| 69 | 0.058 | 0.066 | 0.064 | 0.054 | 0.053 |
| 70 | 0.059 | 0.067 | 0.064 | 0.055 | 0.054 |
| 71 | 0.059 | 0.067 | 0.064 | 0.057 | 0.054 |
| 72 | 0.061 | 0.067 | 0.065 | 0.062 | 0.054 |
| 73 | 0.063 | 0.068 | 0.067 | 0.068 | 0.054 |
| 74 | 0.064 | 0.068 | 0.068 | 0.074 | 0.054 |
| 75 | 0.065 | 0.068 | 0.069 | 0.079 | 0.054 |
| 76 | 0.067 | 0.069 | 0.070 | 0.080 | 0.055 |
| 77 | 0.068 | 0.069 | 0.072 | 0.081 | 0.056 |
| 78 | 0.068 | 0.069 | 0.072 | 0.082 | 0.057 |
| 79 | 0.069 | 0.070 | 0.073 | 0.082 | 0.058 |
| 80 | 0.069 | 0.070 | 0.073 | 0.083 | 0.058 |
| 81 | 0.070 | 0.070 | 0.073 | 0.084 | 0.058 |
| 82 | 0.071 | 0.070 | 0.073 | 0.084 | 0.059 |
| 83 | 0.072 | 0.071 | 0.073 | 0.085 | 0.059 |
| 84 | 0.073 | 0.072 | 0.073 | 0.086 | 0.059 |
| 85 | 0.073 | 0.073 | 0.074 | 0.087 | 0.060 |
| 86 | 0.075 | 0.074 | 0.074 | 0.088 | 0.061 |
| 87 | 0.076 | 0.075 | 0.075 | 0.088 | 0.061 |
| 88 | 0.079 | 0.077 | 0.076 | 0.090 | 0.062 |
| 89 | 0.081 | 0.078 | 0.077 | 0.092 | 0.063 |
| 90 | 0.082 | 0.080 | 0.078 | 0.093 | 0.064 |
| 91 | 0.082 | 0.082 | 0.079 | 0.095 | 0.065 |
| 92 | 0.085 | 0.089 | 0.081 | 0.103 | 0.066 |
| 93 | 0.089 | 0.099 | 0.083 | 0.123 | 0.067 |
| 94 | 0.094 | 0.109 | 0.086 | 0.142 | 0.068 |
| 95 | 0.096 | 0.120 | 0.090 | 0.162 | 0.069 |
| 96 | 0.102 | 0.127 | 0.093 | 0.181 | 0.069 |
| 97 | 0.126 | 0.130 | 0.095 | 0.200 | 0.069 |
| 98 | 0.137 | 0.132 | 0.098 | 0.219 | 0.070 |
| 99 | 0.178 | 0.135 | 0.100 | 0.238 | 0.070 |
| 100 | 0.257 | 0.137 | 0.102 | 0.257 | 0.070 |

***Table S48:*** *Paced tapping with music – Consistency (logit of vector length), music 1 – Paced_music_badine_vecLenLogit*

| | | | | | | |
|---|---|---|---|---|---|---|
| **Task** | Paced tapping with music | | | | | |
| **Outcome measure** | Consistency (logit of vector length), music 1 | | | | | |
| **Variable name** | Paced_music_badine_vecLenLogit | | | | | |
| **Age and gender effects** | | | | | | |
| Age regression (slope) | | 0.006 | | | | |
| Age regression (p) | | 0.249 | | | | |
| Age group (p) | | 0.037 | | | | |
| Gender (p) | | 0.677 | | | | |
| **Group** | | All | Age 18 to 21 | Age 22 to 29 | Age 30 to 54 | Age 55 to 87 |
| N (N female) | | 106 (72) | 27 (18) | 29 (19) | 24 (14) | 26 (21) |
| **Normality** | | | | | | |
| Skewness | | -1.30 | -1.02 | -0.63 | -1.80 | -1.19 |
| Excess kurtosis | | 0.74 | 0.03 | -0.99 | 2.38 | 2.06 |
| **Scores** | | | | | | |
| Mean | | 2.78 | 2.60 | 2.18 | 3.23 | 3.21 |
| SD | | 1.52 | 1.70 | 1.80 | 1.37 | 0.68 |
| Percentiles | | | | | | |
| | 0 | -1.81 | -1.81 | -1.03 | -0.97 | 1.07 |
| | 1 | -1.03 | -1.50 | -1.02 | -0.65 | 1.32 |
| | 2 | -1.01 | -1.18 | -1.02 | -0.32 | 1.57 |
| | 3 | -0.96 | -0.87 | -1.02 | 0.00 | 1.82 |
| | 4 | -0.86 | -0.57 | -1.00 | 0.32 | 2.07 |
| | 5 | -0.75 | -0.38 | -0.96 | 0.50 | 2.17 |
| | 6 | -0.55 | -0.19 | -0.92 | 0.61 | 2.28 |
| | 7 | -0.30 | 0.01 | -0.89 | 0.72 | 2.38 |
| | 8 | 0.01 | 0.16 | -0.86 | 0.82 | 2.49 |
| | 9 | 0.25 | 0.23 | -0.84 | 0.95 | 2.53 |
| | 10 | 0.40 | 0.29 | -0.81 | 1.12 | 2.57 |
| | 11 | 0.43 | 0.36 | -0.76 | 1.29 | 2.61 |
| | 12 | 0.47 | 0.40 | -0.66 | 1.46 | 2.65 |
| | 13 | 0.67 | 0.40 | -0.56 | 1.63 | 2.66 |
| | 14 | 0.86 | 0.41 | -0.46 | 1.93 | 2.66 |
| | 15 | 0.94 | 0.41 | -0.36 | 2.24 | 2.67 |
| | 16 | 0.99 | 0.51 | -0.26 | 2.54 | 2.68 |
| | 17 | 1.06 | 0.66 | -0.16 | 2.85 | 2.69 |
| | 18 | 1.12 | 0.81 | -0.05 | 2.98 | 2.70 |
| | 19 | 1.53 | 0.96 | 0.11 | 2.98 | 2.71 |
| | 20 | 1.64 | 1.02 | 0.26 | 2.99 | 2.71 |
| | 21 | 1.77 | 1.06 | 0.42 | 3.00 | 2.78 |

| | | | | | |
|---|---|---|---|---|---|
| 22 | 2.08 | 1.09 | 0.53 | 3.02 | 2.84 |
| 23 | 2.17 | 1.13 | 0.61 | 3.05 | 2.90 |
| 24 | 2.33 | 1.23 | 0.69 | 3.07 | 2.96 |
| 25 | 2.53 | 1.34 | 0.78 | 3.10 | 2.96 |
| 26 | 2.65 | 1.45 | 0.83 | 3.13 | 2.96 |
| 27 | 2.67 | 1.57 | 0.88 | 3.14 | 2.96 |
| 28 | 2.70 | 1.88 | 0.93 | 3.14 | 2.96 |
| 29 | 2.72 | 2.18 | 1.05 | 3.14 | 2.98 |
| 30 | 2.75 | 2.49 | 1.28 | 3.15 | 2.99 |
| 31 | 2.81 | 2.73 | 1.50 | 3.19 | 3.00 |
| 32 | 2.84 | 2.74 | 1.73 | 3.26 | 3.01 |
| 33 | 2.86 | 2.75 | 1.85 | 3.34 | 3.03 |
| 34 | 2.91 | 2.77 | 1.96 | 3.41 | 3.05 |
| 35 | 2.95 | 2.78 | 2.07 | 3.47 | 3.07 |
| 36 | 2.96 | 2.81 | 2.16 | 3.48 | 3.08 |
| 37 | 2.96 | 2.83 | 2.20 | 3.49 | 3.12 |
| 38 | 2.97 | 2.86 | 2.24 | 3.49 | 3.16 |
| 39 | 2.97 | 2.88 | 2.28 | 3.50 | 3.20 |
| 40 | 3.01 | 2.89 | 2.37 | 3.55 | 3.24 |
| 41 | 3.02 | 2.91 | 2.47 | 3.59 | 3.24 |
| 42 | 3.07 | 2.93 | 2.57 | 3.64 | 3.25 |
| 43 | 3.09 | 2.96 | 2.67 | 3.69 | 3.25 |
| 44 | 3.11 | 3.00 | 2.72 | 3.71 | 3.26 |
| 45 | 3.12 | 3.03 | 2.77 | 3.72 | 3.26 |
| 46 | 3.14 | 3.07 | 2.81 | 3.73 | 3.26 |
| 47 | 3.17 | 3.08 | 2.84 | 3.74 | 3.26 |
| 48 | 3.21 | 3.09 | 2.84 | 3.75 | 3.26 |
| 49 | 3.22 | 3.10 | 2.84 | 3.77 | 3.28 |
| 50 | 3.25 | 3.11 | 2.84 | 3.78 | 3.29 |
| 51 | 3.26 | 3.15 | 2.87 | 3.79 | 3.30 |
| 52 | 3.26 | 3.19 | 2.90 | 3.80 | 3.32 |
| 53 | 3.30 | 3.23 | 2.93 | 3.81 | 3.33 |
| 54 | 3.34 | 3.27 | 2.95 | 3.81 | 3.34 |
| 55 | 3.35 | 3.29 | 2.96 | 3.81 | 3.35 |
| 56 | 3.36 | 3.31 | 2.96 | 3.82 | 3.36 |
| 57 | 3.39 | 3.33 | 2.97 | 3.83 | 3.38 |
| 58 | 3.40 | 3.35 | 3.00 | 3.84 | 3.40 |
| 59 | 3.44 | 3.36 | 3.04 | 3.86 | 3.42 |
| 60 | 3.47 | 3.37 | 3.08 | 3.87 | 3.45 |
| 61 | 3.50 | 3.39 | 3.11 | 3.89 | 3.46 |
| 62 | 3.50 | 3.44 | 3.14 | 3.90 | 3.47 |
| 63 | 3.51 | 3.53 | 3.17 | 3.90 | 3.49 |
| 64 | 3.55 | 3.63 | 3.20 | 3.91 | 3.50 |

| | | | | | |
|---|---|---|---|---|---|
| 65 | 3.65 | 3.73 | 3.21 | 3.92 | 3.50 |
| 66 | 3.69 | 3.77 | 3.21 | 3.93 | 3.50 |
| 67 | 3.71 | 3.78 | 3.21 | 3.94 | 3.50 |
| 68 | 3.73 | 3.79 | 3.22 | 3.96 | 3.51 |
| 69 | 3.76 | 3.80 | 3.26 | 3.97 | 3.51 |
| 70 | 3.78 | 3.80 | 3.30 | 3.98 | 3.51 |
| 71 | 3.79 | 3.81 | 3.34 | 3.98 | 3.52 |
| 72 | 3.80 | 3.82 | 3.36 | 3.98 | 3.52 |
| 73 | 3.81 | 3.82 | 3.38 | 3.98 | 3.55 |
| 74 | 3.82 | 3.84 | 3.39 | 3.98 | 3.58 |
| 75 | 3.84 | 3.86 | 3.40 | 3.99 | 3.61 |
| 76 | 3.85 | 3.88 | 3.48 | 3.99 | 3.64 |
| 77 | 3.88 | 3.89 | 3.56 | 3.99 | 3.66 |
| 78 | 3.89 | 3.91 | 3.64 | 3.99 | 3.67 |
| 79 | 3.89 | 3.92 | 3.71 | 4.02 | 3.69 |
| 80 | 3.92 | 3.93 | 3.75 | 4.06 | 3.70 |
| 81 | 3.94 | 3.94 | 3.79 | 4.09 | 3.73 |
| 82 | 3.95 | 3.94 | 3.84 | 4.13 | 3.75 |
| 83 | 3.98 | 3.94 | 3.84 | 4.16 | 3.77 |
| 84 | 3.98 | 3.95 | 3.85 | 4.17 | 3.79 |
| 85 | 4.01 | 3.96 | 3.85 | 4.19 | 3.82 |
| 86 | 4.08 | 4.00 | 3.87 | 4.20 | 3.84 |
| 87 | 4.09 | 4.03 | 3.94 | 4.21 | 3.86 |
| 88 | 4.11 | 4.06 | 4.01 | 4.22 | 3.89 |
| 89 | 4.13 | 4.09 | 4.08 | 4.23 | 3.94 |
| 90 | 4.15 | 4.11 | 4.14 | 4.23 | 3.99 |
| 91 | 4.16 | 4.13 | 4.18 | 4.24 | 4.04 |
| 92 | 4.19 | 4.16 | 4.23 | 4.26 | 4.08 |
| 93 | 4.22 | 4.17 | 4.27 | 4.29 | 4.09 |
| 94 | 4.23 | 4.19 | 4.30 | 4.32 | 4.10 |
| 95 | 4.26 | 4.20 | 4.33 | 4.35 | 4.11 |
| 96 | 4.35 | 4.21 | 4.36 | 4.37 | 4.12 |
| 97 | 4.37 | 4.34 | 4.38 | 4.39 | 4.13 |
| 98 | 4.42 | 4.48 | 4.40 | 4.41 | 4.14 |
| 99 | 4.44 | 4.62 | 4.41 | 4.42 | 4.14 |
| 100 | 4.76 | 4.76 | 4.42 | 4.44 | 4.15 |

***Table S49:*** *Paced tapping with music – Consistency (logit of vector length), music 2 – Paced_music_ross_vecLenLogit*

| | | | | | | |
|---|---|---|---|---|---|---|
| **Task** | Paced tapping with music | | | | | |
| **Outcome measure** | Consistency (logit of vector length), music 2 | | | | | |
| **Variable name** | Paced_music_ross_vecLenLogit | | | | | |
| **Age and gender effects** | | | | | | |
| Age regression (slope) | | 0.022 | | | | |
| Age regression (p) | | 0.005 | | | | |
| Age group (p) | | 0.010 | | | | |
| Gender (p) | | 0.580 | | | | |
| **Group** | | All | Age 18 to 21 | Age 22 to 29 | Age 30 to 54 | Age 55 to 87 |
| N (N female) | | 102 (70) | 24 (17) | 28 (18) | 25 (15) | 25 (20) |
| **Normality** | | | | | | |
| Skewness | | -0.90 | -0.49 | -0.42 | -0.98 | -1.60 |
| Excess kurtosis | | 0.09 | -0.88 | -0.83 | 0.40 | 3.71 |
| **Scores** | | | | | | |
| Mean | | 2.35 | 1.92 | 1.62 | 2.96 | 2.98 |
| SD | | 1.71 | 1.61 | 2.05 | 1.41 | 1.20 |
| Percentiles | | | | | | |
| | 0 | -2.74 | -1.54 | -2.74 | -0.93 | -1.18 |
| | 1 | -1.88 | -1.36 | -2.51 | -0.55 | -0.52 |
| | 2 | -1.69 | -1.18 | -2.27 | -0.17 | 0.14 |
| | 3 | -1.52 | -1.00 | -2.04 | 0.21 | 0.80 |
| | 4 | -1.17 | -0.82 | -1.86 | 0.59 | 1.46 |
| | 5 | -0.94 | -0.65 | -1.81 | 0.73 | 1.62 |
| | 6 | -0.92 | -0.48 | -1.76 | 0.82 | 1.69 |
| | 7 | -0.75 | -0.32 | -1.71 | 0.91 | 1.76 |
| | 8 | -0.68 | -0.15 | -1.57 | 1.01 | 1.83 |
| | 9 | -0.38 | -0.02 | -1.37 | 1.10 | 1.90 |
| | 10 | -0.14 | 0.01 | -1.17 | 1.20 | 1.95 |
| | 11 | -0.02 | 0.04 | -0.97 | 1.30 | 2.01 |
| | 12 | 0.12 | 0.08 | -0.89 | 1.39 | 2.07 |
| | 13 | 0.14 | 0.11 | -0.82 | 1.48 | 2.10 |
| | 14 | 0.21 | 0.12 | -0.76 | 1.57 | 2.12 |
| | 15 | 0.55 | 0.13 | -0.69 | 1.65 | 2.14 |
| | 16 | 0.61 | 0.13 | -0.61 | 1.73 | 2.16 |
| | 17 | 0.68 | 0.14 | -0.53 | 1.79 | 2.17 |
| | 18 | 0.87 | 0.14 | -0.44 | 1.81 | 2.17 |
| | 19 | 1.04 | 0.15 | -0.37 | 1.82 | 2.18 |
| | 20 | 1.06 | 0.15 | -0.30 | 1.84 | 2.18 |
| | 21 | 1.13 | 0.15 | -0.24 | 1.86 | 2.19 |

| | | | | | |
|---|---|---|---|---|---|
| 22 | 1.20 | 0.21 | -0.17 | 1.92 | 2.23 |
| 23 | 1.25 | 0.41 | -0.01 | 1.98 | 2.27 |
| 24 | 1.42 | 0.62 | 0.18 | 2.04 | 2.30 |
| 25 | 1.46 | 0.82 | 0.37 | 2.10 | 2.34 |
| 26 | 1.54 | 1.03 | 0.54 | 2.13 | 2.38 |
| 27 | 1.58 | 1.06 | 0.56 | 2.15 | 2.41 |
| 28 | 1.62 | 1.08 | 0.58 | 2.18 | 2.44 |
| 29 | 1.66 | 1.09 | 0.59 | 2.20 | 2.47 |
| 30 | 1.71 | 1.11 | 0.63 | 2.26 | 2.51 |
| 31 | 1.81 | 1.13 | 0.69 | 2.33 | 2.54 |
| 32 | 1.85 | 1.15 | 0.75 | 2.40 | 2.57 |
| 33 | 1.85 | 1.17 | 0.81 | 2.46 | 2.60 |
| 34 | 1.87 | 1.18 | 0.90 | 2.49 | 2.64 |
| 35 | 1.96 | 1.21 | 1.00 | 2.50 | 2.69 |
| 36 | 2.10 | 1.26 | 1.10 | 2.51 | 2.74 |
| 37 | 2.10 | 1.31 | 1.20 | 2.52 | 2.79 |
| 38 | 2.13 | 1.36 | 1.28 | 2.58 | 2.84 |
| 39 | 2.17 | 1.41 | 1.37 | 2.69 | 2.90 |
| 40 | 2.19 | 1.50 | 1.46 | 2.80 | 2.96 |
| 41 | 2.26 | 1.60 | 1.53 | 2.90 | 3.01 |
| 42 | 2.38 | 1.70 | 1.55 | 3.00 | 3.05 |
| 43 | 2.46 | 1.80 | 1.57 | 3.08 | 3.06 |
| 44 | 2.48 | 1.88 | 1.59 | 3.15 | 3.06 |
| 45 | 2.50 | 1.94 | 1.61 | 3.23 | 3.06 |
| 46 | 2.57 | 2.00 | 1.63 | 3.30 | 3.06 |
| 47 | 2.63 | 2.06 | 1.64 | 3.34 | 3.08 |
| 48 | 2.66 | 2.13 | 1.66 | 3.38 | 3.10 |
| 49 | 2.70 | 2.26 | 1.66 | 3.41 | 3.11 |
| 50 | 2.76 | 2.39 | 1.67 | 3.45 | 3.13 |
| 51 | 2.78 | 2.51 | 1.67 | 3.50 | 3.14 |
| 52 | 2.80 | 2.64 | 1.68 | 3.55 | 3.15 |
| 53 | 2.83 | 2.68 | 1.74 | 3.59 | 3.15 |
| 54 | 2.84 | 2.70 | 1.80 | 3.64 | 3.16 |
| 55 | 2.89 | 2.72 | 1.86 | 3.65 | 3.19 |
| 56 | 2.95 | 2.74 | 1.96 | 3.66 | 3.23 |
| 57 | 2.97 | 2.75 | 2.11 | 3.66 | 3.26 |
| 58 | 3.02 | 2.75 | 2.25 | 3.67 | 3.30 |
| 59 | 3.06 | 2.76 | 2.40 | 3.69 | 3.34 |
| 60 | 3.10 | 2.77 | 2.48 | 3.71 | 3.39 |
| 61 | 3.15 | 2.77 | 2.54 | 3.74 | 3.44 |
| 62 | 3.20 | 2.79 | 2.60 | 3.76 | 3.48 |
| 63 | 3.26 | 2.81 | 2.65 | 3.79 | 3.52 |
| 64 | 3.30 | 2.83 | 2.69 | 3.80 | 3.54 |

| | | | | | |
|---|---|---|---|---|---|
| 65 | 3.40 | 2.84 | 2.73 | 3.82 | 3.57 |
| 66 | 3.45 | 2.86 | 2.77 | 3.84 | 3.60 |
| 67 | 3.49 | 2.88 | 2.79 | 3.86 | 3.63 |
| 68 | 3.58 | 2.90 | 2.81 | 3.88 | 3.66 |
| 69 | 3.64 | 2.92 | 2.82 | 3.90 | 3.70 |
| 70 | 3.65 | 2.94 | 2.83 | 3.93 | 3.73 |
| 71 | 3.66 | 2.94 | 2.90 | 3.95 | 3.77 |
| 72 | 3.67 | 2.95 | 3.00 | 3.98 | 3.78 |
| 73 | 3.69 | 2.96 | 3.11 | 4.01 | 3.79 |
| 74 | 3.74 | 2.97 | 3.21 | 4.05 | 3.81 |
| 75 | 3.77 | 3.09 | 3.38 | 4.08 | 3.82 |
| 76 | 3.81 | 3.20 | 3.55 | 4.08 | 3.86 |
| 77 | 3.84 | 3.31 | 3.72 | 4.09 | 3.91 |
| 78 | 3.85 | 3.42 | 3.85 | 4.09 | 3.95 |
| 79 | 3.86 | 3.49 | 3.85 | 4.10 | 3.99 |
| 80 | 3.90 | 3.53 | 3.85 | 4.11 | 4.00 |
| 81 | 3.94 | 3.58 | 3.86 | 4.12 | 4.01 |
| 82 | 3.98 | 3.63 | 3.89 | 4.13 | 4.01 |
| 83 | 3.99 | 3.66 | 3.95 | 4.14 | 4.02 |
| 84 | 4.02 | 3.66 | 4.01 | 4.19 | 4.04 |
| 85 | 4.07 | 3.66 | 4.07 | 4.25 | 4.07 |
| 86 | 4.08 | 3.66 | 4.08 | 4.31 | 4.10 |
| 87 | 4.09 | 3.66 | 4.08 | 4.37 | 4.13 |
| 88 | 4.10 | 3.67 | 4.08 | 4.40 | 4.15 |
| 89 | 4.13 | 3.68 | 4.09 | 4.40 | 4.16 |
| 90 | 4.14 | 3.68 | 4.10 | 4.40 | 4.17 |
| 91 | 4.15 | 3.69 | 4.11 | 4.40 | 4.18 |
| 92 | 4.15 | 3.73 | 4.12 | 4.41 | 4.19 |
| 93 | 4.18 | 3.78 | 4.13 | 4.44 | 4.20 |
| 94 | 4.22 | 3.83 | 4.14 | 4.46 | 4.21 |
| 95 | 4.27 | 3.88 | 4.14 | 4.49 | 4.22 |
| 96 | 4.39 | 3.92 | 4.15 | 4.52 | 4.24 |
| 97 | 4.40 | 3.93 | 4.17 | 4.53 | 4.33 |
| 98 | 4.51 | 3.95 | 4.21 | 4.55 | 4.41 |
| 99 | 4.59 | 3.97 | 4.24 | 4.57 | 4.50 |
| 100 | 4.59 | 3.99 | 4.27 | 4.59 | 4.59 |

***Table S50:*** *Paced tapping with music – Accuracy (vector direction %), music 1 – Paced_music_badine_vecDirPct*

| | | | | | | |
|---|---|---|---|---|---|---|
| **Task** | Paced tapping with music | | | | | |
| **Outcome measure** | Accuracy (vector direction %), music 1 | | | | | |
| **Variable name** | Paced_music_badine_vecDirPct | | | | | |
| **Age and gender effects** | | | | | | |
| Age regression (slope) | | -0.008 | | | | |
| Age regression (p) | | 0.742 | | | | |
| Age group (p) | | 0.836 | | | | |
| Gender (p) | | 0.718 | | | | |
| **Group** | | All | Age 18 to 21 | Age 22 to 29 | Age 30 to 54 | Age 55 to 87 |
| N (N female) | | 99 (70) | 25 (17) | 25 (18) | 23 (14) | 26 (21) |
| **Normality** | | | | | | |
| Skewness | | 0.54 | -0.11 | 0.45 | 1.66 | 0.39 |
| Excess kurtosis | | 1.54 | 2.56 | 0.01 | 2.79 | -0.17 |
| **Scores** | | | | | | |
| Mean | | 0.3 | 0.5 | 0.2 | 0.2 | 0.3 |
| SD | | 5.4 | 5.7 | 6.6 | 5.2 | 4.1 |
| Percentiles | | | | | | |
| | 0 | -15.5 | -15.5 | -11.0 | -7.4 | -7.8 |
| | 1 | -11.1 | -13.7 | -10.9 | -7.0 | -7.1 |
| | 2 | -10.3 | -12.0 | -10.7 | -6.6 | -6.4 |
| | 3 | -8.4 | -10.3 | -10.5 | -6.2 | -5.7 |
| | 4 | -7.9 | -8.6 | -10.3 | -5.8 | -4.9 |
| | 5 | -7.5 | -7.6 | -9.7 | -5.4 | -4.8 |
| | 6 | -7.4 | -6.8 | -9.0 | -5.1 | -4.7 |
| | 7 | -6.5 | -6.0 | -8.4 | -4.8 | -4.5 |
| | 8 | -6.2 | -5.2 | -7.7 | -4.5 | -4.4 |
| | 9 | -5.7 | -4.7 | -7.3 | -4.1 | -4.4 |
| | 10 | -5.1 | -4.4 | -7.0 | -3.9 | -4.3 |
| | 11 | -4.9 | -4.0 | -6.8 | -3.6 | -4.3 |
| | 12 | -4.8 | -3.7 | -6.5 | -3.3 | -4.2 |
| | 13 | -4.6 | -3.4 | -6.4 | -3.1 | -4.1 |
| | 14 | -4.4 | -3.2 | -6.3 | -2.9 | -3.9 |
| | 15 | -4.4 | -3.0 | -6.3 | -2.8 | -3.8 |
| | 16 | -4.3 | -2.8 | -6.2 | -2.8 | -3.6 |
| | 17 | -4.2 | -2.6 | -6.1 | -2.7 | -3.5 |
| | 18 | -3.8 | -2.4 | -5.8 | -2.6 | -3.3 |
| | 19 | -3.6 | -2.3 | -5.4 | -2.6 | -3.1 |
| | 20 | -3.5 | -2.1 | -5.0 | -2.6 | -2.9 |
| | 21 | -3.2 | -1.9 | -4.7 | -2.6 | -2.8 |

| | | | | | |
|---|---|---|---|---|---|
| 22 | -2.9 | -1.7 | -4.7 | -2.6 | -2.7 |
| 23 | -2.8 | -1.4 | -4.6 | -2.6 | -2.5 |
| 24 | -2.6 | -1.1 | -4.6 | -2.5 | -2.4 |
| 25 | -2.6 | -0.9 | -4.5 | -2.5 | -2.4 |
| 26 | -2.5 | -0.8 | -4.5 | -2.5 | -2.4 |
| 27 | -2.4 | -0.7 | -4.4 | -2.4 | -2.3 |
| 28 | -2.4 | -0.7 | -4.4 | -2.4 | -2.3 |
| 29 | -2.3 | -0.6 | -4.4 | -2.4 | -2.2 |
| 30 | -2.2 | -0.6 | -4.2 | -2.4 | -2.1 |
| 31 | -2.0 | -0.5 | -4.0 | -2.4 | -1.9 |
| 32 | -1.9 | -0.5 | -3.8 | -2.3 | -1.8 |
| 33 | -1.9 | -0.5 | -3.6 | -2.1 | -1.8 |
| 34 | -1.8 | -0.4 | -3.3 | -1.9 | -1.8 |
| 35 | -1.7 | -0.4 | -2.9 | -1.6 | -1.8 |
| 36 | -1.5 | -0.4 | -2.6 | -1.4 | -1.7 |
| 37 | -1.3 | -0.4 | -2.2 | -1.3 | -1.7 |
| 38 | -1.2 | -0.4 | -2.0 | -1.3 | -1.6 |
| 39 | -1.1 | -0.4 | -2.0 | -1.3 | -1.6 |
| 40 | -1.0 | -0.4 | -1.9 | -1.3 | -1.5 |
| 41 | -0.9 | -0.4 | -1.9 | -1.2 | -1.3 |
| 42 | -0.8 | -0.3 | -1.8 | -1.2 | -1.0 |
| 43 | -0.7 | -0.1 | -1.7 | -1.1 | -0.7 |
| 44 | -0.7 | 0.1 | -1.5 | -1.1 | -0.4 |
| 45 | -0.7 | 0.3 | -1.3 | -1.0 | -0.4 |
| 46 | -0.6 | 0.4 | -1.1 | -1.0 | -0.3 |
| 47 | -0.5 | 0.5 | -0.7 | -1.0 | -0.3 |
| 48 | -0.4 | 0.5 | -0.4 | -1.0 | -0.2 |
| 49 | -0.4 | 0.5 | 0.0 | -1.0 | 0.0 |
| 50 | -0.4 | 0.5 | 0.4 | -0.9 | 0.3 |
| 51 | -0.3 | 0.5 | 0.7 | -0.9 | 0.5 |
| 52 | -0.2 | 0.5 | 1.0 | -0.9 | 0.8 |
| 53 | -0.1 | 0.5 | 1.3 | -0.8 | 0.9 |
| 54 | 0.3 | 0.6 | 1.6 | -0.8 | 1.0 |
| 55 | 0.4 | 0.6 | 1.8 | -0.7 | 1.1 |
| 56 | 0.5 | 0.7 | 1.9 | -0.7 | 1.2 |
| 57 | 0.6 | 0.8 | 2.1 | -0.7 | 1.2 |
| 58 | 0.7 | 0.9 | 2.2 | -0.7 | 1.3 |
| 59 | 0.9 | 1.0 | 2.3 | -0.7 | 1.3 |
| 60 | 1.2 | 1.1 | 2.4 | -0.7 | 1.3 |
| 61 | 1.3 | 1.2 | 2.4 | -0.7 | 1.3 |
| 62 | 1.4 | 1.3 | 2.5 | -0.7 | 1.4 |
| 63 | 1.4 | 1.4 | 2.5 | -0.7 | 1.4 |
| 64 | 1.4 | 1.4 | 2.7 | -0.7 | 1.4 |

| | | | | | |
|---|---|---|---|---|---|
| 65 | 1.5 | 1.4 | 2.8 | -0.6 | 1.4 |
| 66 | 1.6 | 1.4 | 2.9 | -0.6 | 1.5 |
| 67 | 1.7 | 1.5 | 3.0 | -0.6 | 1.5 |
| 68 | 1.8 | 1.6 | 3.0 | -0.5 | 1.6 |
| 69 | 1.9 | 1.7 | 3.0 | -0.4 | 1.6 |
| 70 | 2.0 | 1.8 | 3.0 | -0.3 | 1.6 |
| 71 | 2.2 | 1.9 | 3.1 | -0.2 | 1.6 |
| 72 | 2.3 | 1.9 | 3.1 | -0.1 | 1.7 |
| 73 | 2.4 | 2.0 | 3.2 | 0.1 | 1.8 |
| 74 | 2.7 | 2.1 | 3.2 | 0.5 | 2.0 |
| 75 | 3.0 | 2.2 | 3.3 | 0.9 | 2.2 |
| 76 | 3.0 | 2.5 | 3.8 | 1.3 | 2.4 |
| 77 | 3.2 | 2.9 | 4.2 | 1.7 | 2.5 |
| 78 | 3.4 | 3.2 | 4.6 | 2.1 | 2.6 |
| 79 | 3.6 | 3.6 | 5.1 | 2.5 | 2.8 |
| 80 | 3.8 | 3.8 | 5.2 | 2.8 | 2.9 |
| 81 | 4.0 | 4.0 | 5.3 | 3.2 | 3.2 |
| 82 | 4.1 | 4.1 | 5.4 | 3.5 | 3.5 |
| 83 | 4.3 | 4.3 | 5.4 | 3.6 | 3.7 |
| 84 | 4.4 | 4.5 | 5.6 | 3.7 | 4.0 |
| 85 | 4.6 | 4.8 | 5.7 | 3.9 | 4.1 |
| 86 | 5.2 | 5.0 | 5.9 | 4.0 | 4.2 |
| 87 | 5.4 | 5.3 | 6.0 | 4.0 | 4.3 |
| 88 | 5.5 | 5.4 | 6.1 | 4.1 | 4.4 |
| 89 | 5.7 | 5.5 | 6.2 | 4.1 | 5.1 |
| 90 | 6.2 | 5.5 | 6.3 | 4.2 | 5.9 |
| 91 | 6.6 | 5.5 | 6.4 | 4.4 | 6.7 |
| 92 | 7.5 | 5.8 | 7.0 | 6.2 | 7.4 |
| 93 | 7.7 | 6.4 | 8.6 | 7.9 | 7.5 |
| 94 | 8.4 | 7.1 | 10.2 | 9.7 | 7.5 |
| 95 | 9.8 | 7.7 | 11.8 | 11.5 | 7.6 |
| 96 | 12.3 | 8.6 | 13.2 | 12.7 | 7.7 |
| 97 | 13.3 | 10.4 | 14.0 | 13.6 | 8.1 |
| 98 | 16.1 | 12.3 | 14.7 | 14.4 | 8.6 |
| 99 | 16.2 | 14.2 | 15.5 | 15.3 | 9.1 |
| 100 | 16.2 | 16.1 | 16.2 | 16.2 | 9.5 |

***Table S51:*** *Paced tapping with music – Accuracy (vector direction %), music 2 – Paced_music_ross_vecDirPct*

| | | | | | | |
|---|---|---|---|---|---|---|
| **Task** | Paced tapping with music | | | | | |
| **Outcome measure** | Accuracy (vector direction %), music 2 | | | | | |
| **Variable name** | Paced_music_ross_vecDirPct | | | | | |
| **Age and gender effects** | | | | | | |
| Age regression (slope) | | 0.010 | | | | |
| Age regression (p) | | 0.758 | | | | |
| Age group (p) | | 0.680 | | | | |
| Gender (p) | | 0.165 | | | | |
| **Group** | | All | Age 18 to 21 | Age 22 to 29 | Age 30 to 54 | Age 55 to 87 |
| N (N female) | | 90 (62) | 19 (13) | 23 (15) | 24 (15) | 24 (19) |
| **Normality** | | | | | | |
| Skewness | | -0.66 | -1.10 | -0.48 | -0.35 | -0.69 |
| Excess kurtosis | | 0.77 | 1.83 | -0.52 | 0.08 | 0.15 |
| **Scores** | | | | | | |
| Mean | | -3.4 | -4.1 | -4.0 | -2.0 | -3.8 |
| SD | | 6.6 | 7.4 | 7.6 | 6.6 | 4.9 |
| Percentiles | | | | | | |
| | 0 | -25.0 | -25.0 | -18.4 | -17.9 | -15.8 |
| | 1 | -19.1 | -23.4 | -18.3 | -16.5 | -15.1 |
| | 2 | -18.2 | -21.8 | -18.3 | -15.1 | -14.3 |
| | 3 | -18.0 | -20.2 | -18.2 | -13.7 | -13.6 |
| | 4 | -17.5 | -18.6 | -18.1 | -12.3 | -12.8 |
| | 5 | -16.6 | -17.0 | -18.0 | -11.8 | -12.3 |
| | 6 | -16.0 | -15.5 | -17.8 | -11.7 | -12.0 |
| | 7 | -15.5 | -14.2 | -17.6 | -11.5 | -11.6 |
| | 8 | -14.1 | -12.9 | -17.4 | -11.4 | -11.2 |
| | 9 | -12.6 | -11.6 | -17.1 | -11.2 | -10.9 |
| | 10 | -11.9 | -10.3 | -16.6 | -10.8 | -10.4 |
| | 11 | -11.5 | -8.9 | -15.9 | -10.4 | -10.0 |
| | 12 | -11.1 | -8.7 | -15.3 | -9.9 | -9.6 |
| | 13 | -10.1 | -8.5 | -14.7 | -9.5 | -9.2 |
| | 14 | -9.3 | -8.3 | -13.8 | -8.8 | -8.8 |
| | 15 | -9.0 | -8.1 | -12.5 | -8.1 | -8.4 |
| | 16 | -8.7 | -8.0 | -11.2 | -7.3 | -8.0 |
| | 17 | -8.3 | -7.8 | -9.9 | -6.6 | -7.6 |
| | 18 | -7.9 | -7.7 | -8.6 | -6.2 | -7.4 |
| | 19 | -7.7 | -7.5 | -8.2 | -6.0 | -7.2 |
| | 20 | -7.5 | -7.4 | -8.1 | -5.7 | -7.1 |
| | 21 | -7.2 | -7.2 | -8.0 | -5.5 | -6.9 |

| | | | | | |
|---|---|---|---|---|---|
| 22 | -7.0 | -7.1 | -7.8 | -5.3 | -6.8 |
| 23 | -7.0 | -7.0 | -7.7 | -5.2 | -6.7 |
| 24 | -6.9 | -6.9 | -7.5 | -5.1 | -6.6 |
| 25 | -6.7 | -6.8 | -7.3 | -5.1 | -6.5 |
| 26 | -6.5 | -6.7 | -7.2 | -5.0 | -6.4 |
| 27 | -6.4 | -6.6 | -7.0 | -5.0 | -6.1 |
| 28 | -6.3 | -6.5 | -7.0 | -4.9 | -5.8 |
| 29 | -5.8 | -6.3 | -7.0 | -4.9 | -5.5 |
| 30 | -5.4 | -6.2 | -6.9 | -4.9 | -5.2 |
| 31 | -5.2 | -6.0 | -6.9 | -4.8 | -5.0 |
| 32 | -5.0 | -5.9 | -6.8 | -4.7 | -4.9 |
| 33 | -5.0 | -5.8 | -6.3 | -4.6 | -4.7 |
| 34 | -5.0 | -5.6 | -5.8 | -4.6 | -4.6 |
| 35 | -4.9 | -5.5 | -5.3 | -4.4 | -4.5 |
| 36 | -4.8 | -5.4 | -4.8 | -4.1 | -4.2 |
| 37 | -4.6 | -5.3 | -4.5 | -3.7 | -3.9 |
| 38 | -4.5 | -5.1 | -4.5 | -3.4 | -3.7 |
| 39 | -4.5 | -5.0 | -4.4 | -3.0 | -3.4 |
| 40 | -4.4 | -5.0 | -4.4 | -2.9 | -3.4 |
| 41 | -4.3 | -5.0 | -4.4 | -2.9 | -3.4 |
| 42 | -3.9 | -5.0 | -4.3 | -2.8 | -3.3 |
| 43 | -3.4 | -5.0 | -4.3 | -2.8 | -3.3 |
| 44 | -3.3 | -4.9 | -4.2 | -2.7 | -3.3 |
| 45 | -3.0 | -4.7 | -4.2 | -2.5 | -3.2 |
| 46 | -2.9 | -4.2 | -3.9 | -2.2 | -3.0 |
| 47 | -2.8 | -3.7 | -3.5 | -2.0 | -2.9 |
| 48 | -2.8 | -3.2 | -3.1 | -1.8 | -2.8 |
| 49 | -2.7 | -2.8 | -2.6 | -1.6 | -2.8 |
| 50 | -2.4 | -2.3 | -2.2 | -1.5 | -2.8 |
| 51 | -2.2 | -2.2 | -2.2 | -1.3 | -2.8 |
| 52 | -2.2 | -2.2 | -2.1 | -1.1 | -2.8 |
| 53 | -2.1 | -2.2 | -2.1 | -1.1 | -2.8 |
| 54 | -2.1 | -2.1 | -2.1 | -1.0 | -2.7 |
| 55 | -2.1 | -2.1 | -2.1 | -1.0 | -2.7 |
| 56 | -2.0 | -2.0 | -2.0 | -0.9 | -2.6 |
| 57 | -2.0 | -1.9 | -2.0 | -0.8 | -2.5 |
| 58 | -1.9 | -1.8 | -2.0 | -0.5 | -2.4 |
| 59 | -1.7 | -1.7 | -2.0 | -0.3 | -2.3 |
| 60 | -1.5 | -1.5 | -2.0 | 0.0 | -2.2 |
| 61 | -1.4 | -1.4 | -2.0 | 0.2 | -2.1 |
| 62 | -1.4 | -1.4 | -2.0 | 0.4 | -2.0 |
| 63 | -1.3 | -1.4 | -2.0 | 0.5 | -1.9 |
| 64 | -1.2 | -1.4 | -1.9 | 0.7 | -1.7 |

| | | | | | |
|---|---|---|---|---|---|
| 65 | -1.1 | -1.4 | -1.8 | 0.8 | -1.6 |
| 66 | -1.0 | -1.3 | -1.6 | 1.0 | -1.5 |
| 67 | -0.8 | -1.3 | -1.4 | 1.1 | -1.5 |
| 68 | -0.8 | -1.0 | -1.3 | 1.3 | -1.5 |
| 69 | -0.7 | -0.8 | -1.1 | 1.4 | -1.4 |
| 70 | -0.4 | -0.6 | -0.9 | 1.5 | -1.3 |
| 71 | -0.4 | -0.4 | -0.8 | 1.6 | -1.2 |
| 72 | -0.2 | -0.1 | -0.6 | 1.6 | -1.1 |
| 73 | -0.1 | 0.0 | -0.3 | 1.7 | -0.9 |
| 74 | 0.2 | 0.1 | 0.3 | 1.7 | -0.8 |
| 75 | 0.4 | 0.2 | 0.9 | 1.9 | -0.8 |
| 76 | 0.7 | 0.3 | 1.5 | 2.1 | -0.8 |
| 77 | 0.8 | 0.4 | 2.1 | 2.2 | -0.8 |
| 78 | 1.1 | 0.4 | 2.3 | 2.4 | -0.8 |
| 79 | 1.6 | 0.5 | 2.4 | 2.5 | -0.7 |
| 80 | 1.8 | 0.6 | 2.4 | 2.6 | -0.6 |
| 81 | 1.9 | 0.7 | 2.4 | 2.7 | -0.6 |
| 82 | 2.3 | 0.7 | 2.5 | 2.8 | -0.5 |
| 83 | 2.4 | 0.8 | 3.0 | 2.9 | -0.4 |
| 84 | 2.4 | 1.0 | 3.4 | 3.0 | -0.3 |
| 85 | 2.6 | 1.1 | 3.8 | 3.1 | -0.3 |
| 86 | 2.8 | 1.3 | 4.3 | 3.3 | -0.2 |
| 87 | 3.0 | 1.5 | 4.6 | 3.4 | -0.1 |
| 88 | 3.2 | 1.7 | 5.0 | 4.4 | 0.5 |
| 89 | 3.5 | 1.9 | 5.3 | 5.3 | 1.2 |
| 90 | 3.8 | 2.3 | 5.6 | 6.2 | 1.8 |
| 91 | 3.9 | 2.6 | 5.9 | 7.2 | 2.5 |
| 92 | 4.4 | 3.0 | 6.0 | 7.5 | 2.7 |
| 93 | 5.6 | 3.3 | 6.1 | 7.5 | 2.8 |
| 94 | 6.2 | 3.7 | 6.3 | 7.6 | 3.0 |
| 95 | 7.0 | 4.3 | 6.4 | 7.6 | 3.1 |
| 96 | 7.5 | 5.2 | 6.6 | 7.9 | 3.2 |
| 97 | 7.6 | 6.1 | 6.8 | 8.5 | 3.4 |
| 98 | 7.9 | 7.0 | 7.1 | 9.2 | 3.6 |
| 99 | 9.0 | 7.9 | 7.3 | 9.8 | 3.7 |
| 100 | 10.5 | 8.8 | 7.6 | 10.5 | 3.9 |

***Table S52:*** *Paced tapping with music – Accuracy (vector direction degrees), music 1 – Paced_music_badine_mean_vector_direction*

| | | | | | | |
|---|---|---|---|---|---|---|
| **Task** | | Paced tapping with music | | | | |
| **Outcome measure** | | Accuracy (vector direction degrees), music 1 | | | | |
| **Variable name** | | Paced_music_badine_mean_vector_direction | | | | |
| **Age and gender effects** | | | | | | |
| Age regression (slope) | | -0.013 | | | | |
| Age regression (p) | | 0.887 | | | | |
| Age group (p) | | 0.783 | | | | |
| Gender (p) | | 0.917 | | | | |
| **Group** | | All | Age 18 to 21 | Age 22 to 29 | Age 30 to 54 | Age 55 to 87 |
| N (N female) | | 100 (70) | 25 (17) | 26 (18) | 23 (14) | 26 (21) |
| **Normality** | | | | | | |
| Skewness | | -1.11 | -0.11 | -1.44 | 1.66 | 0.39 |
| Excess kurtosis | | 7.74 | 2.56 | 4.36 | 2.79 | -0.17 |
| **Scores** | | | | | | |
| Mean | | -0.1 | 1.9 | -4.1 | 0.7 | 1.1 |
| SD | | 23.1 | 20.6 | 33.8 | 18.9 | 14.9 |
| Percentiles | | | | | | |
| | 0 | -123.5 | -55.7 | -123.5 | -26.8 | -28.2 |
| | 1 | -56.3 | -49.4 | -102.5 | -25.3 | -25.6 |
| | 2 | -40.1 | -43.2 | -81.6 | -23.8 | -23.0 |
| | 3 | -37.0 | -37.0 | -60.7 | -22.3 | -20.4 |
| | 4 | -30.0 | -30.8 | -39.7 | -20.8 | -17.8 |
| | 5 | -28.2 | -27.4 | -39.0 | -19.5 | -17.3 |
| | 6 | -26.9 | -24.5 | -38.3 | -18.3 | -16.8 |
| | 7 | -26.8 | -21.6 | -37.6 | -17.2 | -16.3 |
| | 8 | -23.3 | -18.8 | -36.9 | -16.1 | -15.8 |
| | 9 | -22.5 | -17.0 | -34.4 | -14.9 | -15.7 |
| | 10 | -20.2 | -15.7 | -31.9 | -13.9 | -15.5 |
| | 11 | -18.0 | -14.4 | -29.4 | -13.0 | -15.4 |
| | 12 | -17.8 | -13.2 | -26.8 | -12.0 | -15.3 |
| | 13 | -17.2 | -12.2 | -25.9 | -11.1 | -14.7 |
| | 14 | -16.3 | -11.5 | -24.9 | -10.4 | -14.2 |
| | 15 | -15.9 | -10.8 | -23.9 | -10.2 | -13.7 |
| | 16 | -15.8 | -10.1 | -23.0 | -9.9 | -13.1 |
| | 17 | -15.4 | -9.4 | -22.8 | -9.7 | -12.5 |
| | 18 | -14.9 | -8.8 | -22.7 | -9.5 | -11.9 |
| | 19 | -13.5 | -8.2 | -22.5 | -9.4 | -11.2 |
| | 20 | -12.9 | -7.6 | -22.4 | -9.3 | -10.6 |
| | 21 | -12.6 | -6.9 | -21.1 | -9.3 | -10.1 |

| | | | | | |
|---|---|---|---|---|---|
| 22 | -11.0 | -6.0 | -19.8 | -9.2 | -9.6 |
| 23 | -10.5 | -5.0 | -18.4 | -9.2 | -9.1 |
| 24 | -9.8 | -4.0 | -17.1 | -9.1 | -8.6 |
| 25 | -9.5 | -3.1 | -16.9 | -9.0 | -8.5 |
| 26 | -9.3 | -2.9 | -16.6 | -8.9 | -8.5 |
| 27 | -8.9 | -2.7 | -16.4 | -8.8 | -8.4 |
| 28 | -8.6 | -2.5 | -16.2 | -8.7 | -8.4 |
| 29 | -8.5 | -2.3 | -16.1 | -8.7 | -7.9 |
| 30 | -8.4 | -2.1 | -16.0 | -8.6 | -7.5 |
| 31 | -7.6 | -2.0 | -15.9 | -8.5 | -7.0 |
| 32 | -7.1 | -1.8 | -15.8 | -8.3 | -6.6 |
| 33 | -6.9 | -1.6 | -15.1 | -7.5 | -6.5 |
| 34 | -6.6 | -1.6 | -14.3 | -6.7 | -6.4 |
| 35 | -6.4 | -1.5 | -13.6 | -5.9 | -6.4 |
| 36 | -5.8 | -1.5 | -12.9 | -5.1 | -6.3 |
| 37 | -5.1 | -1.5 | -11.5 | -4.8 | -6.1 |
| 38 | -4.6 | -1.4 | -10.0 | -4.7 | -5.9 |
| 39 | -4.3 | -1.4 | -8.6 | -4.6 | -5.7 |
| 40 | -3.9 | -1.3 | -7.2 | -4.6 | -5.5 |
| 41 | -3.5 | -1.3 | -7.1 | -4.5 | -4.5 |
| 42 | -3.2 | -1.0 | -7.0 | -4.3 | -3.5 |
| 43 | -2.9 | -0.3 | -6.9 | -4.1 | -2.5 |
| 44 | -2.6 | 0.3 | -6.8 | -3.9 | -1.5 |
| 45 | -2.5 | 1.0 | -6.1 | -3.8 | -1.3 |
| 46 | -2.4 | 1.5 | -5.5 | -3.6 | -1.1 |
| 47 | -2.1 | 1.6 | -4.9 | -3.6 | -1.0 |
| 48 | -1.7 | 1.7 | -4.2 | -3.5 | -0.8 |
| 49 | -1.5 | 1.8 | -2.8 | -3.5 | 0.1 |
| 50 | -1.5 | 1.9 | -1.4 | -3.4 | 1.0 |
| 51 | -1.3 | 1.9 | 0.0 | -3.2 | 1.8 |
| 52 | -1.0 | 1.9 | 1.4 | -3.1 | 2.7 |
| 53 | -0.5 | 2.0 | 2.5 | -2.9 | 3.1 |
| 54 | 0.5 | 2.0 | 3.7 | -2.8 | 3.5 |
| 55 | 1.4 | 2.3 | 4.8 | -2.7 | 3.9 |
| 56 | 1.7 | 2.7 | 6.0 | -2.6 | 4.3 |
| 57 | 1.9 | 3.0 | 6.6 | -2.6 | 4.4 |
| 58 | 2.3 | 3.4 | 7.1 | -2.6 | 4.6 |
| 59 | 3.0 | 3.7 | 7.7 | -2.6 | 4.7 |
| 60 | 3.8 | 4.1 | 8.2 | -2.6 | 4.8 |
| 61 | 4.5 | 4.4 | 8.4 | -2.5 | 4.8 |
| 62 | 4.9 | 4.8 | 8.6 | -2.5 | 4.9 |
| 63 | 4.9 | 5.0 | 8.7 | -2.5 | 4.9 |
| 64 | 5.0 | 5.0 | 8.9 | -2.5 | 4.9 |

| | | | | | |
|---|---|---|---|---|---|
| 65 | 5.3 | 5.1 | 9.4 | -2.3 | 5.1 |
| 66 | 5.7 | 5.1 | 9.8 | -2.2 | 5.3 |
| 67 | 6.0 | 5.3 | 10.3 | -2.1 | 5.4 |
| 68 | 6.2 | 5.7 | 10.7 | -1.9 | 5.6 |
| 69 | 6.7 | 6.0 | 10.8 | -1.6 | 5.7 |
| 70 | 7.0 | 6.4 | 10.9 | -1.2 | 5.8 |
| 71 | 7.9 | 6.8 | 10.9 | -0.8 | 5.9 |
| 72 | 8.3 | 7.0 | 11.0 | -0.5 | 5.9 |
| 73 | 8.6 | 7.3 | 11.2 | 0.2 | 6.6 |
| 74 | 9.3 | 7.5 | 11.5 | 1.7 | 7.2 |
| 75 | 10.6 | 7.8 | 11.7 | 3.2 | 7.8 |
| 76 | 10.8 | 9.1 | 11.9 | 4.8 | 8.5 |
| 77 | 11.2 | 10.4 | 13.6 | 6.3 | 9.0 |
| 78 | 12.1 | 11.7 | 15.2 | 7.6 | 9.5 |
| 79 | 12.7 | 13.0 | 16.9 | 8.9 | 10.1 |
| 80 | 13.4 | 13.7 | 18.6 | 10.2 | 10.6 |
| 81 | 14.4 | 14.3 | 18.8 | 11.5 | 11.5 |
| 82 | 14.6 | 14.9 | 19.1 | 12.6 | 12.5 |
| 83 | 15.3 | 15.5 | 19.4 | 13.0 | 13.4 |
| 84 | 15.7 | 16.3 | 19.6 | 13.5 | 14.4 |
| 85 | 16.2 | 17.2 | 20.2 | 13.9 | 14.7 |
| 86 | 18.7 | 18.2 | 20.8 | 14.3 | 15.0 |
| 87 | 19.6 | 19.1 | 21.3 | 14.6 | 15.4 |
| 88 | 19.7 | 19.6 | 21.9 | 14.7 | 15.7 |
| 89 | 20.1 | 19.7 | 22.3 | 14.9 | 18.5 |
| 90 | 22.1 | 19.8 | 22.6 | 15.1 | 21.3 |
| 91 | 23.6 | 19.9 | 22.9 | 15.8 | 24.0 |
| 92 | 26.8 | 20.7 | 23.3 | 22.2 | 26.8 |
| 93 | 27.7 | 23.1 | 29.3 | 28.5 | 27.0 |
| 94 | 30.0 | 25.4 | 35.2 | 34.9 | 27.2 |
| 95 | 34.8 | 27.7 | 41.2 | 41.2 | 27.4 |
| 96 | 44.3 | 30.8 | 47.2 | 45.8 | 27.6 |
| 97 | 47.5 | 37.6 | 50.0 | 48.9 | 29.3 |
| 98 | 58.0 | 44.4 | 52.8 | 52.0 | 31.0 |
| 99 | 58.2 | 51.2 | 55.6 | 55.1 | 32.6 |
| 100 | 58.4 | 58.0 | 58.4 | 58.2 | 34.3 |

***Table S53:*** *Paced tapping with music – Accuracy (vector direction degrees), music 2 – Paced_music_ross_mean_vector_direction*

| | | | | | | |
|---|---|---|---|---|---|---|
| **Task** | Paced tapping with music | | | | | |
| **Outcome measure** | Accuracy (vector direction degrees), music 2 | | | | | |
| **Variable name** | Paced_music_ross_mean_vector_direction | | | | | |
| **Age and gender effects** | | | | | | |
| Age regression (slope) | | 0.064 | | | | |
| Age regression (p) | | 0.588 | | | | |
| Age group (p) | | 0.786 | | | | |
| Gender (p) | | 0.251 | | | | |
| **Group** | | All | Age 18 to 21 | Age 22 to 29 | Age 30 to 54 | Age 55 to 87 |
| N (N female) | | 92 (63) | 20 (14) | 23 (15) | 25 (15) | 24 (19) |
| **Normality** | | | | | | |
| Skewness | | -1.33 | -1.54 | -0.48 | -1.30 | -0.69 |
| Excess kurtosis | | 2.95 | 2.31 | -0.52 | 2.49 | 0.15 |
| **Scores** | | | | | | |
| Mean | | -14.7 | -20.4 | -14.4 | -11.3 | -13.7 |
| SD | | 28.1 | 35.8 | 27.3 | 31.1 | 17.5 |
| Percentiles | | | | | | |
| | 0 | -125.4 | -125.4 | -66.2 | -110.6 | -57.0 |
| | 1 | -111.9 | -118.7 | -65.9 | -99.5 | -54.3 |
| | 2 | -93.6 | -111.9 | -65.7 | -88.4 | -51.6 |
| | 3 | -72.6 | -105.2 | -65.5 | -77.4 | -48.9 |
| | 4 | -65.5 | -98.4 | -65.3 | -66.3 | -46.2 |
| | 5 | -64.8 | -91.7 | -64.8 | -60.1 | -44.4 |
| | 6 | -63.2 | -85.4 | -64.0 | -54.9 | -43.1 |
| | 7 | -60.2 | -79.3 | -63.3 | -49.6 | -41.8 |
| | 8 | -57.6 | -73.2 | -62.5 | -44.4 | -40.5 |
| | 9 | -55.9 | -67.1 | -61.7 | -42.4 | -39.1 |
| | 10 | -50.8 | -61.0 | -59.6 | -41.9 | -37.6 |
| | 11 | -45.2 | -55.5 | -57.3 | -41.5 | -36.1 |
| | 12 | -42.9 | -50.5 | -55.1 | -41.1 | -34.6 |
| | 13 | -41.1 | -45.5 | -52.9 | -40.0 | -33.1 |
| | 14 | -39.9 | -40.6 | -49.7 | -38.4 | -31.7 |
| | 15 | -36.1 | -35.6 | -45.0 | -36.8 | -30.3 |
| | 16 | -33.5 | -31.6 | -40.4 | -35.3 | -28.9 |
| | 17 | -32.4 | -30.9 | -35.7 | -33.3 | -27.5 |
| | 18 | -31.1 | -30.3 | -31.0 | -30.5 | -26.6 |
| | 19 | -29.6 | -29.6 | -29.7 | -27.8 | -26.0 |
| | 20 | -28.2 | -29.0 | -29.2 | -25.1 | -25.5 |
| | 21 | -27.6 | -28.3 | -28.6 | -22.6 | -24.9 |

| | | | | | |
|---|---|---|---|---|---|
| 22 | -26.9 | -27.8 | -28.1 | -21.8 | -24.4 |
| 23 | -25.4 | -27.2 | -27.6 | -20.9 | -24.1 |
| 24 | -25.2 | -26.6 | -27.0 | -20.0 | -23.8 |
| 25 | -25.0 | -26.1 | -26.4 | -19.1 | -23.4 |
| 26 | -24.6 | -25.5 | -25.9 | -18.8 | -23.1 |
| 27 | -23.9 | -25.1 | -25.3 | -18.6 | -22.1 |
| 28 | -23.2 | -24.7 | -25.1 | -18.3 | -20.9 |
| 29 | -23.0 | -24.3 | -25.1 | -18.0 | -19.8 |
| 30 | -22.1 | -24.0 | -25.0 | -17.9 | -18.7 |
| 31 | -20.2 | -23.6 | -25.0 | -17.8 | -18.0 |
| 32 | -19.0 | -23.2 | -24.6 | -17.6 | -17.5 |
| 33 | -18.2 | -22.6 | -22.7 | -17.5 | -17.1 |
| 34 | -18.1 | -22.1 | -20.9 | -17.3 | -16.7 |
| 35 | -18.0 | -21.5 | -19.0 | -16.9 | -16.1 |
| 36 | -17.8 | -21.0 | -17.1 | -16.6 | -15.2 |
| 37 | -17.5 | -20.5 | -16.3 | -16.3 | -14.2 |
| 38 | -16.9 | -20.0 | -16.2 | -15.5 | -13.2 |
| 39 | -16.4 | -19.6 | -16.0 | -14.2 | -12.3 |
| 40 | -16.3 | -19.1 | -15.8 | -12.9 | -12.1 |
| 41 | -16.0 | -18.6 | -15.7 | -11.5 | -12.1 |
| 42 | -15.5 | -18.2 | -15.5 | -10.6 | -12.0 |
| 43 | -14.6 | -18.1 | -15.3 | -10.5 | -12.0 |
| 44 | -12.2 | -18.0 | -15.2 | -10.3 | -11.7 |
| 45 | -12.0 | -17.9 | -15.0 | -10.2 | -11.3 |
| 46 | -10.8 | -17.8 | -14.1 | -9.9 | -11.0 |
| 47 | -10.3 | -17.8 | -12.6 | -9.1 | -10.6 |
| 48 | -10.1 | -16.6 | -11.0 | -8.3 | -10.2 |
| 49 | -10.0 | -14.8 | -9.5 | -7.4 | -10.2 |
| 50 | -9.7 | -13.0 | -7.9 | -6.6 | -10.1 |
| 51 | -8.9 | -11.2 | -7.8 | -6.0 | -10.1 |
| 52 | -8.1 | -9.3 | -7.7 | -5.4 | -10.0 |
| 53 | -7.8 | -8.1 | -7.6 | -4.7 | -9.9 |
| 54 | -7.6 | -8.0 | -7.5 | -4.1 | -9.7 |
| 55 | -7.5 | -7.9 | -7.4 | -3.8 | -9.6 |
| 56 | -7.4 | -7.7 | -7.4 | -3.7 | -9.4 |
| 57 | -7.3 | -7.6 | -7.3 | -3.5 | -9.2 |
| 58 | -7.2 | -7.4 | -7.3 | -3.3 | -8.8 |
| 59 | -6.8 | -7.0 | -7.3 | -2.6 | -8.4 |
| 60 | -6.0 | -6.5 | -7.3 | -1.6 | -8.0 |
| 61 | -5.4 | -6.1 | -7.2 | -0.7 | -7.6 |
| 62 | -5.0 | -5.6 | -7.2 | 0.3 | -7.1 |
| 63 | -4.9 | -5.1 | -7.2 | 1.1 | -6.7 |
| 64 | -4.7 | -5.0 | -7.0 | 1.6 | -6.2 |

| | | | | | |
|---|---|---|---|---|---|
| 65 | -4.4 | -5.0 | -6.4 | 2.2 | -5.8 |
| 66 | -4.0 | -4.9 | -5.8 | 2.7 | -5.5 |
| 67 | -3.2 | -4.9 | -5.2 | 3.3 | -5.4 |
| 68 | -3.0 | -4.8 | -4.5 | 3.8 | -5.2 |
| 69 | -2.9 | -4.3 | -3.9 | 4.4 | -5.1 |
| 70 | -2.0 | -3.4 | -3.3 | 4.9 | -4.8 |
| 71 | -1.6 | -2.6 | -2.7 | 5.4 | -4.3 |
| 72 | -1.0 | -1.7 | -2.1 | 5.6 | -3.9 |
| 73 | -0.4 | -0.9 | -1.1 | 5.8 | -3.4 |
| 74 | 0.1 | -0.2 | 1.1 | 6.0 | -3.0 |
| 75 | 1.0 | 0.2 | 3.3 | 6.2 | -2.9 |
| 76 | 1.8 | 0.5 | 5.5 | 6.8 | -2.9 |
| 77 | 3.0 | 0.9 | 7.7 | 7.4 | -2.9 |
| 78 | 3.1 | 1.2 | 8.4 | 8.0 | -2.9 |
| 79 | 5.1 | 1.6 | 8.5 | 8.6 | -2.6 |
| 80 | 6.0 | 1.8 | 8.6 | 9.0 | -2.3 |
| 81 | 6.6 | 2.1 | 8.7 | 9.4 | -2.0 |
| 82 | 7.7 | 2.4 | 9.1 | 9.8 | -1.7 |
| 83 | 8.5 | 2.7 | 10.7 | 10.1 | -1.4 |
| 84 | 8.8 | 2.9 | 12.3 | 10.6 | -1.2 |
| 85 | 9.1 | 3.6 | 13.9 | 11.0 | -1.0 |
| 86 | 9.7 | 4.3 | 15.4 | 11.5 | -0.8 |
| 87 | 10.4 | 5.0 | 16.7 | 11.9 | -0.5 |
| 88 | 11.4 | 5.7 | 17.9 | 13.9 | 1.8 |
| 89 | 12.1 | 6.5 | 19.0 | 17.4 | 4.2 |
| 90 | 13.6 | 7.5 | 20.2 | 21.0 | 6.5 |
| 91 | 14.0 | 8.8 | 21.3 | 24.5 | 8.8 |
| 92 | 15.5 | 10.1 | 21.7 | 26.9 | 9.8 |
| 93 | 19.3 | 11.4 | 22.1 | 27.0 | 10.2 |
| 94 | 22.3 | 12.8 | 22.5 | 27.2 | 10.6 |
| 95 | 24.8 | 14.6 | 22.9 | 27.3 | 11.0 |
| 96 | 27.0 | 18.0 | 23.6 | 27.8 | 11.5 |
| 97 | 27.3 | 21.4 | 24.5 | 30.3 | 12.2 |
| 98 | 28.2 | 24.8 | 25.4 | 32.8 | 12.8 |
| 99 | 32.2 | 28.2 | 26.3 | 35.3 | 13.5 |
| 100 | 37.8 | 31.6 | 27.2 | 37.8 | 14.1 |

***Table S54:*** *Synchronization-continuation – Rate (ITI in ms), fast tempo – SyncCont_metro_450_mean_mean_iti*

| **Task** | Synchronization-continuation | | | | |
|---|---|---|---|---|---|
| **Outcome measure** | Rate (ITI in ms), fast tempo | | | | |
| **Variable name** | SyncCont_metro_450_mean_mean_iti | | | | |
| **Age and gender effects** | | | | | |
| Age regression (slope) | 0.255 | | | | |
| Age regression (p) | 0.003 | | | | |
| Age group (p) | 0.107 | | | | |
| Gender (p) | 0.986 | | | | |
| **Group** | All | Age 18 to 21 | Age 22 to 29 | Age 30 to 54 | Age 55 to 87 |
| N (N female) | 108 (74) | 27 (18) | 29 (19) | 26 (16) | 26 (21) |
| **Normality** | | | | | |
| Skewness | 0.17 | 2.09 | -1.02 | -1.32 | 0.13 |
| Excess kurtosis | 4.10 | 6.26 | 0.88 | 2.79 | -0.98 |
| **Scores** | | | | | |
| Mean | 448.8 | 451.1 | 443.4 | 444.2 | 457.3 |
| SD | 21.6 | 24.6 | 21.0 | 21.1 | 17.0 |
| Percentiles | | | | | |
| 0 | 374.6 | 417.4 | 387.2 | 374.6 | 428.8 |
| 1 | 387.7 | 417.9 | 389.4 | 383.7 | 429.5 |
| 2 | 397.0 | 418.4 | 391.7 | 392.7 | 430.1 |
| 3 | 408.4 | 419.0 | 393.9 | 401.8 | 430.8 |
| 4 | 412.7 | 419.6 | 396.7 | 410.9 | 431.4 |
| 5 | 418.1 | 420.9 | 400.2 | 413.3 | 431.5 |
| 6 | 419.8 | 422.2 | 403.8 | 415.8 | 431.6 |
| 7 | 420.6 | 423.6 | 407.3 | 418.2 | 431.7 |
| 8 | 422.1 | 425.1 | 410.8 | 420.7 | 431.8 |
| 9 | 423.4 | 427.1 | 414.4 | 421.4 | 433.3 |
| 10 | 424.2 | 429.1 | 417.9 | 422.1 | 434.7 |
| 11 | 427.8 | 431.2 | 420.7 | 422.8 | 436.2 |
| 12 | 430.2 | 432.5 | 421.5 | 423.5 | 437.6 |
| 13 | 430.5 | 432.9 | 422.2 | 425.3 | 438.7 |
| 14 | 431.4 | 433.4 | 423.0 | 427.0 | 439.8 |
| 15 | 431.9 | 433.9 | 424.7 | 428.8 | 440.9 |
| 16 | 432.3 | 434.2 | 426.7 | 430.5 | 442.0 |
| 17 | 432.8 | 434.5 | 428.7 | 431.5 | 442.5 |
| 18 | 433.3 | 434.8 | 430.5 | 432.6 | 443.0 |
| 19 | 434.2 | 435.0 | 431.2 | 433.6 | 443.5 |
| 20 | 434.7 | 435.3 | 431.8 | 434.6 | 444.0 |
| 21 | 435.0 | 435.6 | 432.5 | 434.7 | 444.0 |
| 22 | 435.6 | 435.8 | 432.8 | 434.7 | 444.1 |

| | | | | | |
|---|---|---|---|---|---|
| 23 | 436.6 | 436.1 | 432.9 | 434.8 | 444.1 |
| 24 | 437.1 | 436.3 | 432.9 | 434.8 | 444.2 |
| 25 | 437.5 | 436.5 | 433.0 | 435.8 | 444.7 |
| 26 | 438.4 | 436.7 | 434.2 | 436.7 | 445.3 |
| 27 | 438.6 | 436.9 | 435.4 | 437.7 | 445.8 |
| 28 | 439.6 | 437.6 | 436.5 | 438.6 | 446.4 |
| 29 | 439.9 | 438.4 | 437.4 | 439.0 | 446.4 |
| 30 | 440.1 | 439.1 | 437.7 | 439.3 | 446.4 |
| 31 | 441.3 | 439.9 | 438.1 | 439.6 | 446.5 |
| 32 | 442.1 | 440.9 | 438.5 | 439.9 | 446.5 |
| 33 | 442.7 | 441.9 | 439.5 | 439.9 | 447.2 |
| 34 | 443.7 | 442.9 | 440.6 | 439.9 | 447.8 |
| 35 | 444.1 | 443.6 | 441.6 | 440.0 | 448.4 |
| 36 | 444.3 | 443.8 | 442.6 | 440.0 | 449.1 |
| 37 | 444.6 | 444.0 | 443.3 | 440.3 | 450.0 |
| 38 | 445.0 | 444.2 | 443.9 | 440.6 | 451.0 |
| 39 | 445.8 | 444.5 | 444.6 | 440.9 | 452.0 |
| 40 | 446.3 | 444.7 | 445.1 | 441.2 | 452.9 |
| 41 | 446.4 | 444.9 | 445.5 | 442.7 | 453.0 |
| 42 | 446.5 | 445.1 | 445.9 | 444.2 | 453.0 |
| 43 | 447.2 | 445.3 | 446.4 | 445.7 | 453.1 |
| 44 | 447.3 | 445.6 | 446.7 | 447.2 | 453.1 |
| 45 | 447.4 | 445.8 | 447.1 | 447.2 | 453.3 |
| 46 | 447.7 | 446.0 | 447.5 | 447.2 | 453.5 |
| 47 | 448.0 | 446.3 | 448.0 | 447.3 | 453.7 |
| 48 | 448.3 | 446.7 | 448.6 | 447.3 | 453.9 |
| 49 | 448.7 | 447.0 | 449.2 | 447.4 | 455.0 |
| 50 | 449.4 | 447.3 | 449.8 | 447.6 | 456.0 |
| 51 | 450.0 | 447.5 | 450.3 | 447.7 | 457.1 |
| 52 | 450.8 | 447.8 | 450.8 | 447.9 | 458.2 |
| 53 | 451.4 | 448.0 | 451.3 | 448.0 | 458.6 |
| 54 | 452.6 | 448.4 | 451.7 | 448.2 | 459.0 |
| 55 | 452.9 | 449.6 | 452.2 | 448.3 | 459.4 |
| 56 | 452.9 | 450.8 | 452.7 | 448.5 | 459.8 |
| 57 | 453.1 | 452.0 | 453.1 | 448.9 | 460.4 |
| 58 | 453.2 | 452.9 | 453.2 | 449.4 | 461.1 |
| 59 | 453.4 | 453.0 | 453.3 | 449.8 | 461.7 |
| 60 | 453.4 | 453.2 | 453.3 | 450.2 | 462.4 |
| 61 | 453.4 | 453.3 | 453.4 | 450.4 | 462.4 |
| 62 | 453.6 | 453.4 | 453.4 | 450.6 | 462.4 |
| 63 | 454.4 | 453.4 | 453.4 | 450.8 | 462.4 |
| 64 | 455.5 | 453.4 | 453.4 | 451.1 | 462.4 |
| 65 | 455.9 | 453.4 | 453.8 | 451.5 | 462.5 |
| 66 | 456.1 | 453.8 | 454.3 | 452.0 | 462.5 |

| | | | | | |
|---|---|---|---|---|---|
| 67 | 456.4 | 454.4 | 454.8 | 452.4 | 462.6 |
| 68 | 456.5 | 455.1 | 455.2 | 452.9 | 462.6 |
| 69 | 457.0 | 455.7 | 455.4 | 453.9 | 463.7 |
| 70 | 457.3 | 456.4 | 455.6 | 455.0 | 464.8 |
| 71 | 458.2 | 457.2 | 455.8 | 456.0 | 465.9 |
| 72 | 458.7 | 457.9 | 456.0 | 457.1 | 467.0 |
| 73 | 459.3 | 458.6 | 456.1 | 457.8 | 467.3 |
| 74 | 459.6 | 458.9 | 456.2 | 458.5 | 467.7 |
| 75 | 459.8 | 459.1 | 456.3 | 459.3 | 468.0 |
| 76 | 459.9 | 459.4 | 456.3 | 460.0 | 468.4 |
| 77 | 460.1 | 459.6 | 456.3 | 460.1 | 469.6 |
| 78 | 460.6 | 460.6 | 456.4 | 460.2 | 470.7 |
| 79 | 461.3 | 461.6 | 456.4 | 460.3 | 471.8 |
| 80 | 462.1 | 462.6 | 456.4 | 460.4 | 473.0 |
| 81 | 462.4 | 463.4 | 456.5 | 460.5 | 474.6 |
| 82 | 462.6 | 463.4 | 456.5 | 460.6 | 476.1 |
| 83 | 463.2 | 463.4 | 456.7 | 460.7 | 477.7 |
| 84 | 463.4 | 463.4 | 456.9 | 460.8 | 479.2 |
| 85 | 465.6 | 463.7 | 457.1 | 461.0 | 480.0 |
| 86 | 466.1 | 464.3 | 457.4 | 461.2 | 480.7 |
| 87 | 467.0 | 464.9 | 458.0 | 461.5 | 481.4 |
| 88 | 467.4 | 465.5 | 458.6 | 461.7 | 482.2 |
| 89 | 468.4 | 466.1 | 459.1 | 462.8 | 482.3 |
| 90 | 469.8 | 466.8 | 459.4 | 463.9 | 482.5 |
| 91 | 474.1 | 467.5 | 459.6 | 465.0 | 482.6 |
| 92 | 476.7 | 468.2 | 459.7 | 466.1 | 482.8 |
| 93 | 478.6 | 471.9 | 460.5 | 466.4 | 483.3 |
| 94 | 480.0 | 477.0 | 465.0 | 466.7 | 483.8 |
| 95 | 481.6 | 482.1 | 469.5 | 466.9 | 484.2 |
| 96 | 482.6 | 487.2 | 473.9 | 467.2 | 484.7 |
| 97 | 484.3 | 500.7 | 476.2 | 470.6 | 484.8 |
| 98 | 485.0 | 515.7 | 476.7 | 473.9 | 484.9 |
| 99 | 487.8 | 530.7 | 477.3 | 477.2 | 484.9 |
| 100 | 545.7 | 545.7 | 477.9 | 480.6 | 485.0 |

***Table S55:*** *Synchronization-continuation – Rate (ITI in ms), fast tempo – SyncCont_metro_450_mean_mean_iti*

| **Task** | Synchronization-continuation | | | | |
|---|---|---|---|---|---|
| **Outcome measure** | Rate (ITI in ms), medium tempo | | | | |
| **Variable name** | SyncCont_metro_600_mean_mean_iti | | | | |
| **Age and gender effects** | | | | | |
| Age regression (slope) | 0.115 | | | | |
| Age regression (p) | 0.286 | | | | |
| Age group (p) | 0.839 | | | | |
| Gender (p) | 0.539 | | | | |
| **Group** | All | Age 18 to 21 | Age 22 to 29 | Age 30 to 54 | Age 55 to 87 |
| N (N female) | 108 (74) | 27 (18) | 29 (19) | 26 (16) | 26 (21) |
| **Normality** | | | | | |
| Skewness | 0.43 | 2.32 | -0.80 | 0.04 | 0.65 |
| Excess kurtosis | 3.74 | 7.14 | 0.46 | -0.34 | -0.03 |
| **Scores** | | | | | |
| Mean | 596.5 | 597.7 | 592.7 | 594.5 | 601.6 |
| SD | 26.1 | 29.2 | 31.7 | 19.7 | 21.3 |
| Percentiles | | | | | |
| 0 | 511.7 | 564.4 | 511.7 | 553.7 | 563.0 |
| 1 | 536.0 | 565.4 | 518.5 | 556.0 | 565.6 |
| 2 | 538.1 | 566.3 | 525.2 | 558.4 | 568.1 |
| 3 | 546.2 | 567.3 | 532.0 | 560.7 | 570.7 |
| 4 | 553.5 | 568.1 | 536.0 | 563.1 | 573.2 |
| 5 | 554.9 | 568.2 | 536.4 | 565.1 | 575.0 |
| 6 | 559.5 | 568.3 | 536.7 | 567.2 | 576.8 |
| 7 | 563.0 | 568.4 | 537.0 | 569.3 | 578.7 |
| 8 | 563.8 | 568.7 | 538.8 | 571.4 | 580.5 |
| 9 | 566.7 | 569.4 | 540.8 | 571.4 | 580.6 |
| 10 | 568.4 | 570.0 | 542.9 | 571.4 | 580.7 |
| 11 | 570.4 | 570.6 | 545.0 | 571.4 | 580.7 |
| 12 | 571.3 | 571.2 | 547.6 | 571.4 | 580.8 |
| 13 | 571.4 | 571.8 | 550.1 | 573.1 | 581.9 |
| 14 | 571.7 | 572.4 | 552.6 | 574.7 | 583.0 |
| 15 | 573.2 | 573.0 | 554.1 | 576.3 | 584.0 |
| 16 | 573.5 | 573.5 | 555.1 | 577.9 | 585.1 |
| 17 | 575.3 | 574.0 | 556.1 | 578.5 | 585.5 |
| 18 | 577.1 | 574.4 | 557.6 | 579.1 | 586.0 |
| 19 | 578.7 | 574.9 | 561.7 | 579.7 | 586.4 |
| 20 | 580.4 | 575.3 | 565.8 | 580.3 | 586.8 |
| 21 | 580.5 | 575.8 | 570.0 | 580.3 | 587.2 |
| 22 | 580.6 | 576.3 | 574.6 | 580.4 | 587.6 |

| | | | | | |
|---|---|---|---|---|---|
| 23 | 580.8 | 576.7 | 579.7 | 580.4 | 588.0 |
| 24 | 581.0 | 577.7 | 584.8 | 580.5 | 588.4 |
| 25 | 581.1 | 578.8 | 589.9 | 580.6 | 588.8 |
| 26 | 583.3 | 579.8 | 590.1 | 580.8 | 589.1 |
| 27 | 585.0 | 580.8 | 590.2 | 580.9 | 589.5 |
| 28 | 585.8 | 581.6 | 590.4 | 581.0 | 589.9 |
| 29 | 586.2 | 582.4 | 590.8 | 581.1 | 590.0 |
| 30 | 586.8 | 583.2 | 591.8 | 581.1 | 590.1 |
| 31 | 587.2 | 583.9 | 592.7 | 581.1 | 590.3 |
| 32 | 588.0 | 584.4 | 593.7 | 581.1 | 590.4 |
| 33 | 588.4 | 585.0 | 595.1 | 582.4 | 590.5 |
| 34 | 589.0 | 585.5 | 596.5 | 583.6 | 590.7 |
| 35 | 589.9 | 586.0 | 598.0 | 584.9 | 590.8 |
| 36 | 590.1 | 586.3 | 599.1 | 586.1 | 590.9 |
| 37 | 590.4 | 586.6 | 599.3 | 586.6 | 591.9 |
| 38 | 590.8 | 586.9 | 599.6 | 587.0 | 592.8 |
| 39 | 591.7 | 587.2 | 599.8 | 587.5 | 593.7 |
| 40 | 592.9 | 587.6 | 599.9 | 587.9 | 594.6 |
| 41 | 593.7 | 587.9 | 600.0 | 588.9 | 594.8 |
| 42 | 593.8 | 588.3 | 600.1 | 589.9 | 595.0 |
| 43 | 594.6 | 589.4 | 600.2 | 590.9 | 595.1 |
| 44 | 595.3 | 590.8 | 600.2 | 592.0 | 595.3 |
| 45 | 595.6 | 592.2 | 600.3 | 592.2 | 595.4 |
| 46 | 596.6 | 593.6 | 600.4 | 592.5 | 595.4 |
| 47 | 597.9 | 595.1 | 600.8 | 592.8 | 595.4 |
| 48 | 599.2 | 596.6 | 601.4 | 593.1 | 595.4 |
| 49 | 599.5 | 598.0 | 602.1 | 594.2 | 595.7 |
| 50 | 599.7 | 599.5 | 602.8 | 595.3 | 595.9 |
| 51 | 600.0 | 599.5 | 603.0 | 596.4 | 596.2 |
| 52 | 600.2 | 599.6 | 603.3 | 597.5 | 596.4 |
| 53 | 600.3 | 599.6 | 603.5 | 598.2 | 597.4 |
| 54 | 600.5 | 599.8 | 603.7 | 598.8 | 598.5 |
| 55 | 600.9 | 600.6 | 603.7 | 599.5 | 599.5 |
| 56 | 602.2 | 601.5 | 603.8 | 600.2 | 600.5 |
| 57 | 602.6 | 602.4 | 603.9 | 600.4 | 601.0 |
| 58 | 602.7 | 603.1 | 604.0 | 600.5 | 601.4 |
| 59 | 602.8 | 603.3 | 604.1 | 600.7 | 601.9 |
| 60 | 603.1 | 603.4 | 604.3 | 600.9 | 602.4 |
| 61 | 603.5 | 603.6 | 604.4 | 601.4 | 602.4 |
| 62 | 603.7 | 603.8 | 604.5 | 601.8 | 602.5 |
| 63 | 603.8 | 603.8 | 604.6 | 602.3 | 602.6 |
| 64 | 603.9 | 603.9 | 604.8 | 602.7 | 602.6 |
| 65 | 604.1 | 604.0 | 604.8 | 602.9 | 603.0 |
| 66 | 604.2 | 604.0 | 604.8 | 603.1 | 603.4 |

| | | | | | |
|---|---|---|---|---|---|
| 67 | 604.3 | 604.0 | 604.9 | 603.2 | 603.8 |
| 68 | 604.6 | 604.1 | 604.9 | 603.4 | 604.2 |
| 69 | 604.8 | 604.1 | 605.2 | 603.8 | 604.3 |
| 70 | 604.8 | 604.3 | 605.5 | 604.3 | 604.4 |
| 71 | 604.8 | 604.4 | 605.7 | 604.7 | 604.5 |
| 72 | 604.9 | 604.6 | 606.3 | 605.1 | 604.6 |
| 73 | 604.9 | 604.8 | 607.1 | 605.9 | 608.1 |
| 74 | 605.2 | 604.8 | 607.9 | 606.8 | 611.6 |
| 75 | 606.4 | 604.8 | 608.7 | 607.6 | 615.1 |
| 76 | 608.1 | 604.8 | 608.8 | 608.4 | 618.6 |
| 77 | 608.5 | 604.8 | 608.9 | 609.0 | 618.8 |
| 78 | 608.9 | 604.8 | 609.1 | 609.5 | 618.9 |
| 79 | 609.6 | 604.9 | 609.6 | 610.1 | 619.1 |
| 80 | 610.4 | 604.9 | 610.6 | 610.6 | 619.2 |
| 81 | 612.2 | 605.1 | 611.7 | 611.2 | 620.5 |
| 82 | 612.9 | 605.9 | 612.8 | 611.8 | 621.8 |
| 83 | 614.3 | 606.6 | 613.8 | 612.4 | 623.1 |
| 84 | 616.5 | 607.4 | 614.9 | 612.9 | 624.4 |
| 85 | 617.8 | 608.1 | 616.0 | 613.3 | 625.2 |
| 86 | 618.6 | 608.7 | 616.8 | 613.8 | 625.9 |
| 87 | 619.1 | 609.2 | 617.1 | 614.2 | 626.7 |
| 88 | 619.5 | 609.8 | 617.4 | 614.6 | 627.4 |
| 89 | 620.8 | 611.7 | 617.8 | 616.1 | 628.5 |
| 90 | 621.1 | 614.6 | 618.1 | 617.7 | 629.5 |
| 91 | 622.6 | 617.6 | 618.4 | 619.2 | 630.6 |
| 92 | 625.7 | 620.5 | 618.8 | 620.7 | 631.7 |
| 93 | 628.4 | 624.9 | 619.5 | 620.8 | 634.4 |
| 94 | 630.7 | 629.8 | 622.4 | 620.8 | 637.2 |
| 95 | 636.2 | 634.7 | 625.3 | 620.9 | 639.9 |
| 96 | 639.9 | 639.7 | 628.2 | 621.0 | 642.7 |
| 97 | 642.2 | 656.4 | 634.1 | 625.4 | 645.2 |
| 98 | 651.2 | 675.3 | 642.2 | 629.8 | 647.6 |
| 99 | 657.9 | 694.2 | 650.3 | 634.2 | 650.1 |
| 100 | 713.1 | 713.1 | 658.4 | 638.6 | 652.6 |

***Table S56:*** *Synchronization-continuation – Rate (ITI in ms), slow tempo – SyncCont_metro_750_mean_mean_iti*

| | | | | | | |
|---|---|---|---|---|---|---|
| **Task** | | Synchronization-continuation | | | | |
| **Outcome measure** | | Rate (ITI in ms), slow tempo | | | | |
| **Variable name** | | SyncCont_metro_750_mean_mean_iti | | | | |
| **Age and gender effects** | | | | | | |
| Age regression (slope) | | -0.275 | | | | |
| Age regression (p) | | 0.177 | | | | |
| Age group (p) | | 0.392 | | | | |
| Gender (p) | | 0.990 | | | | |
| **Group** | | All | Age 18 to 21 | Age 22 to 29 | Age 30 to 54 | Age 55 to 87 |
| N (N female) | | 108 (74) | 27 (18) | 29 (19) | 26 (16) | 26 (21) |
| **Normality** | | | | | | |
| Skewness | | -0.28 | 0.21 | -0.53 | -0.31 | 0.34 |
| Excess kurtosis | | 0.61 | -0.25 | -0.55 | 0.27 | -0.92 |
| **Scores** | | | | | | |
| Mean | | 749.3 | 761.2 | 749.2 | 742.7 | 743.9 |
| SD | | 40.2 | 36.3 | 37.0 | 52.9 | 31.4 |
| Percentiles | | | | | | |
| | 0 | 609.0 | 687.9 | 672.1 | 609.0 | 685.4 |
| | 1 | 663.0 | 693.8 | 672.8 | 622.4 | 690.3 |
| | 2 | 669.7 | 699.6 | 673.5 | 635.7 | 695.2 |
| | 3 | 672.6 | 705.5 | 674.2 | 649.1 | 700.2 |
| | 4 | 677.6 | 710.7 | 676.6 | 662.5 | 705.1 |
| | 5 | 686.3 | 712.4 | 681.1 | 664.2 | 707.5 |
| | 6 | 689.1 | 714.1 | 685.6 | 665.9 | 709.8 |
| | 7 | 694.1 | 715.8 | 690.2 | 667.6 | 712.2 |
| | 8 | 698.3 | 717.1 | 692.8 | 669.3 | 714.6 |
| | 9 | 700.7 | 717.5 | 695.1 | 676.4 | 714.8 |
| | 10 | 703.1 | 717.9 | 697.3 | 683.4 | 715.0 |
| | 11 | 704.8 | 718.4 | 699.3 | 690.5 | 715.3 |
| | 12 | 706.7 | 718.9 | 700.7 | 697.6 | 715.5 |
| | 13 | 707.6 | 719.4 | 702.0 | 698.6 | 715.5 |
| | 14 | 709.3 | 720.0 | 703.3 | 699.6 | 715.5 |
| | 15 | 710.4 | 720.5 | 704.3 | 700.6 | 715.5 |
| | 16 | 710.6 | 722.0 | 705.3 | 701.7 | 715.5 |
| | 17 | 712.5 | 723.9 | 706.2 | 703.2 | 716.3 |
| | 18 | 714.8 | 725.9 | 707.4 | 704.7 | 717.1 |
| | 19 | 715.5 | 727.9 | 709.9 | 706.2 | 717.9 |
| | 20 | 715.7 | 730.6 | 712.4 | 707.7 | 718.7 |
| | 21 | 716.4 | 733.5 | 714.9 | 708.1 | 718.8 |
| | 22 | 717.8 | 736.5 | 718.2 | 708.5 | 718.9 |

| | | | | | |
|---|---|---|---|---|---|
| 23 | 718.7 | 739.4 | 722.0 | 708.9 | 719.1 |
| 24 | 719.0 | 739.8 | 725.9 | 709.3 | 719.2 |
| 25 | 720.2 | 740.0 | 729.8 | 709.6 | 719.5 |
| 26 | 720.7 | 740.2 | 733.2 | 709.9 | 719.9 |
| 27 | 722.3 | 740.4 | 736.6 | 710.2 | 720.2 |
| 28 | 728.1 | 740.7 | 739.9 | 710.4 | 720.5 |
| 29 | 729.8 | 741.0 | 741.9 | 710.8 | 721.0 |
| 30 | 730.2 | 741.2 | 742.0 | 711.2 | 721.5 |
| 31 | 730.9 | 741.7 | 742.1 | 711.7 | 722.0 |
| 32 | 731.8 | 743.0 | 742.1 | 712.1 | 722.5 |
| 33 | 733.5 | 744.3 | 742.2 | 716.7 | 724.4 |
| 34 | 734.5 | 745.5 | 742.3 | 721.4 | 726.3 |
| 35 | 735.0 | 746.4 | 742.4 | 726.1 | 728.2 |
| 36 | 735.8 | 746.6 | 742.8 | 730.8 | 730.2 |
| 37 | 738.2 | 746.9 | 743.9 | 731.4 | 730.5 |
| 38 | 740.1 | 747.1 | 745.0 | 732.0 | 730.8 |
| 39 | 741.2 | 748.8 | 746.1 | 732.5 | 731.1 |
| 40 | 741.8 | 751.5 | 746.6 | 733.1 | 731.4 |
| 41 | 742.0 | 754.3 | 746.7 | 735.3 | 732.2 |
| 42 | 742.1 | 757.1 | 746.9 | 737.5 | 732.9 |
| 43 | 742.5 | 758.0 | 747.1 | 739.8 | 733.7 |
| 44 | 742.9 | 758.1 | 748.1 | 742.0 | 734.4 |
| 45 | 746.3 | 758.1 | 749.1 | 743.1 | 734.5 |
| 46 | 746.5 | 758.2 | 750.0 | 744.3 | 734.5 |
| 47 | 746.7 | 758.7 | 751.3 | 745.5 | 734.6 |
| 48 | 747.1 | 759.2 | 752.7 | 746.6 | 734.6 |
| 49 | 748.6 | 759.8 | 754.2 | 748.5 | 734.8 |
| 50 | 750.5 | 760.3 | 755.6 | 750.4 | 735.0 |
| 51 | 752.6 | 760.9 | 757.3 | 752.3 | 735.2 |
| 52 | 754.6 | 761.4 | 759.0 | 754.2 | 735.4 |
| 53 | 755.4 | 761.9 | 760.7 | 754.4 | 735.6 |
| 54 | 756.7 | 762.4 | 761.8 | 754.5 | 735.8 |
| 55 | 757.8 | 762.4 | 761.9 | 754.6 | 736.0 |
| 56 | 758.2 | 762.4 | 762.1 | 754.8 | 736.2 |
| 57 | 760.1 | 762.5 | 762.3 | 756.4 | 737.8 |
| 58 | 760.4 | 762.5 | 762.8 | 758.1 | 739.4 |
| 59 | 761.5 | 762.7 | 763.4 | 759.8 | 741.0 |
| 60 | 761.6 | 762.8 | 764.0 | 761.4 | 742.6 |
| 61 | 761.9 | 763.0 | 764.8 | 761.5 | 744.6 |
| 62 | 762.3 | 765.0 | 765.9 | 761.5 | 746.6 |
| 63 | 762.4 | 769.1 | 767.1 | 761.5 | 748.5 |
| 64 | 762.4 | 773.3 | 768.3 | 761.5 | 750.5 |
| 65 | 762.8 | 777.4 | 768.9 | 761.7 | 752.1 |
| 66 | 763.4 | 779.8 | 769.4 | 761.9 | 753.7 |

| | | | | | |
|---|---|---|---|---|---|
| 67 | 764.2 | 781.0 | 769.8 | 762.1 | 755.4 |
| 68 | 767.6 | 782.3 | 770.3 | 762.3 | 757.0 |
| 69 | 769.6 | 783.5 | 770.5 | 762.6 | 757.8 |
| 70 | 770.2 | 784.2 | 770.8 | 762.9 | 758.6 |
| 71 | 771.2 | 784.7 | 771.1 | 763.3 | 759.3 |
| 72 | 772.8 | 785.2 | 771.5 | 763.6 | 760.1 |
| 73 | 773.1 | 785.7 | 771.9 | 765.1 | 763.7 |
| 74 | 774.4 | 786.3 | 772.3 | 766.7 | 767.2 |
| 75 | 776.4 | 786.9 | 772.8 | 768.3 | 770.7 |
| 76 | 779.0 | 787.4 | 773.5 | 769.9 | 774.2 |
| 77 | 780.0 | 788.0 | 774.3 | 770.6 | 776.0 |
| 78 | 782.6 | 788.3 | 775.1 | 771.4 | 777.9 |
| 79 | 784.5 | 788.7 | 775.9 | 772.2 | 779.7 |
| 80 | 785.5 | 789.1 | 776.9 | 773.0 | 781.5 |
| 81 | 786.1 | 789.5 | 777.8 | 777.2 | 782.4 |
| 82 | 787.5 | 790.3 | 778.8 | 781.4 | 783.3 |
| 83 | 789.1 | 791.1 | 780.7 | 785.7 | 784.2 |
| 84 | 789.3 | 791.8 | 782.8 | 789.9 | 785.1 |
| 85 | 789.8 | 792.7 | 784.8 | 793.3 | 786.1 |
| 86 | 789.9 | 793.9 | 786.6 | 796.7 | 787.2 |
| 87 | 790.5 | 795.1 | 787.6 | 800.1 | 788.3 |
| 88 | 791.0 | 796.3 | 788.6 | 803.5 | 789.3 |
| 89 | 792.5 | 797.2 | 789.6 | 805.6 | 789.6 |
| 90 | 794.2 | 797.9 | 790.5 | 807.7 | 789.9 |
| 91 | 797.8 | 798.6 | 791.4 | 809.8 | 790.2 |
| 92 | 800.6 | 799.3 | 792.3 | 812.0 | 790.5 |
| 93 | 802.8 | 806.2 | 793.5 | 814.6 | 790.5 |
| 94 | 803.6 | 815.8 | 796.4 | 817.3 | 790.6 |
| 95 | 806.7 | 825.4 | 799.4 | 819.9 | 790.7 |
| 96 | 810.9 | 835.1 | 802.3 | 822.6 | 790.7 |
| 97 | 820.3 | 836.8 | 804.4 | 829.3 | 793.6 |
| 98 | 834.6 | 837.1 | 805.7 | 836.0 | 796.4 |
| 99 | 837.5 | 837.3 | 807.0 | 842.7 | 799.2 |
| 100 | 849.4 | 837.6 | 808.3 | 849.4 | 802.0 |

***Table S57:*** *Synchronization-continuation – Variability (CV of ITI), fast tempo – SyncCont_metro_450_mean_CV_iti*

| | | | | | |
|---|---|---|---|---|---|
| **Task** | Synchronization-continuation | | | | |
| **Outcome measure** | Variability (CV of ITI), fast tempo | | | | |
| **Variable name** | SyncCont_metro_450_mean_CV_iti | | | | |
| **Age and gender effects** | | | | | |
| Age regression (slope) | 0.000 | | | | |
| Age regression (p) | 0.066 | | | | |
| Age group (p) | 0.172 | | | | |
| Gender (p) | 0.864 | | | | |
| **Group** | All | Age 18 to 21 | Age 22 to 29 | Age 30 to 54 | Age 55 to 87 |
| N (N female) | 108 (74) | 27 (18) | 29 (19) | 26 (16) | 26 (21) |
| **Normality** | | | | | |
| Skewness | 1.00 | 0.50 | 1.13 | 0.91 | 0.35 |
| Excess kurtosis | 0.58 | -0.32 | -0.04 | 0.16 | -0.77 |
| **Scores** | | | | | |
| Mean | 0.048 | 0.051 | 0.052 | 0.044 | 0.045 |
| SD | 0.015 | 0.016 | 0.018 | 0.013 | 0.012 |
| Percentiles | | | | | |
| 0 | 0.025 | 0.025 | 0.032 | 0.028 | 0.026 |
| 1 | 0.026 | 0.025 | 0.032 | 0.028 | 0.027 |
| 2 | 0.027 | 0.026 | 0.033 | 0.028 | 0.028 |
| 3 | 0.028 | 0.026 | 0.034 | 0.028 | 0.029 |
| 4 | 0.029 | 0.027 | 0.034 | 0.029 | 0.030 |
| 5 | 0.030 | 0.029 | 0.034 | 0.029 | 0.031 |
| 6 | 0.030 | 0.030 | 0.035 | 0.029 | 0.031 |
| 7 | 0.031 | 0.032 | 0.035 | 0.029 | 0.031 |
| 8 | 0.031 | 0.034 | 0.035 | 0.029 | 0.031 |
| 9 | 0.031 | 0.034 | 0.036 | 0.030 | 0.031 |
| 10 | 0.032 | 0.035 | 0.036 | 0.030 | 0.031 |
| 11 | 0.032 | 0.036 | 0.037 | 0.030 | 0.031 |
| 12 | 0.033 | 0.036 | 0.037 | 0.030 | 0.031 |
| 13 | 0.033 | 0.036 | 0.038 | 0.030 | 0.032 |
| 14 | 0.033 | 0.037 | 0.038 | 0.031 | 0.032 |
| 15 | 0.034 | 0.037 | 0.039 | 0.031 | 0.032 |
| 16 | 0.034 | 0.037 | 0.039 | 0.032 | 0.033 |
| 17 | 0.035 | 0.038 | 0.039 | 0.032 | 0.033 |
| 18 | 0.035 | 0.038 | 0.039 | 0.032 | 0.033 |
| 19 | 0.036 | 0.039 | 0.039 | 0.032 | 0.033 |
| 20 | 0.036 | 0.039 | 0.039 | 0.033 | 0.033 |
| 21 | 0.036 | 0.040 | 0.039 | 0.033 | 0.034 |
| 22 | 0.037 | 0.040 | 0.039 | 0.033 | 0.034 |

| | | | | | |
|---|---|---|---|---|---|
| 23 | 0.037 | 0.041 | 0.039 | 0.034 | 0.035 |
| 24 | 0.037 | 0.041 | 0.039 | 0.034 | 0.035 |
| 25 | 0.037 | 0.041 | 0.039 | 0.035 | 0.035 |
| 26 | 0.038 | 0.041 | 0.039 | 0.035 | 0.036 |
| 27 | 0.039 | 0.042 | 0.039 | 0.036 | 0.036 |
| 28 | 0.039 | 0.042 | 0.039 | 0.036 | 0.037 |
| 29 | 0.039 | 0.042 | 0.040 | 0.036 | 0.037 |
| 30 | 0.039 | 0.042 | 0.040 | 0.036 | 0.037 |
| 31 | 0.039 | 0.042 | 0.040 | 0.036 | 0.037 |
| 32 | 0.040 | 0.042 | 0.040 | 0.036 | 0.037 |
| 33 | 0.040 | 0.042 | 0.041 | 0.036 | 0.037 |
| 34 | 0.041 | 0.043 | 0.042 | 0.036 | 0.037 |
| 35 | 0.041 | 0.043 | 0.043 | 0.036 | 0.038 |
| 36 | 0.041 | 0.043 | 0.043 | 0.037 | 0.038 |
| 37 | 0.041 | 0.043 | 0.043 | 0.037 | 0.039 |
| 38 | 0.042 | 0.044 | 0.044 | 0.038 | 0.040 |
| 39 | 0.042 | 0.044 | 0.044 | 0.039 | 0.041 |
| 40 | 0.042 | 0.044 | 0.044 | 0.040 | 0.041 |
| 41 | 0.042 | 0.045 | 0.045 | 0.040 | 0.042 |
| 42 | 0.043 | 0.045 | 0.046 | 0.041 | 0.042 |
| 43 | 0.043 | 0.046 | 0.046 | 0.041 | 0.042 |
| 44 | 0.044 | 0.046 | 0.047 | 0.041 | 0.042 |
| 45 | 0.044 | 0.046 | 0.047 | 0.041 | 0.042 |
| 46 | 0.044 | 0.046 | 0.047 | 0.041 | 0.043 |
| 47 | 0.044 | 0.047 | 0.047 | 0.041 | 0.043 |
| 48 | 0.045 | 0.047 | 0.047 | 0.041 | 0.044 |
| 49 | 0.046 | 0.048 | 0.047 | 0.042 | 0.044 |
| 50 | 0.046 | 0.048 | 0.047 | 0.042 | 0.044 |
| 51 | 0.046 | 0.048 | 0.047 | 0.042 | 0.045 |
| 52 | 0.047 | 0.048 | 0.048 | 0.042 | 0.045 |
| 53 | 0.047 | 0.048 | 0.048 | 0.042 | 0.045 |
| 54 | 0.047 | 0.049 | 0.048 | 0.043 | 0.045 |
| 55 | 0.047 | 0.050 | 0.048 | 0.043 | 0.046 |
| 56 | 0.047 | 0.051 | 0.049 | 0.044 | 0.046 |
| 57 | 0.048 | 0.052 | 0.049 | 0.044 | 0.047 |
| 58 | 0.048 | 0.053 | 0.049 | 0.045 | 0.047 |
| 59 | 0.048 | 0.053 | 0.049 | 0.046 | 0.048 |
| 60 | 0.048 | 0.053 | 0.050 | 0.047 | 0.048 |
| 61 | 0.049 | 0.053 | 0.050 | 0.047 | 0.048 |
| 62 | 0.049 | 0.053 | 0.050 | 0.047 | 0.049 |
| 63 | 0.049 | 0.053 | 0.050 | 0.047 | 0.049 |
| 64 | 0.050 | 0.053 | 0.050 | 0.047 | 0.049 |
| 65 | 0.050 | 0.053 | 0.051 | 0.047 | 0.049 |
| 66 | 0.050 | 0.054 | 0.051 | 0.047 | 0.049 |

| | | | | | |
|---|---|---|---|---|---|
| 67 | 0.051 | 0.054 | 0.052 | 0.047 | 0.049 |
| 68 | 0.052 | 0.055 | 0.052 | 0.047 | 0.050 |
| 69 | 0.052 | 0.056 | 0.053 | 0.047 | 0.050 |
| 70 | 0.053 | 0.058 | 0.053 | 0.048 | 0.050 |
| 71 | 0.053 | 0.059 | 0.053 | 0.048 | 0.051 |
| 72 | 0.053 | 0.060 | 0.053 | 0.048 | 0.051 |
| 73 | 0.053 | 0.061 | 0.053 | 0.049 | 0.052 |
| 74 | 0.053 | 0.062 | 0.053 | 0.049 | 0.052 |
| 75 | 0.054 | 0.062 | 0.054 | 0.050 | 0.053 |
| 76 | 0.054 | 0.063 | 0.054 | 0.050 | 0.053 |
| 77 | 0.055 | 0.063 | 0.054 | 0.051 | 0.054 |
| 78 | 0.056 | 0.064 | 0.054 | 0.051 | 0.054 |
| 79 | 0.056 | 0.064 | 0.057 | 0.052 | 0.055 |
| 80 | 0.058 | 0.065 | 0.064 | 0.053 | 0.056 |
| 81 | 0.060 | 0.065 | 0.070 | 0.053 | 0.057 |
| 82 | 0.061 | 0.066 | 0.077 | 0.054 | 0.058 |
| 83 | 0.062 | 0.066 | 0.078 | 0.054 | 0.058 |
| 84 | 0.062 | 0.066 | 0.079 | 0.054 | 0.059 |
| 85 | 0.063 | 0.066 | 0.079 | 0.055 | 0.059 |
| 86 | 0.065 | 0.067 | 0.079 | 0.055 | 0.060 |
| 87 | 0.066 | 0.067 | 0.080 | 0.055 | 0.060 |
| 88 | 0.068 | 0.067 | 0.080 | 0.055 | 0.060 |
| 89 | 0.071 | 0.069 | 0.080 | 0.059 | 0.061 |
| 90 | 0.071 | 0.070 | 0.082 | 0.063 | 0.061 |
| 91 | 0.073 | 0.072 | 0.083 | 0.067 | 0.061 |
| 92 | 0.075 | 0.074 | 0.085 | 0.071 | 0.061 |
| 93 | 0.077 | 0.075 | 0.087 | 0.071 | 0.061 |
| 94 | 0.078 | 0.076 | 0.088 | 0.071 | 0.062 |
| 95 | 0.079 | 0.077 | 0.090 | 0.071 | 0.062 |
| 96 | 0.080 | 0.078 | 0.091 | 0.072 | 0.062 |
| 97 | 0.085 | 0.080 | 0.092 | 0.072 | 0.064 |
| 98 | 0.087 | 0.083 | 0.092 | 0.073 | 0.066 |
| 99 | 0.091 | 0.085 | 0.092 | 0.074 | 0.069 |
| 100 | 0.092 | 0.087 | 0.092 | 0.075 | 0.071 |

***Table S58:*** *Synchronization-continuation – Variability (CV of ITI), medium tempo – SyncCont_metro_600_mean_CV_iti*

| | | | | | | |
|---|---|---|---|---|---|---|
| **Task** | | Synchronization-continuation | | | | |
| **Outcome measure** | | Variability (CV of ITI), medium tempo | | | | |
| **Variable name** | | SyncCont_metro_600_mean_CV_iti | | | | |
| **Age and gender effects** | | | | | | |
| Age regression (slope) | | 0.000 | | | | |
| Age regression (p) | | 0.086 | | | | |
| Age group (p) | | 0.287 | | | | |
| Gender (p) | | 0.865 | | | | |
| **Group** | | All | Age 18 to 21 | Age 22 to 29 | Age 30 to 54 | Age 55 to 87 |
| N (N female) | | 108 (74) | 27 (18) | 29 (19) | 26 (16) | 26 (21) |
| **Normality** | | | | | | |
| Skewness | | 1.05 | 1.49 | 0.94 | 0.50 | 1.41 |
| Excess kurtosis | | 1.16 | 2.88 | 0.83 | -0.97 | 2.31 |
| **Scores** | | | | | | |
| Mean | | 0.046 | 0.048 | 0.046 | 0.046 | 0.042 |
| SD | | 0.014 | 0.014 | 0.015 | 0.014 | 0.012 |
| Percentiles | | | | | | |
| | 0 | 0.025 | 0.030 | 0.025 | 0.028 | 0.027 |
| | 1 | 0.027 | 0.031 | 0.026 | 0.028 | 0.028 |
| | 2 | 0.027 | 0.032 | 0.026 | 0.029 | 0.028 |
| | 3 | 0.028 | 0.033 | 0.027 | 0.029 | 0.029 |
| | 4 | 0.029 | 0.033 | 0.028 | 0.030 | 0.029 |
| | 5 | 0.029 | 0.034 | 0.028 | 0.030 | 0.029 |
| | 6 | 0.030 | 0.034 | 0.028 | 0.030 | 0.029 |
| | 7 | 0.030 | 0.035 | 0.029 | 0.030 | 0.030 |
| | 8 | 0.030 | 0.035 | 0.030 | 0.030 | 0.030 |
| | 9 | 0.030 | 0.036 | 0.031 | 0.031 | 0.030 |
| | 10 | 0.031 | 0.036 | 0.032 | 0.031 | 0.030 |
| | 11 | 0.032 | 0.036 | 0.032 | 0.032 | 0.030 |
| | 12 | 0.032 | 0.036 | 0.032 | 0.032 | 0.030 |
| | 13 | 0.032 | 0.036 | 0.032 | 0.032 | 0.031 |
| | 14 | 0.032 | 0.036 | 0.032 | 0.033 | 0.031 |
| | 15 | 0.032 | 0.036 | 0.032 | 0.033 | 0.031 |
| | 16 | 0.032 | 0.037 | 0.032 | 0.033 | 0.032 |
| | 17 | 0.033 | 0.037 | 0.032 | 0.033 | 0.032 |
| | 18 | 0.033 | 0.037 | 0.033 | 0.033 | 0.032 |
| | 19 | 0.034 | 0.037 | 0.033 | 0.033 | 0.032 |
| | 20 | 0.034 | 0.037 | 0.034 | 0.034 | 0.032 |
| | 21 | 0.034 | 0.038 | 0.034 | 0.034 | 0.032 |
| | 22 | 0.034 | 0.038 | 0.034 | 0.034 | 0.032 |

| | | | | | |
|---|---|---|---|---|---|
| 23 | 0.035 | 0.038 | 0.035 | 0.034 | 0.032 |
| 24 | 0.035 | 0.038 | 0.035 | 0.034 | 0.032 |
| 25 | 0.035 | 0.039 | 0.035 | 0.035 | 0.033 |
| 26 | 0.036 | 0.039 | 0.035 | 0.035 | 0.033 |
| 27 | 0.036 | 0.039 | 0.036 | 0.035 | 0.033 |
| 28 | 0.036 | 0.039 | 0.036 | 0.035 | 0.034 |
| 29 | 0.036 | 0.040 | 0.036 | 0.035 | 0.034 |
| 30 | 0.036 | 0.040 | 0.036 | 0.036 | 0.034 |
| 31 | 0.037 | 0.040 | 0.037 | 0.036 | 0.034 |
| 32 | 0.037 | 0.040 | 0.037 | 0.037 | 0.034 |
| 33 | 0.037 | 0.040 | 0.037 | 0.037 | 0.034 |
| 34 | 0.038 | 0.040 | 0.038 | 0.037 | 0.035 |
| 35 | 0.038 | 0.041 | 0.039 | 0.038 | 0.035 |
| 36 | 0.038 | 0.041 | 0.040 | 0.038 | 0.036 |
| 37 | 0.039 | 0.041 | 0.041 | 0.038 | 0.036 |
| 38 | 0.039 | 0.042 | 0.042 | 0.039 | 0.036 |
| 39 | 0.039 | 0.042 | 0.043 | 0.039 | 0.036 |
| 40 | 0.039 | 0.043 | 0.043 | 0.039 | 0.036 |
| 41 | 0.040 | 0.043 | 0.043 | 0.040 | 0.036 |
| 42 | 0.040 | 0.043 | 0.043 | 0.040 | 0.037 |
| 43 | 0.040 | 0.043 | 0.043 | 0.041 | 0.037 |
| 44 | 0.041 | 0.044 | 0.043 | 0.041 | 0.037 |
| 45 | 0.041 | 0.044 | 0.044 | 0.041 | 0.038 |
| 46 | 0.042 | 0.044 | 0.044 | 0.041 | 0.038 |
| 47 | 0.042 | 0.045 | 0.044 | 0.041 | 0.038 |
| 48 | 0.043 | 0.045 | 0.044 | 0.041 | 0.038 |
| 49 | 0.043 | 0.046 | 0.044 | 0.041 | 0.039 |
| 50 | 0.044 | 0.046 | 0.044 | 0.041 | 0.039 |
| 51 | 0.044 | 0.047 | 0.045 | 0.042 | 0.039 |
| 52 | 0.044 | 0.047 | 0.045 | 0.042 | 0.039 |
| 53 | 0.044 | 0.047 | 0.045 | 0.043 | 0.039 |
| 54 | 0.044 | 0.047 | 0.045 | 0.044 | 0.040 |
| 55 | 0.044 | 0.047 | 0.045 | 0.044 | 0.040 |
| 56 | 0.045 | 0.048 | 0.045 | 0.045 | 0.040 |
| 57 | 0.045 | 0.048 | 0.046 | 0.046 | 0.041 |
| 58 | 0.046 | 0.048 | 0.046 | 0.047 | 0.042 |
| 59 | 0.046 | 0.048 | 0.046 | 0.048 | 0.043 |
| 60 | 0.046 | 0.049 | 0.046 | 0.049 | 0.044 |
| 61 | 0.047 | 0.049 | 0.046 | 0.049 | 0.044 |
| 62 | 0.047 | 0.050 | 0.046 | 0.049 | 0.044 |
| 63 | 0.047 | 0.050 | 0.047 | 0.050 | 0.044 |
| 64 | 0.048 | 0.050 | 0.047 | 0.050 | 0.044 |
| 65 | 0.049 | 0.050 | 0.048 | 0.051 | 0.044 |
| 66 | 0.049 | 0.050 | 0.050 | 0.052 | 0.044 |

| | | | | | |
|---|---|---|---|---|---|
| 67 | 0.049 | 0.050 | 0.052 | 0.053 | 0.044 |
| 68 | 0.050 | 0.051 | 0.053 | 0.055 | 0.044 |
| 69 | 0.050 | 0.051 | 0.053 | 0.055 | 0.044 |
| 70 | 0.050 | 0.052 | 0.054 | 0.056 | 0.045 |
| 71 | 0.051 | 0.053 | 0.054 | 0.057 | 0.045 |
| 72 | 0.053 | 0.053 | 0.054 | 0.058 | 0.046 |
| 73 | 0.054 | 0.054 | 0.055 | 0.058 | 0.046 |
| 74 | 0.054 | 0.054 | 0.055 | 0.058 | 0.047 |
| 75 | 0.054 | 0.054 | 0.056 | 0.058 | 0.048 |
| 76 | 0.055 | 0.054 | 0.056 | 0.059 | 0.048 |
| 77 | 0.056 | 0.054 | 0.057 | 0.059 | 0.049 |
| 78 | 0.056 | 0.055 | 0.057 | 0.059 | 0.049 |
| 79 | 0.057 | 0.055 | 0.057 | 0.059 | 0.049 |
| 80 | 0.057 | 0.056 | 0.058 | 0.059 | 0.049 |
| 81 | 0.058 | 0.057 | 0.058 | 0.060 | 0.049 |
| 82 | 0.058 | 0.058 | 0.059 | 0.060 | 0.050 |
| 83 | 0.059 | 0.060 | 0.059 | 0.060 | 0.050 |
| 84 | 0.059 | 0.061 | 0.059 | 0.060 | 0.050 |
| 85 | 0.059 | 0.062 | 0.059 | 0.060 | 0.052 |
| 86 | 0.059 | 0.062 | 0.059 | 0.061 | 0.053 |
| 87 | 0.059 | 0.062 | 0.059 | 0.061 | 0.054 |
| 88 | 0.060 | 0.062 | 0.059 | 0.061 | 0.056 |
| 89 | 0.062 | 0.062 | 0.059 | 0.063 | 0.056 |
| 90 | 0.062 | 0.063 | 0.061 | 0.065 | 0.057 |
| 91 | 0.063 | 0.063 | 0.064 | 0.067 | 0.058 |
| 92 | 0.066 | 0.064 | 0.066 | 0.069 | 0.058 |
| 93 | 0.068 | 0.065 | 0.069 | 0.069 | 0.058 |
| 94 | 0.069 | 0.066 | 0.070 | 0.069 | 0.058 |
| 95 | 0.069 | 0.067 | 0.072 | 0.069 | 0.058 |
| 96 | 0.072 | 0.068 | 0.073 | 0.070 | 0.059 |
| 97 | 0.074 | 0.074 | 0.076 | 0.071 | 0.064 |
| 98 | 0.080 | 0.081 | 0.081 | 0.072 | 0.070 |
| 99 | 0.089 | 0.087 | 0.085 | 0.073 | 0.075 |
| 100 | 0.094 | 0.094 | 0.090 | 0.075 | 0.081 |

***Table S59:*** *Synchronization-continuation – Variability (CV of ITI), slow tempo – SyncCont_metro_750_mean_CV_iti*

| | | | | | | |
|---|---|---|---|---|---|---|
| **Task** | | Synchronization-continuation | | | | |
| **Outcome measure** | | Variability (CV of ITI), slow tempo | | | | |
| **Variable name** | | SyncCont_metro_750_mean_CV_iti | | | | |
| **Age and gender effects** | | | | | | |
| Age regression (slope) | | 0.000 | | | | |
| Age regression (p) | | 0.019 | | | | |
| Age group (p) | | 0.086 | | | | |
| Gender (p) | | 0.721 | | | | |
| **Group** | | All | Age 18 to 21 | Age 22 to 29 | Age 30 to 54 | Age 55 to 87 |
| N (N female) | | 108 (74) | 27 (18) | 29 (19) | 26 (16) | 26 (21) |
| **Normality** | | | | | | |
| Skewness | | 1.22 | 1.60 | 0.60 | 0.48 | 0.43 |
| Excess kurtosis | | 2.59 | 2.78 | -0.03 | -0.03 | -1.18 |
| **Scores** | | | | | | |
| Mean | | 0.048 | 0.053 | 0.048 | 0.049 | 0.042 |
| SD | | 0.014 | 0.017 | 0.012 | 0.015 | 0.009 |
| Percentiles | | | | | | |
| | 0 | 0.024 | 0.028 | 0.026 | 0.024 | 0.031 |
| | 1 | 0.026 | 0.030 | 0.027 | 0.025 | 0.031 |
| | 2 | 0.028 | 0.032 | 0.028 | 0.026 | 0.031 |
| | 3 | 0.029 | 0.034 | 0.030 | 0.027 | 0.031 |
| | 4 | 0.030 | 0.036 | 0.031 | 0.028 | 0.031 |
| | 5 | 0.030 | 0.036 | 0.031 | 0.029 | 0.031 |
| | 6 | 0.031 | 0.037 | 0.032 | 0.029 | 0.031 |
| | 7 | 0.031 | 0.037 | 0.033 | 0.029 | 0.031 |
| | 8 | 0.032 | 0.037 | 0.033 | 0.029 | 0.031 |
| | 9 | 0.033 | 0.038 | 0.034 | 0.031 | 0.032 |
| | 10 | 0.033 | 0.038 | 0.035 | 0.032 | 0.032 |
| | 11 | 0.034 | 0.038 | 0.036 | 0.033 | 0.032 |
| | 12 | 0.034 | 0.039 | 0.036 | 0.034 | 0.032 |
| | 13 | 0.035 | 0.040 | 0.037 | 0.034 | 0.033 |
| | 14 | 0.035 | 0.041 | 0.037 | 0.034 | 0.033 |
| | 15 | 0.035 | 0.042 | 0.037 | 0.034 | 0.033 |
| | 16 | 0.036 | 0.042 | 0.038 | 0.034 | 0.034 |
| | 17 | 0.036 | 0.042 | 0.038 | 0.035 | 0.034 |
| | 18 | 0.037 | 0.042 | 0.038 | 0.035 | 0.034 |
| | 19 | 0.037 | 0.043 | 0.038 | 0.036 | 0.034 |
| | 20 | 0.038 | 0.043 | 0.038 | 0.036 | 0.035 |
| | 21 | 0.038 | 0.043 | 0.038 | 0.037 | 0.035 |
| | 22 | 0.038 | 0.043 | 0.038 | 0.038 | 0.035 |

| | | | | | |
|---|---|---|---|---|---|
| 23 | 0.038 | 0.043 | 0.039 | 0.039 | 0.035 |
| 24 | 0.039 | 0.043 | 0.039 | 0.039 | 0.035 |
| 25 | 0.039 | 0.044 | 0.040 | 0.039 | 0.035 |
| 26 | 0.039 | 0.044 | 0.040 | 0.040 | 0.035 |
| 27 | 0.039 | 0.044 | 0.041 | 0.040 | 0.036 |
| 28 | 0.039 | 0.044 | 0.041 | 0.040 | 0.036 |
| 29 | 0.039 | 0.044 | 0.042 | 0.040 | 0.037 |
| 30 | 0.039 | 0.045 | 0.042 | 0.041 | 0.037 |
| 31 | 0.040 | 0.045 | 0.042 | 0.041 | 0.038 |
| 32 | 0.040 | 0.045 | 0.042 | 0.042 | 0.038 |
| 33 | 0.042 | 0.045 | 0.042 | 0.042 | 0.038 |
| 34 | 0.042 | 0.045 | 0.043 | 0.042 | 0.039 |
| 35 | 0.042 | 0.045 | 0.044 | 0.043 | 0.039 |
| 36 | 0.042 | 0.045 | 0.044 | 0.043 | 0.039 |
| 37 | 0.043 | 0.046 | 0.044 | 0.043 | 0.039 |
| 38 | 0.043 | 0.046 | 0.045 | 0.043 | 0.039 |
| 39 | 0.043 | 0.046 | 0.045 | 0.043 | 0.039 |
| 40 | 0.043 | 0.046 | 0.045 | 0.043 | 0.039 |
| 41 | 0.043 | 0.046 | 0.046 | 0.044 | 0.039 |
| 42 | 0.044 | 0.046 | 0.046 | 0.045 | 0.039 |
| 43 | 0.044 | 0.046 | 0.046 | 0.046 | 0.039 |
| 44 | 0.045 | 0.047 | 0.046 | 0.046 | 0.039 |
| 45 | 0.045 | 0.047 | 0.046 | 0.047 | 0.039 |
| 46 | 0.045 | 0.047 | 0.046 | 0.047 | 0.039 |
| 47 | 0.046 | 0.047 | 0.046 | 0.047 | 0.039 |
| 48 | 0.046 | 0.048 | 0.047 | 0.047 | 0.039 |
| 49 | 0.046 | 0.049 | 0.047 | 0.048 | 0.039 |
| 50 | 0.046 | 0.049 | 0.047 | 0.049 | 0.039 |
| 51 | 0.047 | 0.049 | 0.047 | 0.049 | 0.039 |
| 52 | 0.047 | 0.049 | 0.047 | 0.050 | 0.039 |
| 53 | 0.047 | 0.050 | 0.047 | 0.050 | 0.039 |
| 54 | 0.047 | 0.050 | 0.047 | 0.050 | 0.039 |
| 55 | 0.047 | 0.050 | 0.047 | 0.050 | 0.039 |
| 56 | 0.049 | 0.050 | 0.047 | 0.051 | 0.039 |
| 57 | 0.049 | 0.050 | 0.047 | 0.051 | 0.040 |
| 58 | 0.049 | 0.051 | 0.048 | 0.051 | 0.041 |
| 59 | 0.050 | 0.051 | 0.049 | 0.051 | 0.042 |
| 60 | 0.050 | 0.052 | 0.049 | 0.051 | 0.043 |
| 61 | 0.050 | 0.053 | 0.050 | 0.052 | 0.043 |
| 62 | 0.050 | 0.054 | 0.050 | 0.052 | 0.043 |
| 63 | 0.050 | 0.054 | 0.050 | 0.053 | 0.043 |
| 64 | 0.051 | 0.055 | 0.050 | 0.053 | 0.043 |
| 65 | 0.051 | 0.055 | 0.051 | 0.053 | 0.044 |
| 66 | 0.051 | 0.055 | 0.051 | 0.053 | 0.046 |

| | | | | | |
|---|---|---|---|---|---|
| 67 | 0.052 | 0.055 | 0.051 | 0.053 | 0.048 |
| 68 | 0.053 | 0.055 | 0.051 | 0.053 | 0.049 |
| 69 | 0.053 | 0.056 | 0.052 | 0.053 | 0.049 |
| 70 | 0.053 | 0.056 | 0.052 | 0.053 | 0.049 |
| 71 | 0.053 | 0.056 | 0.053 | 0.053 | 0.049 |
| 72 | 0.054 | 0.056 | 0.053 | 0.054 | 0.049 |
| 73 | 0.054 | 0.056 | 0.053 | 0.054 | 0.049 |
| 74 | 0.054 | 0.056 | 0.054 | 0.054 | 0.049 |
| 75 | 0.054 | 0.057 | 0.054 | 0.054 | 0.050 |
| 76 | 0.055 | 0.057 | 0.054 | 0.054 | 0.050 |
| 77 | 0.055 | 0.057 | 0.054 | 0.055 | 0.050 |
| 78 | 0.055 | 0.058 | 0.054 | 0.057 | 0.051 |
| 79 | 0.055 | 0.058 | 0.054 | 0.058 | 0.052 |
| 80 | 0.055 | 0.059 | 0.055 | 0.060 | 0.053 |
| 81 | 0.056 | 0.060 | 0.055 | 0.061 | 0.053 |
| 82 | 0.057 | 0.060 | 0.055 | 0.061 | 0.053 |
| 83 | 0.058 | 0.060 | 0.056 | 0.062 | 0.053 |
| 84 | 0.058 | 0.060 | 0.057 | 0.063 | 0.053 |
| 85 | 0.059 | 0.060 | 0.058 | 0.064 | 0.054 |
| 86 | 0.060 | 0.061 | 0.060 | 0.066 | 0.054 |
| 87 | 0.060 | 0.062 | 0.062 | 0.067 | 0.054 |
| 88 | 0.061 | 0.063 | 0.065 | 0.069 | 0.055 |
| 89 | 0.063 | 0.065 | 0.067 | 0.069 | 0.055 |
| 90 | 0.065 | 0.070 | 0.068 | 0.069 | 0.055 |
| 91 | 0.068 | 0.074 | 0.069 | 0.069 | 0.055 |
| 92 | 0.069 | 0.078 | 0.069 | 0.069 | 0.055 |
| 93 | 0.069 | 0.081 | 0.070 | 0.069 | 0.056 |
| 94 | 0.069 | 0.083 | 0.071 | 0.069 | 0.057 |
| 95 | 0.072 | 0.086 | 0.072 | 0.069 | 0.057 |
| 96 | 0.076 | 0.088 | 0.073 | 0.069 | 0.058 |
| 97 | 0.079 | 0.092 | 0.074 | 0.073 | 0.058 |
| 98 | 0.084 | 0.097 | 0.075 | 0.077 | 0.058 |
| 99 | 0.088 | 0.101 | 0.076 | 0.081 | 0.058 |
| 100 | 0.106 | 0.106 | 0.076 | 0.085 | 0.058 |

***Table S60:*** *Adaptive tapping – Sensitivity index (d') of perceiving tempo deceleration (IOI + 75ms) – Adaptive_plus_75_dprime2*

| | | | | | |
|---|---|---|---|---|---|
| **Task** | Adaptive tapping | | | | |
| **Outcome measure** | Sensitivity index (d') of perceiving tempo deceleration (IOI + 75ms) | | | | |
| **Variable name** | Adaptive_plus_75_dprime2 | | | | |
| **Age and gender effects** | | | | | |
| Age regression (slope) | 0.000 | | | | |
| Age regression (p) | 1.000 | | | | |
| Age group (p) | 1.000 | | | | |
| Gender (p) | 1.000 | | | | |
| **Group** | All | Age 18 to 21 | Age 22 to 29 | Age 30 to 54 | Age 55 to 87 |
| N (N female) | 105 (72) | 26 (17) | 29 (19) | 26 (16) | 24 (20) |
| **Normality** | | | | | |
| Skewness | -1.74 | -1.06 | -1.87 | -2.47 | -1.05 |
| Excess kurtosis | 2.72 | 0.41 | 2.25 | 6.50 | 0.06 |
| **Scores** | | | | | |
| Mean | 3.63 | 3.47 | 3.68 | 3.70 | 3.66 |
| SD | 0.74 | 0.74 | 0.86 | 0.75 | 0.58 |
| Percentiles | | | | | |
| 0 | 0.82 | 1.45 | 1.25 | 0.82 | 2.22 |
| 1 | 1.25 | 1.62 | 1.27 | 1.24 | 2.30 |
| 2 | 1.31 | 1.79 | 1.28 | 1.67 | 2.38 |
| 3 | 1.53 | 1.96 | 1.30 | 2.10 | 2.46 |
| 4 | 2.15 | 2.13 | 1.41 | 2.53 | 2.54 |
| 5 | 2.22 | 2.23 | 1.67 | 2.62 | 2.59 |
| 6 | 2.24 | 2.32 | 1.92 | 2.70 | 2.62 |
| 7 | 2.37 | 2.42 | 2.18 | 2.79 | 2.65 |
| 8 | 2.52 | 2.52 | 2.24 | 2.87 | 2.67 |
| 9 | 2.53 | 2.52 | 2.27 | 2.87 | 2.71 |
| 10 | 2.55 | 2.52 | 2.29 | 2.87 | 2.76 |
| 11 | 2.63 | 2.53 | 2.36 | 2.87 | 2.81 |
| 12 | 2.69 | 2.53 | 2.53 | 2.87 | 2.86 |
| 13 | 2.79 | 2.57 | 2.70 | 2.97 | 2.91 |
| 14 | 2.87 | 2.61 | 2.86 | 3.07 | 2.99 |
| 15 | 2.90 | 2.65 | 2.96 | 3.17 | 3.07 |
| 16 | 2.91 | 2.69 | 3.03 | 3.27 | 3.15 |
| 17 | 2.91 | 2.75 | 3.10 | 3.30 | 3.24 |
| 18 | 2.91 | 2.80 | 3.17 | 3.34 | 3.27 |
| 19 | 3.10 | 2.86 | 3.29 | 3.37 | 3.27 |
| 20 | 3.25 | 2.91 | 3.40 | 3.40 | 3.27 |
| 21 | 3.27 | 2.91 | 3.52 | 3.44 | 3.27 |
| 22 | 3.27 | 2.91 | 3.60 | 3.48 | 3.28 |

| | | | | | |
|---|---|---|---|---|---|
| 23 | 3.27 | 2.91 | 3.66 | 3.53 | 3.31 |
| 24 | 3.27 | 2.91 | 3.72 | 3.57 | 3.34 |
| 25 | 3.40 | 3.00 | 3.78 | 3.62 | 3.37 |
| 26 | 3.40 | 3.09 | 3.78 | 3.67 | 3.40 |
| 27 | 3.40 | 3.18 | 3.78 | 3.73 | 3.44 |
| 28 | 3.42 | 3.27 | 3.78 | 3.78 | 3.47 |
| 29 | 3.57 | 3.27 | 3.78 | 3.78 | 3.51 |
| 30 | 3.57 | 3.27 | 3.78 | 3.78 | 3.55 |
| 31 | 3.57 | 3.27 | 3.78 | 3.78 | 3.57 |
| 32 | 3.57 | 3.27 | 3.78 | 3.78 | 3.57 |
| 33 | 3.57 | 3.30 | 3.87 | 3.78 | 3.57 |
| 34 | 3.64 | 3.34 | 3.97 | 3.78 | 3.57 |
| 35 | 3.78 | 3.37 | 4.07 | 3.78 | 3.57 |
| 36 | 3.78 | 3.40 | 4.14 | 3.78 | 3.57 |
| 37 | 3.78 | 3.40 | 4.14 | 3.78 | 3.57 |
| 38 | 3.78 | 3.40 | 4.14 | 3.78 | 3.57 |
| 39 | 3.78 | 3.40 | 4.14 | 3.78 | 3.57 |
| 40 | 3.78 | 3.40 | 4.14 | 3.78 | 3.61 |
| 41 | 3.78 | 3.44 | 4.14 | 3.78 | 3.66 |
| 42 | 3.78 | 3.48 | 4.14 | 3.78 | 3.71 |
| 43 | 3.78 | 3.53 | 4.14 | 3.78 | 3.76 |
| 44 | 3.78 | 3.57 | 4.14 | 3.78 | 3.78 |
| 45 | 3.78 | 3.62 | 4.14 | 3.87 | 3.78 |
| 46 | 3.78 | 3.67 | 4.14 | 3.96 | 3.78 |
| 47 | 3.78 | 3.73 | 4.14 | 4.05 | 3.78 |
| 48 | 3.78 | 3.78 | 4.14 | 4.14 | 3.78 |
| 49 | 3.78 | 3.78 | 4.14 | 4.14 | 3.78 |
| 50 | 4.14 | 3.78 | 4.14 | 4.14 | 3.78 |
| 51 | 4.14 | 3.78 | 4.14 | 4.14 | 3.78 |
| 52 | 4.14 | 3.78 | 4.14 | 4.14 | 3.78 |
| 53 | 4.14 | 3.78 | 4.14 | 4.14 | 3.85 |
| 54 | 4.14 | 3.78 | 4.14 | 4.14 | 3.93 |
| 55 | 4.14 | 3.78 | 4.14 | 4.14 | 4.02 |
| 56 | 4.14 | 3.78 | 4.14 | 4.14 | 4.10 |
| 57 | 4.14 | 3.78 | 4.14 | 4.14 | 4.14 |
| 58 | 4.14 | 3.78 | 4.14 | 4.14 | 4.14 |
| 59 | 4.14 | 3.78 | 4.14 | 4.14 | 4.14 |
| 60 | 4.14 | 3.78 | 4.14 | 4.14 | 4.14 |
| 61 | 4.14 | 3.78 | 4.14 | 4.14 | 4.14 |
| 62 | 4.14 | 3.78 | 4.14 | 4.14 | 4.14 |
| 63 | 4.14 | 3.78 | 4.14 | 4.14 | 4.14 |
| 64 | 4.14 | 3.78 | 4.14 | 4.14 | 4.14 |
| 65 | 4.14 | 3.87 | 4.14 | 4.14 | 4.14 |
| 66 | 4.14 | 3.96 | 4.14 | 4.14 | 4.14 |

| | | | | | |
|---|---|---|---|---|---|
| 67 | 4.14 | 4.05 | 4.14 | 4.14 | 4.14 |
| 68 | 4.14 | 4.14 | 4.14 | 4.14 | 4.14 |
| 69 | 4.14 | 4.14 | 4.14 | 4.14 | 4.14 |
| 70 | 4.14 | 4.14 | 4.14 | 4.14 | 4.14 |
| 71 | 4.14 | 4.14 | 4.14 | 4.14 | 4.14 |
| 72 | 4.14 | 4.14 | 4.14 | 4.14 | 4.14 |
| 73 | 4.14 | 4.14 | 4.14 | 4.14 | 4.14 |
| 74 | 4.14 | 4.14 | 4.14 | 4.14 | 4.14 |
| 75 | 4.14 | 4.14 | 4.14 | 4.14 | 4.14 |
| 76 | 4.14 | 4.14 | 4.14 | 4.14 | 4.14 |
| 77 | 4.14 | 4.14 | 4.14 | 4.14 | 4.14 |
| 78 | 4.14 | 4.14 | 4.14 | 4.14 | 4.14 |
| 79 | 4.14 | 4.14 | 4.14 | 4.14 | 4.14 |
| 80 | 4.14 | 4.14 | 4.14 | 4.14 | 4.14 |
| 81 | 4.14 | 4.14 | 4.14 | 4.14 | 4.14 |
| 82 | 4.14 | 4.14 | 4.14 | 4.14 | 4.14 |
| 83 | 4.14 | 4.14 | 4.14 | 4.14 | 4.14 |
| 84 | 4.14 | 4.14 | 4.14 | 4.14 | 4.14 |
| 85 | 4.14 | 4.14 | 4.14 | 4.14 | 4.14 |
| 86 | 4.14 | 4.14 | 4.14 | 4.14 | 4.14 |
| 87 | 4.14 | 4.14 | 4.14 | 4.14 | 4.14 |
| 88 | 4.14 | 4.14 | 4.14 | 4.14 | 4.14 |
| 89 | 4.14 | 4.14 | 4.14 | 4.14 | 4.14 |
| 90 | 4.14 | 4.14 | 4.14 | 4.14 | 4.14 |
| 91 | 4.14 | 4.14 | 4.14 | 4.14 | 4.14 |
| 92 | 4.14 | 4.14 | 4.14 | 4.14 | 4.14 |
| 93 | 4.14 | 4.14 | 4.14 | 4.14 | 4.14 |
| 94 | 4.14 | 4.14 | 4.14 | 4.14 | 4.14 |
| 95 | 4.14 | 4.14 | 4.14 | 4.14 | 4.14 |
| 96 | 4.14 | 4.14 | 4.14 | 4.14 | 4.14 |
| 97 | 4.14 | 4.14 | 4.14 | 4.14 | 4.14 |
| 98 | 4.14 | 4.14 | 4.14 | 4.14 | 4.14 |
| 99 | 4.14 | 4.14 | 4.14 | 4.14 | 4.14 |
| 100 | 4.14 | 4.14 | 4.14 | 4.14 | 4.14 |

***Table S61:*** *Adaptive tapping – Sensitivity index (d') of perceiving tempo deceleration (IOI + 30ms) – Adaptive_plus_30_dprime2*

| | | | | | |
|---|---|---|---|---|---|
| **Task** | Adaptive tapping | | | | |
| **Outcome measure** | Sensitivity index (d') of perceiving tempo deceleration (IOI + 30ms) | | | | |
| **Variable name** | Adaptive_plus_30_dprime2 | | | | |
| **Age and gender effects** | | | | | |
| Age regression (slope) | 0.003 | | | | |
| Age regression (p) | 0.520 | | | | |
| Age group (p) | 0.339 | | | | |
| Gender (p) | 0.589 | | | | |
| **Group** | All | Age 18 to 21 | Age 22 to 29 | Age 30 to 54 | Age 55 to 87 |
| N (N female) | 105 (72) | 26 (17) | 29 (19) | 26 (16) | 24 (20) |
| **Normality** | | | | | |
| Skewness | 0.23 | 0.40 | 0.69 | -0.79 | 0.54 |
| Excess kurtosis | -0.02 | -0.64 | 0.94 | 0.62 | 0.11 |
| **Scores** | | | | | |
| Mean | 1.84 | 1.69 | 1.75 | 2.01 | 1.93 |
| SD | 0.98 | 1.19 | 0.83 | 0.88 | 1.01 |
| Percentiles | | | | | |
| 0 | -0.36 | -0.36 | 0.30 | -0.23 | 0.00 |
| 1 | -0.22 | -0.27 | 0.30 | -0.17 | 0.16 |
| 2 | 0.00 | -0.18 | 0.30 | -0.11 | 0.32 |
| 3 | 0.00 | -0.09 | 0.30 | -0.06 | 0.48 |
| 4 | 0.05 | 0.00 | 0.37 | 0.00 | 0.64 |
| 5 | 0.30 | 0.10 | 0.53 | 0.27 | 0.72 |
| 6 | 0.32 | 0.20 | 0.69 | 0.54 | 0.76 |
| 7 | 0.43 | 0.30 | 0.85 | 0.82 | 0.80 |
| 8 | 0.52 | 0.40 | 0.88 | 1.09 | 0.84 |
| 9 | 0.52 | 0.43 | 0.89 | 1.12 | 0.87 |
| 10 | 0.60 | 0.46 | 0.90 | 1.15 | 0.87 |
| 11 | 0.75 | 0.49 | 0.91 | 1.18 | 0.87 |
| 12 | 0.85 | 0.52 | 0.91 | 1.21 | 0.87 |
| 13 | 0.87 | 0.52 | 0.91 | 1.23 | 0.87 |
| 14 | 0.87 | 0.52 | 0.91 | 1.24 | 0.90 |
| 15 | 0.87 | 0.52 | 0.98 | 1.25 | 0.93 |
| 16 | 0.87 | 0.52 | 1.08 | 1.27 | 0.95 |
| 17 | 0.90 | 0.52 | 1.18 | 1.32 | 0.98 |
| 18 | 0.91 | 0.52 | 1.27 | 1.37 | 1.02 |
| 19 | 0.97 | 0.53 | 1.27 | 1.42 | 1.07 |
| 20 | 1.07 | 0.53 | 1.27 | 1.47 | 1.12 |
| 21 | 1.19 | 0.61 | 1.27 | 1.49 | 1.17 |
| 22 | 1.21 | 0.68 | 1.27 | 1.52 | 1.21 |

| | | | | | |
|---|---|---|---|---|---|
| 23 | 1.21 | 0.76 | 1.27 | 1.54 | 1.23 |
| 24 | 1.27 | 0.83 | 1.27 | 1.57 | 1.24 |
| 25 | 1.27 | 0.84 | 1.27 | 1.63 | 1.25 |
| 26 | 1.27 | 0.85 | 1.27 | 1.70 | 1.27 |
| 27 | 1.27 | 0.86 | 1.27 | 1.76 | 1.33 |
| 28 | 1.27 | 0.87 | 1.27 | 1.83 | 1.40 |
| 29 | 1.27 | 0.87 | 1.27 | 1.83 | 1.47 |
| 30 | 1.28 | 0.87 | 1.27 | 1.83 | 1.54 |
| 31 | 1.36 | 0.87 | 1.27 | 1.83 | 1.57 |
| 32 | 1.47 | 0.87 | 1.27 | 1.83 | 1.57 |
| 33 | 1.48 | 0.96 | 1.28 | 1.83 | 1.57 |
| 34 | 1.52 | 1.04 | 1.30 | 1.83 | 1.57 |
| 35 | 1.57 | 1.13 | 1.31 | 1.83 | 1.57 |
| 36 | 1.57 | 1.21 | 1.34 | 1.83 | 1.57 |
| 37 | 1.57 | 1.28 | 1.39 | 1.86 | 1.57 |
| 38 | 1.57 | 1.34 | 1.44 | 1.89 | 1.57 |
| 39 | 1.57 | 1.41 | 1.48 | 1.92 | 1.57 |
| 40 | 1.57 | 1.47 | 1.51 | 1.96 | 1.60 |
| 41 | 1.57 | 1.49 | 1.53 | 1.96 | 1.63 |
| 42 | 1.57 | 1.52 | 1.55 | 1.96 | 1.66 |
| 43 | 1.67 | 1.54 | 1.57 | 1.96 | 1.70 |
| 44 | 1.71 | 1.57 | 1.57 | 1.96 | 1.72 |
| 45 | 1.71 | 1.57 | 1.57 | 1.99 | 1.72 |
| 46 | 1.74 | 1.57 | 1.57 | 2.01 | 1.73 |
| 47 | 1.82 | 1.57 | 1.59 | 2.04 | 1.73 |
| 48 | 1.83 | 1.57 | 1.63 | 2.07 | 1.74 |
| 49 | 1.83 | 1.57 | 1.67 | 2.07 | 1.76 |
| 50 | 1.83 | 1.57 | 1.71 | 2.07 | 1.78 |
| 51 | 1.83 | 1.57 | 1.75 | 2.07 | 1.80 |
| 52 | 1.83 | 1.58 | 1.78 | 2.07 | 1.82 |
| 53 | 1.83 | 1.61 | 1.81 | 2.07 | 1.85 |
| 54 | 1.83 | 1.64 | 1.83 | 2.07 | 1.88 |
| 55 | 1.83 | 1.67 | 1.83 | 2.07 | 1.91 |
| 56 | 1.86 | 1.70 | 1.83 | 2.07 | 1.94 |
| 57 | 1.96 | 1.73 | 1.83 | 2.07 | 1.96 |
| 58 | 1.96 | 1.76 | 1.83 | 2.07 | 1.97 |
| 59 | 1.96 | 1.80 | 1.83 | 2.07 | 1.98 |
| 60 | 1.97 | 1.83 | 1.83 | 2.07 | 1.99 |
| 61 | 2.00 | 1.83 | 1.83 | 2.13 | 2.00 |
| 62 | 2.00 | 1.83 | 1.83 | 2.19 | 2.02 |
| 63 | 2.04 | 1.83 | 1.83 | 2.25 | 2.03 |
| 64 | 2.07 | 1.83 | 1.83 | 2.31 | 2.05 |
| 65 | 2.07 | 1.87 | 1.85 | 2.36 | 2.07 |
| 66 | 2.07 | 1.91 | 1.89 | 2.41 | 2.10 |

| | | | | | |
|---|---|---|---|---|---|
| 67 | 2.07 | 1.96 | 1.93 | 2.46 | 2.13 |
| 68 | 2.07 | 2.00 | 1.96 | 2.52 | 2.16 |
| 69 | 2.18 | 2.00 | 1.99 | 2.52 | 2.20 |
| 70 | 2.22 | 2.00 | 2.02 | 2.52 | 2.25 |
| 71 | 2.22 | 2.00 | 2.06 | 2.53 | 2.32 |
| 72 | 2.30 | 2.00 | 2.09 | 2.53 | 2.39 |
| 73 | 2.31 | 2.05 | 2.13 | 2.54 | 2.46 |
| 74 | 2.51 | 2.11 | 2.18 | 2.55 | 2.53 |
| 75 | 2.53 | 2.16 | 2.22 | 2.56 | 2.54 |
| 76 | 2.53 | 2.22 | 2.24 | 2.57 | 2.55 |
| 77 | 2.57 | 2.39 | 2.27 | 2.60 | 2.56 |
| 78 | 2.57 | 2.56 | 2.30 | 2.63 | 2.57 |
| 79 | 2.57 | 2.74 | 2.34 | 2.66 | 2.57 |
| 80 | 2.57 | 2.91 | 2.42 | 2.69 | 2.57 |
| 81 | 2.57 | 3.00 | 2.49 | 2.74 | 2.57 |
| 82 | 2.57 | 3.09 | 2.56 | 2.78 | 2.57 |
| 83 | 2.61 | 3.18 | 2.57 | 2.83 | 2.60 |
| 84 | 2.76 | 3.27 | 2.57 | 2.87 | 2.68 |
| 85 | 2.87 | 3.27 | 2.57 | 2.87 | 2.76 |
| 86 | 2.89 | 3.27 | 2.57 | 2.87 | 2.84 |
| 87 | 2.91 | 3.27 | 2.57 | 2.87 | 2.92 |
| 88 | 3.10 | 3.27 | 2.57 | 2.87 | 3.00 |
| 89 | 3.27 | 3.30 | 2.57 | 2.97 | 3.08 |
| 90 | 3.27 | 3.34 | 2.57 | 3.07 | 3.16 |
| 91 | 3.27 | 3.37 | 2.57 | 3.17 | 3.24 |
| 92 | 3.27 | 3.40 | 2.57 | 3.27 | 3.41 |
| 93 | 3.27 | 3.50 | 2.60 | 3.27 | 3.61 |
| 94 | 3.27 | 3.59 | 2.80 | 3.27 | 3.81 |
| 95 | 3.38 | 3.69 | 2.99 | 3.27 | 4.01 |
| 96 | 3.72 | 3.78 | 3.18 | 3.27 | 4.14 |
| 97 | 4.10 | 3.87 | 3.41 | 3.27 | 4.14 |
| 98 | 4.14 | 3.96 | 3.65 | 3.27 | 4.14 |
| 99 | 4.14 | 4.05 | 3.90 | 3.27 | 4.14 |
| 100 | 4.14 | 4.14 | 4.14 | 3.27 | 4.14 |

***Table S62:*** *Adaptive tapping – Sensitivity index (d') of perceiving tempo acceleration (IOI - 75ms) – Adaptive_minus_75_dprime2*

| **Task** | Adaptive tapping | | | | |
|---|---|---|---|---|---|
| **Outcome measure** | Sensitivity index (d') of perceiving tempo acceleration (IOI - 75ms) | | | | |
| **Variable name** | Adaptive_minus_75_dprime2 | | | | |
| **Age and gender effects** | | | | | |
| Age regression (slope) | 0.000 | | | | |
| Age regression (p) | 1.000 | | | | |
| Age group (p) | 1.000 | | | | |
| Gender (p) | 1.000 | | | | |
| **Group** | All | Age 18 to 21 | Age 22 to 29 | Age 30 to 54 | Age 55 to 87 |
| N (N female) | 105 (72) | 26 (17) | 29 (19) | 26 (16) | 24 (20) |
| **Normality** | | | | | |
| Skewness | -2.21 | -1.56 | -1.65 | -2.01 | -1.20 |
| Excess kurtosis | 6.27 | 1.77 | 1.38 | 4.09 | 0.26 |
| **Scores** | | | | | |
| Mean | 3.79 | 3.75 | 3.92 | 3.61 | 3.86 |
| SD | 0.54 | 0.55 | 0.39 | 0.74 | 0.39 |
| Percentiles | | | | | |
| 0 | 1.04 | 2.09 | 2.91 | 1.04 | 2.96 |
| 1 | 2.09 | 2.21 | 2.91 | 1.32 | 2.96 |
| 2 | 2.17 | 2.33 | 2.91 | 1.59 | 2.96 |
| 3 | 2.57 | 2.45 | 2.91 | 1.86 | 2.96 |
| 4 | 2.63 | 2.57 | 2.94 | 2.13 | 2.96 |
| 5 | 2.91 | 2.66 | 3.01 | 2.24 | 3.00 |
| 6 | 2.91 | 2.74 | 3.08 | 2.35 | 3.08 |
| 7 | 2.92 | 2.83 | 3.14 | 2.46 | 3.15 |
| 8 | 2.96 | 2.91 | 3.18 | 2.57 | 3.22 |
| 9 | 3.03 | 3.00 | 3.21 | 2.75 | 3.27 |
| 10 | 3.20 | 3.09 | 3.25 | 2.92 | 3.27 |
| 11 | 3.27 | 3.18 | 3.29 | 3.09 | 3.27 |
| 12 | 3.27 | 3.27 | 3.38 | 3.27 | 3.27 |
| 13 | 3.27 | 3.27 | 3.46 | 3.27 | 3.27 |
| 14 | 3.27 | 3.27 | 3.54 | 3.27 | 3.33 |
| 15 | 3.27 | 3.27 | 3.61 | 3.27 | 3.40 |
| 16 | 3.27 | 3.27 | 3.67 | 3.27 | 3.47 |
| 17 | 3.27 | 3.27 | 3.73 | 3.27 | 3.54 |
| 18 | 3.27 | 3.27 | 3.78 | 3.27 | 3.60 |
| 19 | 3.27 | 3.27 | 3.78 | 3.27 | 3.65 |
| 20 | 3.51 | 3.27 | 3.78 | 3.27 | 3.70 |
| 21 | 3.57 | 3.34 | 3.78 | 3.27 | 3.75 |
| 22 | 3.57 | 3.42 | 3.78 | 3.27 | 3.78 |

| | | | | | |
|---|---|---|---|---|---|
| 23 | 3.57 | 3.49 | 3.78 | 3.27 | 3.78 |
| 24 | 3.57 | 3.57 | 3.78 | 3.27 | 3.78 |
| 25 | 3.57 | 3.62 | 3.78 | 3.34 | 3.78 |
| 26 | 3.57 | 3.67 | 3.78 | 3.42 | 3.78 |
| 27 | 3.78 | 3.73 | 3.78 | 3.49 | 3.78 |
| 28 | 3.78 | 3.78 | 3.78 | 3.57 | 3.78 |
| 29 | 3.78 | 3.78 | 3.83 | 3.57 | 3.78 |
| 30 | 3.78 | 3.78 | 3.93 | 3.57 | 3.78 |
| 31 | 3.78 | 3.78 | 4.03 | 3.57 | 3.78 |
| 32 | 3.78 | 3.78 | 4.13 | 3.57 | 3.78 |
| 33 | 3.78 | 3.78 | 4.14 | 3.57 | 3.78 |
| 34 | 3.78 | 3.78 | 4.14 | 3.57 | 3.78 |
| 35 | 3.78 | 3.78 | 4.14 | 3.57 | 3.78 |
| 36 | 3.78 | 3.78 | 4.14 | 3.57 | 3.78 |
| 37 | 3.78 | 3.78 | 4.14 | 3.57 | 3.78 |
| 38 | 3.78 | 3.78 | 4.14 | 3.57 | 3.78 |
| 39 | 3.78 | 3.78 | 4.14 | 3.57 | 3.78 |
| 40 | 3.78 | 3.78 | 4.14 | 3.57 | 3.78 |
| 41 | 3.78 | 3.78 | 4.14 | 3.62 | 3.78 |
| 42 | 3.78 | 3.78 | 4.14 | 3.67 | 3.78 |
| 43 | 3.78 | 3.78 | 4.14 | 3.73 | 3.78 |
| 44 | 3.78 | 3.78 | 4.14 | 3.78 | 3.83 |
| 45 | 3.78 | 3.78 | 4.14 | 3.78 | 3.91 |
| 46 | 4.08 | 3.78 | 4.14 | 3.78 | 3.99 |
| 47 | 4.14 | 3.78 | 4.14 | 3.78 | 4.07 |
| 48 | 4.14 | 3.78 | 4.14 | 3.78 | 4.14 |
| 49 | 4.14 | 3.87 | 4.14 | 3.78 | 4.14 |
| 50 | 4.14 | 3.96 | 4.14 | 3.78 | 4.14 |
| 51 | 4.14 | 4.05 | 4.14 | 3.78 | 4.14 |
| 52 | 4.14 | 4.14 | 4.14 | 3.78 | 4.14 |
| 53 | 4.14 | 4.14 | 4.14 | 3.78 | 4.14 |
| 54 | 4.14 | 4.14 | 4.14 | 3.78 | 4.14 |
| 55 | 4.14 | 4.14 | 4.14 | 3.78 | 4.14 |
| 56 | 4.14 | 4.14 | 4.14 | 3.78 | 4.14 |
| 57 | 4.14 | 4.14 | 4.14 | 3.87 | 4.14 |
| 58 | 4.14 | 4.14 | 4.14 | 3.96 | 4.14 |
| 59 | 4.14 | 4.14 | 4.14 | 4.05 | 4.14 |
| 60 | 4.14 | 4.14 | 4.14 | 4.14 | 4.14 |
| 61 | 4.14 | 4.14 | 4.14 | 4.14 | 4.14 |
| 62 | 4.14 | 4.14 | 4.14 | 4.14 | 4.14 |
| 63 | 4.14 | 4.14 | 4.14 | 4.14 | 4.14 |
| 64 | 4.14 | 4.14 | 4.14 | 4.14 | 4.14 |
| 65 | 4.14 | 4.14 | 4.14 | 4.14 | 4.14 |
| 66 | 4.14 | 4.14 | 4.14 | 4.14 | 4.14 |

| | | | | | |
|---|---|---|---|---|---|
| 67 | 4.14 | 4.14 | 4.14 | 4.14 | 4.14 |
| 68 | 4.14 | 4.14 | 4.14 | 4.14 | 4.14 |
| 69 | 4.14 | 4.14 | 4.14 | 4.14 | 4.14 |
| 70 | 4.14 | 4.14 | 4.14 | 4.14 | 4.14 |
| 71 | 4.14 | 4.14 | 4.14 | 4.14 | 4.14 |
| 72 | 4.14 | 4.14 | 4.14 | 4.14 | 4.14 |
| 73 | 4.14 | 4.14 | 4.14 | 4.14 | 4.14 |
| 74 | 4.14 | 4.14 | 4.14 | 4.14 | 4.14 |
| 75 | 4.14 | 4.14 | 4.14 | 4.14 | 4.14 |
| 76 | 4.14 | 4.14 | 4.14 | 4.14 | 4.14 |
| 77 | 4.14 | 4.14 | 4.14 | 4.14 | 4.14 |
| 78 | 4.14 | 4.14 | 4.14 | 4.14 | 4.14 |
| 79 | 4.14 | 4.14 | 4.14 | 4.14 | 4.14 |
| 80 | 4.14 | 4.14 | 4.14 | 4.14 | 4.14 |
| 81 | 4.14 | 4.14 | 4.14 | 4.14 | 4.14 |
| 82 | 4.14 | 4.14 | 4.14 | 4.14 | 4.14 |
| 83 | 4.14 | 4.14 | 4.14 | 4.14 | 4.14 |
| 84 | 4.14 | 4.14 | 4.14 | 4.14 | 4.14 |
| 85 | 4.14 | 4.14 | 4.14 | 4.14 | 4.14 |
| 86 | 4.14 | 4.14 | 4.14 | 4.14 | 4.14 |
| 87 | 4.14 | 4.14 | 4.14 | 4.14 | 4.14 |
| 88 | 4.14 | 4.14 | 4.14 | 4.14 | 4.14 |
| 89 | 4.14 | 4.14 | 4.14 | 4.14 | 4.14 |
| 90 | 4.14 | 4.14 | 4.14 | 4.14 | 4.14 |
| 91 | 4.14 | 4.14 | 4.14 | 4.14 | 4.14 |
| 92 | 4.14 | 4.14 | 4.14 | 4.14 | 4.14 |
| 93 | 4.14 | 4.14 | 4.14 | 4.14 | 4.14 |
| 94 | 4.14 | 4.14 | 4.14 | 4.14 | 4.14 |
| 95 | 4.14 | 4.14 | 4.14 | 4.14 | 4.14 |
| 96 | 4.14 | 4.14 | 4.14 | 4.14 | 4.14 |
| 97 | 4.14 | 4.14 | 4.14 | 4.14 | 4.14 |
| 98 | 4.14 | 4.14 | 4.14 | 4.14 | 4.14 |
| 99 | 4.14 | 4.14 | 4.14 | 4.14 | 4.14 |
| 100 | 4.14 | 4.14 | 4.14 | 4.14 | 4.14 |

***Table S63:*** *Adaptive tapping – Sensitivity index (d') perceiving a tempo acceleration (IOI - 30ms) – Adaptive_minus_30_dprime2*

| | | | | | | |
|---|---|---|---|---|---|---|
| **Task** | | Adaptive tapping | | | | |
| **Outcome measure** | | Sensitivity index (d') perceiving a tempo acceleration (IOI - 30ms) | | | | |
| **Variable name** | | Adaptive_minus_30_dprime2 | | | | |
| **Age and gender effects** | | | | | | |
| Age regression (slope) | | 0.004 | | | | |
| Age regression (p) | | 0.313 | | | | |
| Age group (p) | | 0.373 | | | | |
| Gender (p) | | 0.377 | | | | |
| **Group** | | All | Age 18 to 21 | Age 22 to 29 | Age 30 to 54 | Age 55 to 87 |
| N (N female) | | 105 (72) | 26 (17) | 29 (19) | 26 (16) | 24 (20) |
| **Normality** | | | | | | |
| Skewness | | -0.09 | 0.76 | 0.09 | -0.44 | -1.04 |
| Excess kurtosis | | -0.07 | 0.56 | 0.18 | -0.94 | 0.57 |
| **Scores** | | | | | | |
| Mean | | 1.69 | 1.54 | 1.79 | 1.56 | 1.86 |
| SD | | 0.88 | 0.83 | 0.97 | 0.92 | 0.78 |
| Percentiles | | | | | | |
| | 0 | -0.36 | 0.00 | -0.36 | 0.00 | 0.00 |
| | 1 | 0.00 | 0.13 | -0.26 | 0.00 | 0.00 |
| | 2 | 0.00 | 0.26 | -0.16 | 0.00 | 0.00 |
| | 3 | 0.00 | 0.39 | -0.06 | 0.00 | 0.00 |
| | 4 | 0.00 | 0.52 | 0.10 | 0.00 | 0.00 |
| | 5 | 0.00 | 0.60 | 0.35 | 0.00 | 0.13 |
| | 6 | 0.00 | 0.69 | 0.59 | 0.00 | 0.33 |
| | 7 | 0.00 | 0.78 | 0.84 | 0.00 | 0.53 |
| | 8 | 0.16 | 0.87 | 0.87 | 0.00 | 0.73 |
| | 9 | 0.52 | 0.87 | 0.87 | 0.00 | 0.87 |
| | 10 | 0.67 | 0.87 | 0.87 | 0.00 | 0.88 |
| | 11 | 0.87 | 0.87 | 0.87 | 0.00 | 0.88 |
| | 12 | 0.87 | 0.87 | 0.87 | 0.00 | 0.88 |
| | 13 | 0.87 | 0.87 | 0.87 | 0.13 | 0.89 |
| | 14 | 0.87 | 0.87 | 0.87 | 0.27 | 0.97 |
| | 15 | 0.87 | 0.87 | 0.87 | 0.40 | 1.05 |
| | 16 | 0.87 | 0.87 | 0.87 | 0.53 | 1.14 |
| | 17 | 0.87 | 0.87 | 0.87 | 0.62 | 1.22 |
| | 18 | 0.87 | 0.87 | 0.87 | 0.70 | 1.28 |
| | 19 | 0.87 | 0.87 | 0.88 | 0.79 | 1.33 |
| | 20 | 0.87 | 0.87 | 0.90 | 0.87 | 1.38 |
| | 21 | 0.88 | 0.87 | 0.91 | 0.88 | 1.43 |
| | 22 | 0.89 | 0.87 | 0.96 | 0.89 | 1.48 |

| | | | | | |
|---|---|---|---|---|---|
| 23 | 0.91 | 0.87 | 1.04 | 0.90 | 1.50 |
| 24 | 0.91 | 0.87 | 1.13 | 0.91 | 1.52 |
| 25 | 1.21 | 0.88 | 1.21 | 1.00 | 1.54 |
| 26 | 1.21 | 0.88 | 1.21 | 1.08 | 1.57 |
| 27 | 1.21 | 0.88 | 1.21 | 1.17 | 1.62 |
| 28 | 1.22 | 0.89 | 1.21 | 1.25 | 1.68 |
| 29 | 1.25 | 0.97 | 1.22 | 1.26 | 1.74 |
| 30 | 1.26 | 1.05 | 1.23 | 1.26 | 1.80 |
| 31 | 1.27 | 1.13 | 1.25 | 1.26 | 1.83 |
| 32 | 1.27 | 1.21 | 1.27 | 1.27 | 1.83 |
| 33 | 1.27 | 1.21 | 1.32 | 1.27 | 1.83 |
| 34 | 1.27 | 1.21 | 1.37 | 1.27 | 1.83 |
| 35 | 1.27 | 1.21 | 1.43 | 1.27 | 1.83 |
| 36 | 1.27 | 1.21 | 1.48 | 1.27 | 1.86 |
| 37 | 1.37 | 1.23 | 1.51 | 1.27 | 1.89 |
| 38 | 1.47 | 1.24 | 1.53 | 1.27 | 1.92 |
| 39 | 1.48 | 1.25 | 1.56 | 1.27 | 1.95 |
| 40 | 1.54 | 1.27 | 1.57 | 1.27 | 1.96 |
| 41 | 1.57 | 1.27 | 1.57 | 1.33 | 1.96 |
| 42 | 1.57 | 1.27 | 1.57 | 1.38 | 1.96 |
| 43 | 1.57 | 1.27 | 1.58 | 1.44 | 1.96 |
| 44 | 1.57 | 1.27 | 1.65 | 1.50 | 1.97 |
| 45 | 1.57 | 1.27 | 1.72 | 1.51 | 2.00 |
| 46 | 1.57 | 1.27 | 1.80 | 1.53 | 2.02 |
| 47 | 1.57 | 1.27 | 1.83 | 1.55 | 2.05 |
| 48 | 1.69 | 1.27 | 1.83 | 1.57 | 2.07 |
| 49 | 1.74 | 1.34 | 1.83 | 1.63 | 2.07 |
| 50 | 1.83 | 1.42 | 1.83 | 1.70 | 2.07 |
| 51 | 1.83 | 1.49 | 1.84 | 1.76 | 2.07 |
| 52 | 1.83 | 1.57 | 1.86 | 1.83 | 2.07 |
| 53 | 1.83 | 1.57 | 1.87 | 1.86 | 2.07 |
| 54 | 1.83 | 1.57 | 1.91 | 1.89 | 2.07 |
| 55 | 1.84 | 1.57 | 1.96 | 1.92 | 2.07 |
| 56 | 1.90 | 1.57 | 2.01 | 1.96 | 2.07 |
| 57 | 1.96 | 1.57 | 2.06 | 1.96 | 2.09 |
| 58 | 1.96 | 1.57 | 2.07 | 1.96 | 2.12 |
| 59 | 1.96 | 1.57 | 2.07 | 1.96 | 2.15 |
| 60 | 1.97 | 1.57 | 2.07 | 1.96 | 2.19 |
| 61 | 2.03 | 1.57 | 2.07 | 1.97 | 2.22 |
| 62 | 2.07 | 1.57 | 2.07 | 1.98 | 2.22 |
| 63 | 2.07 | 1.57 | 2.07 | 1.99 | 2.22 |
| 64 | 2.07 | 1.57 | 2.07 | 2.00 | 2.22 |
| 65 | 2.07 | 1.60 | 2.10 | 2.02 | 2.22 |
| 66 | 2.07 | 1.63 | 2.14 | 2.03 | 2.23 |

| | | | | | |
|---|---|---|---|---|---|
| 67 | 2.07 | 1.67 | 2.18 | 2.05 | 2.26 |
| 68 | 2.18 | 1.70 | 2.22 | 2.07 | 2.28 |
| 69 | 2.22 | 1.71 | 2.24 | 2.11 | 2.30 |
| 70 | 2.22 | 1.72 | 2.26 | 2.14 | 2.31 |
| 71 | 2.22 | 1.73 | 2.29 | 2.18 | 2.31 |
| 72 | 2.22 | 1.74 | 2.30 | 2.22 | 2.31 |
| 73 | 2.29 | 1.76 | 2.30 | 2.24 | 2.31 |
| 74 | 2.30 | 1.78 | 2.31 | 2.26 | 2.31 |
| 75 | 2.31 | 1.80 | 2.31 | 2.28 | 2.31 |
| 76 | 2.31 | 1.83 | 2.31 | 2.30 | 2.31 |
| 77 | 2.31 | 1.92 | 2.31 | 2.37 | 2.31 |
| 78 | 2.31 | 2.02 | 2.31 | 2.43 | 2.31 |
| 79 | 2.31 | 2.12 | 2.34 | 2.50 | 2.31 |
| 80 | 2.35 | 2.22 | 2.42 | 2.57 | 2.31 |
| 81 | 2.53 | 2.29 | 2.49 | 2.57 | 2.31 |
| 82 | 2.57 | 2.37 | 2.56 | 2.57 | 2.31 |
| 83 | 2.57 | 2.44 | 2.57 | 2.57 | 2.34 |
| 84 | 2.57 | 2.52 | 2.57 | 2.57 | 2.40 |
| 85 | 2.57 | 2.53 | 2.57 | 2.57 | 2.46 |
| 86 | 2.57 | 2.54 | 2.60 | 2.57 | 2.52 |
| 87 | 2.57 | 2.56 | 2.68 | 2.57 | 2.57 |
| 88 | 2.57 | 2.57 | 2.76 | 2.57 | 2.57 |
| 89 | 2.57 | 2.57 | 2.85 | 2.57 | 2.57 |
| 90 | 2.57 | 2.57 | 2.95 | 2.57 | 2.57 |
| 91 | 2.65 | 2.57 | 3.06 | 2.57 | 2.57 |
| 92 | 2.81 | 2.57 | 3.17 | 2.57 | 2.63 |
| 93 | 2.87 | 2.66 | 3.27 | 2.60 | 2.70 |
| 94 | 2.90 | 2.74 | 3.27 | 2.63 | 2.78 |
| 95 | 2.91 | 2.83 | 3.27 | 2.66 | 2.86 |
| 96 | 2.95 | 2.91 | 3.27 | 2.69 | 2.92 |
| 97 | 3.23 | 3.13 | 3.41 | 2.74 | 2.93 |
| 98 | 3.27 | 3.35 | 3.65 | 2.78 | 2.94 |
| 99 | 3.76 | 3.57 | 3.90 | 2.83 | 2.95 |
| 100 | 4.14 | 3.78 | 4.14 | 2.87 | 2.96 |

***Table S64:*** *Adaptive tapping – Variability (CV of ITI) of continuation tapping (isochronous condition) – Adaptive_iso_600_CV_iti*

| | | | | | |
|---|---|---|---|---|---|
| **Task** | Adaptive tapping | | | | |
| **Outcome measure** | Variability (CV of ITI) of continuation tapping (isochronous condition) | | | | |
| **Variable name** | Adaptive_iso_600_CV_iti | | | | |
| **Age and gender effects** | | | | | |
| Age regression (slope) | 0.000 | | | | |
| Age regression (p) | 0.054 | | | | |
| Age group (p) | 0.213 | | | | |
| Gender (p) | 0.222 | | | | |
| **Group** | All | Age 18 to 21 | Age 22 to 29 | Age 30 to 54 | Age 55 to 87 |
| N (N female) | 105 (72) | 26 (17) | 29 (19) | 26 (16) | 24 (20) |
| **Normality** | | | | | |
| Skewness | 2.39 | 1.12 | 1.70 | 2.07 | 0.22 |
| Excess kurtosis | 8.42 | 1.03 | 3.10 | 4.52 | -0.28 |
| **Scores** | | | | | |
| Mean | 0.057 | 0.058 | 0.059 | 0.062 | 0.049 |
| SD | 0.020 | 0.015 | 0.019 | 0.029 | 0.010 |
| Percentiles | | | | | |
| 0 | 0.031 | 0.035 | 0.037 | 0.033 | 0.031 |
| 1 | 0.033 | 0.037 | 0.037 | 0.033 | 0.032 |
| 2 | 0.033 | 0.038 | 0.037 | 0.033 | 0.032 |
| 3 | 0.034 | 0.039 | 0.037 | 0.033 | 0.033 |
| 4 | 0.035 | 0.040 | 0.037 | 0.033 | 0.033 |
| 5 | 0.036 | 0.040 | 0.039 | 0.035 | 0.034 |
| 6 | 0.037 | 0.041 | 0.040 | 0.036 | 0.034 |
| 7 | 0.037 | 0.041 | 0.042 | 0.037 | 0.034 |
| 8 | 0.038 | 0.042 | 0.042 | 0.039 | 0.035 |
| 9 | 0.039 | 0.042 | 0.042 | 0.039 | 0.035 |
| 10 | 0.039 | 0.042 | 0.043 | 0.039 | 0.036 |
| 11 | 0.040 | 0.043 | 0.043 | 0.040 | 0.036 |
| 12 | 0.041 | 0.043 | 0.043 | 0.040 | 0.037 |
| 13 | 0.042 | 0.043 | 0.043 | 0.040 | 0.038 |
| 14 | 0.042 | 0.044 | 0.043 | 0.041 | 0.038 |
| 15 | 0.042 | 0.045 | 0.044 | 0.041 | 0.038 |
| 16 | 0.042 | 0.045 | 0.044 | 0.041 | 0.039 |
| 17 | 0.043 | 0.046 | 0.044 | 0.042 | 0.039 |
| 18 | 0.043 | 0.046 | 0.044 | 0.042 | 0.040 |
| 19 | 0.043 | 0.046 | 0.044 | 0.042 | 0.040 |
| 20 | 0.044 | 0.047 | 0.044 | 0.042 | 0.041 |
| 21 | 0.044 | 0.047 | 0.044 | 0.042 | 0.042 |

| | | | | | |
|---|---|---|---|---|---|
| 22 | 0.044 | 0.047 | 0.044 | 0.043 | 0.042 |
| 23 | 0.044 | 0.047 | 0.045 | 0.043 | 0.043 |
| 24 | 0.044 | 0.047 | 0.045 | 0.044 | 0.043 |
| 25 | 0.044 | 0.047 | 0.046 | 0.044 | 0.043 |
| 26 | 0.045 | 0.047 | 0.047 | 0.044 | 0.044 |
| 27 | 0.046 | 0.047 | 0.048 | 0.044 | 0.044 |
| 28 | 0.046 | 0.048 | 0.049 | 0.044 | 0.044 |
| 29 | 0.046 | 0.049 | 0.049 | 0.045 | 0.044 |
| 30 | 0.046 | 0.049 | 0.049 | 0.046 | 0.044 |
| 31 | 0.047 | 0.050 | 0.049 | 0.047 | 0.045 |
| 32 | 0.047 | 0.051 | 0.049 | 0.048 | 0.045 |
| 33 | 0.048 | 0.052 | 0.049 | 0.048 | 0.045 |
| 34 | 0.048 | 0.052 | 0.049 | 0.048 | 0.045 |
| 35 | 0.049 | 0.053 | 0.050 | 0.048 | 0.046 |
| 36 | 0.049 | 0.053 | 0.050 | 0.048 | 0.046 |
| 37 | 0.049 | 0.053 | 0.050 | 0.049 | 0.046 |
| 38 | 0.050 | 0.053 | 0.050 | 0.049 | 0.046 |
| 39 | 0.050 | 0.053 | 0.050 | 0.049 | 0.046 |
| 40 | 0.050 | 0.053 | 0.050 | 0.050 | 0.046 |
| 41 | 0.050 | 0.054 | 0.050 | 0.050 | 0.046 |
| 42 | 0.050 | 0.054 | 0.050 | 0.050 | 0.046 |
| 43 | 0.050 | 0.054 | 0.050 | 0.050 | 0.046 |
| 44 | 0.050 | 0.054 | 0.051 | 0.050 | 0.047 |
| 45 | 0.051 | 0.054 | 0.051 | 0.050 | 0.048 |
| 46 | 0.051 | 0.054 | 0.051 | 0.050 | 0.049 |
| 47 | 0.051 | 0.054 | 0.051 | 0.050 | 0.049 |
| 48 | 0.051 | 0.054 | 0.052 | 0.050 | 0.050 |
| 49 | 0.051 | 0.054 | 0.053 | 0.050 | 0.050 |
| 50 | 0.052 | 0.054 | 0.054 | 0.050 | 0.050 |
| 51 | 0.053 | 0.055 | 0.054 | 0.050 | 0.050 |
| 52 | 0.053 | 0.055 | 0.054 | 0.051 | 0.050 |
| 53 | 0.053 | 0.055 | 0.055 | 0.051 | 0.051 |
| 54 | 0.054 | 0.055 | 0.056 | 0.052 | 0.051 |
| 55 | 0.054 | 0.055 | 0.057 | 0.053 | 0.051 |
| 56 | 0.054 | 0.055 | 0.059 | 0.053 | 0.051 |
| 57 | 0.054 | 0.056 | 0.060 | 0.055 | 0.051 |
| 58 | 0.055 | 0.057 | 0.060 | 0.056 | 0.051 |
| 59 | 0.055 | 0.058 | 0.061 | 0.057 | 0.051 |
| 60 | 0.055 | 0.059 | 0.061 | 0.058 | 0.051 |
| 61 | 0.055 | 0.060 | 0.061 | 0.059 | 0.051 |
| 62 | 0.056 | 0.060 | 0.061 | 0.060 | 0.052 |
| 63 | 0.057 | 0.060 | 0.061 | 0.061 | 0.052 |
| 64 | 0.058 | 0.060 | 0.061 | 0.062 | 0.052 |
| 65 | 0.059 | 0.060 | 0.062 | 0.063 | 0.052 |

| | | | | | |
|---|---|---|---|---|---|
| 66 | 0.060 | 0.060 | 0.062 | 0.064 | 0.053 |
| 67 | 0.060 | 0.061 | 0.062 | 0.066 | 0.053 |
| 68 | 0.061 | 0.061 | 0.063 | 0.067 | 0.053 |
| 69 | 0.061 | 0.061 | 0.063 | 0.067 | 0.053 |
| 70 | 0.061 | 0.061 | 0.064 | 0.067 | 0.053 |
| 71 | 0.062 | 0.062 | 0.064 | 0.067 | 0.054 |
| 72 | 0.062 | 0.062 | 0.064 | 0.067 | 0.054 |
| 73 | 0.062 | 0.063 | 0.065 | 0.068 | 0.054 |
| 74 | 0.063 | 0.064 | 0.065 | 0.069 | 0.054 |
| 75 | 0.064 | 0.065 | 0.065 | 0.069 | 0.055 |
| 76 | 0.065 | 0.066 | 0.065 | 0.070 | 0.055 |
| 77 | 0.065 | 0.066 | 0.065 | 0.071 | 0.055 |
| 78 | 0.065 | 0.066 | 0.065 | 0.072 | 0.056 |
| 79 | 0.066 | 0.067 | 0.065 | 0.073 | 0.056 |
| 80 | 0.066 | 0.067 | 0.065 | 0.073 | 0.056 |
| 81 | 0.067 | 0.067 | 0.065 | 0.075 | 0.056 |
| 82 | 0.067 | 0.067 | 0.065 | 0.077 | 0.056 |
| 83 | 0.067 | 0.067 | 0.066 | 0.078 | 0.056 |
| 84 | 0.068 | 0.067 | 0.067 | 0.080 | 0.057 |
| 85 | 0.069 | 0.068 | 0.068 | 0.081 | 0.057 |
| 86 | 0.071 | 0.069 | 0.069 | 0.082 | 0.057 |
| 87 | 0.072 | 0.070 | 0.071 | 0.083 | 0.058 |
| 88 | 0.073 | 0.071 | 0.073 | 0.084 | 0.059 |
| 89 | 0.075 | 0.074 | 0.075 | 0.085 | 0.060 |
| 90 | 0.078 | 0.076 | 0.078 | 0.087 | 0.061 |
| 91 | 0.080 | 0.078 | 0.081 | 0.089 | 0.062 |
| 92 | 0.082 | 0.080 | 0.083 | 0.091 | 0.063 |
| 93 | 0.085 | 0.084 | 0.086 | 0.099 | 0.064 |
| 94 | 0.090 | 0.087 | 0.091 | 0.107 | 0.065 |
| 95 | 0.093 | 0.090 | 0.095 | 0.115 | 0.066 |
| 96 | 0.098 | 0.094 | 0.100 | 0.123 | 0.067 |
| 97 | 0.102 | 0.095 | 0.105 | 0.134 | 0.068 |
| 98 | 0.120 | 0.097 | 0.111 | 0.144 | 0.069 |
| 99 | 0.123 | 0.098 | 0.116 | 0.155 | 0.070 |
| 100 | 0.166 | 0.099 | 0.122 | 0.166 | 0.072 |

***Table S65:*** *Adaptive tapping – Adaptation index (acceleration trials) – Adaptive_adaptation_index_acceleration*

| | | | | | | |
|---|---|---|---|---|---|---|
| **Task** | | Adaptive tapping | | | | |
| **Outcome measure** | | Adaptation index (acceleration trials) | | | | |
| **Variable name** | | Adaptive_adaptation_index_acceleration | | | | |
| **Age and gender effects** | | | | | | |
| Age regression (slope) | | 0.001 | | | | |
| Age regression (p) | | 0.749 | | | | |
| Age group (p) | | 0.653 | | | | |
| Gender (p) | | 0.691 | | | | |
| **Group** | | All | Age 18 to 21 | Age 22 to 29 | Age 30 to 54 | Age 55 to 87 |
| N (N female) | | 105 (72) | 26 (17) | 29 (19) | 26 (16) | 24 (20) |
| **Normality** | | | | | | |
| Skewness | | 2.18 | 0.17 | 2.05 | 2.47 | 1.20 |
| Excess kurtosis | | 8.00 | -0.46 | 5.37 | 7.69 | 1.55 |
| **Scores** | | | | | | |
| Mean | | 1.442 | 1.432 | 1.407 | 1.495 | 1.439 |
| SD | | 0.597 | 0.417 | 0.655 | 0.770 | 0.501 |
| Percentiles | | | | | | |
| | 0 | 0.403 | 0.767 | 0.403 | 0.580 | 0.784 |
| | 1 | 0.580 | 0.772 | 0.493 | 0.580 | 0.797 |
| | 2 | 0.593 | 0.777 | 0.582 | 0.581 | 0.809 |
| | 3 | 0.728 | 0.782 | 0.671 | 0.581 | 0.821 |
| | 4 | 0.770 | 0.786 | 0.734 | 0.582 | 0.833 |
| | 5 | 0.785 | 0.788 | 0.762 | 0.643 | 0.839 |
| | 6 | 0.788 | 0.790 | 0.789 | 0.704 | 0.842 |
| | 7 | 0.801 | 0.792 | 0.816 | 0.764 | 0.846 |
| | 8 | 0.822 | 0.794 | 0.838 | 0.825 | 0.849 |
| | 9 | 0.830 | 0.829 | 0.858 | 0.854 | 0.853 |
| | 10 | 0.843 | 0.864 | 0.879 | 0.882 | 0.861 |
| | 11 | 0.866 | 0.899 | 0.897 | 0.911 | 0.869 |
| | 12 | 0.889 | 0.935 | 0.908 | 0.939 | 0.877 |
| | 13 | 0.915 | 0.944 | 0.920 | 0.948 | 0.885 |
| | 14 | 0.935 | 0.953 | 0.931 | 0.957 | 0.911 |
| | 15 | 0.937 | 0.962 | 0.949 | 0.966 | 0.937 |
| | 16 | 0.959 | 0.971 | 0.969 | 0.975 | 0.964 |
| | 17 | 0.973 | 0.986 | 0.989 | 0.980 | 0.990 |
| | 18 | 0.989 | 1.001 | 1.010 | 0.985 | 1.002 |
| | 19 | 0.999 | 1.016 | 1.042 | 0.990 | 1.005 |
| | 20 | 1.005 | 1.031 | 1.074 | 0.994 | 1.008 |
| | 21 | 1.012 | 1.041 | 1.106 | 1.004 | 1.011 |
| | 22 | 1.029 | 1.050 | 1.122 | 1.014 | 1.024 |

| | | | | | |
|---|---|---|---|---|---|
| 23 | 1.033 | 1.060 | 1.125 | 1.023 | 1.064 |
| 24 | 1.069 | 1.070 | 1.129 | 1.033 | 1.103 |
| 25 | 1.120 | 1.107 | 1.132 | 1.063 | 1.143 |
| 26 | 1.133 | 1.144 | 1.139 | 1.094 | 1.183 |
| 27 | 1.155 | 1.180 | 1.145 | 1.124 | 1.207 |
| 28 | 1.156 | 1.217 | 1.152 | 1.155 | 1.230 |
| 29 | 1.160 | 1.223 | 1.156 | 1.171 | 1.252 |
| 30 | 1.173 | 1.228 | 1.157 | 1.188 | 1.275 |
| 31 | 1.182 | 1.234 | 1.157 | 1.205 | 1.286 |
| 32 | 1.189 | 1.240 | 1.158 | 1.221 | 1.288 |
| 33 | 1.201 | 1.249 | 1.161 | 1.230 | 1.291 |
| 34 | 1.213 | 1.258 | 1.164 | 1.239 | 1.293 |
| 35 | 1.219 | 1.267 | 1.168 | 1.248 | 1.298 |
| 36 | 1.228 | 1.276 | 1.171 | 1.257 | 1.310 |
| 37 | 1.238 | 1.300 | 1.174 | 1.263 | 1.323 |
| 38 | 1.249 | 1.323 | 1.177 | 1.269 | 1.335 |
| 39 | 1.260 | 1.347 | 1.180 | 1.274 | 1.348 |
| 40 | 1.262 | 1.371 | 1.184 | 1.280 | 1.351 |
| 41 | 1.271 | 1.383 | 1.188 | 1.281 | 1.354 |
| 42 | 1.279 | 1.395 | 1.193 | 1.281 | 1.356 |
| 43 | 1.282 | 1.407 | 1.197 | 1.282 | 1.359 |
| 44 | 1.284 | 1.419 | 1.201 | 1.283 | 1.361 |
| 45 | 1.293 | 1.421 | 1.205 | 1.294 | 1.364 |
| 46 | 1.299 | 1.423 | 1.209 | 1.306 | 1.366 |
| 47 | 1.325 | 1.425 | 1.215 | 1.317 | 1.368 |
| 48 | 1.340 | 1.427 | 1.222 | 1.329 | 1.371 |
| 49 | 1.349 | 1.427 | 1.229 | 1.332 | 1.373 |
| 50 | 1.355 | 1.427 | 1.236 | 1.335 | 1.375 |
| 51 | 1.360 | 1.428 | 1.243 | 1.338 | 1.377 |
| 52 | 1.366 | 1.428 | 1.250 | 1.341 | 1.379 |
| 53 | 1.370 | 1.444 | 1.257 | 1.345 | 1.384 |
| 54 | 1.372 | 1.459 | 1.262 | 1.348 | 1.390 |
| 55 | 1.384 | 1.475 | 1.262 | 1.352 | 1.396 |
| 56 | 1.409 | 1.490 | 1.263 | 1.355 | 1.402 |
| 57 | 1.421 | 1.518 | 1.263 | 1.411 | 1.409 |
| 58 | 1.427 | 1.546 | 1.272 | 1.467 | 1.417 |
| 59 | 1.432 | 1.573 | 1.282 | 1.523 | 1.424 |
| 60 | 1.443 | 1.601 | 1.293 | 1.579 | 1.432 |
| 61 | 1.465 | 1.606 | 1.305 | 1.579 | 1.439 |
| 62 | 1.487 | 1.610 | 1.324 | 1.579 | 1.441 |
| 63 | 1.494 | 1.615 | 1.342 | 1.579 | 1.444 |
| 64 | 1.500 | 1.620 | 1.360 | 1.579 | 1.446 |
| 65 | 1.511 | 1.626 | 1.389 | 1.585 | 1.449 |
| 66 | 1.519 | 1.632 | 1.423 | 1.591 | 1.461 |

| | | | | | |
|---|---|---|---|---|---|
| 67 | 1.538 | 1.638 | 1.456 | 1.597 | 1.477 |
| 68 | 1.565 | 1.645 | 1.485 | 1.602 | 1.492 |
| 69 | 1.577 | 1.654 | 1.489 | 1.608 | 1.508 |
| 70 | 1.579 | 1.664 | 1.493 | 1.613 | 1.517 |
| 71 | 1.598 | 1.674 | 1.496 | 1.619 | 1.518 |
| 72 | 1.602 | 1.684 | 1.499 | 1.624 | 1.519 |
| 73 | 1.613 | 1.685 | 1.500 | 1.626 | 1.520 |
| 74 | 1.619 | 1.687 | 1.501 | 1.629 | 1.522 |
| 75 | 1.624 | 1.689 | 1.502 | 1.631 | 1.534 |
| 76 | 1.634 | 1.691 | 1.515 | 1.634 | 1.545 |
| 77 | 1.648 | 1.707 | 1.527 | 1.660 | 1.557 |
| 78 | 1.684 | 1.724 | 1.540 | 1.686 | 1.569 |
| 79 | 1.695 | 1.740 | 1.555 | 1.712 | 1.618 |
| 80 | 1.720 | 1.757 | 1.574 | 1.738 | 1.680 |
| 81 | 1.743 | 1.765 | 1.593 | 1.750 | 1.741 |
| 82 | 1.765 | 1.774 | 1.612 | 1.761 | 1.803 |
| 83 | 1.786 | 1.782 | 1.639 | 1.773 | 1.845 |
| 84 | 1.809 | 1.790 | 1.667 | 1.785 | 1.856 |
| 85 | 1.849 | 1.808 | 1.695 | 1.845 | 1.866 |
| 86 | 1.872 | 1.825 | 1.750 | 1.905 | 1.877 |
| 87 | 1.910 | 1.843 | 1.869 | 1.965 | 1.887 |
| 88 | 1.936 | 1.860 | 1.989 | 2.025 | 1.898 |
| 89 | 1.987 | 1.880 | 2.108 | 2.026 | 1.910 |
| 90 | 2.027 | 1.899 | 2.191 | 2.027 | 1.921 |
| 91 | 2.029 | 1.918 | 2.260 | 2.028 | 1.932 |
| 92 | 2.106 | 1.937 | 2.329 | 2.028 | 2.023 |
| 93 | 2.319 | 1.960 | 2.392 | 2.163 | 2.150 |
| 94 | 2.400 | 1.984 | 2.421 | 2.298 | 2.276 |
| 95 | 2.468 | 2.007 | 2.450 | 2.433 | 2.402 |
| 96 | 2.491 | 2.030 | 2.480 | 2.568 | 2.516 |
| 97 | 2.559 | 2.124 | 2.714 | 3.063 | 2.609 |
| 98 | 2.863 | 2.217 | 3.103 | 3.558 | 2.703 |
| 99 | 3.840 | 2.310 | 3.491 | 4.052 | 2.796 |
| 100 | 4.547 | 2.404 | 3.880 | 4.547 | 2.889 |

***Table S66:*** *Adaptive tapping – Adaptation index (deceleration trials) – Adaptive_adaptation_index_deceleration*

| | | | | | |
|---|---|---|---|---|---|
| **Task** | Adaptive tapping | | | | |
| **Outcome measure** | Adaptation index (deceleration trials) | | | | |
| **Variable name** | Adaptive_adaptation_index_deceleration | | | | |
| **Age and gender effects** | | | | | |
| Age regression (slope) | -0.005 | | | | |
| Age regression (p) | 0.032 | | | | |
| Age group (p) | 0.205 | | | | |
| Gender (p) | 0.301 | | | | |
| **Group** | All | Age 18 to 21 | Age 22 to 29 | Age 30 to 54 | Age 55 to 87 |
| N (N female) | 105 (72) | 26 (17) | 29 (19) | 26 (16) | 24 (20) |
| **Normality** | | | | | |
| Skewness | -0.36 | 0.54 | -2.18 | 0.92 | 0.34 |
| Excess kurtosis | 5.15 | 0.42 | 6.33 | 1.40 | -0.89 |
| **Scores** | | | | | |
| Mean | 1.263 | 1.390 | 1.186 | 1.347 | 1.128 |
| SD | 0.522 | 0.338 | 0.599 | 0.660 | 0.381 |
| Percentiles | | | | | |
| 0 | -1.169 | 0.826 | -1.169 | 0.309 | 0.536 |
| 1 | 0.310 | 0.832 | -0.748 | 0.316 | 0.563 |
| 2 | 0.337 | 0.838 | -0.326 | 0.323 | 0.591 |
| 3 | 0.353 | 0.844 | 0.096 | 0.330 | 0.619 |
| 4 | 0.475 | 0.850 | 0.352 | 0.338 | 0.647 |
| 5 | 0.545 | 0.870 | 0.388 | 0.399 | 0.663 |
| 6 | 0.592 | 0.889 | 0.423 | 0.460 | 0.672 |
| 7 | 0.634 | 0.909 | 0.458 | 0.521 | 0.681 |
| 8 | 0.659 | 0.928 | 0.502 | 0.582 | 0.689 |
| 9 | 0.675 | 0.955 | 0.548 | 0.602 | 0.696 |
| 10 | 0.696 | 0.983 | 0.593 | 0.623 | 0.696 |
| 11 | 0.699 | 1.010 | 0.634 | 0.643 | 0.697 |
| 12 | 0.718 | 1.037 | 0.666 | 0.664 | 0.697 |
| 13 | 0.764 | 1.046 | 0.697 | 0.700 | 0.698 |
| 14 | 0.799 | 1.056 | 0.728 | 0.735 | 0.698 |
| 15 | 0.812 | 1.066 | 0.773 | 0.771 | 0.699 |
| 16 | 0.822 | 1.076 | 0.823 | 0.806 | 0.699 |
| 17 | 0.842 | 1.096 | 0.874 | 0.836 | 0.700 |
| 18 | 0.889 | 1.117 | 0.921 | 0.865 | 0.713 |
| 19 | 0.914 | 1.137 | 0.951 | 0.895 | 0.733 |
| 20 | 0.923 | 1.157 | 0.981 | 0.924 | 0.754 |
| 21 | 0.928 | 1.164 | 1.011 | 0.944 | 0.774 |
| 22 | 0.928 | 1.171 | 1.030 | 0.963 | 0.791 |

| | | | | | |
|---|---|---|---|---|---|
| 23 | 0.989 | 1.177 | 1.040 | 0.982 | 0.797 |
| 24 | 1.001 | 1.184 | 1.050 | 1.001 | 0.803 |
| 25 | 1.009 | 1.187 | 1.060 | 1.006 | 0.809 |
| 26 | 1.020 | 1.190 | 1.066 | 1.010 | 0.815 |
| 27 | 1.021 | 1.192 | 1.072 | 1.015 | 0.834 |
| 28 | 1.026 | 1.195 | 1.078 | 1.020 | 0.855 |
| 29 | 1.040 | 1.207 | 1.090 | 1.020 | 0.875 |
| 30 | 1.063 | 1.219 | 1.110 | 1.020 | 0.896 |
| 31 | 1.076 | 1.232 | 1.130 | 1.021 | 0.908 |
| 32 | 1.077 | 1.244 | 1.150 | 1.021 | 0.913 |
| 33 | 1.079 | 1.250 | 1.160 | 1.035 | 0.919 |
| 34 | 1.086 | 1.255 | 1.168 | 1.049 | 0.924 |
| 35 | 1.104 | 1.261 | 1.177 | 1.062 | 0.932 |
| 36 | 1.124 | 1.266 | 1.185 | 1.076 | 0.947 |
| 37 | 1.135 | 1.269 | 1.197 | 1.081 | 0.962 |
| 38 | 1.143 | 1.271 | 1.208 | 1.086 | 0.977 |
| 39 | 1.150 | 1.274 | 1.219 | 1.090 | 0.992 |
| 40 | 1.155 | 1.276 | 1.226 | 1.095 | 0.997 |
| 41 | 1.160 | 1.284 | 1.233 | 1.106 | 1.001 |
| 42 | 1.176 | 1.293 | 1.239 | 1.117 | 1.004 |
| 43 | 1.184 | 1.301 | 1.245 | 1.128 | 1.008 |
| 44 | 1.192 | 1.310 | 1.248 | 1.140 | 1.017 |
| 45 | 1.216 | 1.313 | 1.251 | 1.145 | 1.033 |
| 46 | 1.241 | 1.316 | 1.255 | 1.151 | 1.049 |
| 47 | 1.245 | 1.319 | 1.258 | 1.156 | 1.064 |
| 48 | 1.253 | 1.322 | 1.261 | 1.162 | 1.079 |
| 49 | 1.256 | 1.344 | 1.264 | 1.185 | 1.088 |
| 50 | 1.256 | 1.367 | 1.268 | 1.209 | 1.098 |
| 51 | 1.266 | 1.389 | 1.276 | 1.232 | 1.107 |
| 52 | 1.268 | 1.411 | 1.284 | 1.256 | 1.117 |
| 53 | 1.277 | 1.416 | 1.292 | 1.277 | 1.121 |
| 54 | 1.287 | 1.421 | 1.300 | 1.299 | 1.123 |
| 55 | 1.300 | 1.426 | 1.306 | 1.320 | 1.126 |
| 56 | 1.312 | 1.430 | 1.313 | 1.341 | 1.129 |
| 57 | 1.321 | 1.449 | 1.319 | 1.346 | 1.132 |
| 58 | 1.328 | 1.468 | 1.332 | 1.351 | 1.135 |
| 59 | 1.348 | 1.487 | 1.346 | 1.356 | 1.139 |
| 60 | 1.364 | 1.506 | 1.359 | 1.361 | 1.142 |
| 61 | 1.383 | 1.510 | 1.371 | 1.405 | 1.149 |
| 62 | 1.405 | 1.513 | 1.380 | 1.448 | 1.174 |
| 63 | 1.421 | 1.516 | 1.389 | 1.492 | 1.199 |
| 64 | 1.432 | 1.520 | 1.397 | 1.536 | 1.224 |
| 65 | 1.457 | 1.523 | 1.407 | 1.546 | 1.249 |
| 66 | 1.484 | 1.527 | 1.416 | 1.555 | 1.260 |

| | | | | | |
|---|---|---|---|---|---|
| 67 | 1.490 | 1.530 | 1.426 | 1.565 | 1.267 |
| 68 | 1.502 | 1.533 | 1.435 | 1.575 | 1.274 |
| 69 | 1.517 | 1.538 | 1.446 | 1.600 | 1.281 |
| 70 | 1.528 | 1.542 | 1.457 | 1.625 | 1.305 |
| 71 | 1.533 | 1.546 | 1.468 | 1.650 | 1.352 |
| 72 | 1.536 | 1.550 | 1.475 | 1.675 | 1.400 |
| 73 | 1.549 | 1.554 | 1.480 | 1.678 | 1.447 |
| 74 | 1.563 | 1.558 | 1.485 | 1.680 | 1.491 |
| 75 | 1.565 | 1.561 | 1.490 | 1.682 | 1.500 |
| 76 | 1.566 | 1.565 | 1.512 | 1.685 | 1.509 |
| 77 | 1.567 | 1.567 | 1.533 | 1.690 | 1.518 |
| 78 | 1.570 | 1.569 | 1.555 | 1.696 | 1.528 |
| 79 | 1.574 | 1.571 | 1.569 | 1.701 | 1.536 |
| 80 | 1.576 | 1.573 | 1.573 | 1.707 | 1.544 |
| 81 | 1.586 | 1.579 | 1.577 | 1.763 | 1.551 |
| 82 | 1.619 | 1.585 | 1.581 | 1.818 | 1.559 |
| 83 | 1.676 | 1.591 | 1.621 | 1.874 | 1.564 |
| 84 | 1.680 | 1.597 | 1.666 | 1.930 | 1.565 |
| 85 | 1.686 | 1.620 | 1.712 | 1.976 | 1.565 |
| 86 | 1.697 | 1.643 | 1.747 | 2.022 | 1.566 |
| 87 | 1.725 | 1.666 | 1.759 | 2.068 | 1.566 |
| 88 | 1.766 | 1.689 | 1.771 | 2.114 | 1.567 |
| 89 | 1.795 | 1.734 | 1.782 | 2.117 | 1.568 |
| 90 | 1.806 | 1.779 | 1.789 | 2.120 | 1.569 |
| 91 | 1.848 | 1.824 | 1.794 | 2.122 | 1.569 |
| 92 | 1.897 | 1.869 | 1.798 | 2.125 | 1.587 |
| 93 | 1.924 | 1.886 | 1.803 | 2.152 | 1.612 |
| 94 | 1.934 | 1.902 | 1.805 | 2.180 | 1.636 |
| 95 | 1.936 | 1.918 | 1.806 | 2.208 | 1.661 |
| 96 | 2.086 | 1.935 | 1.808 | 2.236 | 1.698 |
| 97 | 2.123 | 2.023 | 1.825 | 2.510 | 1.757 |
| 98 | 2.227 | 2.111 | 1.853 | 2.785 | 1.817 |
| 99 | 2.285 | 2.199 | 1.882 | 3.060 | 1.877 |
| 100 | 3.334 | 2.287 | 1.910 | 3.334 | 1.936 |

***Table S67:*** *BTI – Beat Tracking Index – BTI*

| | | | | | | |
|---|---|---|---|---|---|---|
| **Task** | BTI | | | | | |
| **Outcome measure** | Beat Tracking Index | | | | | |
| **Variable name** | BTI | | | | | |
| **Age and gender effects** | | | | | | |
| Age regression (slope) | | 0.001 | | | | |
| Age regression (p) | | 0.896 | | | | |
| Age group (p) | | 0.682 | | | | |
| Gender (p) | | 0.556 | | | | |
| **Group** | | All | Age 18 to 21 | Age 22 to 29 | Age 30 to 54 | Age 55 to 87 |
| N (N female) | | 102 (70) | 24 (17) | 28 (18) | 25 (15) | 25 (20) |
| **Normality** | | | | | | |
| Skewness | | -0.50 | -0.75 | -0.27 | -0.55 | 0.07 |
| Excess kurtosis | | 0.06 | 0.80 | -0.72 | -0.14 | -0.26 |
| **Scores** | | | | | | |
| Mean | | -0.008 | 0.038 | -0.164 | 0.089 | 0.025 |
| SD | | 0.806 | 0.772 | 0.990 | 0.817 | 0.592 |
| Percentiles | | | | | | |
| | 0 | -2.180 | -2.136 | -2.180 | -1.791 | -1.148 |
| | 1 | -2.135 | -1.870 | -2.159 | -1.724 | -1.082 |
| | 2 | -2.096 | -1.605 | -2.138 | -1.656 | -1.016 |
| | 3 | -1.786 | -1.339 | -2.117 | -1.589 | -0.950 |
| | 4 | -1.611 | -1.074 | -2.063 | -1.522 | -0.884 |
| | 5 | -1.504 | -0.950 | -1.932 | -1.400 | -0.861 |
| | 6 | -1.362 | -0.902 | -1.800 | -1.267 | -0.847 |
| | 7 | -1.136 | -0.854 | -1.668 | -1.134 | -0.833 |
| | 8 | -0.980 | -0.807 | -1.576 | -1.002 | -0.820 |
| | 9 | -0.966 | -0.772 | -1.512 | -0.947 | -0.792 |
| | 10 | -0.951 | -0.770 | -1.447 | -0.932 | -0.757 |
| | 11 | -0.892 | -0.767 | -1.383 | -0.917 | -0.722 |
| | 12 | -0.866 | -0.764 | -1.277 | -0.902 | -0.687 |
| | 13 | -0.813 | -0.762 | -1.167 | -0.861 | -0.664 |
| | 14 | -0.798 | -0.730 | -1.056 | -0.795 | -0.651 |
| | 15 | -0.792 | -0.698 | -0.958 | -0.728 | -0.638 |
| | 16 | -0.775 | -0.665 | -0.913 | -0.661 | -0.625 |
| | 17 | -0.773 | -0.633 | -0.868 | -0.616 | -0.597 |
| | 18 | -0.769 | -0.610 | -0.822 | -0.614 | -0.537 |
| | 19 | -0.744 | -0.594 | -0.798 | -0.612 | -0.478 |
| | 20 | -0.660 | -0.578 | -0.797 | -0.610 | -0.419 |
| | 21 | -0.619 | -0.562 | -0.796 | -0.602 | -0.365 |
| | 22 | -0.617 | -0.536 | -0.795 | -0.563 | -0.335 |
| | 23 | -0.615 | -0.484 | -0.791 | -0.524 | -0.305 |
| | 24 | -0.607 | -0.432 | -0.786 | -0.485 | -0.275 |

| | | | | | |
|---|---|---|---|---|---|
| 25 | -0.590 | -0.380 | -0.781 | -0.446 | -0.246 |
| 26 | -0.544 | -0.327 | -0.776 | -0.369 | -0.244 |
| 27 | -0.508 | -0.317 | -0.774 | -0.291 | -0.243 |
| 28 | -0.450 | -0.310 | -0.773 | -0.214 | -0.241 |
| 29 | -0.424 | -0.304 | -0.772 | -0.136 | -0.240 |
| 30 | -0.367 | -0.297 | -0.754 | -0.104 | -0.214 |
| 31 | -0.349 | -0.276 | -0.709 | -0.080 | -0.184 |
| 32 | -0.314 | -0.243 | -0.663 | -0.057 | -0.153 |
| 33 | -0.278 | -0.210 | -0.618 | -0.034 | -0.123 |
| 34 | -0.244 | -0.177 | -0.590 | -0.009 | -0.112 |
| 35 | -0.230 | -0.145 | -0.570 | 0.017 | -0.112 |
| 36 | -0.190 | -0.117 | -0.549 | 0.043 | -0.112 |
| 37 | -0.141 | -0.089 | -0.529 | 0.068 | -0.112 |
| 38 | -0.119 | -0.060 | -0.509 | 0.082 | -0.108 |
| 39 | -0.112 | -0.032 | -0.488 | 0.085 | -0.102 |
| 40 | -0.108 | -0.017 | -0.467 | 0.088 | -0.096 |
| 41 | -0.097 | -0.004 | -0.445 | 0.090 | -0.089 |
| 42 | -0.087 | 0.009 | -0.421 | 0.100 | -0.083 |
| 43 | -0.076 | 0.022 | -0.396 | 0.123 | -0.078 |
| 44 | -0.048 | 0.039 | -0.372 | 0.145 | -0.073 |
| 45 | -0.027 | 0.061 | -0.339 | 0.168 | -0.068 |
| 46 | -0.021 | 0.083 | -0.298 | 0.187 | -0.061 |
| 47 | -0.009 | 0.104 | -0.258 | 0.188 | -0.050 |
| 48 | 0.012 | 0.123 | -0.218 | 0.188 | -0.039 |
| 49 | 0.054 | 0.127 | -0.187 | 0.189 | -0.027 |
| 50 | 0.087 | 0.131 | -0.158 | 0.190 | -0.016 |
| 51 | 0.095 | 0.135 | -0.128 | 0.206 | -0.013 |
| 52 | 0.097 | 0.139 | -0.103 | 0.222 | -0.009 |
| 53 | 0.108 | 0.143 | -0.099 | 0.239 | -0.006 |
| 54 | 0.120 | 0.147 | -0.095 | 0.255 | -0.003 |
| 55 | 0.126 | 0.151 | -0.091 | 0.278 | 0.022 |
| 56 | 0.135 | 0.155 | -0.066 | 0.303 | 0.051 |
| 57 | 0.149 | 0.169 | -0.016 | 0.328 | 0.080 |
| 58 | 0.157 | 0.195 | 0.034 | 0.353 | 0.108 |
| 59 | 0.175 | 0.220 | 0.084 | 0.366 | 0.124 |
| 60 | 0.189 | 0.245 | 0.097 | 0.374 | 0.133 |
| 61 | 0.197 | 0.268 | 0.097 | 0.381 | 0.143 |
| 62 | 0.237 | 0.279 | 0.097 | 0.389 | 0.152 |
| 63 | 0.263 | 0.291 | 0.098 | 0.396 | 0.162 |
| 64 | 0.277 | 0.302 | 0.106 | 0.401 | 0.173 |
| 65 | 0.304 | 0.313 | 0.115 | 0.406 | 0.184 |
| 66 | 0.346 | 0.333 | 0.123 | 0.412 | 0.195 |
| 67 | 0.382 | 0.355 | 0.177 | 0.417 | 0.208 |
| 68 | 0.407 | 0.378 | 0.322 | 0.422 | 0.228 |

| | | | | | |
|---|---|---|---|---|---|
| 69 | 0.415 | 0.400 | 0.467 | 0.427 | 0.247 |
| 70 | 0.430 | 0.422 | 0.611 | 0.432 | 0.266 |
| 71 | 0.467 | 0.443 | 0.674 | 0.442 | 0.290 |
| 72 | 0.498 | 0.465 | 0.687 | 0.474 | 0.337 |
| 73 | 0.554 | 0.486 | 0.701 | 0.507 | 0.385 |
| 74 | 0.576 | 0.507 | 0.714 | 0.539 | 0.433 |
| 75 | 0.591 | 0.523 | 0.730 | 0.572 | 0.480 |
| 76 | 0.600 | 0.540 | 0.746 | 0.584 | 0.508 |
| 77 | 0.618 | 0.557 | 0.762 | 0.596 | 0.535 |
| 78 | 0.656 | 0.573 | 0.779 | 0.608 | 0.563 |
| 79 | 0.674 | 0.610 | 0.799 | 0.620 | 0.591 |
| 80 | 0.708 | 0.654 | 0.818 | 0.660 | 0.597 |
| 81 | 0.717 | 0.697 | 0.837 | 0.705 | 0.598 |
| 82 | 0.759 | 0.741 | 0.875 | 0.751 | 0.600 |
| 83 | 0.774 | 0.780 | 0.929 | 0.796 | 0.602 |
| 84 | 0.777 | 0.811 | 0.983 | 0.830 | 0.614 |
| 85 | 0.806 | 0.842 | 1.038 | 0.858 | 0.632 |
| 86 | 0.842 | 0.873 | 1.061 | 0.887 | 0.649 |
| 87 | 0.895 | 0.904 | 1.078 | 0.915 | 0.667 |
| 88 | 0.926 | 0.933 | 1.094 | 0.949 | 0.681 |
| 89 | 1.020 | 0.963 | 1.109 | 0.989 | 0.691 |
| 90 | 1.046 | 0.992 | 1.112 | 1.028 | 0.701 |
| 91 | 1.089 | 1.022 | 1.114 | 1.067 | 0.711 |
| 92 | 1.099 | 1.042 | 1.117 | 1.113 | 0.722 |
| 93 | 1.108 | 1.058 | 1.133 | 1.170 | 0.737 |
| 94 | 1.118 | 1.073 | 1.167 | 1.227 | 0.751 |
| 95 | 1.203 | 1.089 | 1.202 | 1.285 | 0.766 |
| 96 | 1.246 | 1.108 | 1.237 | 1.336 | 0.802 |
| 97 | 1.289 | 1.133 | 1.255 | 1.357 | 0.944 |
| 98 | 1.331 | 1.158 | 1.267 | 1.378 | 1.087 |
| 99 | 1.372 | 1.182 | 1.279 | 1.400 | 1.230 |
| 100 | 1.421 | 1.207 | 1.291 | 1.421 | 1.373 |
