## Supplementary materials for "Mobile version of the Battery for the Assessment of Auditory Sensorimotor and Timing Abilities (BAASTA): Implementation and adult norms"

#### Comparison of the results obtained with the BAASTA norm sample (Test) with the original version of BAASTA (*Dalla Bella et al. (2017)*

| | | **BAASTA Norms 2023** | | **Dalla Bella et al. (2017)** | | |
|---|---|---|---|---|---|---|
| **Task** | **Variable** | **N** | **Mean (SD)** | **N** | **Mean (SD)** | **p** |
| Beat Alignment Test | Sensitivity index (d') for all trials | 108 | 2.76 (1.03) | 20 | 2.67 (1.2) | 1.000 |
| Beat Alignment Test | Sensitivity index (d') for fast tempo trials | 108 | 2.22 (0.87) | 20 | 2.21 (1.14) | 1.000 |
| Beat Alignment Test | Sensitivity index (d') for medium tempo trials | 108 | 2.43 (0.87) | 20 | 2.42 (0.88) | 1.000 |
| Beat Alignment Test | Sensitivity index (d') for slow tempo trials | 108 | 2.65 (0.91) | 20 | 2.42 (0.96) | 1.000 |
| Anisochrony detection with music | Threshold (% of IOI) [a] | 98 | 12 (7.1) | 19 | 8 (3.8) | 0.807 |
| Anisochrony detection with tones | Threshold (% of IOI) [a] | 100 | 12.4 (5.1) | 20 | 10.8 (4) | 1.000 |
| Duration discrimination | Threshold (% of IOI) [a] | 101 | 29.1 (12.7) | 18 | 23 (6.6) | 1.000 |
| Adaptive tapping | Adaptation index (acceleration trials) | 105 | 1.442 (0.597) | 20 | 1.28 (0.453) | 1.000 |
| Adaptive tapping | Adaptation index (deceleration trials) | 105 | 1.263 (0.522) | 20 | 1.413 (0.57) | 1.000 |
| Adaptive tapping | Sensitivity index (d') perceiving a tempo acceleration (IOI - 30ms) | 105 | 1.69 (0.88) | 20 | 2.17 (0.65) | 0.986 |
| Adaptive tapping | Sensitivity index (d') of perceiving tempo acceleration (IOI - 75ms) | 105 | 3.79 (0.54) | 20 | 3.75 (0.49) | 1.000 |
| Adaptive tapping | Sensitivity index (d') of perceiving tempo deceleration (IOI + 30ms) | 105 | 1.84 (0.98) | 20 | 2.37 (0.68) | 0.122 |
| Adaptive tapping | Sensitivity index (d') of perceiving tempo deceleration (IOI + 75ms) | 105 | 3.63 (0.74) | 20 | 3.59 (0.66) | 1.000 |
| Paced tapping with tones | Accuracy (vector direction %), fast tempo | 97 | -7.1 (5.9) | 19 | -4.7 (6.4) | 1.000 |
| Paced tapping with tones | Accuracy (vector direction %), medium tempo | 106 | -6.5 (5.6) | 18 | -4.3 (5.2) | 1.000 |
| Paced tapping with tones | Accuracy (vector direction %), slow tempo | 104 | -5.3 (4.7) | 20 | -5 (6.4) | 1.000 |

| | | | | | | |
|---|---|---|---|---|---|---|
| Paced tapping with music | Accuracy (vector direction %), music 1 | 99 | 0.3 (5.4) | 20 | 2.5 (5.4) | 1.000 |
| Paced tapping with music | Accuracy (vector direction %), music 2 | 90 | -3.4 (6.6) | 20 | -1.1 (8.9) | 1.000 |
| Synchronization-continuation | Variability (CV of ITI), fast tempo | 108 | 0.048 (0.015) | 20 | 0.043 (0.015) | 1.000 |
| Synchronization-continuation | Rate (ITI in ms), fast tempo | 108 | 448.8 (21.6) | 20 | 445.4 (16.4) | 1.000 |
| Synchronization-continuation | Variability (CV of ITI), medium tempo | 108 | 0.046 (0.014) | 20 | 0.037 (0.01) | 0.119 |
| Synchronization-continuation | Rate (ITI in ms), medium tempo | 108 | 596.5 (26.1) | 20 | 592.8 (16.5) | 1.000 |
| Synchronization-continuation | Variability (CV of ITI), slow tempo | 108 | 0.048 (0.014) | 20 | 0.039 (0.013) | 0.063 |
| Synchronization-continuation | Rate (ITI in ms), slow tempo | 108 | 749.3 (40.2) | 20 | 759.2 (32) | 1.000 |
| Unpaced tapping | Variability (CV of ITI) of fast tapping | 107 | 0.061 (0.029) | 20 | 0.051 (0.021) | 1.000 |
| Unpaced tapping | Rate (ITI in ms) of fast tapping | 107 | 333.9 (134.5) | 20 | 287.7 (117.5) | 1.000 |
| Unpaced tapping | Variability (CV of ITI) of slow tapping | 103 | 0.064 (0.061) | 20 | 0.048 (0.014) | 1.000 |
| Unpaced tapping | Rate (ITI in ms) of slow tapping | 103 | 1230.1 (465.8) | 20 | 1342.5 (371.3) | 1.000 |
| Unpaced tapping | Variability (CV of ITI) of spontaneous tapping, initial trial | 106 | 0.057 (0.039) | 20 | 0.044 (0.015) | 0.307 |
| Unpaced tapping | Rate (ITI in ms) of spontaneous tapping, initial trial | 106 | 685.3 (243) | 20 | 611.5 (153.4) | 1.000 |
| Paced tapping with tones | Consistency (logit of vector length), fast tempo | 107 | 2.77 (0.86) | 20 | 3.07 (0.81) | 1.000 |
| Paced tapping with tones | Consistency (logit of vector length), medium tempo | 108 | 2.97 (0.78) | 20 | 3.1 (1) | 1.000 |
| Paced tapping with tones | Consistency (logit of vector length), slow tempo | 107 | 2.94 (0.95) | 20 | 2.98 (0.89) | 1.000 |
| Paced tapping with music | Consistency (logit of vector length), music 1 | 106 | 2.78 (1.52) | 20 | 2.79 (1.42) | 1.000 |
| Paced tapping with music | Consistency (logit of vector length), music 2 | 102 | 2.35 (1.71) | 20 | 2.49 (1.64) | 1.000 |

*p* values were calculated using Wilcoxon-Mann-Whitney tests and corrected using the Bonferroni method.

[a] In Dalla Bella et al. (2017), perceptual thresholds were estimated using a maximum likelihood procedure, whereas in the present version of BAASTA a 2-down/1-up staircase procedure is used.

### QUESTIONNAIRES (formal / non-formal musical training)

**ENGLISH VERSION**

**General Survey. Any information that can be linked to your identity will be used for administrative purpose. These informations will not be transcribed in the datafiles. This form will remain confidential and away from your experimental datas.**

Name: _________________________________ Age: _____ Gender: ______________

Phone: _________________________________ Email: ___________________________________

Do you wish to be contacted for future studies? (Circle) YES / NO

Are you right or left handed (When using a pen) ? Right-Handed / Left-Handed / Ambidextrous (Both)

What is your main language? ______________________

Which other language do you speak fluently? _________________________________

If you currently have any auditory issues, please list them below (e.g.: otitis, hearing impairment, etc.): ____________________________

**I. <u>Formal musical training</u>**:

1. Have you ever taken any musical lesson? (All type of lessons included, such as elementary school music class)?

   (Circle) YES / NO

   * If **YES**, please fill section #2 and #3, if **NOT**, please fill section #4

2. Please, write which i*nstrument you've learned to play or what kind of singing training you have been following*, taking a different line for each instrument (or voice). "Individual lessons" refers to private lessons. "Group lessons" refers to group classes or schools lessons.

| Instrument/Voice | Individual lessons (How long?) | How old were you? | Group lessons (How long?) | How old were you? |
|---|---|---|---|---|
| 1) | | | | |
| 2) | | | | |
| 3) | | | | |
| 4) | | | | |

3. Please, write down any training you may have in *musical theory*:

| Type (for example, composition) | Individual lessons (How long?) | How old were you? | Group lessons (How long?) | How old were you? |
|---|---|---|---|---|
| 1) | | | | |
| 2) | | | | |

**II**. **<u>Non-formal musical training & participation to musical activities:</u>**

4. Do you have any self-educated musical instrument training, on your own or with a group (with any formal lesson with a teacher)?

| Instrument/Voice | Individual lessons (How long?) | How old were you? | Group lessons (How long?) | How old were you? |
|---|---|---|---|---|
| 1) | | | | |
| 2) | | | | |
| 3) | | | | |
| 4) | | | | |

5. If you once practiced music, are you still practicing (for example, as a spare-time, formal lessons, performances, etc…)?

(Circle) YES / NO

**If NO:** When did you stop? ______________________

**Si OUI:** Spare-time ( Write down **how many hours per month** and the **instrument**): ______________________

Formal lessons ( Write down **how many hours per month** and the **instrument**): ______________________

Non-professional performances ( Write down **how many hours per month** and the **instrument**): ______________________

Professional performances (paid)
( Write down **how many hours per month** and the **instrument**): ______________________

6. Do you consider yourself as a professional musician? (Circle) YES / NO

7. Do you listen to music in a regular way? (Circle) YES / NO

If YES, how many hours per day? ______________________

If YES, what type of music do you listen to (for example, classic, rock)?

______________________

8. If you hear a musical note, can you *name* which note it is without using any other note as reference? YES / NO

**FRENCH VERSION**

**Questionnaire général. Les informations permettant de vous identifier servent à des fins administratives seulement. Ces informations ne seront pas conservées dans les fichiers de données. Ce formulaire restera confidentiel et tenu à l'écart de vos données expérimentales.**

Nom: _________________________________ Age: _____ Genre: ______________

Téléphone: _________________________________ Courriel: ___________________________________

Souhaitez-vous être contacté(e) pour d'autres études dans le futur? (Encerclez) OUI / NON

Etes-vous gaucher(ère) ou droitier(ère) (Lorsque vous utilisez un crayon) ? (Encerclez) Droitier(ère) / Gaucher(ère) / Ambidextre

Quelle est votre langue maternelle? ______________________

Quelle(s) autre(s) langue(s) parlez-vous couramment? _________________________________

Si vous présentez actuellement des problèmes d'audition, veuillez en faire la liste (par exemple, otite, perte auditive, etc.): ____________________________

**I. <u>Formation musicale formelle</u>:**

1. Avez-vous déjà pris des leçons de musique (tous les types de leçons comptent, par exemple, des cours en groupe à l'école)?

   (Encerclez) OUI / NON

   * Si **OUI**, veuillez remplir la section #2 et #3; Si **NON**, passez à la section #4

2. Veuillez indiquer quel *instrument ou entraînement de la voix* vous avez suivi, en prenant une ligne différente pour chaque instrument (ou voix). "Leçons individuelles" réfèrent à des cours particulier, "Leçons en groupe" réfèrent à des cours de groupe ou à l'école.

| Instrument/Voix | Leçons individuelles (nombre d'années) | À quel(s) âge(s)? | Leçons en groupe (nombre d'années) | À quel(s) âge(s)? |
|---|---|---|---|---|
| 1) | | | | |
| 2) | | | | |
| 3) | | | | |
| 4) | | | | |

3. Veuillez indiquer votre formation en *théorie musicale*, si vous en avez eue:

| Type (par exemple, composition) | Leçons individuelles (nombre d'années) | À quel(s) âge(s)? | Leçons en groupe (nombre d'années) | À quel(s) âge(s)? |
|---|---|---|---|---|
| 1) | | | | |
| 2) | | | | |

**II**. **Formation musicale informelle et participation actuelle à des activités musicales**:

4. Avez-vous appris à jouer d'un instrument de façon autodidacte, seul ou en groupe (c'est-à-dire sans leçons formelles avec un professeur)?

| Instrument/Voix | Leçons individuelles (nombre d'années) | À quel(s) âge(s)? | Leçons en groupe (nombre d'années) | À quel(s) âge(s)? |
|---|---|---|---|---|
| 1) | | | | |
| 2) | | | | |
| 3) | | | | |
| 4) | | | | |

5. Si vous avez un jour pratiqué la musique, est-ce toujours le cas (par exemple, en loisir, leçons formelles, représentations)?

(Encerclez) OUI / NON

**Si NON:** Depuis quand avez-vous arrêté? ____________________

**Si OUI:** En loisir (indiquez combien d'**heures par mois** et l'**instrument**): ____________________

Leçons formelles (indiquez combien d'**heures par mois** et l'**instrument**): ____________________

Représentations amateures (indiquez combien d'**heures par mois** et l'**instrument**): ____________________

Représentations professionnelles (rémunérées) (indiquez combien d'**heures par mois** et l'**instrument**): ____________________

6. Vous considérez-vous comme un(e) musicien(ne) professionnel(le)? (Encerclez): OUI / NON

7. Écoutez-vous de la musique régulièrement? (Encerclez): OUI / NON

Si OUI, combien d'heures par jour? ____________________

Si OUI, quel type de musique (par exemple, classique, rock)?

____________________

8. Si vous entendez une note de musique, pouvez-vous donner le *nom* de la note sans devoir recourir à une autre note comme référence? OUI / NON
